# Phosphorylation-dependent Remodeling of the CLOCK–BMAL1–nucleosome Complex

**DOI:** 10.64898/2026.06.30.735537

**Authors:** Dima Amairy, Hanife Pekel, Şeref Gül, Ozge Sensoy

**Affiliations:** Graduate School of Engineering and Natural Sciences, Istanbul Medipol University, Istanbul, Turkey; Department of Pharmacy Services, Vocational School of Health Services, Istanbul Medipol University, Istanbul, Türkiye; Regenerative and Restorative Medicine Research Center (REMER), Research Institute for Health Sciences and Technologies (SABITA), Istanbul Medipol University, Istanbul, Türkiye; Institute of Life Sciences and Biotechnology, Bezmialem Vakif University, Istanbul, Türkiye

**Keywords:** circadian clock, CLOCK/BMAL1, phosphorylation, nucleosome, coarse-grained molecular dynamics simulation, allostery, transcription factors

## Abstract

Phosphorylation of the CLOCK/BMAL1 complex is a reversible post-translational modification that plays a central role in regulating circadian oscillations; however, its underlying mechanistic basis remains poorly understood. Although biochemical studies have shown that phosphorylation modulates CLOCK/BMAL1 binding to DNA, yet it remains unclear whether these effects are confined to local perturbation or also propagate through allosteric effects. Moreover, the influence of phosphorylation on histone dynamics and transcription factor-nucleosome interactions has not been systematically investigated.

Here, we address these questions using atomistic trajectories obtained through backmapping of coarse-grained molecular dynamics simulations based on the recently resolved cryo-EM structure of the CLOCK/BMAL1 and nucleosome complex. We investigated three experimentally identified phosphorylation states: CLOCK bHLH^S38/42^, BMAL1 bHLH^S78^, and simultaneous phosphorylation of both proteins. Our results demonstrate that phosphorylation regulates CLOCK and BMAL1 asymmetrically. Whereas phosphorylation weakens the interaction of the modified bHLH domain with the E-box, BMAL1 phosphorylation simultaneously enhances DNA engagement by the CLOCK bHLH domain, an effect that persists in the doubly phosphorylated complex. Steered pulling simulations showed that phosphorylation of the CLOCK bHLH domain results in mechanical stability comparable to that of the unphosphorylated system. In contrast, BMAL1 phosphorylation appears to reduce the mechanical stability of the bHLH–DNA interaction.

Beyond modulating DNA binding, phosphorylation remodels protein–histone interactions by altering contacts between the CLOCK PAS-B domain and histone H3 and between the BMAL1 PAS-A domain and the H2A/H2B acidic patch, while simultaneously rewiring residue-correlation and allosteric communication networks throughout the heterodimer. Importantly, phosphorylation increases the separation between the CLOCK HI loop and the H3 α1–L1 elbow, supporting a structural model in which phosphorylation provides a more permissive environment for CRY1 recruitment to the chromatin-bound CLOCK–BMAL1 complex, thereby facilitating displacement-type transcriptional repression.

Collectively, our findings reveal that phosphorylation regulates the CLOCK–BMAL1 complex through coordinated remodeling of DNA binding, nucleosome interactions, and long-range allosteric communication, providing a mechanistic framework for circadian transcriptional repression and a foundation for the rational design of therapeutics targeting the molecular circadian clock.

## Introduction

Circadian rhythms in mammals orchestrate a wide range of physiological functions, including sleep-wake cycle, hormone secretion, and immune responses, all of which exhibit rhythmic fluctuations across the 24-hours.^1–4^ These rhythms are essential for maintaining whole-body homeostasis. The mammalian circadian clock is primarily governed by a transcriptional-translational feedback loop, in which BMAL1 and CLOCK form the positive arm. BMAL1 and CLOCK forms a heterodimer through their PER–ARNT–SIM (PAS; PAS-A and PAS-B) domains, whereas their basic helix–loop–helix (bHLH) domains mediate binding to E-box sequences in the regulatory regions of target genes, thereby initiating transcription (Figure 1).^5–7^ Cryptochromes (CRY1/CRY2) and Periods (PER1/PER2/PER3), whose expression is activated by the CLOCK-BMAL1 complex, form the negative arm of the loop by inhibiting the CLOCK-BMAL1 driven transcription.^8,9^ CRY proteins, the primary repressor, can directly bind to CLOCK and BMAL1 and inhibit their transactivation activity while the complex is bound to E-box elements.^10^ In addition, through a CRY-dependent mechanism, PER facilitates the dissociation of the CLOCK–BMAL1 complex from the E-box DNA.^11^

**Figure 1.**
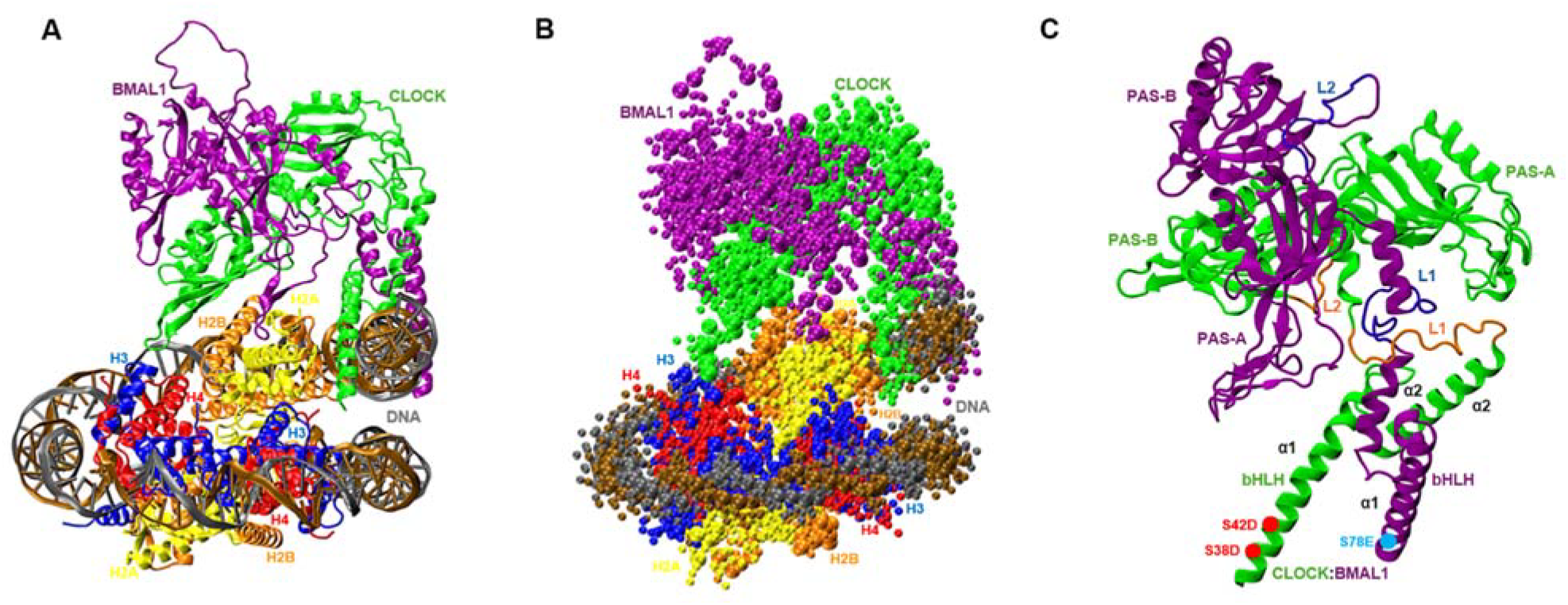
Three-dimensional structure of CLOCK-BMAL1/nucleosome complex (PDB ID:8OSK)^12^ shown in **A.**atomistic and **B**. coarse-grained representation. **C**. bHLH, PAS-A/B domains, and linkers that connect these domains are shown on the 3D structure of CLOCK and BMAL1. The Cα atoms of the phosphorylated residues are shown by red and blue circles.

Post-translational modifications (PTMs) play critical roles in regulating the repression and reactivation phases of the circadian feedback loop. For instance, phosphorylation and ubiquitination of CRY and PER proteins regulate their stability and degradation, thereby contributing to the reactivation of CLOCK–BMAL1–mediated transcription.^13–15^ During the repression phase, the CRY–PER complex recruits casein kinases to the nucleus, leading to phosphorylation of CLOCK and reducing the affinity of CLOCK–BMAL1 for E-box sequences.^16,17^ Mass spectrometry-based analyses of mouse liver identified Ser38 and Ser42 of CLOCK as phosphorylation sites during PER mediated repression.^18^ Substitution of these residues with glutamate to mimic constitutive phosphorylation attenuated CLOCK–BMAL1 transactivation activity in subsequent transactivation assays.^19^ The functional consequences of phosphorylation at these CLOCK residues and at BMAL1-Ser78, known to impair E-box binding when mutated to glutamate,^6^ were further characterized using rescue complementation assays and mathematical modeling.^16^ Biochemical and quantitative mass spectrometry analyses using biotinylated E-box sequences further demonstrated that these substitutions significantly reduced CLOCK– BMAL1 binding to DNA. Moreover, simultaneous phospho-mimetic substitutions at all three serine residues (CLOCK^S38D/S42D^ and BMAL1^S78E^) abolished binding to the E-box.^16^ Although these findings establish an important role for phosphorylation in regulating CLOCK–BMAL1 activity, they were largely obtained using isolated DNA or cellular complementation systems. Because E-box elements are embedded within nucleosomal DNA and CLOCK–BMAL1 engages chromatinized target sites in vivo,^20^ it remains unclear how phosphorylation regulates CLOCK–BMAL1 within the structural and dynamic constraints imposed by the nucleosome. Whether phosphorylation-induced perturbations at the bHLH domains propagate to histones and distal regulatory domains remains unknown.

A recently resolved cryo-EM structure offers detailed insights into the recognition of nucleosome-bound E-box elements by two evolutionarily distinct bHLH transcription factor complexes, CLOCK–BMAL1 and MYC– MAX.^12^ The nature of CLOCK–BMAL1’s interactions with histones depends on both the protein composition and the specific location of the DNA motifs, and these interactions can be modulated by nearby histone modifications. CLOCK-BMAL1 interacts directly with histone surfaces through its PAS domains, particularly targeting the C-terminal helix of H2B and the junction of the α1 helix in H3. Notably, the HI loop of CLOCK, a region essential for binding to CRY1 and CRY2,^21,22^ lies at the histone interaction interface. This spatial relationship suggests that nucleosome engagement may influence the accessibility of the CLOCK surface recognized by CRYs and provides a potential structural connection between chromatin binding and CRY-mediated repression.

Coarse-grained (CG) models simplify molecular systems by grouping atoms into beads, thereby reducing the degree of freedom in the system. This reduction also eliminates high-frequency vibrational modes, enabling the use of longer timesteps and resulting in smoother energy landscapes. As a result, CG molecular dynamics (MD) simulations are well-suited for investigating the dynamics of large biological systems such as nuclear pore complexes and nucleosomes.^23^ Advances in CG modeling have further enabled detailed characterization of DNA and nucleosome dynamics over timescales that remain challenging for conventional all-atom simulations.^24^ Here, we used CG-MD simulations based on the nucleosome-bound CLOCK–BMAL1 structure to investigate how phosphorylation at CLOCK^S38/S42^ and BMAL1^S78^ remodels the dynamics of the chromatin-bound complex. We characterized phosphorylation-dependent changes in E-box engagement, histone interactions, and allosteric communication within the CLOCK–BMAL1–nucleosome assembly. We further examined how these changes influence the structural accessibility of the CLOCK HI loop and its potential interaction with CRY1. By investigating phosphorylation within an explicit nucleosomal context, our study provides a mechanistic framework for understanding how localized PTMs propagate through the CLOCK– BMAL1 complex to reshape its interactions with DNA and the nucleosome, while providing a structural basis for future efforts to pharmacologically modulate these regulatory interfaces.

## Methods

### Molecular modeling of the systems

The cryo-EM structure of the CLOCK–BMAL1–nucleosome complex is available (PDB ID: 8OSK).^12^ In this structure, the regions D92–S106 and S224–D247 of CLOCK and K130–L149, D219–S244, S264–D281, and W297–C315 of BMAL1 are unresolved. These missing regions were modeled using AlphaFold2.^25^ Two copies of histone H3 (Arg41–Ala136) were included based on the histone H3 sequence (UniProt ID: P68431), with an AlphaFold2 confidence score of 86.06. Two copies of histone H4 (Leu23–Gly103) were included based on UniProt ID: P62805, with a confidence score of 89.81. Two copies of histone H2A (Lys16–Lys119) were included based on UniProt ID: P04908, with a confidence score of 90.75. Two copies of histone H2B (Arg34– Lys126) were included based on UniProt ID: P06899, with a confidence score of 85.50. CLOCK residues Asp30–Glu384 were modeled based on UniProt ID: O08785, with a confidence score of 88.39 and iPTM of 0.52, whereas BMAL1 residues Ala80–Val447 were modeled based on UniProt ID: Q9WTL8, with a confidence score of 80.33 and iPTM of 0.53. The resulting model contained a total of 12 chains, comprising a synthetically constructed double-stranded DNA, core human histone proteins, and the Mus-musculus transcription factors CLOCK and BMAL1. The AlphaFold2-modeled structures of the histones, CLOCK, and BMAL1 were subsequently repositioned by aligning the corresponding resolved regions to those of the template structure (PDB ID: 8OSK).^12^ The aligned protein coordinates were then combined with the DNA structure retained from the reference model, thereby preserving the structural arrangement observed in the experimentally resolved complex. The resulting protein–DNA complex was subsequently energy-minimized and structurally refined using Maestro–Schrödinger,^26^ then used as the starting model for the simulations.

### System Preparation

As detailed in the previous section, the model structure of the CLOCK/BMAL1–nucleosome system was prepared at atomistic resolution. The DNA component of the complex was subsequently converted to a coarse-grained (CG) representation using the martinize-dna.py, with the ds-stiff option applied to maintain a rigid double-stranded DNA structure. Similarly, the protein component was coarse-grained using the martinize.py, with the ElNeDyn elastic network model enabled (-ff elnedyn22).^27^ Default values were used for the force constant and upper cutoff parameters. Backbone restraints were assigned according to the secondary-structure elements predicted by DSSP.^28^ All system preparation steps were performed using GROMACS version 2023.^29^ The resulting CG DNA–protein complex was placed at the center of a 22 × 22 × 22 nm^3^ dodecahedral simulation box. The system was solvated using the Martini water model,^29,30^ and subsequently neutralized and ionized to a final NaCl concentration of 0.15 M. The resulting solvated and ionized CG representation of the DNA–protein complex was then used as the starting system for the molecular dynamics simulations.

### Coarse-Grained Molecular Dynamics Simulations and Backmapping

Coarse-grained molecular dynamic simulations were run with GROMACS version 2023.^29^ Energy minimization was conducted with the steepest descent algorithm until the maximum force fell below 5000 kJ/mol/nm, with 30,000 steps and an initial step size of 0.01 ps (integrator = steep). The potential energy reached approximately −1.64×10^6^ kJ/mol, with maximum force values converging around 4.5×10^3^ kJ/mol/nm. Minimization was followed by NPT equilibration using the velocity rescale thermostat (τ_t_ = 2.0, ref_t = 300 K) and C-rescale barostat (τ_p_ = 3.0, ref_p = 1.0 bar) for 250 ns with a 20 fs timestep (250,000 steps), under periodic boundary conditions and with a Verlet cutoff scheme. Production simulations were then performed for 10 μs with a timestep of 20 fs (500 million steps) using the Parrinello-Rahman barostat and the same thermostat.^31^ All interactions were handled using the reaction-field method for electrostatics (εr = 15, rcoulomb = 1.1 nm) and a cutoff scheme for van der Waals interactions (rvdw = 1.1 nm) with potential-shift-verlet modifiers. Trajectories and energy outputs were saved at appropriate intervals, and energy drift was minimal, with conserved energy deviation around −1.63×10^−3^ kJ/mol/ps per atom, confirming simulation stability. Each system was simulated using five independent replicates of 10 μs each, yielding a total of 200 μs molecular dynamics data. For network analysis, CG trajectories were backmapped using cg2at metholodogy.^32^ Using the CHARMM36 (July 2020 updated) force field,^33^ and the Martini 2.0 mapping scheme,^30^ each frame taken from the CG simulations was converted into an all-atom representation. The process was repeated for each frame along the trajectory, with each frame being aligned to the reference atomistic structure. The TIP3P water model was used for solvation. The time-line plots for each replicate are provided in Supplementary Figure 2-15 and results presented herein correspond to average of the individual replicates.

### Root Mean Square Deviation

To evaluate the structural stability of the CLOCK-BMAL1 and histones during the simulations, the root mean square deviation (RMSD) analysis was carried out. Calculations were done using the gmx rms tool in GROMACS,^29^ and RMSD values were calculated for the backbone beads of each subunit separately. All trajectory frames were aligned relative to the DNA before calculation to eliminate global translational and rotational motions. The RMSD at time t was computed using the standard equation:

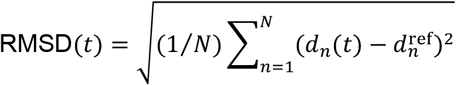

where:

*N*: denotes the number of atoms,

*d*_*n*_(*t*): the position of atom *n* at time *t*,

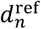: the corresponding position in the reference structure.

### Number of Contacts

The intermolecular interactions within the systems were quantified by calculating the number of contacts between the centers of mass (COMs) of selected groups of CLOCK/BMAL1, DNA, and nucleosome. Contact analysis was performed using a distance cutoff of 0.6 nm. The data obtained from each replicate was averaged and reported for the corresponding system.

### Dynamic Cross Correlation Map

Dynamic cross-correlation maps (DCCMs) were created with the Bio3D package to discover correlated and anti-correlated motions between residues for the merged trajectories.^34^ The analysis was performed on backbone beads, with the correlations being calculated according to the following expression:

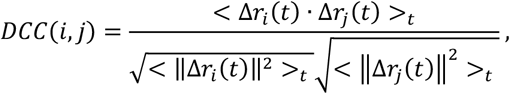

where *r*_*i*_*(t*) and *r*_*j*_*(t)* are the coordinates of atoms i and j at time t, and ⟨·⟩t refers to the time average. The displacement vectors Δ*r*_*i*_*(t)* and Δ*r*_*j*_*(t)* are defined as the deviation from the time-averaged positions. Correlation coefficients are in a range between −1 to 1, where a positive value indicates correlated motion, and a negative value indicates anti-correlated motion. The final DCCMs were obtained using the meta trajectories obtained by merging individual replicates.

### Principal Component Analysis (PCA)

Principal component analysis was performed to examine dominant collective motions and structural variations across the trajectories. For this, the trajectories were first merged and aligned with respect to the backbone atoms of the reference structure. Covariance matrices, which define positional changes of atoms, were computed by the following formula:

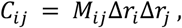

where *C*_*ij*_ represents the covariance between atoms i and j, and Δr refers to the deviations from the time-averaged position.

The covariance matrix was then diagonalized, yielding a set of eigenvalues and eigenvectors that describe both the amplitude and direction of the principal motions, accordingly. This relationship can be defined as:

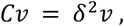

where *Cv* defines the covariance matrix, *δ*^2^ refers to the eigenvalues, and *v* the eigenvectors.

To further investigate the dynamic global motions, the two-dimensional principal component projections were created for each system by aligning the trajectories to their reference structures. Covariance matrices were computed as well from the displacements of backbone atoms, which were then diagonalized using the gmx covar and gmx anaeig modules of GROMACS,^29^ yielding the corresponding eigenvalues and eigenvectors. By defining the first two principal components, each system was then projected onto a two-dimensional space. For the distribution of conformational states to be visualized along the main eigenvector, the ProDy Python library was used thus allowing for an examination and comparison of large-scale dynamic behaviors across wild-type and phosphorylated systems.^35^

### Global Network Analysis

Global network analysis was performed on backmapped trajectories to characterize residue-level interaction networks. Representative structures together with the corresponding trajectories were used as input to generate protein structure networks (PSNs). Each amino acid residue was considered as a node in the PSN representation, and the strong non-covalent interactions between residue pairs were presented as links connecting the corresponding nodes.^36–38^

The interaction strength between each pair of residues *i* and *j* was calculated as:

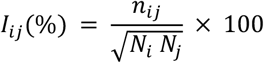

where *n*_*ij*_ is the number of side-chain heavy atom pairs that belongs to residues *i* and *j* which occur within a distance of 4.5 Å, while *N*_*i*_ and *N*_*j*_ denote the residue-specific normalization parameters that take into consideration variations in side-chain size and their inherent tendency to form interactions with neighboring residues. Only residue-residue interactions that satisfied *I*_*ij*_ ≥ *I*_*min*_ were kept as network linkages. The PSN process automatically calculated the value of Imin based on the critical interaction strength, Icritical. This critical number is the point at which when no interaction-strength threshold is applied (*I*_*min*_ = 0), the largest connected group of residues decreases to about half of its size as the interaction-strength threshold is increased.^36,38^

Each residue-residue interaction was also considered by its link frequency, which is computed as follows:

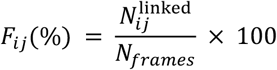

where 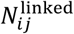 represents the number of trajectory frames where residues *i* and *j* met the interaction condition *I*_*ij*_ ≥ *I*_*min*_ and *N*_*frames*_ is the total number of trajectory frames examined. *F*_*ij*_ indicates the percentage of the trajectory with a certain residue-residue interaction. Since no analysis parameters were changed manually, the default PSN-MD link-frequency cutoff of 50% was applied. This means that a link was retained in the final network if it was present in at least 50% of the conformational ensemble investigated.^38^

The WebPSN server was then used to evaluate the generated PSNs to find hubs, communities, and structural communication pathways. Hubs stand for strongly connected regions within the residue interaction network as they were characterized as residues participating in at least four network links, while Communities stand for the connected regions of the network where residues exhibited relatively few connections with residues outside the community but strong connections to other residues within the same community. However, sets of three fully connected nodes (k = 3 cliques) serve as the basis for WebPSN community construction. These cliques are combined when they have at least one connection in common.^38,39^

The structural communication of the systems studied was described by using the shortest communication pathways occurring between the pairs of network nodes. Then, WebPSN finds by default the shortest path between two endpoint residues using the minimum number of intermediate residues by searching for pathways between all possible node pairs. The associated residue movements data from the trajectories were then used to filter these pathways.^38,40^ By using linear mutual information (LMI), the corresponding dynamic associations were then analyzed for PSN files by using the equation below:

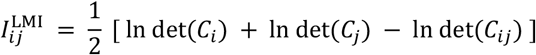

where *C*_*ij*_ denotes the joint covariance matrix which indicates the combined fluctuations of residues *i* and *j, C*_*i*_ and *C*_*j*_ denote the covariance matrices which indicate their corresponding positional fluctuations. Thus, LMI determines the degree to which one residue’s motion is statistically associated with another residue’s motion along the trajectory.^40^

The calculated LMI is then converted into a normalized motion-correlation coefficient using:

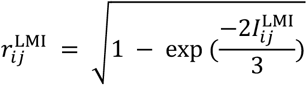

where 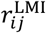 ranges from 0 to 1 such that values near 0 suggest weak or absent correlated motion and those near 1 indicate increasingly strong correlated motion. The denominator of 3 represents the three Cartesian dimensions (*x, y, and z*) that describe atomic positional fluctuations. With the default PSN-MD parameters, a motion-correlation coefficient cutoff of 0.8 was used, and a communication pathway was preserved if at least one internal node of the pathway showed associated motion above this threshold with one of the two pathway endpoints.^38,40^

Finally, the global metapath was constructed by the total pool of the dynamically filtered shortest communication pathways. Rather than a single communication pathway, the global metapath represents the nodes and links that are found most frequently among the filtered pathway ensemble, thus summarizing the predominant residue-to-residue communication routes within the analyzed system. The default PSN settings used 20% link-recurrence cutoff for constructing the global metapath.^38^

### Steered Molecular Dynamics Simulations

According to the PER-dependent CLOCK/BMAL1 displacement mechanism, PER binds to the CLOCK/BMAL1 complex and recruits CK1δ, which phosphorylates CLOCK and BMAL1 at the identified phosphorylation sites, ultimately promoting the dissociation of the transcription factor complex from the nucleosome. To investigate how phosphorylation affects the dissociation behavior of the CLOCK and BMAL1 bHLH domains from the E-box, constant-velocity steered molecular dynamics (SMD) pulling simulations were performed. For this purpose, representative structures were selected from the unphosphorylated and double-phosphomimetic systems based on the contact-number distributions presented in Figures 2A and 2B. In both unphosphorylated and double-phosphomimetic systems, the low-contact state was more populated for the CLOCK bHLH domain. In contrast, the BMAL1 bHLH domain predominantly adopted the high-contact state in the unphosphorylated system, whereas the low-contact state was more populated in the double-phosphomimetic system. Importantly, because the objective of the pulling simulations was to specifically examine the dissociation pattern of the bHLH domains from the E-box, representative structures were selected such that no contacts were present between the PAS domains and the nucleosome. To obtain such a state, the PAS domains were first detached from the histones using steered molecular dynamics simulations, with a pulling velocity of 0.005 nm ps^−1^ and a force constant of 1800 kJ mol^−1^ nm^−2^ along the z direction. As a result, the selected representative structures exhibited a slight deviation in their initial E-box contact numbers from the maximum contact values shown in Figures 2A and 2B. Nevertheless, the overall binding patterns observed in the corresponding contact distributions were preserved. That is to say, the number of contacts formed by the CLOCK bHLH domain is comparable between the unphosphorylated and phosphorylated systems. In contrast, the BMAL1 bHLH domain is less in the unphosphorylated system than in the phosphorylated starting system.

Prior to the production pulling simulations, the pulling velocity and force constant were optimized through a series of preliminary tests. Because the pulling simulations were performed using a coarse-grained representation, the force constant was selected to avoid introducing excessive mechanical forces that could compromise the structural integrity of secondary-structural elements. When force constants higher than 500 kJ mol^−1^ nm^−2^ were tested, partial unwinding of the helices within the bHLH domains was observed (Supplementary Video). Accordingly, a pulling velocity of 0.01 nm ps^−1^ and a force constant of 500 kJ mol^−1^ nm^−2^ were selected for all subsequent simulations. The CLOCK and BMAL1 bHLH domains were defined as the pulling groups, while the nucleosome was used as the reference group. Simulations were repeated ten times for each system for examining reproducibility of the results. Each replicate was initiated with a different initial velocity distribution. All pulling simulations were carried out under the same ionic conditions, temperature, and pressure as the production molecular dynamics simulations.

**Figure 2.**
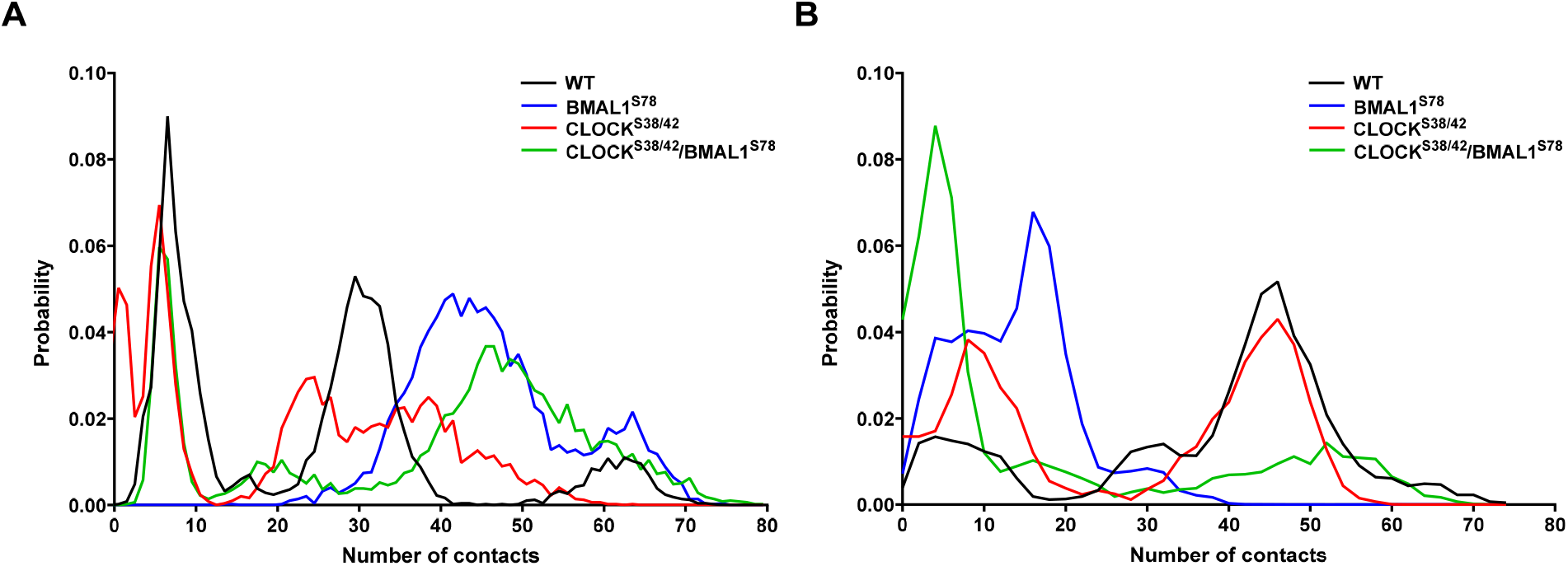
Probability distributions of the contact numbers between the E-box DNA and **A.**the CLOCK bHLH domain and **B**. the BMAL1 bHLH domain, calculated over the five replicates of simulation trajectories.

### Generation of CRY1 bound CLOCK-BMAL1-Nucleosome Model

A structural model of CRY1 associated with the CLOCK–BMAL1–nucleosome complex was generated using the AlphaFold Server to explore a possible binding mode of CRY1 within the nucleosome-bound CLOCK– BMAL1 complex.^41^ The modeled system contained the photolyase homology region (PHR) of human CRY1 (residues 1–506; UniProt Q16526), the bHLH–PAS regions of CLOCK (residues 30–384; UniProt O15516) and BMAL1 (residues 80–447; UniProt Q9WTL8), and the histone and DNA components corresponding to the experimentally determined CLOCK–BMAL1–nucleosome complex (PDB ID: 8OSK). The protein and DNA sequences were provided to the AlphaFold Server in the same component composition and order as in 8OSK, with CRY1 included as an additional component. The highest-ranked CRY1-containing model had a ranking score of 0.53, with predicted TM (pTM) and interface predicted TM (ipTM) scores of 0.48 and 0.46, respectively. Model confidence was further evaluated using predicted local distance difference test (pLDDT) scores and predicted aligned error (PAE). PAE was used to assess confidence in the relative positioning of residues and components within the complex. Lower PAE values indicate higher confidence in their relative spatial arrangement, whereas higher values indicate greater uncertainty in their relative positioning. The structure was visualized according to pLDDT values to illustrate local prediction confidence (Supplementary Figure 1).

### Results

In this study, we focused on phosphorylation at CLOCK^S38/S42^ and BMAL1^S78^, which have previously been shown to play critical roles in modulating circadian period length.^16^ Consistent with the experimental design of Otobe et al.,^16^ phosphorylation was modeled using phosphomimetic substitutions, in which the corresponding serine residues were mutated to acidic amino acids: Ser38 and Ser42 of CLOCK were substituted with aspartate, whereas Ser78 of BMAL1 was substituted with glutamate.

### Phosphorylation exhibits asymmetric effects, with BMAL1^S78^ phosphorylation enhancing the interaction between the CLOCK bHLH domain and the E-box

The phosphorylation sites in CLOCK and BMAL1 are both located within the bHLH domains and lie in similar proximity to the DNA-binding interface (Figure 1C). To investigate how these phosphorylation patterns influence the dynamics of the CLOCK/BMAL1 and nucleosome complex, we performed coarse-grained molecular dynamics (CG-MD) simulations on four distinct systems: (i) unphosphorylated, containing no phosphomimetic substitutions; (ii) BMAL1^S78E^, containing a phosphomimetic substitution at BMAL1 Ser78; (iii) CLOCK^S38/S42D^, containing phosphomimetic substitutions at CLOCK Ser38 and Ser42; and (iv) CLOCK^S38/S42D^/BMAL1^S78E^, containing phosphomimetic substitutions in both proteins.

In the unphosphorylated simulations, both CLOCK and BMAL1 bHLH domains exhibited bimodal binding mode with the E-box, indicating the presence of two distinct DNA-binding states. For CLOCK, contact distributions reveal two distinct binding states centered at approximately 10 and 30 contacts with the low-contact state being more sampled (Figure 2A, black). BMAL1 displayed a similar bimodal distribution; however, the high-contact state was more dominant, indicating stronger E-Box engagement relative to CLOCK under unphosphorylated conditions (Figure 2B, black). Phosphorylation produced markedly distinct effects on the DNA-binding behavior of the two proteins. In the CLOCK^S38/S42D^ system, the contact-number distribution of the CLOCK bHLH domain became broader with a reduced maximum around the high-contact state (Figure 2A, red). Likewise, the BMAL1 bHLH domain exhibited enhanced sampling of states with fewer E-box contacts (Figure 2B, red), suggesting a phosphorylation-induced weakening of DNA engagement. In contrast, BMAL1^S78E^ disrupted the bimodal binding pattern and enhanced the E-box engagement by the CLOCK bHLH domain (Figure 2A, blue) while reducing it with the BMAL1 bHLH domain (Figure 2B, blue). In the double phosphomimetic system (CLOCK^S38/S42D^/BMAL1^S78E^), the bimodal binding mode was maintained, whereas the high-contact state of the CLOCK bHLH domain became broader and shifted toward higher contact numbers (Figure 2A, green). Conversely, E-box contacts of the BMAL1 bHLH domain were reduced (Figure 2B, green). Together, these findings reveal that phosphorylation at structurally analogous positions within the CLOCK and BMAL1 bHLH domains produces asymmetric regulatory outcomes. Rather than exerting equivalent effects on

DNA binding, BMAL1^S78E^ phosphorylation appears to allosterically enhance CLOCK–DNA engagement. It is important to note that a previous study by Otobe and colleagues reported that phosphorylation of BMAL1 at S78 reduces the interaction between the CLOCK bHLH domain and the E-box.^16^ In contrast, our simulations indicate an increase in CLOCK–DNA contacts upon BMAL1^S78E^ phosphorylation. One possible explanation for this discrepancy is the different molecular systems employed in the two studies. Unlike previous work, our simulations explicitly include histones and the nucleosomal environment. Because the CLOCK bHLH domain interacts not only with DNA but also with histones, phosphorylation-induced perturbations originating in BMAL1 may be transmitted through the nucleosome, leading to a redistribution of interactions that ultimately enhances CLOCK bHLH engagement with the E-box.

According to the PER-dependent CLOCK/BMAL1 displacement mechanism, binding of PER to the CLOCK/BMAL1 complex and recruitment of CK1δ to facilitate phosphorylation of CLOCK/BMAL1, ultimately promoting dissociation of the CLOCK/BMAL1 complex from the nucleosome. To examine the effect of phosphorylation on complex dissociation, constant-velocity-steered molecular dynamics (SMD) simulations were performed using the unphosphorylated and double-phosphomimetic systems, as described in the Methods section. During the pulling simulations, the number of contacts between the CLOCK/BMAL1 bHLH domains and the E-box was monitored as a function of time, and the resulting contact profiles are presented in Figure 3B and 3C. The force peak observed during the SMD pulling simulations represents the first major mechanical resistance encountered as the pulling group begins to dissociate from its interaction interface. As shown in Figure 3B and C, both the unphosphorylated and double-phosphomimetic systems exhibited a similar initial peak for the CLOCK bHLH domain. However, complete dissociation of the contacts between the CLOCK bHLH domain and the E-box occurred at a later stage in the unphosphorylated system. In contrast, a distinct difference was observed for the BMAL1 bHLH domain. In the unphosphorylated system, the maximum number of E-box contacts reached approximately threefold or more of the initial contact number, whereas this increase was limited to approximately 1.5-fold in the double-phosphomimetic system (Figure 3C). Finally, it is worth noting that, despite exhibiting a higher number of contacts with the E-box, the BMAL1 bHLH domain dissociated earlier during the pulling simulations than the CLOCK bHLH domain. This apparent discrepancy may arise from the additional interactions formed by the CLOCK bHLH domain with the DNA and histone components of the nucleosome, which may provide additional resistance to its dissociation during the pulling process.

**Figure 3.**
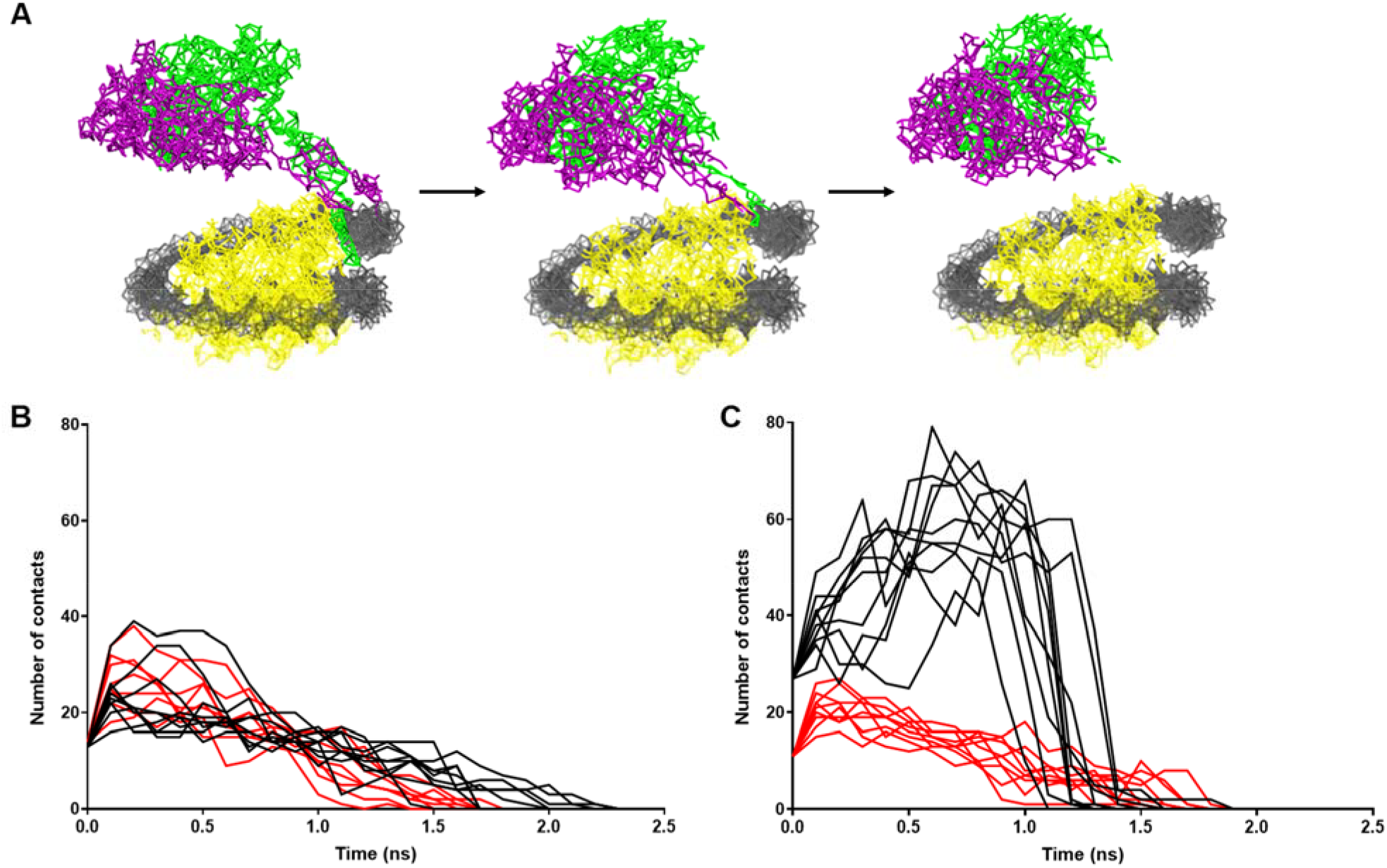
**A**. Schematic depiction of dissociation of CLOCK/BMAL1 bHLH domains from E-box during SMD pulling simulations. Gray, yellow, purple and green represent the DNA, histones, BMAL1 and CLOCK, respectively. Plots show the number of contacts formed between E-box and **B**. CLOCK bHLH domain, **C**. BMAL1 bHLH domain throughout the SMD pulling simulations. Black replicates pertain to unphosphorylated systems, red replicates pertain to double-phosphorylated systems.

Next, to examine the relative orientation of the CLOCK and BMAL1 bHLH domains we described an angle using one reference point on the CLOCK bHLH domain, one reference point on the BMAL1 bHLH domain, and a third point located at their intersection/interface (Figure 4A). More specifically, it can be interpreted as a measure of the opening/closing or packing geometry of the two bHLH domains within the CLOCK/BMAL1 heterodimer. Accordingly, the angle was reduced and exhibited a broader distribution relative to the unphosphorylated system in the BMAL1^S78E^ phosphorylation, whereas CLOCK^S38/S42D^ phosphorylation resulted in a slightly more open angle comparable to that observed in the unphosphorylated system (Figure 4B). In the double-phosphomimetic system, the angle distribution was also broader and exhibited two distinct peaks. The dominant peak corresponded to a slightly smaller angle than that observed in the unphosphorylated system, whereas the less frequently sampled state was characterized by a larger angle.

**Figure 4.**
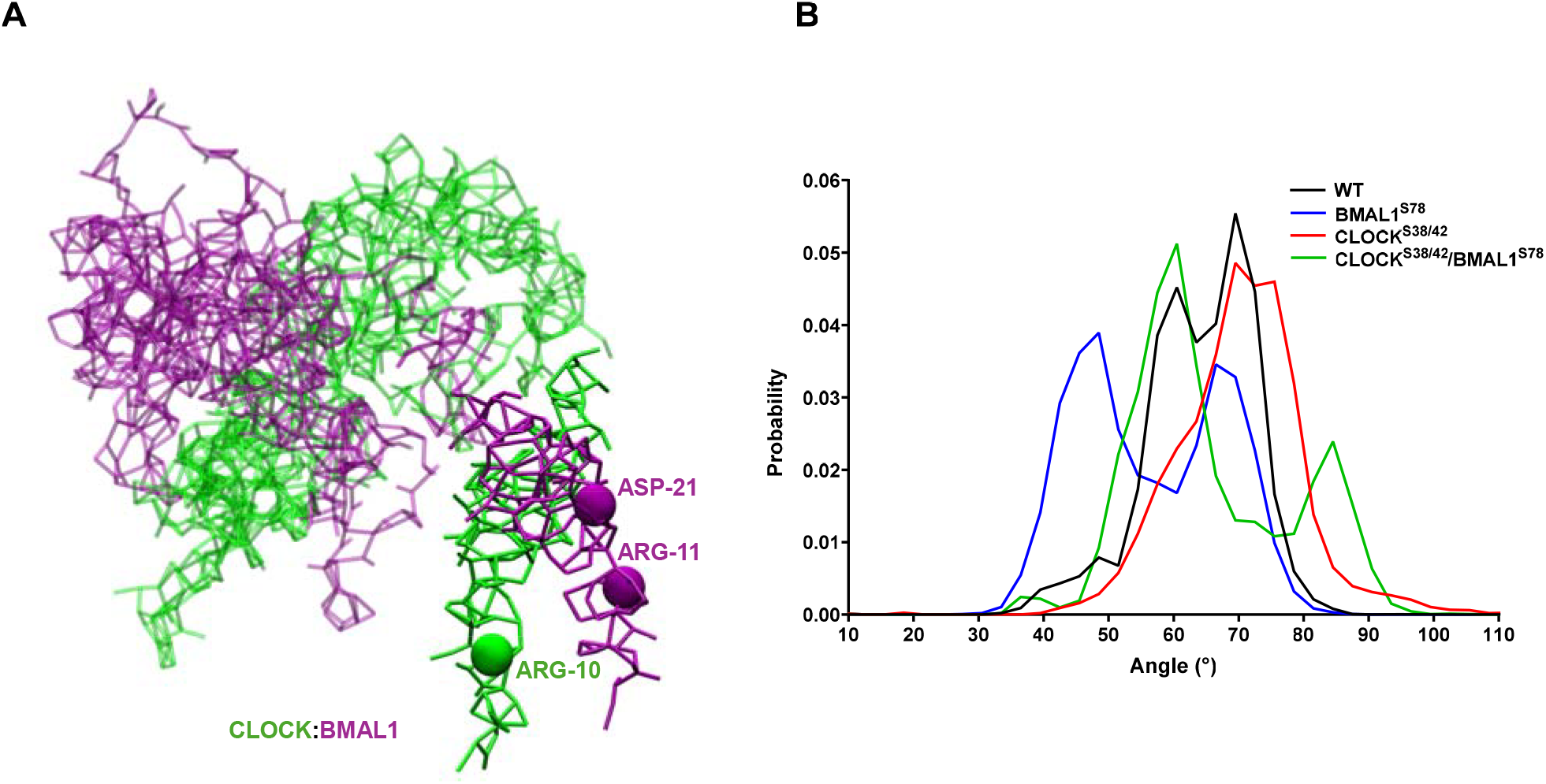
**A**. The Cα atoms of the amino acids used to describe the angle are depicted by balls. **B**. Probability distributions of the angle described between the CLOCK and BMAL1 bHLH domain, calculated over the five replicates of simulation trajectories.

### Phosphorylation remodels residue-residue correlations within the CLOCK/BMAL1 complex

To investigate whether phosphorylation alters residue–residue correlations, we examined dynamic cross-correlation patterns across the systems studied. This type of analysis provides information on the extent to which different regions of the protein exhibit correlated, anti-correlated, or uncorrelated motions. Positive correlations indicate that two regions tend to fluctuate in the same direction, reflecting coordinated motions, whereas negative correlations indicate anti-correlated motions, in which the two regions fluctuate in opposite directions. Regions showing little or no correlation exhibit largely independent motions.

Our analysis showed that phosphorylation of CLOCK increased anti-correlated motions between the bHLH domain and PAS-A and PAS-B domains. Moreover, the pronounced anticorrelated motions observed in the unphosphorylated complex between the PAS-A and PAS-B domains were largely abolished following CLOCK phosphorylation (Figure 5A). These findings suggest that phosphorylation disrupts long-range interdomain communication within CLOCK, potentially modulating the ability of the complex to engage nucleosomal DNA. Interestingly, phosphorylation of BMAL1 also substantially altered the dynamic correlation landscape of CLOCK. In the BMAL1^S78E^ system, positive correlations within the PAS-A domain decreased whereas that of between the PAS-A and PAS-B domains were increased (Figure 5B), indicating that phosphorylation-induced perturbations are transmitted across the heterodimer interface and can allosterically reshape the conformational dynamics of the partner protein.

**Figure 5.**
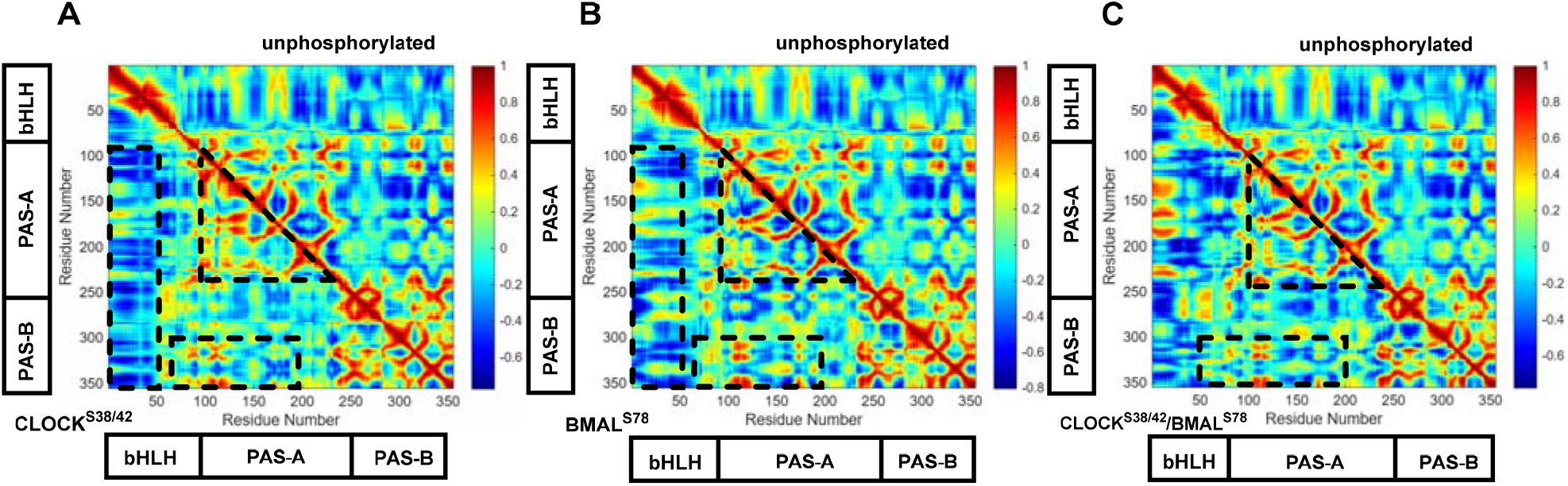
Dynamic cross-correlation matrices of CLOCK pertaining to **A.** CLOCK^S38/S42D^, **B**. BMAL^S78E^, and **C**. CLOCK^S38/S42D^-BMAL^S78E^ calculated over the five replicates of simulation trajectories. The regions that display different correlation compared to unphosphorylated system are shown in dashed rectangle.

In the double phosphomimetic system, the positive correlation between the bHLH and the PAS-A domains, as well as between the PAS-A and PAS-B domains, is enhanced relative to the unphosphorylated system (Figure 5C). Notably, the internal correlation within the PAS-A domain is consistently reduced, regardless of phosphorylation status. Together, these findings suggest that phosphorylation alters the pattern of coordinated and coupled motions between the bHLH and PAS-A/PAS-B domains, as well as between the PAS-A and PAS-B domains. Such changes in interdomain dynamics may, in turn, influence how these domains interact with the nucleosome.

We next examined phosphorylation-dependent correlation changes within BMAL1 itself. BMAL1 phosphorylation enhanced anticorrelated motions both between the bHLH and the PAS-A domain and between the PAS-A and PAS-B domains (Figure 6B). Additionally, positive correlations within the PAS-B domain became more pronounced, suggesting increased internal coordination within this region. Consistent with reciprocal effects observed in CLOCK, CLOCK phosphorylation also altered the dynamic correlation network within BMAL1. Specifically, CLOCK^S38/S42D^ increased the correlated motion between BMAL1 bHLH domain and the PAS-B domain, whereas positive correlations within the PAS-B domain decreased (Figure 6A). Importantly, the double phosphomimetic system converged in the CLOCK^S38/S42D^/BMAL^S78E^ system, indicating that phosphorylation-induced dynamic perturbations propagate bidirectionally across the heterodimer interface (Figure 6C). Overall, residue–residue correlation network analysis revealed that phosphorylation extensively rewires dynamic communication pathways within the CLOCK–BMAL1 complex. Rather than producing purely local structural effects, phosphorylation modulates long-range allosteric coupling between DNA-binding and regulatory domains, thereby reshaping the dynamical architecture of the heterodimer.

**Figure 6.**
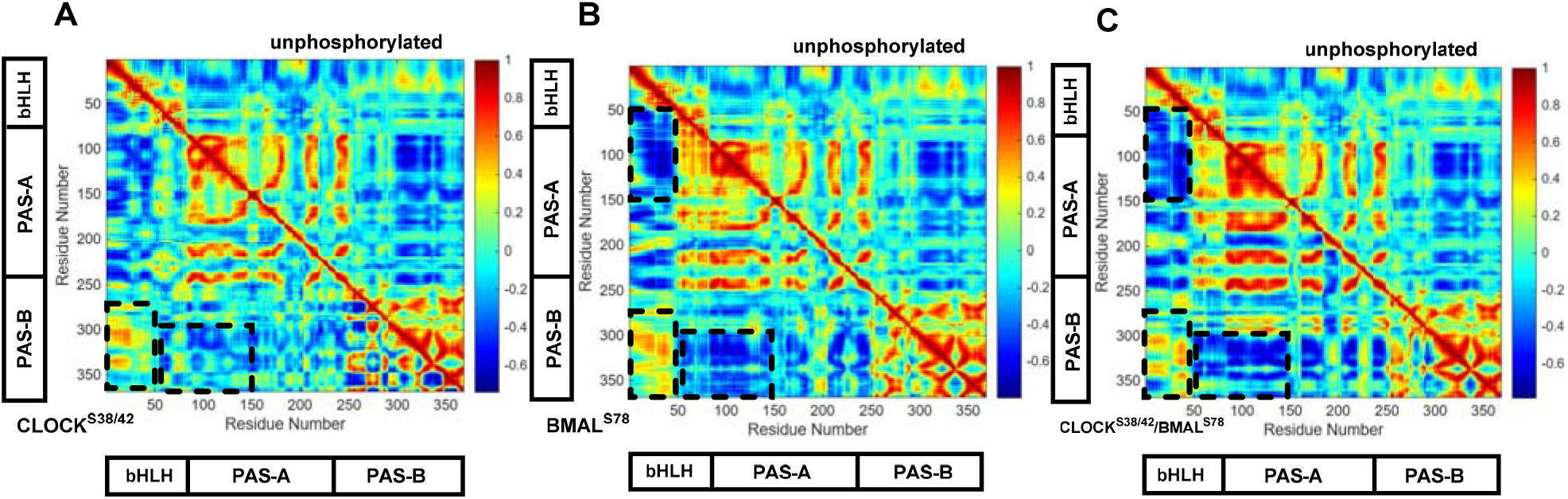
Dynamic cross-correlation matrices of BMAL1 pertaining to **A.** CLOCK^38/42^, **B**. BMAL^S78^, and **C**. CLOCK^S38/S42^/BMAL^S78^ calculated over the five replicates of simulation trajectories. The regions that display different correlation compared to unphosphorylated system are shown in dashed rectangle.

Overall, CLOCK and BMAL1 phosphorylation differentially modulate the dynamic coupling between the bHLH and PAS domains of the CLOCK/BMAL1 heterodimer. Upon CLOCK phosphorylation, the anticorrelated motions between the CLOCK bHLH domain and the PAS-A/PAS-B region are enhanced, whereas the PAS-A and PAS-B domains exhibit increased correlated motions. In contrast, BMAL1 phosphorylation enhances the anticorrelated motions between the BMAL1 bHLH domain and PAS-A, accompanied by increased anticorrelation between the PAS-A and PAS-B domains. Thus, the two phosphorylation events induce distinct patterns of correlated and anticorrelated motions within the heterodimer (Figure 7).

**Figure 7.**
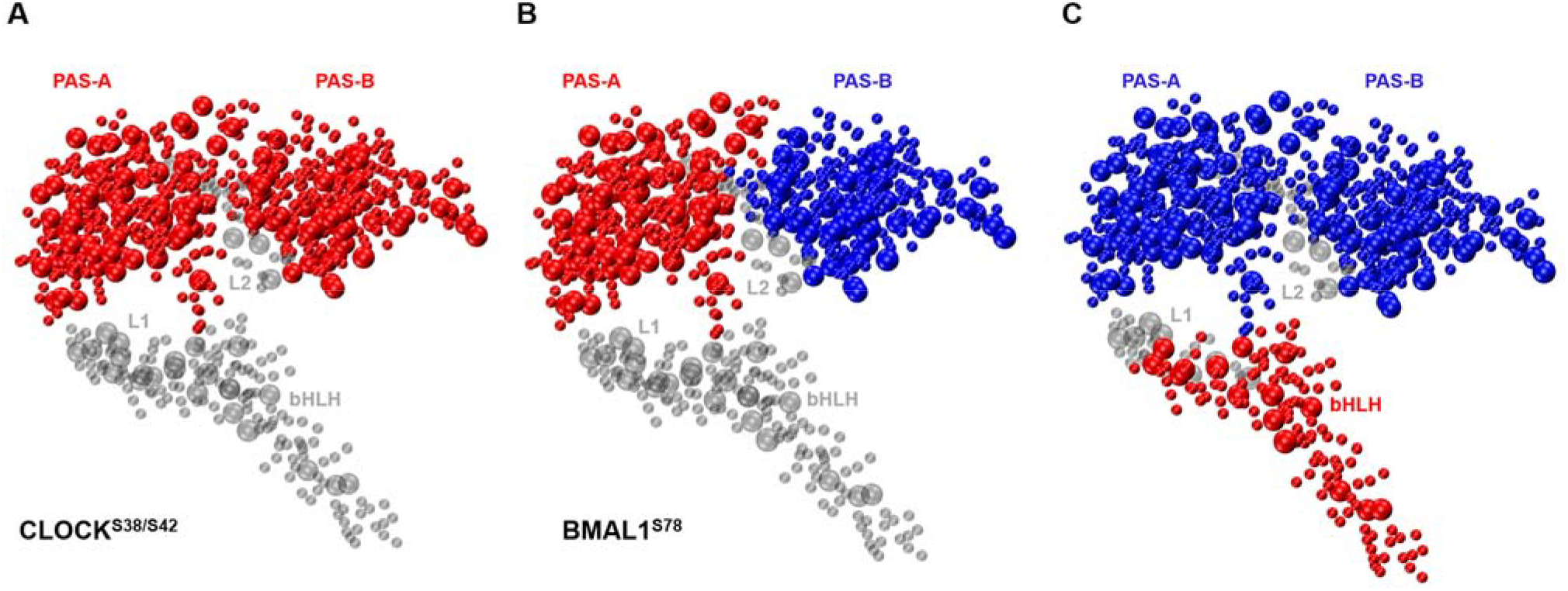
Structural mapping of cross correlation-modulations upon phosphorylation. **A.** Correlated motions between PAS-A/PAS-B domains of CLOCK are shown in red in CLOCK^S38/S42^. **B**. Anticorrelated motions between PAS-A and PAS-B of BMAL1 are shown in red and blue in BMAL1^S78^ **C**. Anticorrelated motions between the bHLH domain and PAS domains are shown in red and blue, respectively. The L1 and L2 linkers are shown in gray.

### Phosphorylation of CLOCK and BMAL1 bHLH Domains Remodels Nucleosomal Interactions

In addition to phosphorylation-dependent modulation effects observed at the E-box, we next investigated whether phosphorylation alters interactions between CLOCK/BMAL1 and histone octamer, as well as histone– DNA contacts within the nucleosome. Analysis of the trajectories revealed that phosphorylation reduces interactions between the CLOCK bHLH domain and histones except the double phosphorylated system as it displays a similar but broader maximum (Figure 8A). In contrast, no detectable modulation was observed for the BMAL1 bHLH domain, which remained spatially separated from the histone core owing to its positioning on the opposite side of the DNA relative to the nucleosome surface.

**Figure 8.**
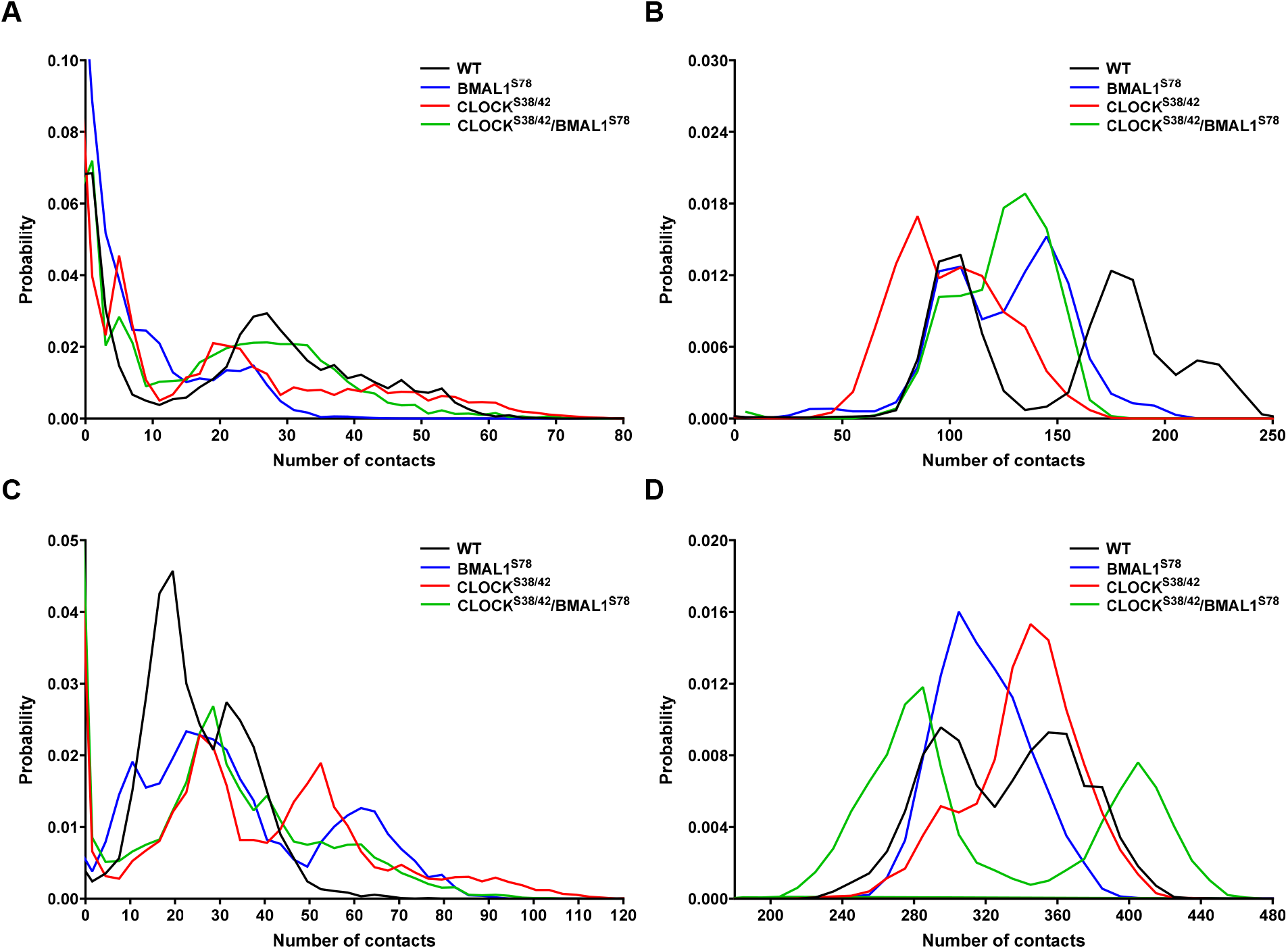
Probability distributions of the number of contacts formed between histones and **A.**the CLOCK bHLH domain, **B**. CLOCK PAS-B, **C**. BMAL1 PAS-A, and **D**. DNA calculated over the five replicates of simulation trajectories.

We further analyzed the effects of phosphorylation on histone interactions mediated by PAS domains. Neither the CLOCK PAS-A domain nor the BMAL1 PAS-B domain formed measurable contacts with histones, regardless of the phosphorylation state. In contrast, persistent interactions were observed for CLOCK PAS-B as well as the BMAL1 PAS-A domains. CLOCK PAS-B domain exhibited a bimodal contact distribution with peaks centered at approximately 100 and 175 contacts. Upon phosphorylation, the overall number of contacts decreased, and the bimodal character of the distribution diminished, consistent with a redistribution toward weaker histone engagement (Figure 8B). Conversely, phosphorylated systems showed increased contact between the BMAL1 PAS-A domain and histones, suggesting differential allosteric responses of the two proteins to phosphorylation within the bHLH region (Figure 8C).

Finally, we assessed whether phosphorylation modulates histone–DNA interactions within the nucleosome. In the unphosphorylated complex, histone-DNA contacts exhibited a bimodal distribution characterized by two closely spaced peaks. Phosphorylation altered this equilibrium in a modification-specific manner. In the CLOCK^S38/S42D^ system, the dominant population shifted toward the higher-contact state, whereas BMAL1^S78E^ system favored the lower-contact state (Figure 8D). Notably, in the double phosphomimetic system (CLOCK^S38/S42D^/BMAL1^S78E^), the two peaks became more clearly separated, indicating enhanced partitioning between distinct nucleosomal-contact states.

Collectively, these findings suggest that phosphorylation reshapes not only CLOCK–BMAL1 interactions with DNA, but also broader nucleosomal interaction network. The observed redistribution of protein–histone and histone–DNA contacts suggest that phosphorylation propagates structural effects throughout the entire nucleosome and CLOCK/BMAL1 assembly, thereby modulating its global dynamical landscape.

### Phosphorylation rewires allosteric communication networks within the CLOCK/BMAL1 complex

Motivated by the phosphorylation-dependent changes observed in residue–residue cross-correlation analysis, we further examined whether phosphorylation affects allosteric communication networks within the CLOCK-BMAL1 complex. To this end, we identified global allosteric communication pathways defined as ensembles of shortest paths that collectively capture the distribution of communication flow across the protein. In this framework, pathways with higher occurrence frequencies are interpreted as making greater contributions to intramolecular signal propagation.

In the unphosphorylated complex, prominent communication pathways were observed between the two bHLH domains, as well as between the bHLH and PAS-A domains, with occurrence frequencies of approximately 50–60%, which is represented by light blue color (Figure 9A). Additionally, pathways originating from the PAS-A domain and extending toward the PAS-A/PAS-B interface are observed with higher frequencies (∼60–70%, green). Direct communication pathways connecting the PAS-A and PAS-B domains are also present, albeit with lower frequencies (Figure 9A, black and dark blue lines). Together, these results indicate that, under unphosphorylated conditions, allosteric communication is hierarchically organized, predominantly flows from the bHLH domains to PAS-A, and subsequently toward the PAS-A/PAS-B interface, while direct PAS-A/PAS-B coupling remains relatively limited.

**Figure 9.**
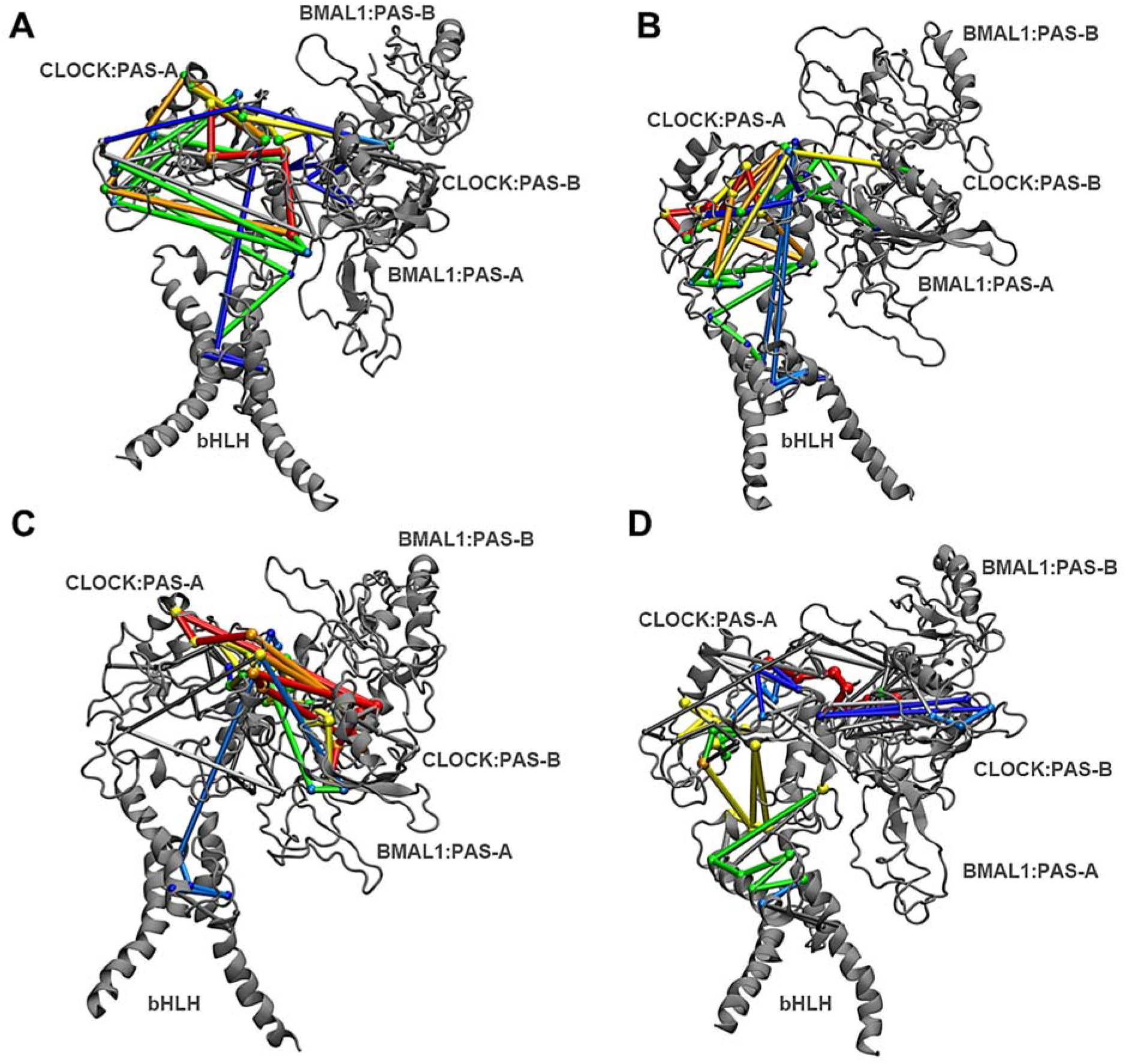
Allosteric communications pathways calculated using the five replicates of the systems studied. **A.** unphosphorylated CLOCK/BMAL1, **B**. CLOCK^S38/S42D^, **C**. BMAL1^S78E^ and **D**. CLOCK^S38/S42D^/BMAL1^S78E^. The color code reports on the frequency of presence of the networks: dark blue: %40-%50, light blue: %50-%60, green: %60-%70, yellow: %70-%80, orange: %80-%90, red: %90-%100.

CLOCK^S38/S42D^ induced a marked redistribution of these communication networks. The most prominent effect was an increase in the frequency of intra-PAS-A communication pathways relative to the unphosphorylated system, accompanied by a reduction in pathways connecting PAS-A and PAS-B (Figure 9B). In contrast, BMAL1^S78^ phosphorylation exhibited opposing reorganization of the allosteric network. Specifically, phosphorylation enhanced high-frequency pathways (>80%, orange) between the PAS-A and PAS-B domains, as well as within both PAS domains (Figure 9C). Concurrently, pathways within the PAS-A domain and those connecting the bHLH domain to PAS-A or to the PAS-A/PAS-B interface were reduced. These findings suggest that BMAL1 phosphorylation shifts the dominant communication flow away from the DNA-binding interface and toward enhanced coupling between regulatory PAS domains.

Finally, the double phosphomimetic system CLOCK^S38/S42^/BMAL1^S78^ displayed a partially convergent communication profile. Like BMAL1^S78E^ alone, pathways connecting PAS-A and PAS-B were redistributed, accompanied by an increased frequency of intra-PAS-B communication pathways (Figure 9D). Together, allosteric pathway analysis demonstrates that phosphorylation extensively rewires communication flow within the CLOCK–BMAL1 complex, producing modification-specific shifts in coupling between DNA-binding and regulatory domains.

### Phosphorylation-induced modulation of protein dynamics reshapes global conformational motions

Post-translational modifications of core clock proteins play important roles in coordinating the transitions between activation and repression phases of the circadian cycle.^42^ For instance, BMAL1^S90^ phosphorylation is required for efficient K537 acetylation and subsequent recruitment of CRY1/2 to BMAL1. However, whether phosphorylation-dependent remodeling of CLOCK influences CRY-binding has not been established.^43^

To address this gap, we performed principal component analysis (PCA) to investigate the impact of phosphorylation on the global dynamics of the complex. First, we found that, in the unphosphorylated system, the CLOCK HI loop does not remain stably engaged with the nucleosome but instead undergoes transient excursions away from the nucleosomal surface. These fluctuations sample conformations in which the HI loop becomes more accessible for potential CRY1 binding. This observation is consistent with previous biochemical evidence showing that CRY1 can simultaneously interact with the CLOCK HI loop and nucleosomal DNA, thereby enabling CRY1 to engage the CLOCK/BMAL1–nucleosome complex without necessarily requiring complete displacement of CLOCK/BMAL1 from the nucleosome. This mechanism is reminiscent of a blocking-type mode of transcriptional repression, in which the transcriptional activator remains associated with its regulatory site while its activity is inhibited by the repressor.^21,44^

Interestingly, phosphomimetic substitutions substantially remodel large-scale conformational motions within the assembly, particularly by altering the spatial relationship between CLOCK/BMAL1 and specific histone regions. These effects were dependent on the phosphorylation pattern. For instance, phosphorylation at CLOCK^S38/S42^ increases the distance between the CLOCK HI loop – a critical region for CRY1 and CRY2 binding – and the H3α1 L1 elbow (Lys^79^ and Thr^80^) (Supplementary Figure 9,10). This increased separation suggests that CLOCK phosphorylation alters the conformational landscape of the HI-loop region and may increase the accessibility of this CRY-binding interface. In contrast, BMAL1^S78^ phosphorylation alone did not induce a comparable effect. Notably, the double phosphomimetic system CLOCK^S38/S42^/BMAL1^S78^ exhibited the opposite behavior, displaying reduced separation between these regions relative to the unphosphorylated system, indicating that the effects of phosphorylation are not simply additive (Supplementary Figure 10,11).

The increased separation between the CLOCK HI loop and the nucleosome observed in the CLOCK^S38/S42^ simulations prompted us to examine whether CRY1 could be structurally accommodated by nucleosome-bound CLOCK–BMAL1. We therefore generated a three-dimensional model of a CRY1–CLOCK–BMAL1– nucleosome complex using the AlphaFold Server (Figure 10).^41^ Given the relatively low global confidence metrics (pTM = 0.48 and ipTM = 0.46), the model was not considered a definitive representation of the complete complex. Instead, it was used to examine whether CRY1 could be structurally accommodated by nucleosome-bound CLOCK–BMAL1 in a configuration compatible with blocking-type repression. Interestingly, the predicted model positioned the secondary-pocket region of CRY1 toward the CLOCK HI loop while simultaneously placing another surface of the CRY1 PHR domain in contact with nucleosomal DNA (Figure 10).

**Figure 10.**
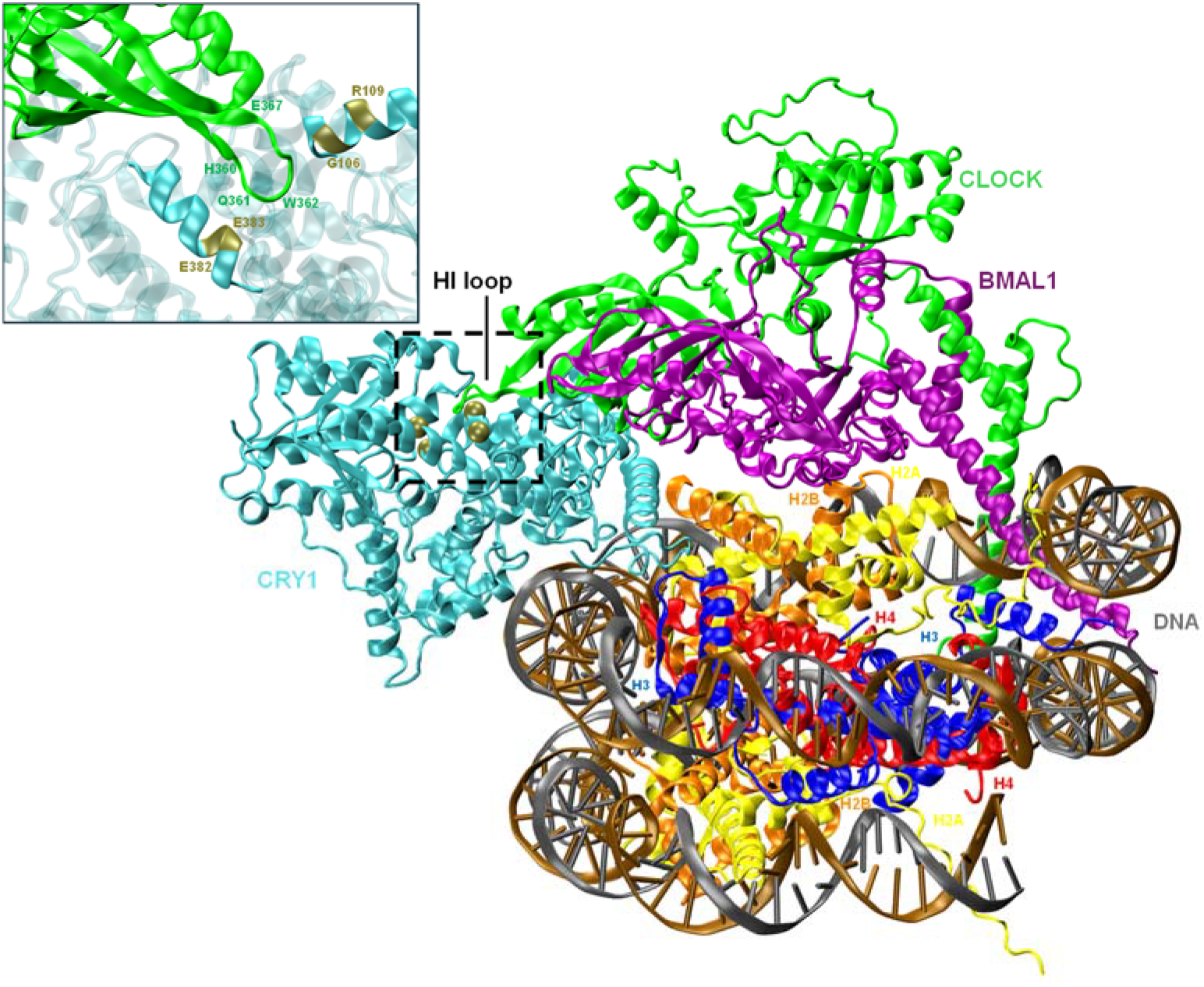
The three-dimensional structural model of the CRY1–CLOCK/BMAL1–nucleosome complex. The model is generated by using AlphaFold 3.^41^

Considering that the CLOCK HI loop constitutes an established CRY-binding interface, these dynamics suggest that phosphorylation could shift the conformational ensemble toward states in which the HI loop is more accessible to CRY1. Such remodeling would be compatible with formation or stabilization of the DNA-bound CRY1–CLOCK–BMAL1 complex characteristic of late, displacement-type repression.^10,17,42^ We emphasize that these observations do not establish CLOCK^S38/S42^ phosphorylation as a requirement for CRY1 recruitment. Rather, they suggest a potential structural mechanism through which phosphorylation-dependent remodeling of the CLOCK–nucleosome interface could modulate accessibility of the CRY1-binding surface.

Phosphorylation also increased the distance between BMAL1 PAS-A domain and the nucleosomal acidic patch, composed of H2A (Glu^61^, Asp^90^ and Glu^92^) and H2B (Glu^105^ and His^109^) (Supplementary Figure 12). Given that the acidic patch serves as a major docking platform for numerous chromatin-associated factors, these findings suggest that phosphorylation may indirectly influence chromatin factor accessibility and nucleosome engagement by altering the positioning of CLOCK/BMAL1 relative to the nucleosome surface. Consistent with these large-scale conformational changes, phosphorylation also increased the structural fluctuations of histones H3 and H2A/H2B, as revealed by root mean square deviation (RMSD) analysis (Supplementary Figure 13-15). Together, global conformational dynamics analysis indicates that phosphorylation not only modulates local interaction networks within the CLOCK/BMAL1 complex but also propagates structural perturbations throughout the nucleosome-bound assembly.

## Conclusion and Discussion

Regulation of the circadian rhythm is a highly complex process in which post-translational modifications play critical roles. Among these, phosphorylation exerts a particularly profound influence. Numerous biochemical studies have demonstrated that phosphorylation modulates the binding of transcription factors to DNA.^45,46^ However, despite offering valuable insights into binding regulation, these studies largely lack atomistic-level mechanistic detail. Moreover, the potential contribution of histones to this process has remained unexplored. This is notable given that the cryo-EM structure of the CLOCK/BMAL1 complex reveals close spatial proximity to histones, suggesting an additional and previously underappreciated layer of regulatory control. Motivated by these unresolved questions, we conducted coarse grained simulations of the CLOCK/BMAL1-nucleosome complex derived from cryo-EM data, incorporating phosphorylation states, which are identified by experimental studies probing PER-dependent repression of CLOCK–BMAL1 transcriptional activity.

Our analyses reveal that phosphorylation induces coordinated changes across multiple structural layers of the assembly, including DNA engagement, histone interactions, interdomain communication, and long-range allosteric signaling networks. One of the most striking findings of this study is the asymmetric nature of phosphorylation-dependent regulation between CLOCK and BMAL1. Although the phosphorylation sites examined here occupy structurally analogous positions within the bHLH domains, they produce markedly distinct dynamical outcomes. CLOCK phosphorylation largely preserves the bimodal E-box binding behavior observed in the unphosphorylated complex, while BMAL1 phosphorylation substantially redistributes DNA-contact populations and promotes a more compact DNA-engaged conformation at CLOCK bHLH. Importantly, our simulations further suggest that BMAL1 phosphorylation acts in a dominant manner within the heterodimer, reshaping not only BMAL1 dynamics itself but also the conformational behavior and residue-correlation networks of CLOCK.

Beyond direct modulation of DNA interactions, our results indicate that phosphorylation propagates structural perturbations throughout the nucleosome-bound CLOCK-BMAL1 complex. Phosphomimetic substitutions altered contacts between CLOCK domains and histones, reshaped histone–DNA interaction states, and increased fluctuations within H2A/H2B and H3 histones. These observations are particularly intriguing considering recent 8OSK structure, in which the H3 α1–L1 region contacts the CLOCK PAS-B HI loop, a surface that also constitutes an established binding interface for CRY proteins.^12^ The phosphorylation-dependent increase in separation between the CLOCK HI loop and the H3 α1–L1 elbow observed in our simulations suggests that phosphorylation may remodel the structural accessibility of the CRY interaction interface. Motivated by this observation, we constructed a structural model of the CRY1–CLOCK–BMAL1– nucleosome complex. Consistent with previous biochemical and structural studies,^21,22^ the model positions the CRY1 secondary pocket in contact with the CLOCK HI loop, while the photolyase homology region is oriented toward the nucleosomal DNA. Although the ability of CRY proteins to directly engage DNA remains controversial, our model is compatible with previous reports of sequence-independent proposed CRY1–DNA interactions.^44,47^ Given the limited global confidence of the multicomponent prediction, we interpret this model as demonstrating the structural feasibility of such an arrangement rather than defining the architecture of the repressive complex. Furthermore, recent studies have suggested that, during displacement-type repression in the mouse liver, CRY and PER are recruited to chromatin-bound CLOCK–BMAL1, where they recruit casein kinases before phosphorylation-dependent dissociation of the complex from DNA.^17,48^ In this context, our results suggest a potentially complementary structural role for phosphorylation before or during CRY association: phosphorylation-dependent remodeling of the CLOCK–nucleosome interface may shift the conformational ensemble toward states in which the HI loop is less constrained by the histone surface and therefore more accessible to CRY1. This model does not imply that CLOCK^S38/S42^ phosphorylation is required for CRY recruitment; rather, it suggests that such phosphorylation-dependent remodeling may facilitate the structural accessibility of an established CRY-binding interface and thereby contribute to the transition from transcriptional activation toward repression, possibly via displacement-type.

Previous studies investigating the allosteric regulation of CRY proteins have demonstrated that CRY1 possesses extensive long-range allosteric communication, whereas CRY2 lacks comparable allosteric coupling.^22,49^ To our knowledge, however, the allosteric communication networks of CLOCK and BMAL1 have not previously been investigated. Here, we showed that such allosteric regulation exists both on BMAL1 and CLOCK. Our analysis demonstrates that phosphorylation also impacts the communication network within the CLOCK/BMAL1 heterodimer. Residue-correlation and allosteric pathway analyses revealed that phosphorylation disrupts long-range coupling between DNA-binding and regulatory PAS domains while simultaneously redistributing communication flow toward domain-specific subnetworks. Interestingly, both CLOCK and BMAL1 phosphorylation altered interaction between bHLH and PAS domains as well as PAS-A and PAS-B domains. Although these effects require experimental validation, they raise the possibility that phosphorylation of CLOCK and BMAL1 serves partially specialized regulatory functions within the heterodimer: one predominantly modulating allosteric signal propagation and conformational coordination, and the other more directly influencing chromatin engagement.

The PAS domains of CLOCK and BMAL1 have recently emerged as attractive targets for pharmacological intervention.^50–52^ Small molecules targeting CRY proteins, REV-ERBs, and casein kinases have already demonstrated the therapeutic potential against metabolic disease, cancer, inflammation, and sleep disorders.^50,53–55^ More recently, ligands targeting the CLOCK PAS-A domain have been shown to suppress CLOCK/BMAL1 transcriptional activity, further highlighting the functional importance of PAS-domain conformational dynamics.^56^ In this context, our proposed CRY1–CLOCK–BMAL1–nucleosome model suggests that phosphorylation may function as a structural priming event that facilitates subsequent CRY recruitment to the chromatin-bound complex by increasing the conformational space surrounding the CLOCK HI loop. Although this model requires experimental validation, it establishes a testable mechanistic framework linking phosphorylation-dependent conformational remodeling to CRY-mediated transcriptional repression.

Collectively, our findings demonstrate that phosphorylation remodels the CLOCK–BMAL1–nucleosome complex through coordinated changes in DNA binding, nucleosome interactions, and long-range allosteric communication. By integrating these multiscale structural effects into a unified mechanistic model, this work advances our understanding of chromatin-dependent circadian transcription and provides a foundation for the rational design of therapeutics targeting the molecular circadian clock.

## Data Availability Statement

Softwares used in the study are owned by their respective developers. The trajectories pertaining to unphosphorylated and phosphorylated systems are deposited in the OSF repository, which can be accessed via the link: https://osf.io/m7q4k/overview?view_only=3e66d7ab41a14745be173c726a14e46a

## Supporting Information

The model pertaining to CRY1-bound CLOCK-BMAL1-nucleosome complex, timeline plots of contacts numbers between the E-box and bHLH domains of CLOCK and BMAL1, the fork angle, contacts between histones and CLOCK (bHLH and PAS-B), and BMAL1 (PAS-A), as well as DNA contacts, for five replicates of each system. Representative conformations obtained from PCA, and RMSDs of histones H2A/H2B and H3 are also provided.

## Acknowledgements

This work was supported by Research Fund of Istanbul University (FYL-2022-38660; FYL-2023-40057; FYL-2023-40061).

## Author Contributions

Ş.G and O.S designed the study. D.A and H.P performed modeling, molecular dynamics simulations, and analyses of the trajectories. D.A and H.P wrote the first draft, and all the authors finalized the manuscript.

## Notes

The authors declare no competing financial interest.

