## Supplementary File v2 for "Phosphorylation-dependent Remodeling of the CLOCK–BMAL1–nucleosome Complex"

**CLOCK–BMAL1–nucleosome Complex**

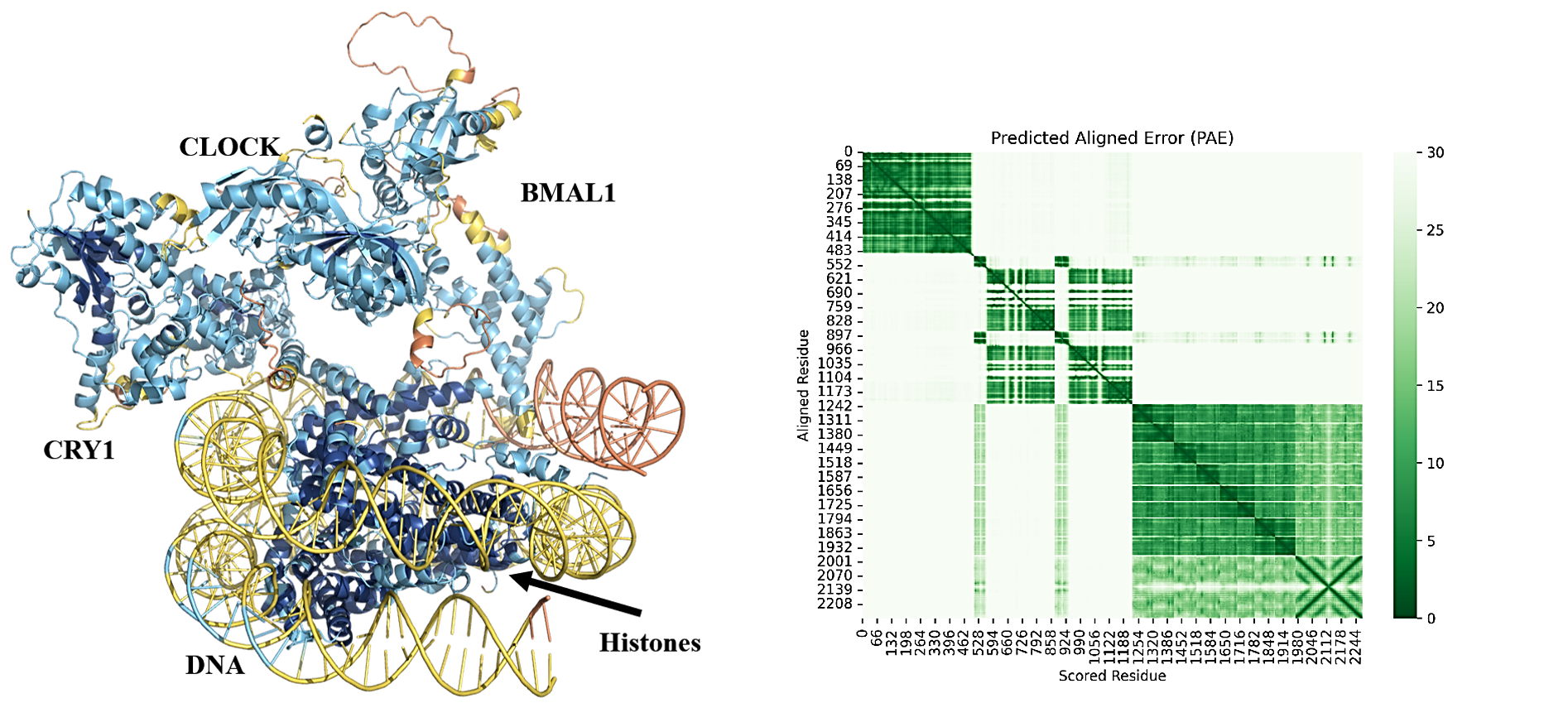

**Figure S1**: **AlphaFold model and prediction confidence of the CRY1–CLOCK–BMAL1–nucleosome complex.** The highest-ranked model had pTM and ipTM scores of 0.48 and 0.46, respectively. The model is colored according to predicted local distance difference test (pLDDT) scores, with blue regions representing higher and yellow-to-orange regions representing lower local prediction confidence. The corresponding predicted aligned error (PAE) plot is shown to illustrate confidence in the relative positioning of the components. Although the global confidence metrics indicate uncertainty in the overall architecture of the multicomponent complex, the model provides a structural framework for examining the predicted positioning of CRY1 relative to CLOCK–BMAL1 and nucleosomal DNA.

DNA"E-box" - CLOCK_bHLH (nonphosphorylated)

Number of contacts

| Frame | 1st | 2nd | 3rd | 4th | 5th |
| --- | --- | --- | --- | --- | --- |
| 0 | 16 | 16 | 6 | 11 | 21 |
| 0.1 | 16 | 16 | 6 | 11 | 16 |
| 0.2 | 16 | 16 | 6 | 17 | 17 |
| 0.3 | 17 | 16 | 7 | 14 | 12 |
| 0.4 | 17 | 14 | 6 | 16 | 17 |
| 0.5 | 14 | 16 | 7 | 17 | 17 |
| 0.6 | 19 | 15 | 7 | 13 | 15 |
| 0.7 | 11 | 12 | 11 | 15 | 13 |
| 0.8 | 13 | 16 | 7 | 14 | 21 |
| 0.9 | 13 | 14 | 7 | 12 | 18 |
| 1 | 12 | 17 | 9 | 17 | 17 |
| 1.1 | 14 | 14 | 8 | 12 | 15 |
| 1.2 | 15 | 17 | 11 | 13 | 14 |
| 1.3 | 17 | 17 | 7 | 5 | 17 |
| 1.4 | 17 | 17 | 11 | 9 | 18 |
| 1.5 | 20 | 15 | 12 | 11 | 14 |
| 1.6 | 21 | 19 | 11 | 13 | 17 |
| 1.7 | 27 | 18 | 9 | 12 | 15 |
| 1.8 | 29 | 16 | 13 | 9 | 15 |
| 1.9 | 26 | 21 | 11 | 8 | 18 |
| 2 | 26 | 20 | 13 | 8 | 15 |
| 2.1 | 28 | 16 | 11 | 9 | 24 |
| 2.2 | 28 | 13 | 15 | 5 | 22 |
| 2.3 | 24 | 16 | 13 | 12 | 21 |
| 2.4 | 29 | 15 | 10 | 12 | 19 |
| 2.5 | 28 | 17 | 11 | 12 | 22 |
| 2.6 | 30 | 16 | 14 | 9 | 21 |
| 2.7 | 18 | 19 | 12 | 13 | 20 |
| 2.8 | 19 | 13 | 12 | 12 | 25 |
| 2.9 | 13 | 18 | 11 | 13 | 18 |
| 3 | 35 | 12 | 10 | 10 | 20 |
| 3.1 | 37 | 17 | 12 | 9 | 22 |
| 3.2 | 26 | 17 | 11 | 8 | 22 |
| 3.3 | 36 | 16 | 12 | 8 | 27 |
| 3.4 | 31 | 14 | 12 | 5 | 21 |
| 3.5 | 31 | 16 | 9 | 6 | 25 |
| 3.6 | 29 | 18 | 12 | 7 | 30 |
| 3.7 | 32 | 17 | 12 | 7 | 28 |
| 3.8 | 34 | 17 | 13 | 7 | 17 |
| 3.9 | 35 | 15 | 18 | 9 | 20 |
| 4 | 34 | 19 | 13 | 11 | 24 |
| 4.1 | 35 | 18 | 13 | 7 | 19 |
| 4.2 | 30 | 18 | 10 | 9 | 18 |
| 4.3 | 32 | 18 | 16 | 6 | 24 |
| 4.4 | 29 | 15 | 11 | 11 | 30 |
| 4.5 | 26 | 16 | 11 | 7 | 33 |
| 4.6 | 31 | 16 | 8 | 11 | 26 |
| 4.7 | 27 | 17 | 8 | 10 | 30 |
| 4.8 | 40 | 15 | 6 | 7 | 29 |
| 4.9 | 39 | 16 | 8 | 9 | 32 |
| 5 | 34 | 15 | 8 | 8 | 27 |
| 5.1 | 28 | 12 | 12 | 10 | 29 |
| 5.2 | 43 | 14 | 10 | 9 | 27 |
| 5.3 | 34 | 17 | 8 | 4 | 29 |
| 5.4 | 29 | 13 | 9 | 4 | 33 |
| 5.5 | 41 | 17 | 6 | 7 | 28 |
| 5.6 | 38 | 12 | 9 | 12 | 31 |
| 5.7 | 39 | 15 | 9 | 11 | 32 |
| 5.8 | 33 | 26 | 10 | 6 | 28 |
| 5.9 | 28 | 17 | 9 | 9 | 28 |
| 6 | 28 | 24 | 6 | 5 | 32 |
| 6.1 | 36 | 19 | 8 | 4 | 27 |
| 6.2 | 25 | 37 | 9 | 8 | 29 |
| 6.3 | 38 | 32 | 10 | 9 | 30 |
| 6.4 | 35 | 36 | 10 | 9 | 32 |
| 6.5 | 38 | 33 | 12 | 9 | 33 |
| 6.6 | 29 | 33 | 11 | 7 | 33 |
| 6.7 | 40 | 27 | 8 | 8 | 30 |
| 6.8 | 39 | 34 | 10 | 9 | 34 |
| 6.9 | 34 | 41 | 11 | 8 | 27 |
| 7 | 40 | 32 | 10 | 7 | 31 |
| 7.1 | 35 | 28 | 8 | 6 | 33 |
| 7.2 | 30 | 31 | 10 | 5 | 35 |
| 7.3 | 44 | 41 | 8 | 6 | 31 |
| 7.4 | 40 | 37 | 11 | 12 | 32 |
| 7.5 | 38 | 33 | 12 | 6 | 29 |
| 7.6 | 35 | 38 | 10 | 6 | 27 |
| 7.7 | 31 | 36 | 10 | 6 | 29 |
| 7.8 | 39 | 36 | 11 | 7 | 30 |
| 7.9 | 31 | 37 | 5 | 6 | 27 |
| 8 | 40 | 33 | 11 | 6 | 25 |
| 8.1 | 31 | 36 | 10 | 6 | 28 |
| 8.2 | 35 | 30 | 13 | 7 | 23 |
| 8.3 | 34 | 32 | 15 | 7 | 26 |
| 8.4 | 25 | 35 | 10 | 4 | 23 |
| 8.5 | 27 | 32 | 10 | 12 | 28 |
| 8.6 | 29 | 33 | 11 | 7 | 28 |
| 8.7 | 30 | 32 | 10 | 8 | 26 |
| 8.8 | 30 | 31 | 10 | 6 | 28 |
| 8.9 | 27 | 36 | 8 | 10 | 26 |
| 9 | 30 | 33 | 8 | 9 | 30 |
| 9.1 | 33 | 31 | 11 | 10 | 29 |
| 9.2 | 35 | 32 | 13 | 8 | 27 |
| 9.3 | 24 | 33 | 11 | 8 | 29 |
| 9.4 | 34 | 32 | 11 | 6 | 28 |
| 9.5 | 26 | 32 | 10 | 9 | 28 |
| 9.6 | 31 | 30 | 12 | 8 | 25 |
| 9.7 | 26 | 31 | 8 | 8 | 30 |
| 9.8 | 34 | 26 | 10 | 5 | 30 |
| 9.9 | 28 | 25 | 10 | 7 | 23 |
| 10 | 23 | 30 | 13 | 6 | 26 |
| 10.1 | 35 | 30 | 7 | 6 | 28 |
| 10.2 | 38 | 31 | 10 | 7 | 25 |
| 10.3 | 29 | 32 | 11 | 6 | 32 |
| 10.4 | 30 | 16 | 12 | 8 | 26 |
| 10.5 | 27 | 36 | 10 | 7 | 31 |
| 10.6 | 28 | 35 | 11 | 5 | 27 |
| 10.7 | 33 | 27 | 10 | 9 | 24 |
| 10.8 | 30 | 31 | 9 | 8 | 28 |
| 10.9 | 31 | 29 | 12 | 9 | 28 |
| 11 | 29 | 25 | 11 | 6 | 27 |
| 11.1 | 30 | 28 | 9 | 8 | 30 |
| 11.2 | 34 | 33 | 9 | 7 | 29 |
| 11.3 | 33 | 36 | 11 | 9 | 29 |
| 11.4 | 34 | 28 | 13 | 12 | 30 |
| 11.5 | 33 | 28 | 11 | 8 | 27 |
| 11.6 | 26 | 34 | 12 | 8 | 31 |
| 11.7 | 32 | 32 | 14 | 8 | 31 |
| 11.8 | 33 | 28 | 11 | 9 | 27 |
| 11.9 | 30 | 33 | 9 | 12 | 33 |
| 12 | 24 | 36 | 11 | 6 | 31 |
| 12.1 | 28 | 35 | 10 | 9 | 26 |
| 12.2 | 34 | 35 | 13 | 9 | 33 |
| 12.3 | 25 | 31 | 13 | 6 | 28 |
| 12.4 | 33 | 32 | 11 | 9 | 27 |
| 12.5 | 30 | 35 | 11 | 6 | 27 |
| 12.6 | 25 | 26 | 9 | 7 | 28 |
| 12.7 | 28 | 31 | 12 | 7 | 30 |
| 12.8 | 28 | 39 | 9 | 7 | 29 |
| 12.9 | 23 | 31 | 9 | 10 | 31 |
| 13 | 30 | 32 | 11 | 7 | 30 |
| 13.1 | 24 | 29 | 11 | 8 | 25 |
| 13.2 | 31 | 33 | 10 | 7 | 26 |
| 13.3 | 30 | 34 | 6 | 10 | 31 |
| 13.4 | 33 | 32 | 8 | 9 | 36 |
| 13.5 | 26 | 29 | 8 | 7 | 26 |
| 13.6 | 22 | 33 | 11 | 6 | 31 |
| 13.7 | 29 | 32 | 7 | 8 | 31 |
| 13.8 | 19 | 29 | 7 | 10 | 28 |
| 13.9 | 16 | 37 | 7 | 8 | 30 |
| 14 | 33 | 34 | 5 | 8 | 33 |
| 14.1 | 32 | 29 | 9 | 7 | 30 |
| 14.2 | 29 | 29 | 6 | 6 | 30 |
| 14.3 | 31 | 41 | 8 | 7 | 31 |
| 14.4 | 32 | 39 | 6 | 6 | 30 |
| 14.5 | 33 | 36 | 7 | 6 | 31 |
| 14.6 | 30 | 33 | 7 | 7 | 32 |
| 14.7 | 29 | 31 | 6 | 6 | 27 |
| 14.8 | 33 | 36 | 7 | 7 | 26 |
| 14.9 | 27 | 38 | 11 | 5 | 30 |
| 15 | 33 | 41 | 7 | 5 | 30 |
| 15.1 | 34 | 29 | 7 | 6 | 28 |
| 15.2 | 27 | 33 | 8 | 6 | 28 |
| 15.3 | 25 | 28 | 7 | 7 | 30 |
| 15.4 | 32 | 39 | 7 | 6 | 31 |
| 15.5 | 27 | 36 | 8 | 7 | 28 |
| 15.6 | 29 | 28 | 8 | 7 | 32 |
| 15.7 | 35 | 28 | 8 | 6 | 27 |
| 15.8 | 43 | 34 | 8 | 7 | 31 |
| 15.9 | 30 | 35 | 8 | 7 | 28 |
| 16 | 27 | 33 | 7 | 6 | 34 |
| 16.1 | 29 | 32 | 6 | 10 | 30 |
| 16.2 | 33 | 36 | 9 | 8 | 31 |
| 16.3 | 37 | 33 | 11 | 7 | 28 |
| 16.4 | 32 | 32 | 8 | 6 | 29 |
| 16.5 | 34 | 31 | 8 | 8 | 30 |
| 16.6 | 29 | 35 | 8 | 5 | 27 |
| 16.7 | 28 | 31 | 7 | 5 | 28 |
| 16.8 | 37 | 37 | 10 | 7 | 32 |
| 16.9 | 30 | 31 | 7 | 6 | 30 |
| 17 | 34 | 39 | 7 | 5 | 33 |
| 17.1 | 24 | 37 | 7 | 8 | 31 |
| 17.2 | 24 | 38 | 6 | 8 | 31 |
| 17.3 | 23 | 37 | 8 | 9 | 31 |
| 17.4 | 30 | 33 | 6 | 10 | 33 |
| 17.5 | 31 | 32 | 6 | 11 | 33 |
| 17.6 | 29 | 34 | 6 | 10 | 29 |
| 17.7 | 32 | 41 | 9 | 9 | 27 |
| 17.8 | 31 | 32 | 9 | 9 | 33 |
| 17.9 | 31 | 34 | 7 | 12 | 34 |
| 18 | 27 | 28 | 6 | 9 | 31 |
| 18.1 | 33 | 29 | 6 | 10 | 28 |
| 18.2 | 22 | 28 | 7 | 11 | 30 |
| 18.3 | 33 | 30 | 6 | 12 | 27 |
| 18.4 | 27 | 29 | 6 | 8 | 30 |
| 18.5 | 30 | 34 | 5 | 8 | 29 |
| 18.6 | 29 | 33 | 6 | 10 | 29 |
| 18.7 | 25 | 27 | 6 | 8 | 30 |
| 18.8 | 32 | 28 | 6 | 9 | 29 |
| 18.9 | 30 | 25 | 8 | 8 | 32 |
| 19 | 24 | 31 | 7 | 5 | 28 |
| 19.1 | 36 | 31 | 6 | 7 | 33 |
| 19.2 | 25 | 29 | 6 | 10 | 34 |
| 19.3 | 29 | 31 | 6 | 7 | 33 |
| 19.4 | 32 | 36 | 10 | 8 | 32 |
| 19.5 | 37 | 30 | 7 | 10 | 33 |
| 19.6 | 36 | 29 | 6 | 9 | 35 |
| 19.7 | 23 | 31 | 7 | 6 | 32 |
| 19.8 | 26 | 39 | 7 | 5 | 32 |
| 19.9 | 24 | 29 | 7 | 9 | 34 |

DNA"E-box" - CLOCK_bHLH (BMAL1S78)

Number of contacts

| Frame | 1st | 2nd | 3rd | 4th | 5th |
| --- | --- | --- | --- | --- | --- |
| 0 | 23 | 14 | 27 | 22 | 36 |
| 0.1 | 22 | 15 | 27 | 26 | 38 |
| 0.2 | 26 | 27 | 24 | 40 | 38 |
| 0.3 | 47 | 27 | 32 | 38 | 37 |
| 0.4 | 45 | 27 | 45 | 53 | 43 |
| 0.5 | 43 | 24 | 30 | 50 | 50 |
| 0.6 | 49 | 30 | 23 | 54 | 50 |
| 0.7 | 38 | 29 | 22 | 48 | 39 |
| 0.8 | 51 | 28 | 26 | 60 | 37 |
| 0.9 | 44 | 29 | 34 | 53 | 48 |
| 1 | 44 | 26 | 51 | 56 | 41 |
| 1.1 | 43 | 31 | 54 | 52 | 41 |
| 1.2 | 44 | 27 | 51 | 64 | 48 |
| 1.3 | 51 | 28 | 44 | 63 | 35 |
| 1.4 | 47 | 26 | 47 | 62 | 42 |
| 1.5 | 46 | 29 | 55 | 45 | 47 |
| 1.6 | 50 | 27 | 49 | 64 | 36 |
| 1.7 | 42 | 26 | 47 | 63 | 34 |
| 1.8 | 39 | 31 | 49 | 55 | 44 |
| 1.9 | 47 | 26 | 43 | 61 | 30 |
| 2 | 47 | 26 | 52 | 60 | 45 |
| 2.1 | 35 | 26 | 44 | 59 | 44 |
| 2.2 | 35 | 30 | 46 | 53 | 54 |
| 2.3 | 39 | 35 | 50 | 50 | 30 |
| 2.4 | 43 | 30 | 43 | 52 | 48 |
| 2.5 | 49 | 35 | 47 | 56 | 40 |
| 2.6 | 39 | 31 | 56 | 52 | 49 |
| 2.7 | 42 | 38 | 45 | 53 | 38 |
| 2.8 | 41 | 41 | 50 | 54 | 49 |
| 2.9 | 43 | 29 | 46 | 51 | 43 |
| 3 | 44 | 31 | 53 | 56 | 42 |
| 3.1 | 53 | 36 | 53 | 54 | 60 |
| 3.2 | 52 | 32 | 47 | 55 | 40 |
| 3.3 | 50 | 39 | 51 | 51 | 46 |
| 3.4 | 47 | 29 | 41 | 54 | 45 |
| 3.5 | 45 | 31 | 50 | 53 | 43 |
| 3.6 | 42 | 38 | 45 | 53 | 42 |
| 3.7 | 38 | 33 | 36 | 55 | 50 |
| 3.8 | 43 | 35 | 48 | 55 | 48 |
| 3.9 | 45 | 35 | 50 | 55 | 43 |
| 4 | 42 | 32 | 56 | 59 | 50 |
| 4.1 | 39 | 37 | 46 | 53 | 34 |
| 4.2 | 46 | 48 | 46 | 55 | 40 |
| 4.3 | 42 | 40 | 49 | 52 | 46 |
| 4.4 | 40 | 42 | 53 | 53 | 40 |
| 4.5 | 41 | 45 | 48 | 55 | 46 |
| 4.6 | 51 | 47 | 40 | 52 | 36 |
| 4.7 | 42 | 45 | 52 | 56 | 40 |
| 4.8 | 44 | 44 | 47 | 57 | 36 |
| 4.9 | 40 | 49 | 47 | 49 | 47 |
| 5 | 46 | 46 | 41 | 53 | 47 |
| 5.1 | 49 | 43 | 45 | 54 | 43 |
| 5.2 | 45 | 32 | 51 | 52 | 46 |
| 5.3 | 48 | 33 | 57 | 48 | 45 |
| 5.4 | 48 | 46 | 47 | 51 | 39 |
| 5.5 | 57 | 45 | 48 | 50 | 38 |
| 5.6 | 57 | 54 | 45 | 49 | 48 |
| 5.7 | 48 | 43 | 51 | 48 | 37 |
| 5.8 | 60 | 56 | 46 | 50 | 43 |
| 5.9 | 53 | 44 | 51 | 63 | 43 |
| 6 | 59 | 49 | 40 | 53 | 42 |
| 6.1 | 46 | 53 | 48 | 54 | 47 |
| 6.2 | 40 | 46 | 44 | 54 | 38 |
| 6.3 | 44 | 31 | 45 | 52 | 42 |
| 6.4 | 53 | 31 | 43 | 54 | 41 |
| 6.5 | 43 | 30 | 46 | 44 | 33 |
| 6.6 | 52 | 33 | 49 | 53 | 51 |
| 6.7 | 50 | 33 | 47 | 53 | 47 |
| 6.8 | 51 | 38 | 53 | 57 | 42 |
| 6.9 | 56 | 47 | 44 | 54 | 44 |
| 7 | 53 | 67 | 40 | 50 | 45 |
| 7.1 | 55 | 59 | 49 | 52 | 41 |
| 7.2 | 47 | 64 | 49 | 56 | 39 |
| 7.3 | 44 | 60 | 55 | 49 | 40 |
| 7.4 | 51 | 63 | 50 | 57 | 42 |
| 7.5 | 47 | 66 | 56 | 57 | 45 |
| 7.6 | 41 | 66 | 39 | 51 | 43 |
| 7.7 | 51 | 68 | 48 | 49 | 43 |
| 7.8 | 52 | 61 | 44 | 53 | 40 |
| 7.9 | 40 | 67 | 45 | 55 | 44 |
| 8 | 46 | 61 | 52 | 57 | 44 |
| 8.1 | 44 | 55 | 55 | 56 | 40 |
| 8.2 | 40 | 70 | 37 | 56 | 41 |
| 8.3 | 47 | 62 | 51 | 54 | 44 |
| 8.4 | 51 | 61 | 50 | 51 | 41 |
| 8.5 | 47 | 61 | 42 | 57 | 41 |
| 8.6 | 46 | 66 | 48 | 55 | 44 |
| 8.7 | 47 | 66 | 50 | 58 | 45 |
| 8.8 | 44 | 62 | 46 | 53 | 50 |
| 8.9 | 51 | 66 | 51 | 46 | 44 |
| 9 | 48 | 58 | 52 | 48 | 36 |
| 9.1 | 49 | 59 | 45 | 45 | 42 |
| 9.2 | 50 | 59 | 46 | 57 | 40 |
| 9.3 | 55 | 60 | 47 | 53 | 34 |
| 9.4 | 45 | 63 | 50 | 53 | 32 |
| 9.5 | 40 | 57 | 51 | 48 | 35 |
| 9.6 | 46 | 65 | 52 | 52 | 31 |
| 9.7 | 47 | 68 | 44 | 51 | 48 |
| 9.8 | 39 | 57 | 47 | 46 | 35 |
| 9.9 | 55 | 62 | 44 | 50 | 37 |
| 10 | 46 | 58 | 52 | 47 | 46 |
| 10.1 | 51 | 54 | 42 | 50 | 42 |
| 10.2 | 46 | 60 | 49 | 52 | 40 |
| 10.3 | 42 | 64 | 51 | 49 | 43 |
| 10.4 | 43 | 62 | 46 | 53 | 40 |
| 10.5 | 57 | 60 | 47 | 57 | 35 |
| 10.6 | 52 | 58 | 50 | 46 | 43 |
| 10.7 | 52 | 63 | 46 | 49 | 37 |
| 10.8 | 40 | 66 | 43 | 43 | 43 |
| 10.9 | 46 | 57 | 54 | 43 | 36 |
| 11 | 41 | 63 | 41 | 55 | 41 |
| 11.1 | 41 | 69 | 49 | 49 | 39 |
| 11.2 | 41 | 65 | 51 | 45 | 42 |
| 11.3 | 41 | 68 | 52 | 33 | 40 |
| 11.4 | 46 | 60 | 50 | 37 | 50 |
| 11.5 | 42 | 59 | 44 | 32 | 42 |
| 11.6 | 53 | 65 | 42 | 34 | 46 |
| 11.7 | 44 | 66 | 45 | 32 | 45 |
| 11.8 | 46 | 63 | 49 | 35 | 45 |
| 11.9 | 50 | 64 | 53 | 35 | 46 |
| 12 | 50 | 63 | 55 | 36 | 42 |
| 12.1 | 48 | 63 | 52 | 39 | 39 |
| 12.2 | 50 | 68 | 45 | 33 | 43 |
| 12.3 | 43 | 64 | 57 | 27 | 44 |
| 12.4 | 47 | 59 | 48 | 37 | 41 |
| 12.5 | 47 | 64 | 42 | 30 | 46 |
| 12.6 | 50 | 61 | 49 | 37 | 45 |
| 12.7 | 49 | 62 | 42 | 34 | 36 |
| 12.8 | 46 | 65 | 51 | 32 | 36 |
| 12.9 | 44 | 65 | 43 | 35 | 43 |
| 13 | 40 | 56 | 48 | 30 | 49 |
| 13.1 | 42 | 64 | 51 | 32 | 47 |
| 13.2 | 48 | 66 | 43 | 31 | 48 |
| 13.3 | 48 | 56 | 52 | 27 | 41 |
| 13.4 | 47 | 61 | 49 | 35 | 43 |
| 13.5 | 44 | 65 | 46 | 31 | 42 |
| 13.6 | 52 | 64 | 42 | 35 | 38 |
| 13.7 | 50 | 67 | 48 | 30 | 41 |
| 13.8 | 52 | 69 | 45 | 42 | 40 |
| 13.9 | 50 | 55 | 47 | 37 | 45 |
| 14 | 51 | 59 | 44 | 39 | 37 |
| 14.1 | 43 | 62 | 43 | 37 | 39 |
| 14.2 | 44 | 72 | 51 | 37 | 43 |
| 14.3 | 44 | 69 | 47 | 39 | 47 |
| 14.4 | 48 | 63 | 49 | 35 | 50 |
| 14.5 | 48 | 62 | 47 | 39 | 47 |
| 14.6 | 53 | 66 | 50 | 46 | 43 |
| 14.7 | 46 | 62 | 40 | 46 | 48 |
| 14.8 | 50 | 64 | 47 | 44 | 40 |
| 14.9 | 47 | 63 | 45 | 45 | 31 |
| 15 | 40 | 64 | 53 | 34 | 47 |
| 15.1 | 44 | 61 | 49 | 38 | 39 |
| 15.2 | 45 | 59 | 51 | 41 | 38 |
| 15.3 | 63 | 61 | 47 | 37 | 43 |
| 15.4 | 46 | 52 | 53 | 36 | 47 |
| 15.5 | 44 | 64 | 40 | 39 | 42 |
| 15.6 | 44 | 64 | 47 | 31 | 51 |
| 15.7 | 45 | 53 | 43 | 40 | 38 |
| 15.8 | 37 | 64 | 42 | 44 | 47 |
| 15.9 | 49 | 64 | 58 | 38 | 47 |
| 16 | 42 | 59 | 48 | 38 | 42 |
| 16.1 | 50 | 61 | 48 | 36 | 38 |
| 16.2 | 44 | 59 | 47 | 34 | 41 |
| 16.3 | 53 | 63 | 54 | 36 | 44 |
| 16.4 | 54 | 62 | 45 | 33 | 48 |
| 16.5 | 47 | 64 | 46 | 38 | 42 |
| 16.6 | 41 | 69 | 42 | 39 | 38 |
| 16.7 | 50 | 61 | 51 | 39 | 48 |
| 16.8 | 45 | 58 | 50 | 46 | 38 |
| 16.9 | 38 | 55 | 51 | 41 | 45 |
| 17 | 47 | 61 | 48 | 35 | 39 |
| 17.1 | 54 | 66 | 43 | 39 | 51 |
| 17.2 | 44 | 57 | 49 | 31 | 49 |
| 17.3 | 44 | 55 | 48 | 36 | 47 |
| 17.4 | 45 | 52 | 48 | 41 | 37 |
| 17.5 | 54 | 64 | 51 | 42 | 48 |
| 17.6 | 42 | 65 | 41 | 45 | 49 |
| 17.7 | 42 | 61 | 46 | 44 | 43 |
| 17.8 | 47 | 65 | 51 | 30 | 38 |
| 17.9 | 44 | 51 | 48 | 31 | 40 |
| 18 | 46 | 66 | 52 | 33 | 45 |
| 18.1 | 48 | 61 | 46 | 36 | 42 |
| 18.2 | 50 | 62 | 54 | 33 | 39 |
| 18.3 | 62 | 64 | 47 | 40 | 37 |
| 18.4 | 52 | 63 | 45 | 36 | 43 |
| 18.5 | 64 | 59 | 48 | 35 | 36 |
| 18.6 | 63 | 60 | 53 | 38 | 42 |
| 18.7 | 52 | 62 | 48 | 38 | 40 |
| 18.8 | 60 | 58 | 52 | 33 | 45 |
| 18.9 | 60 | 61 | 47 | 37 | 38 |
| 19 | 64 | 63 | 47 | 37 | 39 |
| 19.1 | 54 | 56 | 52 | 35 | 47 |
| 19.2 | 48 | 59 | 42 | 36 | 51 |
| 19.3 | 59 | 63 | 48 | 27 | 52 |
| 19.4 | 51 | 64 | 53 | 38 | 42 |
| 19.5 | 66 | 62 | 51 | 40 | 43 |
| 19.6 | 57 | 64 | 46 | 33 | 49 |
| 19.7 | 57 | 67 | 51 | 33 | 44 |
| 19.8 | 50 | 62 | 48 | 35 | 45 |
| 19.9 | 56 | 61 | 47 | 45 | 45 |

DNA"E-box" - CLOCK_bHLH (CLOCKS38/42)

Number of contacts

| Frame | 1st | 2nd | 3rd | 4th | 5th |
| --- | --- | --- | --- | --- | --- |
| 0 | 20 | 4 | 23 | 22 | 16 |
| 0.1 | 22 | 4 | 23 | 22 | 17 |
| 0.2 | 19 | 3 | 24 | 27 | 18 |
| 0.3 | 18 | 6 | 24 | 27 | 17 |
| 0.4 | 9 | 7 | 30 | 35 | 33 |
| 0.5 | 20 | 9 | 24 | 32 | 33 |
| 0.6 | 20 | 6 | 33 | 29 | 47 |
| 0.7 | 16 | 5 | 27 | 33 | 38 |
| 0.8 | 14 | 5 | 29 | 28 | 31 |
| 0.9 | 23 | 6 | 28 | 34 | 28 |
| 1 | 15 | 6 | 24 | 32 | 33 |
| 1.1 | 10 | 8 | 28 | 22 | 21 |
| 1.2 | 4 | 5 | 31 | 35 | 25 |
| 1.3 | 4 | 6 | 25 | 26 | 15 |
| 1.4 | 4 | 7 | 26 | 31 | 31 |
| 1.5 | 4 | 8 | 31 | 28 | 30 |
| 1.6 | 4 | 5 | 26 | 26 | 24 |
| 1.7 | 4 | 9 | 33 | 26 | 32 |
| 1.8 | 5 | 7 | 27 | 24 | 36 |
| 1.9 | 6 | 6 | 29 | 29 | 31 |
| 2 | 5 | 6 | 24 | 32 | 25 |
| 2.1 | 6 | 6 | 23 | 28 | 32 |
| 2.2 | 6 | 6 | 23 | 29 | 27 |
| 2.3 | 5 | 7 | 19 | 27 | 23 |
| 2.4 | 5 | 7 | 20 | 24 | 35 |
| 2.5 | 6 | 9 | 28 | 27 | 42 |
| 2.6 | 5 | 8 | 24 | 34 | 36 |
| 2.7 | 5 | 9 | 33 | 29 | 36 |
| 2.8 | 5 | 6 | 21 | 18 | 33 |
| 2.9 | 6 | 8 | 27 | 24 | 34 |
| 3 | 7 | 6 | 24 | 29 | 33 |
| 3.1 | 3 | 8 | 27 | 29 | 31 |
| 3.2 | 1 | 10 | 26 | 30 | 32 |
| 3.3 | 5 | 8 | 29 | 23 | 36 |
| 3.4 | 10 | 7 | 33 | 20 | 35 |
| 3.5 | 8 | 8 | 20 | 20 | 36 |
| 3.6 | 6 | 9 | 29 | 17 | 35 |
| 3.7 | 7 | 8 | 28 | 17 | 35 |
| 3.8 | 6 | 6 | 20 | 22 | 36 |
| 3.9 | 8 | 8 | 30 | 20 | 35 |
| 4 | 8 | 6 | 22 | 21 | 32 |
| 4.1 | 8 | 8 | 22 | 18 | 35 |
| 4.2 | 8 | 8 | 25 | 20 | 34 |
| 4.3 | 6 | 8 | 19 | 21 | 31 |
| 4.4 | 6 | 6 | 24 | 19 | 31 |
| 4.5 | 8 | 7 | 25 | 16 | 34 |
| 4.6 | 6 | 7 | 18 | 24 | 35 |
| 4.7 | 6 | 8 | 16 | 19 | 39 |
| 4.8 | 4 | 7 | 27 | 23 | 47 |
| 4.9 | 5 | 7 | 25 | 25 | 49 |
| 5 | 4 | 8 | 27 | 23 | 53 |
| 5.1 | 5 | 8 | 25 | 26 | 47 |
| 5.2 | 6 | 7 | 27 | 23 | 40 |
| 5.3 | 6 | 5 | 32 | 20 | 52 |
| 5.4 | 8 | 8 | 21 | 27 | 48 |
| 5.5 | 5 | 8 | 30 | 26 | 47 |
| 5.6 | 4 | 7 | 28 | 23 | 49 |
| 5.7 | 6 | 7 | 23 | 25 | 47 |
| 5.8 | 3 | 7 | 28 | 27 | 52 |
| 5.9 | 5 | 6 | 26 | 20 | 47 |
| 6 | 6 | 7 | 27 | 25 | 44 |
| 6.1 | 6 | 4 | 23 | 20 | 45 |
| 6.2 | 6 | 6 | 29 | 19 | 53 |
| 6.3 | 6 | 5 | 27 | 21 | 52 |
| 6.4 | 6 | 8 | 21 | 20 | 43 |
| 6.5 | 5 | 6 | 24 | 21 | 49 |
| 6.6 | 5 | 8 | 26 | 22 | 53 |
| 6.7 | 4 | 5 | 26 | 20 | 38 |
| 6.8 | 5 | 6 | 25 | 18 | 43 |
| 6.9 | 4 | 7 | 27 | 20 | 42 |
| 7 | 5 | 9 | 31 | 21 | 47 |
| 7.1 | 6 | 4 | 27 | 20 | 45 |
| 7.2 | 4 | 5 | 31 | 23 | 46 |
| 7.3 | 5 | 2 | 32 | 23 | 47 |
| 7.4 | 6 | 1 | 30 | 18 | 33 |
| 7.5 | 5 | 9 | 25 | 14 | 39 |
| 7.6 | 5 | 9 | 29 | 20 | 42 |
| 7.7 | 4 | 8 | 31 | 23 | 49 |
| 7.8 | 5 | 10 | 27 | 18 | 49 |
| 7.9 | 6 | 9 | 26 | 19 | 41 |
| 8 | 5 | 11 | 31 | 21 | 49 |
| 8.1 | 5 | 9 | 30 | 18 | 46 |
| 8.2 | 3 | 8 | 31 | 16 | 57 |
| 8.3 | 5 | 7 | 31 | 19 | 42 |
| 8.4 | 7 | 4 | 30 | 19 | 40 |
| 8.5 | 8 | 3 | 32 | 18 | 51 |
| 8.6 | 7 | 6 | 28 | 18 | 52 |
| 8.7 | 6 | 10 | 33 | 22 | 39 |
| 8.8 | 5 | 4 | 34 | 20 | 46 |
| 8.9 | 5 | 3 | 22 | 23 | 48 |
| 9 | 5 | 4 | 27 | 21 | 43 |
| 9.1 | 6 | 4 | 25 | 21 | 51 |
| 9.2 | 4 | 3 | 25 | 26 | 45 |
| 9.3 | 6 | 4 | 30 | 22 | 42 |
| 9.4 | 4 | 2 | 32 | 24 | 45 |
| 9.5 | 3 | 3 | 36 | 25 | 44 |
| 9.6 | 2 | 3 | 32 | 23 | 53 |
| 9.7 | 3 | 3 | 34 | 23 | 51 |
| 9.8 | 2 | 6 | 31 | 26 | 49 |
| 9.9 | 4 | 5 | 33 | 16 | 52 |
| 10 | 4 | 5 | 31 | 23 | 47 |
| 10.1 | 10 | 3 | 31 | 21 | 42 |
| 10.2 | 10 | 3 | 32 | 23 | 50 |
| 10.3 | 8 | 3 | 34 | 25 | 46 |
| 10.4 | 4 | 2 | 39 | 24 | 46 |
| 10.5 | 1 | 3 | 32 | 26 | 48 |
| 10.6 | 1 | 3 | 32 | 27 | 53 |
| 10.7 | 0 | 4 | 32 | 28 | 46 |
| 10.8 | 0 | 3 | 31 | 28 | 53 |
| 10.9 | 1 | 3 | 32 | 26 | 48 |
| 11 | 0 | 4 | 36 | 26 | 46 |
| 11.1 | 0 | 3 | 35 | 25 | 37 |
| 11.2 | 0 | 3 | 33 | 24 | 41 |
| 11.3 | 0 | 3 | 34 | 27 | 42 |
| 11.4 | 2 | 2 | 30 | 27 | 46 |
| 11.5 | 1 | 7 | 36 | 25 | 48 |
| 11.6 | 2 | 5 | 34 | 22 | 53 |
| 11.7 | 0 | 2 | 25 | 27 | 53 |
| 11.8 | 1 | 2 | 29 | 24 | 47 |
| 11.9 | 0 | 2 | 29 | 26 | 42 |
| 12 | 1 | 2 | 34 | 27 | 46 |
| 12.1 | 1 | 2 | 27 | 21 | 45 |
| 12.2 | 0 | 2 | 30 | 24 | 45 |
| 12.3 | 0 | 2 | 32 | 25 | 42 |
| 12.4 | 1 | 3 | 26 | 23 | 46 |
| 12.5 | 0 | 2 | 32 | 24 | 45 |
| 12.6 | 0 | 2 | 28 | 27 | 46 |
| 12.7 | 0 | 2 | 33 | 22 | 38 |
| 12.8 | 0 | 2 | 25 | 25 | 50 |
| 12.9 | 0 | 2 | 32 | 25 | 41 |
| 13 | 1 | 2 | 23 | 23 | 44 |
| 13.1 | 0 | 2 | 40 | 24 | 46 |
| 13.2 | 0 | 2 | 29 | 26 | 49 |
| 13.3 | 0 | 2 | 25 | 20 | 47 |
| 13.4 | 0 | 2 | 27 | 24 | 50 |
| 13.5 | 0 | 2 | 30 | 23 | 43 |
| 13.6 | 1 | 2 | 28 | 21 | 46 |
| 13.7 | 0 | 2 | 27 | 26 | 42 |
| 13.8 | 0 | 2 | 27 | 23 | 45 |
| 13.9 | 0 | 1 | 31 | 24 | 38 |
| 14 | 0 | 2 | 39 | 29 | 48 |
| 14.1 | 0 | 2 | 44 | 28 | 51 |
| 14.2 | 0 | 2 | 36 | 22 | 49 |
| 14.3 | 0 | 2 | 44 | 23 | 49 |
| 14.4 | 0 | 2 | 36 | 27 | 51 |
| 14.5 | 0 | 2 | 34 | 23 | 55 |
| 14.6 | 0 | 2 | 44 | 24 | 45 |
| 14.7 | 0 | 2 | 38 | 21 | 48 |
| 14.8 | 0 | 1 | 36 | 24 | 47 |
| 14.9 | 0 | 2 | 39 | 32 | 47 |
| 15 | 1 | 2 | 41 | 31 | 51 |
| 15.1 | 1 | 2 | 39 | 28 | 50 |
| 15.2 | 1 | 2 | 31 | 30 | 45 |
| 15.3 | 1 | 2 | 38 | 30 | 43 |
| 15.4 | 1 | 2 | 36 | 30 | 52 |
| 15.5 | 0 | 2 | 48 | 21 | 47 |
| 15.6 | 0 | 2 | 39 | 21 | 42 |
| 15.7 | 0 | 2 | 30 | 17 | 54 |
| 15.8 | 0 | 2 | 46 | 18 | 43 |
| 15.9 | 1 | 2 | 45 | 19 | 52 |
| 16 | 1 | 2 | 40 | 19 | 45 |
| 16.1 | 1 | 2 | 48 | 16 | 49 |
| 16.2 | 0 | 2 | 35 | 18 | 47 |
| 16.3 | 1 | 2 | 40 | 23 | 49 |
| 16.4 | 1 | 2 | 49 | 22 | 52 |
| 16.5 | 1 | 3 | 48 | 21 | 46 |
| 16.6 | 1 | 2 | 38 | 19 | 46 |
| 16.7 | 1 | 2 | 41 | 28 | 45 |
| 16.8 | 1 | 2 | 44 | 24 | 52 |
| 16.9 | 1 | 2 | 34 | 24 | 40 |
| 17 | 0 | 2 | 43 | 24 | 52 |
| 17.1 | 2 | 2 | 43 | 28 | 50 |
| 17.2 | 1 | 2 | 45 | 27 | 42 |
| 17.3 | 1 | 2 | 37 | 25 | 43 |
| 17.4 | 0 | 2 | 44 | 24 | 45 |
| 17.5 | 0 | 1 | 42 | 25 | 49 |
| 17.6 | 0 | 2 | 43 | 25 | 45 |
| 17.7 | 0 | 2 | 40 | 30 | 49 |
| 17.8 | 0 | 2 | 41 | 25 | 39 |
| 17.9 | 0 | 2 | 36 | 23 | 46 |
| 18 | 0 | 2 | 38 | 25 | 39 |
| 18.1 | 1 | 2 | 50 | 26 | 46 |
| 18.2 | 1 | 2 | 35 | 27 | 32 |
| 18.3 | 2 | 2 | 39 | 23 | 45 |
| 18.4 | 0 | 2 | 48 | 25 | 42 |
| 18.5 | 0 | 2 | 43 | 25 | 47 |
| 18.6 | 0 | 2 | 38 | 22 | 46 |
| 18.7 | 0 | 2 | 45 | 23 | 52 |
| 18.8 | 0 | 2 | 47 | 23 | 44 |
| 18.9 | 0 | 2 | 32 | 24 | 41 |
| 19 | 0 | 3 | 36 | 23 | 44 |
| 19.1 | 0 | 2 | 37 | 23 | 44 |
| 19.2 | 0 | 3 | 36 | 20 | 49 |
| 19.3 | 1 | 4 | 30 | 28 | 45 |
| 19.4 | 1 | 3 | 37 | 19 | 41 |
| 19.5 | 0 | 5 | 31 | 24 | 46 |
| 19.6 | 1 | 4 | 30 | 28 | 48 |
| 19.7 | 1 | 5 | 38 | 24 | 48 |
| 19.8 | 1 | 4 | 33 | 23 | 45 |
| 19.9 | 2 | 3 | 37 | 22 | 53 |

DNA"E-box" - CLOCK_bHLH (CLOCKS38/42/BMAL1S78)

Number of contacts

| Frame | 1st | 2nd | 3rd | 4th | 5th |
| --- | --- | --- | --- | --- | --- |
| 0 | 31 | 42 | 22 | 9 | 9 |
| 0.1 | 31 | 42 | 22 | 9 | 8 |
| 0.2 | 31 | 35 | 43 | 3 | 6 |
| 0.3 | 32 | 38 | 39 | 5 | 9 |
| 0.4 | 31 | 39 | 41 | 6 | 10 |
| 0.5 | 36 | 45 | 44 | 4 | 9 |
| 0.6 | 40 | 35 | 46 | 6 | 7 |
| 0.7 | 39 | 28 | 40 | 8 | 12 |
| 0.8 | 36 | 27 | 45 | 8 | 10 |
| 0.9 | 44 | 33 | 46 | 4 | 2 |
| 1 | 35 | 25 | 45 | 5 | 12 |
| 1.1 | 38 | 42 | 47 | 6 | 18 |
| 1.2 | 40 | 53 | 47 | 5 | 15 |
| 1.3 | 41 | 51 | 47 | 9 | 17 |
| 1.4 | 37 | 53 | 48 | 6 | 19 |
| 1.5 | 41 | 36 | 41 | 4 | 16 |
| 1.6 | 38 | 48 | 37 | 5 | 20 |
| 1.7 | 43 | 39 | 43 | 6 | 25 |
| 1.8 | 42 | 40 | 44 | 3 | 42 |
| 1.9 | 37 | 45 | 43 | 5 | 46 |
| 2 | 50 | 57 | 33 | 5 | 50 |
| 2.1 | 47 | 46 | 38 | 7 | 43 |
| 2.2 | 47 | 52 | 43 | 6 | 39 |
| 2.3 | 33 | 44 | 41 | 5 | 38 |
| 2.4 | 41 | 47 | 40 | 6 | 49 |
| 2.5 | 36 | 45 | 40 | 6 | 43 |
| 2.6 | 33 | 40 | 37 | 5 | 42 |
| 2.7 | 38 | 42 | 28 | 7 | 44 |
| 2.8 | 32 | 44 | 24 | 5 | 44 |
| 2.9 | 50 | 31 | 25 | 6 | 45 |
| 3 | 50 | 41 | 27 | 7 | 46 |
| 3.1 | 47 | 52 | 30 | 5 | 51 |
| 3.2 | 47 | 31 | 37 | 8 | 50 |
| 3.3 | 45 | 47 | 27 | 5 | 44 |
| 3.4 | 50 | 51 | 25 | 7 | 53 |
| 3.5 | 50 | 40 | 27 | 6 | 49 |
| 3.6 | 51 | 49 | 30 | 5 | 46 |
| 3.7 | 47 | 32 | 23 | 5 | 48 |
| 3.8 | 47 | 35 | 26 | 6 | 46 |
| 3.9 | 58 | 37 | 29 | 5 | 47 |
| 4 | 46 | 43 | 23 | 4 | 42 |
| 4.1 | 53 | 25 | 27 | 6 | 49 |
| 4.2 | 48 | 26 | 35 | 5 | 55 |
| 4.3 | 48 | 31 | 27 | 6 | 46 |
| 4.4 | 40 | 28 | 28 | 5 | 41 |
| 4.5 | 56 | 33 | 31 | 7 | 40 |
| 4.6 | 43 | 33 | 33 | 7 | 48 |
| 4.7 | 41 | 39 | 27 | 6 | 54 |
| 4.8 | 48 | 39 | 25 | 7 | 49 |
| 4.9 | 58 | 45 | 31 | 8 | 46 |
| 5 | 50 | 34 | 27 | 7 | 46 |
| 5.1 | 44 | 37 | 30 | 6 | 51 |
| 5.2 | 39 | 23 | 24 | 6 | 52 |
| 5.3 | 58 | 25 | 30 | 6 | 44 |
| 5.4 | 51 | 48 | 27 | 11 | 45 |
| 5.5 | 54 | 33 | 28 | 8 | 36 |
| 5.6 | 47 | 36 | 32 | 7 | 43 |
| 5.7 | 45 | 26 | 31 | 7 | 37 |
| 5.8 | 48 | 42 | 21 | 8 | 47 |
| 5.9 | 47 | 43 | 22 | 8 | 44 |
| 6 | 46 | 41 | 22 | 4 | 49 |
| 6.1 | 48 | 40 | 21 | 8 | 41 |
| 6.2 | 51 | 43 | 26 | 7 | 54 |
| 6.3 | 50 | 23 | 24 | 7 | 45 |
| 6.4 | 51 | 42 | 23 | 7 | 52 |
| 6.5 | 36 | 40 | 28 | 6 | 48 |
| 6.6 | 43 | 44 | 25 | 6 | 45 |
| 6.7 | 48 | 34 | 27 | 9 | 53 |
| 6.8 | 48 | 29 | 24 | 9 | 41 |
| 6.9 | 50 | 41 | 25 | 10 | 49 |
| 7 | 47 | 52 | 26 | 8 | 42 |
| 7.1 | 46 | 42 | 23 | 9 | 44 |
| 7.2 | 46 | 42 | 22 | 8 | 34 |
| 7.3 | 39 | 37 | 23 | 6 | 47 |
| 7.4 | 52 | 25 | 24 | 8 | 48 |
| 7.5 | 47 | 31 | 25 | 6 | 50 |
| 7.6 | 52 | 33 | 29 | 8 | 51 |
| 7.7 | 44 | 41 | 23 | 8 | 48 |
| 7.8 | 46 | 46 | 24 | 9 | 47 |
| 7.9 | 49 | 46 | 27 | 9 | 49 |
| 8 | 50 | 40 | 25 | 7 | 47 |
| 8.1 | 44 | 35 | 25 | 8 | 40 |
| 8.2 | 51 | 29 | 22 | 7 | 46 |
| 8.3 | 53 | 38 | 20 | 8 | 38 |
| 8.4 | 42 | 37 | 24 | 8 | 35 |
| 8.5 | 56 | 27 | 23 | 7 | 39 |
| 8.6 | 53 | 41 | 25 | 9 | 27 |
| 8.7 | 48 | 42 | 30 | 6 | 27 |
| 8.8 | 43 | 35 | 24 | 9 | 22 |
| 8.9 | 40 | 32 | 22 | 7 | 21 |
| 9 | 41 | 30 | 25 | 7 | 21 |
| 9.1 | 61 | 31 | 21 | 6 | 21 |
| 9.2 | 44 | 29 | 27 | 7 | 28 |
| 9.3 | 46 | 31 | 27 | 7 | 22 |
| 9.4 | 55 | 25 | 22 | 6 | 21 |
| 9.5 | 53 | 24 | 25 | 6 | 20 |
| 9.6 | 51 | 21 | 29 | 7 | 23 |
| 9.7 | 55 | 20 | 26 | 8 | 22 |
| 9.8 | 50 | 23 | 30 | 6 | 24 |
| 9.9 | 49 | 26 | 30 | 7 | 22 |
| 10 | 45 | 25 | 31 | 8 | 26 |
| 10.1 | 48 | 28 | 25 | 9 | 26 |
| 10.2 | 47 | 30 | 25 | 6 | 23 |
| 10.3 | 52 | 35 | 30 | 8 | 26 |
| 10.4 | 40 | 42 | 22 | 7 | 22 |
| 10.5 | 49 | 43 | 27 | 7 | 20 |
| 10.6 | 44 | 53 | 23 | 8 | 25 |
| 10.7 | 46 | 50 | 28 | 8 | 24 |
| 10.8 | 44 | 34 | 23 | 6 | 24 |
| 10.9 | 48 | 35 | 23 | 7 | 20 |
| 11 | 49 | 37 | 25 | 7 | 23 |
| 11.1 | 47 | 33 | 22 | 8 | 21 |
| 11.2 | 43 | 36 | 24 | 9 | 33 |
| 11.3 | 50 | 33 | 28 | 11 | 21 |
| 11.4 | 44 | 41 | 29 | 9 | 19 |
| 11.5 | 48 | 43 | 23 | 7 | 21 |
| 11.6 | 43 | 33 | 26 | 6 | 19 |
| 11.7 | 40 | 37 | 22 | 6 | 22 |
| 11.8 | 57 | 30 | 26 | 9 | 25 |
| 11.9 | 45 | 45 | 25 | 7 | 20 |
| 12 | 42 | 49 | 34 | 4 | 24 |
| 12.1 | 43 | 41 | 26 | 6 | 20 |
| 12.2 | 48 | 46 | 43 | 8 | 21 |
| 12.3 | 52 | 42 | 45 | 7 | 20 |
| 12.4 | 52 | 43 | 46 | 6 | 19 |
| 12.5 | 51 | 53 | 53 | 8 | 22 |
| 12.6 | 46 | 45 | 45 | 8 | 21 |
| 12.7 | 45 | 44 | 43 | 7 | 21 |
| 12.8 | 44 | 47 | 45 | 7 | 21 |
| 12.9 | 50 | 39 | 47 | 7 | 31 |
| 13 | 47 | 50 | 54 | 7 | 25 |
| 13.1 | 43 | 45 | 56 | 7 | 17 |
| 13.2 | 56 | 49 | 40 | 7 | 23 |
| 13.3 | 45 | 27 | 47 | 8 | 24 |
| 13.4 | 43 | 41 | 45 | 8 | 24 |
| 13.5 | 46 | 27 | 44 | 8 | 24 |
| 13.6 | 54 | 40 | 46 | 7 | 22 |
| 13.7 | 40 | 36 | 49 | 7 | 21 |
| 13.8 | 53 | 42 | 45 | 6 | 23 |
| 13.9 | 40 | 39 | 53 | 7 | 25 |
| 14 | 54 | 45 | 42 | 8 | 29 |
| 14.1 | 45 | 40 | 48 | 6 | 31 |
| 14.2 | 44 | 33 | 54 | 7 | 51 |
| 14.3 | 49 | 35 | 52 | 6 | 41 |
| 14.4 | 46 | 41 | 54 | 8 | 41 |
| 14.5 | 49 | 47 | 45 | 7 | 51 |
| 14.6 | 52 | 42 | 45 | 7 | 43 |
| 14.7 | 54 | 37 | 43 | 7 | 47 |
| 14.8 | 46 | 33 | 47 | 12 | 43 |
| 14.9 | 41 | 29 | 53 | 9 | 48 |
| 15 | 49 | 43 | 46 | 6 | 46 |
| 15.1 | 46 | 37 | 43 | 8 | 40 |
| 15.2 | 48 | 40 | 45 | 7 | 49 |
| 15.3 | 53 | 36 | 55 | 8 | 50 |
| 15.4 | 49 | 41 | 43 | 6 | 25 |
| 15.5 | 42 | 38 | 53 | 8 | 24 |
| 15.6 | 48 | 46 | 46 | 7 | 32 |
| 15.7 | 52 | 36 | 50 | 8 | 29 |
| 15.8 | 45 | 41 | 53 | 8 | 32 |
| 15.9 | 43 | 36 | 42 | 7 | 23 |
| 16 | 52 | 40 | 46 | 5 | 26 |
| 16.1 | 53 | 42 | 50 | 9 | 26 |
| 16.2 | 42 | 39 | 40 | 7 | 32 |
| 16.3 | 44 | 46 | 49 | 6 | 42 |
| 16.4 | 43 | 34 | 50 | 7 | 47 |
| 16.5 | 35 | 44 | 46 | 8 | 42 |
| 16.6 | 41 | 32 | 54 | 6 | 48 |
| 16.7 | 50 | 42 | 49 | 8 | 43 |
| 16.8 | 44 | 46 | 50 | 6 | 39 |
| 16.9 | 42 | 41 | 47 | 6 | 27 |
| 17 | 43 | 49 | 51 | 7 | 26 |
| 17.1 | 42 | 44 | 49 | 9 | 27 |
| 17.2 | 48 | 41 | 45 | 8 | 30 |
| 17.3 | 45 | 37 | 55 | 8 | 29 |
| 17.4 | 46 | 42 | 54 | 8 | 27 |
| 17.5 | 39 | 41 | 54 | 8 | 23 |
| 17.6 | 47 | 39 | 46 | 8 | 24 |
| 17.7 | 50 | 45 | 56 | 7 | 28 |
| 17.8 | 48 | 51 | 53 | 7 | 27 |
| 17.9 | 49 | 47 | 47 | 9 | 25 |
| 18 | 53 | 37 | 49 | 8 | 21 |
| 18.1 | 36 | 37 | 52 | 9 | 30 |
| 18.2 | 42 | 40 | 41 | 6 | 31 |
| 18.3 | 39 | 41 | 44 | 8 | 38 |
| 18.4 | 43 | 48 | 42 | 6 | 42 |
| 18.5 | 52 | 38 | 46 | 8 | 42 |
| 18.6 | 43 | 34 | 52 | 7 | 40 |
| 18.7 | 53 | 47 | 47 | 7 | 30 |
| 18.8 | 48 | 35 | 44 | 9 | 41 |
| 18.9 | 51 | 46 | 47 | 8 | 51 |
| 19 | 41 | 38 | 52 | 7 | 35 |
| 19.1 | 53 | 37 | 46 | 8 | 51 |
| 19.2 | 46 | 39 | 50 | 7 | 43 |
| 19.3 | 48 | 39 | 46 | 7 | 40 |
| 19.4 | 56 | 41 | 49 | 7 | 46 |
| 19.5 | 43 | 43 | 45 | 7 | 47 |
| 19.6 | 39 | 50 | 55 | 7 | 51 |
| 19.7 | 47 | 44 | 53 | 4 | 55 |
| 19.8 | 39 | 42 | 48 | 7 | 58 |
| 19.9 | 45 | 42 | 46 | 6 | 69 |

**Figure S2.** Timeline plots pertaining to number of contacts between the E-box and CLOCK bHLH for the systems studied.

DNA "E-box" - BMAL1_bHLH (nonphosphorylated)

Number of contacts

| Frame | 1st | 2nd | 3rd | 4th | 5th |
| --- | --- | --- | --- | --- | --- |
| 0 | 15 | 8 | 13 | 5 | 69 |
| 0.1 | 15 | 8 | 13 | 5 | 38 |
| 0.2 | 15 | 8 | 15 | 3 | 58 |
| 0.3 | 15 | 8 | 14 | 3 | 49 |
| 0.4 | 12 | 11 | 7 | 3 | 62 |
| 0.5 | 14 | 12 | 4 | 9 | 65 |
| 0.6 | 16 | 12 | 8 | 10 | 62 |
| 0.7 | 12 | 12 | 4 | 14 | 54 |
| 0.8 | 15 | 15 | 3 | 9 | 61 |
| 0.9 | 10 | 9 | 7 | 13 | 63 |
| 1 | 13 | 14 | 3 | 4 | 69 |
| 1.1 | 6 | 16 | 15 | 8 | 66 |
| 1.2 | 20 | 9 | 0 | 6 | 69 |
| 1.3 | 18 | 12 | 6 | 11 | 62 |
| 1.4 | 15 | 16 | 10 | 6 | 64 |
| 1.5 | 11 | 9 | 12 | 17 | 62 |
| 1.6 | 14 | 14 | 9 | 8 | 54 |
| 1.7 | 11 | 11 | 9 | 7 | 51 |
| 1.8 | 14 | 12 | 1 | 15 | 50 |
| 1.9 | 9 | 8 | 3 | 16 | 53 |
| 2 | 16 | 10 | 4 | 14 | 54 |
| 2.1 | 15 | 9 | 10 | 16 | 45 |
| 2.2 | 14 | 12 | 7 | 11 | 68 |
| 2.3 | 13 | 10 | 3 | 5 | 60 |
| 2.4 | 9 | 13 | 7 | 13 | 64 |
| 2.5 | 14 | 11 | 7 | 6 | 64 |
| 2.6 | 7 | 18 | 5 | 4 | 64 |
| 2.7 | 7 | 18 | 8 | 8 | 57 |
| 2.8 | 6 | 10 | 9 | 13 | 59 |
| 2.9 | 9 | 14 | 9 | 5 | 62 |
| 3 | 8 | 11 | 7 | 9 | 59 |
| 3.1 | 11 | 12 | 12 | 12 | 46 |
| 3.2 | 10 | 14 | 10 | 3 | 48 |
| 3.3 | 9 | 21 | 16 | 4 | 38 |
| 3.4 | 10 | 17 | 18 | 8 | 45 |
| 3.5 | 8 | 30 | 22 | 8 | 51 |
| 3.6 | 7 | 18 | 10 | 5 | 51 |
| 3.7 | 7 | 28 | 8 | 5 | 46 |
| 3.8 | 11 | 14 | 9 | 6 | 64 |
| 3.9 | 10 | 14 | 5 | 5 | 64 |
| 4 | 9 | 16 | 9 | 10 | 50 |
| 4.1 | 2 | 18 | 5 | 14 | 39 |
| 4.2 | 3 | 11 | 16 | 10 | 31 |
| 4.3 | 3 | 31 | 26 | 9 | 48 |
| 4.4 | 2 | 26 | 26 | 7 | 37 |
| 4.5 | 1 | 29 | 20 | 4 | 37 |
| 4.6 | 10 | 27 | 29 | 6 | 37 |
| 4.7 | 11 | 19 | 30 | 4 | 44 |
| 4.8 | 3 | 9 | 5 | 5 | 40 |
| 4.9 | 4 | 8 | 10 | 14 | 37 |
| 5 | 3 | 16 | 4 | 5 | 38 |
| 5.1 | 3 | 23 | 7 | 11 | 41 |
| 5.2 | 3 | 30 | 10 | 7 | 43 |
| 5.3 | 3 | 26 | 4 | 11 | 41 |
| 5.4 | 3 | 10 | 10 | 5 | 42 |
| 5.5 | 4 | 36 | 14 | 9 | 39 |
| 5.6 | 4 | 18 | 5 | 10 | 45 |
| 5.7 | 5 | 30 | 7 | 20 | 46 |
| 5.8 | 3 | 29 | 7 | 9 | 40 |
| 5.9 | 2 | 30 | 12 | 8 | 45 |
| 6 | 3 | 29 | 13 | 10 | 41 |
| 6.1 | 3 | 31 | 10 | 15 | 37 |
| 6.2 | 3 | 24 | 5 | 10 | 46 |
| 6.3 | 3 | 25 | 5 | 7 | 43 |
| 6.4 | 4 | 30 | 10 | 6 | 47 |
| 6.5 | 4 | 16 | 8 | 6 | 40 |
| 6.6 | 7 | 27 | 13 | 14 | 42 |
| 6.7 | 3 | 28 | 25 | 11 | 43 |
| 6.8 | 6 | 26 | 9 | 4 | 46 |
| 6.9 | 7 | 27 | 15 | 12 | 42 |
| 7 | 5 | 20 | 6 | 6 | 43 |
| 7.1 | 8 | 28 | 7 | 8 | 33 |
| 7.2 | 4 | 29 | 12 | 4 | 41 |
| 7.3 | 1 | 27 | 15 | 12 | 45 |
| 7.4 | 4 | 26 | 10 | 11 | 43 |
| 7.5 | 5 | 29 | 19 | 5 | 42 |
| 7.6 | 4 | 27 | 33 | 8 | 39 |
| 7.7 | 2 | 30 | 29 | 13 | 37 |
| 7.8 | 4 | 29 | 31 | 8 | 56 |
| 7.9 | 4 | 27 | 22 | 7 | 48 |
| 8 | 4 | 29 | 31 | 9 | 53 |
| 8.1 | 4 | 23 | 26 | 16 | 46 |
| 8.2 | 4 | 15 | 21 | 7 | 46 |
| 8.3 | 5 | 22 | 28 | 11 | 59 |
| 8.4 | 3 | 40 | 30 | 14 | 46 |
| 8.5 | 2 | 35 | 27 | 11 | 53 |
| 8.6 | 4 | 43 | 28 | 13 | 53 |
| 8.7 | 1 | 42 | 28 | 14 | 52 |
| 8.8 | 3 | 48 | 31 | 11 | 55 |
| 8.9 | 2 | 43 | 20 | 4 | 45 |
| 9 | 4 | 45 | 26 | 5 | 39 |
| 9.1 | 5 | 51 | 30 | 12 | 41 |
| 9.2 | 2 | 49 | 24 | 16 | 42 |
| 9.3 | 9 | 43 | 36 | 9 | 39 |
| 9.4 | 9 | 47 | 28 | 5 | 35 |
| 9.5 | 12 | 44 | 27 | 5 | 46 |
| 9.6 | 2 | 45 | 30 | 2 | 38 |
| 9.7 | 6 | 48 | 27 | 4 | 43 |
| 9.8 | 7 | 51 | 31 | 2 | 46 |
| 9.9 | 6 | 48 | 35 | 6 | 39 |
| 10 | 3 | 46 | 29 | 7 | 44 |
| 10.1 | 3 | 57 | 28 | 2 | 41 |
| 10.2 | 1 | 44 | 28 | 4 | 35 |
| 10.3 | 1 | 48 | 29 | 10 | 46 |
| 10.4 | 5 | 8 | 33 | 3 | 46 |
| 10.5 | 4 | 49 | 33 | 8 | 42 |
| 10.6 | 6 | 51 | 26 | 9 | 42 |
| 10.7 | 3 | 55 | 28 | 4 | 38 |
| 10.8 | 4 | 53 | 26 | 8 | 36 |
| 10.9 | 6 | 53 | 22 | 7 | 38 |
| 11 | 4 | 48 | 29 | 7 | 35 |
| 11.1 | 4 | 50 | 30 | 7 | 42 |
| 11.2 | 3 | 51 | 30 | 8 | 39 |
| 11.3 | 2 | 49 | 22 | 9 | 40 |
| 11.4 | 3 | 43 | 28 | 8 | 37 |
| 11.5 | 4 | 39 | 28 | 8 | 35 |
| 11.6 | 10 | 38 | 28 | 7 | 42 |
| 11.7 | 7 | 51 | 30 | 8 | 41 |
| 11.8 | 4 | 42 | 31 | 6 | 40 |
| 11.9 | 3 | 44 | 32 | 8 | 43 |
| 12 | 1 | 47 | 20 | 7 | 40 |
| 12.1 | 1 | 43 | 26 | 8 | 37 |
| 12.2 | 2 | 47 | 31 | 8 | 33 |
| 12.3 | 0 | 41 | 30 | 9 | 41 |
| 12.4 | 4 | 43 | 26 | 9 | 39 |
| 12.5 | 3 | 44 | 28 | 15 | 38 |
| 12.6 | 4 | 45 | 32 | 9 | 37 |
| 12.7 | 6 | 42 | 28 | 20 | 40 |
| 12.8 | 4 | 40 | 26 | 17 | 44 |
| 12.9 | 8 | 39 | 25 | 8 | 44 |
| 13 | 7 | 41 | 26 | 6 | 46 |
| 13.1 | 8 | 48 | 24 | 8 | 41 |
| 13.2 | 5 | 48 | 37 | 10 | 37 |
| 13.3 | 3 | 41 | 42 | 9 | 42 |
| 13.4 | 3 | 47 | 39 | 7 | 43 |
| 13.5 | 1 | 47 | 45 | 9 | 44 |
| 13.6 | 4 | 39 | 46 | 10 | 46 |
| 13.7 | 2 | 48 | 45 | 8 | 37 |
| 13.8 | 1 | 40 | 48 | 8 | 38 |
| 13.9 | 3 | 42 | 50 | 8 | 43 |
| 14 | 2 | 41 | 46 | 7 | 42 |
| 14.1 | 1 | 37 | 48 | 7 | 43 |
| 14.2 | 2 | 41 | 43 | 10 | 46 |
| 14.3 | 3 | 49 | 40 | 10 | 39 |
| 14.4 | 2 | 48 | 51 | 10 | 38 |
| 14.5 | 2 | 52 | 45 | 6 | 40 |
| 14.6 | 1 | 51 | 44 | 10 | 36 |
| 14.7 | 5 | 47 | 55 | 11 | 43 |
| 14.8 | 9 | 50 | 43 | 10 | 45 |
| 14.9 | 11 | 50 | 44 | 8 | 43 |
| 15 | 4 | 45 | 57 | 7 | 35 |
| 15.1 | 8 | 46 | 40 | 9 | 37 |
| 15.2 | 5 | 49 | 48 | 8 | 41 |
| 15.3 | 7 | 50 | 45 | 6 | 38 |
| 15.4 | 4 | 52 | 39 | 8 | 42 |
| 15.5 | 3 | 50 | 47 | 7 | 39 |
| 15.6 | 2 | 51 | 49 | 9 | 45 |
| 15.7 | 1 | 49 | 44 | 10 | 44 |
| 15.8 | 4 | 50 | 45 | 8 | 45 |
| 15.9 | 0 | 43 | 47 | 7 | 45 |
| 16 | 2 | 57 | 47 | 7 | 41 |
| 16.1 | 4 | 53 | 47 | 9 | 39 |
| 16.2 | 4 | 56 | 44 | 8 | 47 |
| 16.3 | 4 | 47 | 43 | 6 | 47 |
| 16.4 | 5 | 51 | 46 | 8 | 48 |
| 16.5 | 3 | 52 | 45 | 7 | 43 |
| 16.6 | 5 | 47 | 44 | 8 | 47 |
| 16.7 | 5 | 53 | 41 | 7 | 42 |
| 16.8 | 5 | 55 | 52 | 7 | 44 |
| 16.9 | 2 | 47 | 43 | 11 | 44 |
| 17 | 7 | 50 | 52 | 10 | 46 |
| 17.1 | 4 | 53 | 44 | 6 | 38 |
| 17.2 | 4 | 49 | 54 | 6 | 43 |
| 17.3 | 3 | 53 | 40 | 6 | 43 |
| 17.4 | 3 | 45 | 47 | 9 | 47 |
| 17.5 | 6 | 53 | 48 | 7 | 37 |
| 17.6 | 3 | 48 | 35 | 6 | 38 |
| 17.7 | 3 | 52 | 43 | 9 | 42 |
| 17.8 | 2 | 47 | 43 | 7 | 37 |
| 17.9 | 2 | 45 | 43 | 7 | 47 |
| 18 | 3 | 44 | 47 | 7 | 37 |
| 18.1 | 0 | 46 | 43 | 8 | 44 |
| 18.2 | 1 | 48 | 51 | 8 | 44 |
| 18.3 | 1 | 50 | 40 | 8 | 42 |
| 18.4 | 1 | 44 | 44 | 6 | 44 |
| 18.5 | 2 | 45 | 44 | 6 | 49 |
| 18.6 | 3 | 53 | 53 | 6 | 41 |
| 18.7 | 5 | 48 | 49 | 5 | 47 |
| 18.8 | 5 | 48 | 45 | 7 | 46 |
| 18.9 | 0 | 47 | 44 | 7 | 46 |
| 19 | 4 | 52 | 43 | 7 | 43 |
| 19.1 | 1 | 41 | 41 | 5 | 43 |
| 19.2 | 4 | 52 | 54 | 8 | 41 |
| 19.3 | 4 | 45 | 44 | 9 | 39 |
| 19.4 | 2 | 48 | 47 | 6 | 52 |
| 19.5 | 4 | 40 | 45 | 7 | 37 |
| 19.6 | 2 | 57 | 43 | 8 | 49 |
| 19.7 | 2 | 52 | 40 | 7 | 44 |
| 19.8 | 26 | 45 | 44 | 10 | 47 |
| 19.9 | 5 | 53 | 55 | 8 | 50 |

DNA "E-box" - BMAL1_bHLH (BMAL1S78)

Number of contacts

| Frame | 1st | 2nd | 3rd | 4th | 5th |
| --- | --- | --- | --- | --- | --- |
| 0 | 4 | 8 | 4 | 65 | 13 |
| 0.1 | 5 | 8 | 6 | 8 | 12 |
| 0.2 | 1 | 1 | 3 | 5 | 2 |
| 0.3 | 2 | 2 | 2 | 6 | 3 |
| 0.4 | 3 | 3 | 3 | 10 | 3 |
| 0.5 | 5 | 3 | 5 | 5 | 1 |
| 0.6 | 4 | 7 | 8 | 8 | 1 |
| 0.7 | 4 | 13 | 11 | 8 | 2 |
| 0.8 | 7 | 16 | 10 | 9 | 1 |
| 0.9 | 7 | 11 | 6 | 9 | 1 |
| 1 | 10 | 5 | 6 | 8 | 1 |
| 1.1 | 9 | 5 | 8 | 12 | 1 |
| 1.2 | 9 | 5 | 9 | 6 | 1 |
| 1.3 | 10 | 4 | 7 | 7 | 4 |
| 1.4 | 9 | 5 | 16 | 9 | 5 |
| 1.5 | 11 | 4 | 22 | 8 | 2 |
| 1.6 | 10 | 7 | 19 | 16 | 2 |
| 1.7 | 6 | 2 | 16 | 15 | 1 |
| 1.8 | 11 | 2 | 20 | 19 | 1 |
| 1.9 | 11 | 6 | 16 | 17 | 1 |
| 2 | 10 | 6 | 21 | 16 | 2 |
| 2.1 | 6 | 2 | 18 | 17 | 1 |
| 2.2 | 9 | 3 | 17 | 16 | 1 |
| 2.3 | 6 | 4 | 24 | 16 | 1 |
| 2.4 | 7 | 6 | 19 | 14 | 1 |
| 2.5 | 1 | 6 | 21 | 16 | 1 |
| 2.6 | 7 | 4 | 17 | 17 | 1 |
| 2.7 | 7 | 17 | 19 | 18 | 1 |
| 2.8 | 5 | 15 | 17 | 13 | 4 |
| 2.9 | 6 | 13 | 14 | 12 | 3 |
| 3 | 4 | 7 | 13 | 15 | 3 |
| 3.1 | 9 | 12 | 15 | 18 | 2 |
| 3.2 | 10 | 13 | 16 | 17 | 5 |
| 3.3 | 8 | 10 | 18 | 16 | 3 |
| 3.4 | 8 | 12 | 16 | 13 | 2 |
| 3.5 | 5 | 12 | 19 | 13 | 3 |
| 3.6 | 8 | 14 | 20 | 14 | 3 |
| 3.7 | 4 | 15 | 18 | 17 | 4 |
| 3.8 | 6 | 10 | 19 | 17 | 4 |
| 3.9 | 7 | 13 | 18 | 13 | 3 |
| 4 | 6 | 9 | 18 | 16 | 4 |
| 4.1 | 7 | 4 | 19 | 15 | 4 |
| 4.2 | 6 | 16 | 19 | 23 | 1 |
| 4.3 | 6 | 9 | 29 | 17 | 5 |
| 4.4 | 9 | 5 | 17 | 16 | 3 |
| 4.5 | 7 | 4 | 22 | 16 | 4 |
| 4.6 | 7 | 3 | 20 | 16 | 5 |
| 4.7 | 7 | 3 | 21 | 18 | 5 |
| 4.8 | 8 | 4 | 21 | 15 | 6 |
| 4.9 | 7 | 4 | 19 | 19 | 6 |
| 5 | 7 | 3 | 23 | 13 | 3 |
| 5.1 | 6 | 5 | 18 | 19 | 3 |
| 5.2 | 7 | 6 | 21 | 13 | 6 |
| 5.3 | 5 | 8 | 23 | 12 | 8 |
| 5.4 | 8 | 7 | 24 | 17 | 4 |
| 5.5 | 15 | 3 | 17 | 19 | 4 |
| 5.6 | 12 | 2 | 14 | 20 | 4 |
| 5.7 | 13 | 4 | 18 | 16 | 5 |
| 5.8 | 13 | 3 | 17 | 19 | 5 |
| 5.9 | 14 | 2 | 14 | 20 | 6 |
| 6 | 10 | 4 | 16 | 17 | 7 |
| 6.1 | 4 | 4 | 18 | 17 | 5 |
| 6.2 | 7 | 3 | 17 | 14 | 6 |
| 6.3 | 6 | 4 | 15 | 13 | 15 |
| 6.4 | 7 | 1 | 17 | 12 | 4 |
| 6.5 | 9 | 2 | 19 | 14 | 8 |
| 6.6 | 5 | 5 | 20 | 16 | 4 |
| 6.7 | 5 | 3 | 17 | 17 | 3 |
| 6.8 | 8 | 5 | 27 | 18 | 5 |
| 6.9 | 7 | 5 | 17 | 19 | 3 |
| 7 | 8 | 7 | 22 | 21 | 5 |
| 7.1 | 8 | 12 | 23 | 18 | 5 |
| 7.2 | 7 | 12 | 20 | 22 | 5 |
| 7.3 | 7 | 7 | 27 | 19 | 5 |
| 7.4 | 7 | 8 | 17 | 15 | 7 |
| 7.5 | 4 | 11 | 18 | 15 | 6 |
| 7.6 | 8 | 13 | 25 | 14 | 6 |
| 7.7 | 10 | 10 | 21 | 13 | 5 |
| 7.8 | 8 | 6 | 20 | 19 | 7 |
| 7.9 | 5 | 9 | 19 | 16 | 6 |
| 8 | 7 | 10 | 18 | 18 | 2 |
| 8.1 | 6 | 10 | 28 | 16 | 6 |
| 8.2 | 6 | 14 | 23 | 17 | 8 |
| 8.3 | 6 | 10 | 20 | 17 | 2 |
| 8.4 | 8 | 9 | 14 | 18 | 3 |
| 8.5 | 6 | 8 | 16 | 17 | 6 |
| 8.6 | 6 | 9 | 16 | 20 | 6 |
| 8.7 | 8 | 4 | 22 | 16 | 7 |
| 8.8 | 9 | 10 | 25 | 17 | 4 |
| 8.9 | 8 | 8 | 21 | 16 | 8 |
| 9 | 7 | 8 | 19 | 16 | 10 |
| 9.1 | 7 | 12 | 23 | 16 | 4 |
| 9.2 | 10 | 7 | 21 | 14 | 6 |
| 9.3 | 8 | 14 | 24 | 15 | 7 |
| 9.4 | 4 | 5 | 17 | 16 | 5 |
| 9.5 | 9 | 9 | 23 | 17 | 4 |
| 9.6 | 8 | 16 | 24 | 17 | 6 |
| 9.7 | 7 | 10 | 19 | 17 | 1 |
| 9.8 | 6 | 11 | 19 | 18 | 13 |
| 9.9 | 6 | 11 | 21 | 20 | 16 |
| 10 | 6 | 15 | 21 | 19 | 20 |
| 10.1 | 10 | 13 | 19 | 14 | 6 |
| 10.2 | 9 | 11 | 25 | 16 | 5 |
| 10.3 | 7 | 11 | 25 | 16 | 3 |
| 10.4 | 6 | 10 | 22 | 14 | 5 |
| 10.5 | 7 | 6 | 23 | 19 | 5 |
| 10.6 | 8 | 10 | 15 | 17 | 7 |
| 10.7 | 8 | 6 | 21 | 17 | 6 |
| 10.8 | 7 | 10 | 22 | 18 | 18 |
| 10.9 | 6 | 9 | 18 | 20 | 11 |
| 11 | 7 | 14 | 21 | 19 | 5 |
| 11.1 | 6 | 8 | 20 | 18 | 5 |
| 11.2 | 7 | 10 | 22 | 19 | 5 |
| 11.3 | 5 | 6 | 17 | 23 | 2 |
| 11.4 | 7 | 9 | 25 | 25 | 9 |
| 11.5 | 9 | 10 | 20 | 22 | 4 |
| 11.6 | 8 | 14 | 19 | 23 | 4 |
| 11.7 | 7 | 7 | 17 | 21 | 4 |
| 11.8 | 8 | 11 | 24 | 16 | 6 |
| 11.9 | 7 | 5 | 21 | 20 | 3 |
| 12 | 9 | 11 | 17 | 19 | 4 |
| 12.1 | 8 | 10 | 25 | 20 | 3 |
| 12.2 | 11 | 11 | 17 | 7 | 5 |
| 12.3 | 9 | 10 | 20 | 24 | 1 |
| 12.4 | 10 | 11 | 16 | 18 | 3 |
| 12.5 | 6 | 5 | 20 | 28 | 3 |
| 12.6 | 8 | 8 | 19 | 18 | 4 |
| 12.7 | 7 | 8 | 26 | 18 | 3 |
| 12.8 | 7 | 5 | 23 | 17 | 4 |
| 12.9 | 4 | 6 | 21 | 20 | 4 |
| 13 | 6 | 10 | 26 | 19 | 3 |
| 13.1 | 4 | 9 | 23 | 19 | 7 |
| 13.2 | 3 | 8 | 19 | 18 | 5 |
| 13.3 | 7 | 8 | 16 | 22 | 6 |
| 13.4 | 5 | 7 | 16 | 12 | 2 |
| 13.5 | 5 | 6 | 21 | 20 | 2 |
| 13.6 | 5 | 9 | 18 | 14 | 2 |
| 13.7 | 4 | 13 | 19 | 20 | 3 |
| 13.8 | 8 | 10 | 17 | 23 | 1 |
| 13.9 | 5 | 14 | 21 | 27 | 1 |
| 14 | 6 | 15 | 19 | 19 | 1 |
| 14.1 | 7 | 10 | 17 | 20 | 1 |
| 14.2 | 4 | 9 | 14 | 16 | 2 |
| 14.3 | 5 | 8 | 21 | 11 | 2 |
| 14.4 | 7 | 10 | 25 | 16 | 1 |
| 14.5 | 6 | 4 | 20 | 23 | 3 |
| 14.6 | 4 | 10 | 17 | 16 | 3 |
| 14.7 | 5 | 11 | 18 | 20 | 1 |
| 14.8 | 7 | 6 | 18 | 20 | 5 |
| 14.9 | 6 | 9 | 24 | 21 | 6 |
| 15 | 6 | 10 | 20 | 23 | 7 |
| 15.1 | 5 | 11 | 17 | 17 | 3 |
| 15.2 | 5 | 9 | 22 | 14 | 3 |
| 15.3 | 12 | 8 | 21 | 22 | 3 |
| 15.4 | 9 | 10 | 21 | 22 | 0 |
| 15.5 | 5 | 11 | 18 | 18 | 4 |
| 15.6 | 11 | 10 | 19 | 18 | 4 |
| 15.7 | 11 | 17 | 26 | 25 | 3 |
| 15.8 | 10 | 10 | 22 | 19 | 4 |
| 15.9 | 10 | 10 | 19 | 23 | 1 |
| 16 | 8 | 6 | 15 | 17 | 4 |
| 16.1 | 8 | 7 | 24 | 18 | 5 |
| 16.2 | 7 | 10 | 20 | 24 | 6 |
| 16.3 | 9 | 10 | 24 | 17 | 5 |
| 16.4 | 9 | 9 | 22 | 22 | 4 |
| 16.5 | 7 | 9 | 19 | 19 | 4 |
| 16.6 | 9 | 9 | 21 | 18 | 4 |
| 16.7 | 8 | 9 | 22 | 18 | 7 |
| 16.8 | 5 | 6 | 22 | 19 | 2 |
| 16.9 | 9 | 9 | 23 | 19 | 6 |
| 17 | 7 | 10 | 22 | 23 | 6 |
| 17.1 | 10 | 11 | 18 | 18 | 5 |
| 17.2 | 6 | 5 | 28 | 20 | 5 |
| 17.3 | 6 | 5 | 21 | 22 | 2 |
| 17.4 | 7 | 4 | 20 | 23 | 2 |
| 17.5 | 5 | 4 | 21 | 21 | 4 |
| 17.6 | 7 | 4 | 24 | 21 | 3 |
| 17.7 | 7 | 4 | 19 | 18 | 4 |
| 17.8 | 9 | 3 | 22 | 24 | 3 |
| 17.9 | 8 | 6 | 21 | 24 | 2 |
| 18 | 6 | 4 | 26 | 16 | 2 |
| 18.1 | 8 | 5 | 16 | 16 | 4 |
| 18.2 | 10 | 7 | 18 | 17 | 5 |
| 18.3 | 11 | 6 | 16 | 17 | 3 |
| 18.4 | 8 | 6 | 17 | 18 | 4 |
| 18.5 | 7 | 6 | 17 | 19 | 3 |
| 18.6 | 11 | 6 | 18 | 21 | 5 |
| 18.7 | 8 | 4 | 14 | 15 | 5 |
| 18.8 | 14 | 4 | 19 | 23 | 3 |
| 18.9 | 7 | 4 | 20 | 20 | 6 |
| 19 | 6 | 12 | 16 | 20 | 5 |
| 19.1 | 9 | 3 | 19 | 22 | 2 |
| 19.2 | 6 | 5 | 20 | 19 | 3 |
| 19.3 | 11 | 4 | 16 | 22 | 2 |
| 19.4 | 8 | 3 | 20 | 18 | 1 |
| 19.5 | 8 | 3 | 23 | 23 | 1 |
| 19.6 | 11 | 4 | 19 | 23 | 3 |
| 19.7 | 9 | 6 | 25 | 27 | 4 |
| 19.8 | 8 | 14 | 22 | 20 | 4 |
| 19.9 | 7 | 5 | 16 | 19 | 5 |

DNA "E-box" - BMAL1_bHLH (CLOCKS38/42)

Number of contacts

| Frame | 1st | 2nd | 3rd | 4th | 5th |
| --- | --- | --- | --- | --- | --- |
| 0 | 24 | 14 | 11 | 2 | 8 |
| 0.1 | 29 | 15 | 15 | 2 | 8 |
| 0.2 | 41 | 12 | 8 | 11 | 3 |
| 0.3 | 43 | 11 | 17 | 10 | 10 |
| 0.4 | 42 | 14 | 11 | 17 | 7 |
| 0.5 | 47 | 11 | 13 | 15 | 16 |
| 0.6 | 46 | 10 | 7 | 13 | 13 |
| 0.7 | 46 | 2 | 5 | 11 | 11 |
| 0.8 | 46 | 2 | 5 | 20 | 13 |
| 0.9 | 49 | 4 | 7 | 12 | 14 |
| 1 | 36 | 2 | 2 | 19 | 11 |
| 1.1 | 55 | 2 | 5 | 12 | 25 |
| 1.2 | 55 | 4 | 5 | 11 | 21 |
| 1.3 | 45 | 3 | 7 | 15 | 18 |
| 1.4 | 44 | 8 | 10 | 12 | 27 |
| 1.5 | 53 | 9 | 6 | 13 | 20 |
| 1.6 | 52 | 6 | 14 | 7 | 20 |
| 1.7 | 51 | 4 | 7 | 17 | 24 |
| 1.8 | 52 | 11 | 9 | 19 | 20 |
| 1.9 | 46 | 8 | 7 | 20 | 27 |
| 2 | 47 | 4 | 7 | 18 | 25 |
| 2.1 | 46 | 3 | 9 | 14 | 31 |
| 2.2 | 46 | 5 | 7 | 13 | 27 |
| 2.3 | 48 | 5 | 5 | 12 | 25 |
| 2.4 | 48 | 12 | 8 | 12 | 30 |
| 2.5 | 48 | 4 | 7 | 10 | 20 |
| 2.6 | 47 | 4 | 10 | 16 | 25 |
| 2.7 | 47 | 9 | 6 | 15 | 23 |
| 2.8 | 46 | 6 | 8 | 12 | 24 |
| 2.9 | 46 | 4 | 5 | 15 | 24 |
| 3 | 46 | 3 | 9 | 14 | 25 |
| 3.1 | 49 | 14 | 8 | 12 | 28 |
| 3.2 | 48 | 31 | 6 | 11 | 16 |
| 3.3 | 51 | 31 | 7 | 8 | 22 |
| 3.4 | 50 | 14 | 6 | 13 | 4 |
| 3.5 | 51 | 8 | 7 | 11 | 10 |
| 3.6 | 49 | 4 | 7 | 13 | 11 |
| 3.7 | 45 | 4 | 9 | 11 | 15 |
| 3.8 | 48 | 1 | 9 | 14 | 13 |
| 3.9 | 47 | 5 | 7 | 16 | 11 |
| 4 | 49 | 4 | 7 | 13 | 6 |
| 4.1 | 47 | 1 | 7 | 7 | 17 |
| 4.2 | 47 | 2 | 5 | 17 | 15 |
| 4.3 | 48 | 1 | 9 | 17 | 15 |
| 4.4 | 46 | 2 | 8 | 17 | 12 |
| 4.5 | 45 | 3 | 9 | 15 | 3 |
| 4.6 | 45 | 2 | 7 | 16 | 3 |
| 4.7 | 44 | 2 | 10 | 11 | 2 |
| 4.8 | 46 | 3 | 8 | 8 | 5 |
| 4.9 | 53 | 3 | 8 | 10 | 8 |
| 5 | 48 | 1 | 6 | 7 | 11 |
| 5.1 | 49 | 0 | 5 | 8 | 14 |
| 5.2 | 45 | 4 | 7 | 13 | 15 |
| 5.3 | 41 | 4 | 8 | 13 | 10 |
| 5.4 | 56 | 4 | 10 | 10 | 12 |
| 5.5 | 48 | 1 | 8 | 19 | 7 |
| 5.6 | 49 | 3 | 7 | 15 | 9 |
| 5.7 | 41 | 3 | 27 | 11 | 11 |
| 5.8 | 50 | 2 | 9 | 11 | 14 |
| 5.9 | 48 | 3 | 6 | 12 | 16 |
| 6 | 46 | 4 | 13 | 15 | 20 |
| 6.1 | 42 | 3 | 8 | 16 | 18 |
| 6.2 | 50 | 0 | 13 | 14 | 16 |
| 6.3 | 53 | 2 | 16 | 15 | 22 |
| 6.4 | 46 | 3 | 8 | 14 | 17 |
| 6.5 | 50 | 3 | 7 | 14 | 11 |
| 6.6 | 43 | 1 | 8 | 15 | 21 |
| 6.7 | 46 | 8 | 5 | 15 | 18 |
| 6.8 | 47 | 3 | 8 | 14 | 19 |
| 6.9 | 49 | 3 | 6 | 12 | 25 |
| 7 | 48 | 2 | 6 | 18 | 19 |
| 7.1 | 44 | 5 | 10 | 8 | 19 |
| 7.2 | 43 | 5 | 9 | 14 | 10 |
| 7.3 | 40 | 3 | 7 | 15 | 22 |
| 7.4 | 45 | 3 | 9 | 10 | 21 |
| 7.5 | 48 | 6 | 5 | 11 | 19 |
| 7.6 | 47 | 2 | 8 | 17 | 17 |
| 7.7 | 48 | 2 | 13 | 14 | 17 |
| 7.8 | 47 | 5 | 12 | 13 | 16 |
| 7.9 | 47 | 3 | 8 | 16 | 19 |
| 8 | 48 | 5 | 8 | 14 | 22 |
| 8.1 | 38 | 4 | 9 | 13 | 21 |
| 8.2 | 51 | 3 | 6 | 14 | 19 |
| 8.3 | 58 | 3 | 8 | 11 | 15 |
| 8.4 | 44 | 2 | 13 | 15 | 20 |
| 8.5 | 52 | 3 | 7 | 12 | 16 |
| 8.6 | 45 | 2 | 11 | 14 | 15 |
| 8.7 | 42 | 0 | 8 | 12 | 15 |
| 8.8 | 47 | 3 | 9 | 13 | 20 |
| 8.9 | 45 | 5 | 22 | 12 | 14 |
| 9 | 44 | 7 | 38 | 16 | 22 |
| 9.1 | 51 | 14 | 52 | 13 | 19 |
| 9.2 | 46 | 4 | 42 | 11 | 13 |
| 9.3 | 44 | 2 | 52 | 9 | 14 |
| 9.4 | 45 | 3 | 43 | 10 | 22 |
| 9.5 | 54 | 10 | 48 | 12 | 17 |
| 9.6 | 45 | 11 | 50 | 9 | 14 |
| 9.7 | 44 | 4 | 49 | 10 | 21 |
| 9.8 | 48 | 1 | 47 | 11 | 11 |
| 9.9 | 51 | 7 | 42 | 9 | 16 |
| 10 | 48 | 8 | 50 | 13 | 15 |
| 10.1 | 46 | 5 | 42 | 9 | 14 |
| 10.2 | 49 | 1 | 49 | 10 | 17 |
| 10.3 | 47 | 6 | 44 | 15 | 17 |
| 10.4 | 47 | 5 | 48 | 17 | 19 |
| 10.5 | 57 | 5 | 38 | 10 | 20 |
| 10.6 | 45 | 2 | 46 | 9 | 13 |
| 10.7 | 48 | 3 | 37 | 12 | 10 |
| 10.8 | 44 | 0 | 47 | 12 | 20 |
| 10.9 | 47 | 3 | 45 | 8 | 15 |
| 11 | 51 | 2 | 44 | 16 | 10 |
| 11.1 | 51 | 3 | 46 | 10 | 15 |
| 11.2 | 46 | 2 | 45 | 12 | 9 |
| 11.3 | 48 | 4 | 47 | 14 | 19 |
| 11.4 | 49 | 10 | 42 | 10 | 17 |
| 11.5 | 49 | 21 | 47 | 14 | 18 |
| 11.6 | 53 | 15 | 48 | 11 | 16 |
| 11.7 | 45 | 4 | 48 | 11 | 13 |
| 11.8 | 51 | 5 | 47 | 16 | 15 |
| 11.9 | 49 | 2 | 45 | 10 | 14 |
| 12 | 41 | 4 | 46 | 10 | 17 |
| 12.1 | 47 | 6 | 48 | 13 | 13 |
| 12.2 | 45 | 4 | 44 | 12 | 19 |
| 12.3 | 43 | 6 | 41 | 15 | 20 |
| 12.4 | 43 | 5 | 48 | 4 | 20 |
| 12.5 | 57 | 6 | 44 | 14 | 16 |
| 12.6 | 49 | 4 | 44 | 9 | 19 |
| 12.7 | 52 | 4 | 42 | 12 | 12 |
| 12.8 | 45 | 4 | 50 | 12 | 19 |
| 12.9 | 48 | 3 | 48 | 9 | 16 |
| 13 | 45 | 7 | 47 | 8 | 19 |
| 13.1 | 46 | 5 | 48 | 7 | 20 |
| 13.2 | 38 | 4 | 47 | 10 | 23 |
| 13.3 | 49 | 6 | 44 | 7 | 8 |
| 13.4 | 52 | 6 | 49 | 11 | 21 |
| 13.5 | 48 | 6 | 55 | 6 | 17 |
| 13.6 | 46 | 4 | 52 | 11 | 20 |
| 13.7 | 45 | 7 | 50 | 12 | 26 |
| 13.8 | 45 | 6 | 49 | 8 | 25 |
| 13.9 | 46 | 6 | 50 | 14 | 21 |
| 14 | 42 | 2 | 43 | 9 | 26 |
| 14.1 | 56 | 5 | 48 | 10 | 19 |
| 14.2 | 47 | 6 | 44 | 12 | 30 |
| 14.3 | 51 | 5 | 45 | 13 | 33 |
| 14.4 | 54 | 7 | 42 | 12 | 25 |
| 14.5 | 48 | 5 | 39 | 14 | 28 |
| 14.6 | 58 | 6 | 33 | 13 | 29 |
| 14.7 | 50 | 6 | 41 | 10 | 30 |
| 14.8 | 50 | 3 | 47 | 8 | 13 |
| 14.9 | 44 | 10 | 46 | 13 | 17 |
| 15 | 43 | 6 | 46 | 10 | 31 |
| 15.1 | 45 | 5 | 46 | 8 | 27 |
| 15.2 | 41 | 5 | 39 | 10 | 26 |
| 15.3 | 43 | 5 | 46 | 13 | 30 |
| 15.4 | 42 | 4 | 47 | 14 | 22 |
| 15.5 | 43 | 6 | 47 | 9 | 18 |
| 15.6 | 50 | 4 | 51 | 11 | 18 |
| 15.7 | 43 | 6 | 54 | 9 | 22 |
| 15.8 | 34 | 6 | 55 | 12 | 10 |
| 15.9 | 45 | 8 | 38 | 14 | 8 |
| 16 | 40 | 6 | 43 | 12 | 7 |
| 16.1 | 47 | 6 | 42 | 8 | 9 |
| 16.2 | 48 | 9 | 50 | 9 | 12 |
| 16.3 | 48 | 2 | 49 | 13 | 14 |
| 16.4 | 50 | 3 | 48 | 14 | 8 |
| 16.5 | 48 | 3 | 46 | 12 | 6 |
| 16.6 | 49 | 5 | 46 | 14 | 8 |
| 16.7 | 47 | 4 | 53 | 11 | 8 |
| 16.8 | 45 | 4 | 47 | 11 | 21 |
| 16.9 | 54 | 6 | 51 | 6 | 15 |
| 17 | 46 | 2 | 48 | 13 | 16 |
| 17.1 | 46 | 6 | 55 | 18 | 6 |
| 17.2 | 46 | 15 | 50 | 9 | 9 |
| 17.3 | 43 | 2 | 49 | 14 | 16 |
| 17.4 | 50 | 3 | 42 | 10 | 7 |
| 17.5 | 39 | 4 | 42 | 10 | 10 |
| 17.6 | 50 | 7 | 50 | 9 | 10 |
| 17.7 | 54 | 3 | 49 | 12 | 10 |
| 17.8 | 45 | 7 | 39 | 9 | 8 |
| 17.9 | 47 | 8 | 46 | 12 | 13 |
| 18 | 48 | 5 | 58 | 10 | 8 |
| 18.1 | 51 | 6 | 45 | 9 | 14 |
| 18.2 | 48 | 6 | 41 | 20 | 8 |
| 18.3 | 48 | 5 | 48 | 15 | 11 |
| 18.4 | 41 | 8 | 45 | 9 | 8 |
| 18.5 | 44 | 5 | 44 | 13 | 9 |
| 18.6 | 47 | 5 | 46 | 16 | 9 |
| 18.7 | 49 | 18 | 52 | 14 | 14 |
| 18.8 | 46 | 1 | 42 | 14 | 8 |
| 18.9 | 50 | 1 | 47 | 14 | 10 |
| 19 | 54 | 0 | 52 | 14 | 13 |
| 19.1 | 50 | 2 | 46 | 15 | 6 |
| 19.2 | 47 | 2 | 44 | 17 | 15 |
| 19.3 | 52 | 19 | 44 | 10 | 15 |
| 19.4 | 53 | 11 | 50 | 13 | 21 |
| 19.5 | 52 | 0 | 45 | 8 | 17 |
| 19.6 | 44 | 3 | 47 | 10 | 20 |
| 19.7 | 42 | 3 | 49 | 9 | 19 |
| 19.8 | 49 | 2 | 48 | 12 | 12 |
| 19.9 | 45 | 2 | 49 | 9 | 16 |

DNA "E-box" - BMAL1_bHLH (CLOCKS38/42/BMAL1S78)

Number of contacts

| Frame | 1st | 2nd | 3rd | 4th | 5th |
| --- | --- | --- | --- | --- | --- |
| 0 | 5 | 7 | 4 | 5 | 4 |
| 0.1 | 5 | 7 | 4 | 5 | 5 |
| 0.2 | 2 | 4 | 5 | 2 | 14 |
| 0.3 | 4 | 5 | 3 | 4 | 6 |
| 0.4 | 4 | 4 | 7 | 3 | 8 |
| 0.5 | 4 | 5 | 7 | 23 | 8 |
| 0.6 | 3 | 11 | 16 | 19 | 4 |
| 0.7 | 2 | 4 | 12 | 17 | 3 |
| 0.8 | 8 | 3 | 5 | 16 | 9 |
| 0.9 | 4 | 0 | 9 | 18 | 5 |
| 1 | 4 | 0 | 12 | 3 | 6 |
| 1.1 | 10 | 0 | 8 | 5 | 5 |
| 1.2 | 3 | 1 | 17 | 4 | 3 |
| 1.3 | 6 | 0 | 12 | 8 | 3 |
| 1.4 | 9 | 1 | 5 | 5 | 4 |
| 1.5 | 7 | 1 | 5 | 7 | 3 |
| 1.6 | 4 | 2 | 13 | 10 | 4 |
| 1.7 | 5 | 1 | 19 | 6 | 6 |
| 1.8 | 6 | 2 | 13 | 6 | 6 |
| 1.9 | 3 | 1 | 10 | 6 | 13 |
| 2 | 6 | 3 | 11 | 4 | 5 |
| 2.1 | 6 | 2 | 7 | 9 | 4 |
| 2.2 | 9 | 5 | 9 | 8 | 6 |
| 2.3 | 3 | 4 | 6 | 6 | 6 |
| 2.4 | 3 | 2 | 8 | 5 | 6 |
| 2.5 | 4 | 2 | 12 | 7 | 4 |
| 2.6 | 3 | 1 | 8 | 3 | 3 |
| 2.7 | 7 | 2 | 3 | 4 | 6 |
| 2.8 | 4 | 2 | 5 | 4 | 3 |
| 2.9 | 5 | 1 | 7 | 3 | 6 |
| 3 | 6 | 1 | 6 | 4 | 8 |
| 3.1 | 5 | 7 | 5 | 11 | 4 |
| 3.2 | 7 | 12 | 5 | 2 | 4 |
| 3.3 | 5 | 3 | 4 | 4 | 7 |
| 3.4 | 5 | 1 | 6 | 4 | 4 |
| 3.5 | 7 | 2 | 6 | 5 | 3 |
| 3.6 | 5 | 2 | 6 | 3 | 4 |
| 3.7 | 3 | 2 | 4 | 2 | 3 |
| 3.8 | 4 | 2 | 6 | 3 | 4 |
| 3.9 | 4 | 1 | 4 | 5 | 7 |
| 4 | 4 | 1 | 5 | 2 | 7 |
| 4.1 | 2 | 1 | 7 | 5 | 8 |
| 4.2 | 2 | 1 | 6 | 3 | 6 |
| 4.3 | 5 | 1 | 7 | 4 | 3 |
| 4.4 | 5 | 1 | 5 | 6 | 6 |
| 4.5 | 6 | 1 | 3 | 3 | 4 |
| 4.6 | 4 | 1 | 4 | 3 | 6 |
| 4.7 | 6 | 2 | 4 | 3 | 5 |
| 4.8 | 4 | 1 | 3 | 3 | 6 |
| 4.9 | 3 | 2 | 4 | 4 | 7 |
| 5 | 5 | 1 | 3 | 3 | 8 |
| 5.1 | 5 | 1 | 3 | 4 | 6 |
| 5.2 | 3 | 1 | 4 | 3 | 7 |
| 5.3 | 1 | 1 | 7 | 3 | 4 |
| 5.4 | 2 | 1 | 5 | 7 | 8 |
| 5.5 | 5 | 2 | 6 | 5 | 5 |
| 5.6 | 4 | 2 | 4 | 3 | 7 |
| 5.7 | 4 | 1 | 2 | 8 | 5 |
| 5.8 | 3 | 1 | 3 | 6 | 5 |
| 5.9 | 4 | 2 | 4 | 5 | 6 |
| 6 | 4 | 2 | 3 | 5 | 6 |
| 6.1 | 4 | 2 | 4 | 4 | 4 |
| 6.2 | 4 | 1 | 4 | 6 | 6 |
| 6.3 | 5 | 2 | 2 | 6 | 6 |
| 6.4 | 4 | 2 | 4 | 4 | 4 |
| 6.5 | 3 | 1 | 3 | 5 | 6 |
| 6.6 | 4 | 2 | 2 | 6 | 4 |
| 6.7 | 3 | 1 | 2 | 4 | 7 |
| 6.8 | 4 | 1 | 5 | 5 | 5 |
| 6.9 | 4 | 1 | 3 | 5 | 4 |
| 7 | 5 | 1 | 4 | 8 | 6 |
| 7.1 | 1 | 1 | 6 | 10 | 3 |
| 7.2 | 5 | 1 | 7 | 6 | 9 |
| 7.3 | 8 | 2 | 5 | 7 | 5 |
| 7.4 | 3 | 1 | 6 | 7 | 6 |
| 7.5 | 4 | 1 | 5 | 6 | 6 |
| 7.6 | 4 | 1 | 3 | 4 | 7 |
| 7.7 | 3 | 1 | 5 | 6 | 5 |
| 7.8 | 5 | 2 | 3 | 5 | 6 |
| 7.9 | 4 | 1 | 6 | 8 | 5 |
| 8 | 4 | 1 | 4 | 7 | 5 |
| 8.1 | 5 | 1 | 6 | 4 | 8 |
| 8.2 | 4 | 1 | 4 | 5 | 5 |
| 8.3 | 6 | 1 | 2 | 6 | 7 |
| 8.4 | 8 | 3 | 4 | 7 | 5 |
| 8.5 | 5 | 1 | 4 | 8 | 4 |
| 8.6 | 8 | 2 | 3 | 8 | 5 |
| 8.7 | 4 | 1 | 3 | 5 | 3 |
| 8.8 | 18 | 1 | 3 | 4 | 4 |
| 8.9 | 5 | 1 | 6 | 9 | 3 |
| 9 | 9 | 1 | 4 | 8 | 4 |
| 9.1 | 3 | 1 | 4 | 6 | 3 |
| 9.2 | 5 | 2 | 4 | 4 | 4 |
| 9.3 | 5 | 2 | 4 | 6 | 2 |
| 9.4 | 5 | 2 | 3 | 2 | 3 |
| 9.5 | 4 | 2 | 3 | 4 | 3 |
| 9.6 | 5 | 2 | 3 | 6 | 3 |
| 9.7 | 8 | 1 | 2 | 8 | 3 |
| 9.8 | 4 | 2 | 3 | 6 | 3 |
| 9.9 | 7 | 1 | 4 | 4 | 3 |
| 10 | 5 | 1 | 8 | 6 | 3 |
| 10.1 | 7 | 3 | 3 | 4 | 3 |
| 10.2 | 6 | 5 | 3 | 3 | 2 |
| 10.3 | 4 | 3 | 7 | 7 | 4 |
| 10.4 | 4 | 0 | 5 | 6 | 3 |
| 10.5 | 7 | 0 | 5 | 7 | 3 |
| 10.6 | 4 | 0 | 6 | 5 | 2 |
| 10.7 | 3 | 1 | 3 | 5 | 4 |
| 10.8 | 2 | 1 | 3 | 6 | 2 |
| 10.9 | 4 | 10 | 4 | 9 | 3 |
| 11 | 5 | 1 | 3 | 3 | 2 |
| 11.1 | 4 | 0 | 6 | 7 | 8 |
| 11.2 | 7 | 9 | 6 | 6 | 4 |
| 11.3 | 5 | 3 | 5 | 6 | 3 |
| 11.4 | 4 | 7 | 2 | 7 | 4 |
| 11.5 | 6 | 5 | 7 | 4 | 4 |
| 11.6 | 4 | 11 | 3 | 5 | 4 |
| 11.7 | 3 | 2 | 4 | 7 | 4 |
| 11.8 | 5 | 2 | 6 | 6 | 4 |
| 11.9 | 4 | 1 | 3 | 7 | 4 |
| 12 | 5 | 1 | 4 | 5 | 4 |
| 12.1 | 8 | 2 | 3 | 5 | 3 |
| 12.2 | 4 | 2 | 3 | 6 | 4 |
| 12.3 | 4 | 8 | 4 | 8 | 3 |
| 12.4 | 6 | 1 | 4 | 6 | 4 |
| 12.5 | 7 | 2 | 21 | 4 | 5 |
| 12.6 | 5 | 1 | 18 | 6 | 4 |
| 12.7 | 3 | 3 | 23 | 5 | 3 |
| 12.8 | 3 | 4 | 22 | 6 | 4 |
| 12.9 | 3 | 3 | 24 | 7 | 6 |
| 13 | 3 | 2 | 24 | 5 | 2 |
| 13.1 | 5 | 1 | 18 | 11 | 4 |
| 13.2 | 6 | 2 | 24 | 7 | 4 |
| 13.3 | 4 | 1 | 23 | 7 | 4 |
| 13.4 | 7 | 1 | 22 | 7 | 3 |
| 13.5 | 4 | 2 | 17 | 6 | 6 |
| 13.6 | 5 | 1 | 21 | 4 | 4 |
| 13.7 | 5 | 2 | 20 | 5 | 5 |
| 13.8 | 6 | 2 | 22 | 8 | 5 |
| 13.9 | 6 | 2 | 17 | 5 | 5 |
| 14 | 5 | 1 | 21 | 5 | 18 |
| 14.1 | 5 | 1 | 23 | 7 | 18 |
| 14.2 | 4 | 1 | 22 | 6 | 15 |
| 14.3 | 7 | 1 | 20 | 10 | 17 |
| 14.4 | 3 | 1 | 19 | 5 | 21 |
| 14.5 | 4 | 1 | 17 | 3 | 16 |
| 14.6 | 5 | 4 | 23 | 5 | 17 |
| 14.7 | 5 | 3 | 19 | 4 | 17 |
| 14.8 | 4 | 2 | 21 | 2 | 14 |
| 14.9 | 5 | 2 | 20 | 6 | 10 |
| 15 | 8 | 2 | 18 | 6 | 14 |
| 15.1 | 5 | 2 | 18 | 4 | 4 |
| 15.2 | 5 | 2 | 22 | 2 | 4 |
| 15.3 | 4 | 2 | 20 | 7 | 3 |
| 15.4 | 5 | 2 | 16 | 7 | 5 |
| 15.5 | 5 | 1 | 20 | 3 | 3 |
| 15.6 | 9 | 2 | 25 | 4 | 3 |
| 15.7 | 3 | 1 | 21 | 1 | 5 |
| 15.8 | 3 | 1 | 21 | 5 | 4 |
| 15.9 | 5 | 2 | 24 | 4 | 5 |
| 16 | 8 | 2 | 26 | 4 | 4 |
| 16.1 | 9 | 1 | 23 | 4 | 7 |
| 16.2 | 5 | 2 | 20 | 5 | 8 |
| 16.3 | 6 | 2 | 21 | 4 | 6 |
| 16.4 | 4 | 2 | 22 | 4 | 5 |
| 16.5 | 5 | 1 | 13 | 4 | 5 |
| 16.6 | 18 | 2 | 3 | 3 | 5 |
| 16.7 | 20 | 5 | 4 | 2 | 5 |
| 16.8 | 23 | 2 | 6 | 5 | 6 |
| 16.9 | 16 | 4 | 2 | 8 | 3 |
| 17 | 13 | 2 | 7 | 4 | 3 |
| 17.1 | 14 | 5 | 4 | 1 | 3 |
| 17.2 | 19 | 1 | 8 | 3 | 4 |
| 17.3 | 12 | 1 | 6 | 5 | 3 |
| 17.4 | 16 | 1 | 6 | 2 | 4 |
| 17.5 | 15 | 3 | 5 | 3 | 3 |
| 17.6 | 14 | 1 | 6 | 4 | 3 |
| 17.7 | 4 | 8 | 4 | 6 | 5 |
| 17.8 | 5 | 5 | 8 | 4 | 4 |
| 17.9 | 5 | 3 | 6 | 3 | 4 |
| 18 | 12 | 6 | 9 | 4 | 2 |
| 18.1 | 8 | 1 | 8 | 6 | 3 |
| 18.2 | 9 | 2 | 10 | 7 | 4 |
| 18.3 | 4 | 4 | 6 | 5 | 3 |
| 18.4 | 6 | 2 | 16 | 4 | 2 |
| 18.5 | 6 | 7 | 8 | 4 | 3 |
| 18.6 | 5 | 7 | 5 | 5 | 3 |
| 18.7 | 6 | 1 | 6 | 5 | 6 |
| 18.8 | 4 | 2 | 7 | 6 | 7 |
| 18.9 | 7 | 2 | 10 | 3 | 5 |
| 19 | 6 | 4 | 6 | 4 | 3 |
| 19.1 | 4 | 4 | 6 | 6 | 1 |
| 19.2 | 6 | 4 | 8 | 6 | 2 |
| 19.3 | 5 | 5 | 7 | 4 | 2 |
| 19.4 | 5 | 2 | 5 | 4 | 1 |
| 19.5 | 4 | 5 | 6 | 4 | 1 |
| 19.6 | 5 | 2 | 4 | 6 | 1 |
| 19.7 | 6 | 2 | 4 | 12 | 3 |
| 19.8 | 5 | 4 | 7 | 2 | 1 |
| 19.9 | 5 | 2 | 6 | 3 | 1 |

**Figure S3.** Timeline plots pertaining to number of contacts between the E-box and BMAL1 bHLH for the systems studied.

ARG-10, ARG-11, ASP-21 (nonphosphorylated)

Angle

| Frame | 1st | 2nd | 3rd | 4th | 5th |
| --- | --- | --- | --- | --- | --- |
| 0 | 69.879 | 65.4462 | 70.0504 | 80.0107 | 71.7256 |
| 1 | 69.879 | 65.4462 | 70.0504 | 80.0107 | 71.7256 |
| 2 | 69.879 | 65.4462 | 70.0504 | 80.0186 | 61.7798 |
| 3 | 69.879 | 65.4462 | 62.3609 | 76.3562 | 61.9316 |
| 4 | 66.5262 | 65.4462 | 66.1342 | 69.4906 | 68.2314 |
| 5 | 70.752 | 78.9657 | 60.773 | 68.4975 | 71.4541 |
| 6 | 70.5761 | 76.7589 | 62.6506 | 61.2103 | 68.4048 |
| 7 | 70.95 | 72.6825 | 72.6165 | 73.9996 | 71.7563 |
| 8 | 72.4816 | 80.1191 | 78.7536 | 68.6518 | 71.1388 |
| 9 | 68.8724 | 74.8289 | 64.1208 | 62.953 | 70.3961 |
| 10 | 69.2134 | 78.7672 | 73.1167 | 62.4916 | 69.3499 |
| 11 | 69.5209 | 80.951 | 63.1136 | 74.0751 | 68.0303 |
| 12 | 75.2368 | 85.3047 | 67.3336 | 71.0636 | 69.5661 |
| 13 | 68.7314 | 81.1899 | 74.1104 | 62.8399 | 71.4584 |
| 14 | 69.7423 | 80.4762 | 71.1813 | 56.5301 | 73.4091 |
| 15 | 71.2523 | 71.7413 | 73.0172 | 63.2982 | 66.2433 |
| 16 | 69.9877 | 75.0553 | 65.8184 | 65.4646 | 69.5019 |
| 17 | 65.8271 | 79.3403 | 64.5343 | 65.0045 | 62.6951 |
| 18 | 63.7479 | 75.275 | 65.867 | 64.4878 | 75.9567 |
| 19 | 67.3097 | 70.34 | 66.2716 | 48.2612 | 71.9907 |
| 20 | 69.0566 | 82.0699 | 75.0883 | 45.1611 | 72.4199 |
| 21 | 65.6488 | 76.9399 | 72.9911 | 52.7556 | 66.1372 |
| 22 | 67.0369 | 74.7047 | 67.923 | 53.4232 | 70.4015 |
| 23 | 67.0179 | 83.0789 | 67.2769 | 55.905 | 67.4102 |
| 24 | 64.4411 | 75.2357 | 71.1539 | 64.3717 | 66.3226 |
| 25 | 67.9987 | 74.5719 | 63.5892 | 73.4907 | 72.9303 |
| 26 | 64.6909 | 75.0607 | 76.3161 | 58.8529 | 70.5668 |
| 27 | 66.8681 | 79.2932 | 82.5453 | 64.1142 | 70.7853 |
| 28 | 87.7905 | 73.6712 | 66.8253 | 67.8839 | 65.4157 |
| 29 | 74.4321 | 76.8609 | 64.1612 | 60.2936 | 68.0102 |
| 30 | 93.1113 | 76.9055 | 67.5763 | 66.5551 | 71.3917 |
| 31 | 77.1434 | 78.0547 | 65.3777 | 64.0915 | 69.1963 |
| 32 | 80.6888 | 72.3304 | 71.1652 | 61.0539 | 66.2755 |
| 33 | 78.687 | 72.65 | 70.4469 | 60.5164 | 67.8383 |
| 34 | 67.3823 | 67.5675 | 71.7513 | 60.9771 | 67.542 |
| 35 | 70.1006 | 71.1979 | 71.4319 | 62.836 | 65.1888 |
| 36 | 69.6543 | 67.8665 | 57.963 | 63.5532 | 63.2166 |
| 37 | 73.607 | 73.5531 | 73.9511 | 62.0205 | 59.0524 |
| 38 | 67.0677 | 71.1376 | 69.0878 | 57.5891 | 60.8849 |
| 39 | 55.6323 | 72.3811 | 65.8151 | 59.629 | 64.0355 |
| 40 | 54.657 | 71.937 | 78.2826 | 53.1413 | 62.0729 |
| 41 | 76.8581 | 73.881 | 66.1268 | 62.9962 | 64.2653 |
| 42 | 67.9588 | 72.9239 | 77.3127 | 55.2788 | 64.9264 |
| 43 | 75.065 | 76.2313 | 64.0879 | 65.3923 | 65.4739 |
| 44 | 79.6742 | 70.0891 | 65.7903 | 53.8853 | 66.0805 |
| 45 | 70.8892 | 69.443 | 67.0194 | 51.1747 | 56.6656 |
| 46 | 68.6502 | 75.0952 | 71.2268 | 67.7587 | 58.782 |
| 47 | 69.7523 | 73.0529 | 69.8089 | 60.6154 | 58.9431 |
| 48 | 65.2558 | 82.8349 | 69.4878 | 68.7304 | 61.238 |
| 49 | 79.1307 | 79.3862 | 75.6352 | 69.158 | 59.9535 |
| 50 | 80.6048 | 84.41 | 69.9064 | 65.0576 | 55.9148 |
| 51 | 79.8926 | 77.9935 | 80.2369 | 52.56 | 59.1482 |
| 52 | 83.2554 | 74.2638 | 78.3377 | 53.6017 | 57.1301 |
| 53 | 79.3588 | 68.5206 | 76.2089 | 57.8672 | 57.0255 |
| 54 | 75.1712 | 73.3752 | 73.9118 | 56.7637 | 56.4453 |
| 55 | 72.5956 | 73.4391 | 81.1582 | 63.0107 | 63.5412 |
| 56 | 73.2662 | 73.8978 | 76.324 | 70.3493 | 60.1145 |
| 57 | 67.4292 | 72.1787 | 76.7742 | 66.3651 | 62.2356 |
| 58 | 72.8077 | 73.6079 | 70.4347 | 59.9269 | 66.6849 |
| 59 | 75.6645 | 64.9206 | 80.8731 | 55.2919 | 58.7295 |
| 60 | 69.9904 | 70.0769 | 79.834 | 65.4842 | 59.0826 |
| 61 | 69.9495 | 61.9405 | 78.6887 | 54.7174 | 55.8664 |
| 62 | 72.22 | 69.1669 | 70.2085 | 57.5085 | 57.9735 |
| 63 | 78.914 | 76.4456 | 80.539 | 61.0068 | 57.1369 |
| 64 | 72.4705 | 65.8944 | 81.2653 | 69.9399 | 58.3753 |
| 65 | 80.8574 | 72.7761 | 68.236 | 71.1574 | 57.6392 |
| 66 | 68.3659 | 70.2776 | 83.3579 | 58.4554 | 56.3981 |
| 67 | 71.5966 | 64.6062 | 75.6622 | 54.7967 | 56.3679 |
| 68 | 76.5192 | 70.7763 | 75.1908 | 60.9634 | 61.0573 |
| 69 | 69.8751 | 70.458 | 73.4078 | 81.3077 | 59.1652 |
| 70 | 67.3462 | 72.7313 | 74.1061 | 67.1758 | 56.6351 |
| 71 | 72.7107 | 66.4382 | 74.6027 | 69.0363 | 60.9205 |
| 72 | 69.8299 | 67.5378 | 73.6316 | 70.4368 | 64.0609 |
| 73 | 71.0843 | 73.9503 | 78.6048 | 70.9685 | 59.6012 |
| 74 | 71.5327 | 71.0875 | 70.7537 | 68.7241 | 59.7951 |
| 75 | 74.8694 | 66.6185 | 69.6225 | 75.4192 | 61.5285 |
| 76 | 73.7235 | 68.9397 | 73.5649 | 68.5112 | 64.6959 |
| 77 | 64.8518 | 71.138 | 74.5289 | 72.5667 | 59.6469 |
| 78 | 68.1975 | 67.7453 | 84.8738 | 71.3398 | 57.4277 |
| 79 | 63.5047 | 69.7248 | 85.1055 | 75.1858 | 58.4187 |
| 80 | 64.711 | 67.7948 | 82.2835 | 76.5775 | 55.5917 |
| 81 | 77.3596 | 66.3535 | 83.4234 | 68.7634 | 56.8499 |
| 82 | 69.4389 | 76.5311 | 83.2803 | 77.8527 | 55.8291 |
| 83 | 65.7202 | 67.2334 | 85.6658 | 61.0131 | 57.8395 |
| 84 | 75.3026 | 71.1866 | 73.6621 | 64.5382 | 57.3993 |
| 85 | 57.5986 | 62.4148 | 75.3029 | 63.312 | 59.5551 |
| 86 | 60.9808 | 66.8943 | 72.2968 | 72.0442 | 58.5905 |
| 87 | 77.2673 | 65.5838 | 71.8773 | 75.3849 | 58.4661 |
| 88 | 68.7426 | 65.3341 | 63.4172 | 69.7379 | 62.5354 |
| 89 | 85.1661 | 63.4063 | 69.0499 | 67.0062 | 58.8439 |
| 90 | 74.0415 | 61.7249 | 73.9701 | 70.3933 | 57.1938 |
| 91 | 62.9478 | 55.8156 | 71.5095 | 73.6426 | 56.2743 |
| 92 | 63.4657 | 61.2716 | 71.4075 | 68.5769 | 59.4255 |
| 93 | 61.3808 | 60.6068 | 69.0181 | 75.4534 | 58.2394 |
| 94 | 56.5794 | 62.9184 | 67.2455 | 81.0433 | 60.0082 |
| 95 | 55.8602 | 58.2792 | 71.2853 | 73.5079 | 61.1061 |
| 96 | 55.4451 | 63.3306 | 67.3799 | 78.2244 | 58.4091 |
| 97 | 59.3488 | 62.2775 | 72.1607 | 74.5706 | 55.8607 |
| 98 | 58.3574 | 60.2837 | 67.515 | 72.7226 | 56.9548 |
| 99 | 56.9345 | 58.9908 | 71.3635 | 75.5722 | 58.5703 |
| 100 | 62.3378 | 63.4661 | 73.9247 | 70.9116 | 59.7479 |
| 101 | 69.0082 | 56.304 | 63.4535 | 67.7864 | 61.9879 |
| 102 | 68.3799 | 55.9217 | 63.7195 | 77.0844 | 62.851 |
| 103 | 59.2843 | 58.8336 | 65.5931 | 68.4358 | 60.6264 |
| 104 | 66.6889 | 63.7485 | 71.619 | 69.6407 | 58.1943 |
| 105 | 71.0736 | 65.4462 | 72.7721 | 67.5408 | 56.177 |
| 106 | 67.8837 | 54.9155 | 66.2494 | 66.739 | 58.8387 |
| 107 | 66.9075 | 58.8742 | 68.2704 | 69.3604 | 57.4421 |
| 108 | 60.1187 | 59.925 | 69.3868 | 75.3117 | 62.2548 |
| 109 | 54.1495 | 58.549 | 57.8208 | 71.1631 | 59.9881 |
| 110 | 56.4123 | 53.25 | 67.9253 | 69.0854 | 58.3441 |
| 111 | 66.5528 | 54.8656 | 79.8654 | 76.3656 | 60.0845 |
| 112 | 71.2144 | 57.3059 | 71.606 | 76.1484 | 63.3544 |
| 113 | 65.5689 | 54.7499 | 65.3056 | 73.9433 | 58.3561 |
| 114 | 69.9334 | 55.687 | 68.5439 | 77.8903 | 59.0005 |
| 115 | 71.7295 | 56.7822 | 68.9397 | 76.1674 | 59.006 |
| 116 | 50.699 | 57.3667 | 67.8499 | 69.7872 | 59.8448 |
| 117 | 55.9707 | 58.1389 | 68.521 | 79.9424 | 60.2319 |
| 118 | 52.3042 | 52.9752 | 64.744 | 69.6855 | 64.4974 |
| 119 | 69.5672 | 54.0527 | 64.1167 | 73.5843 | 60.8081 |
| 120 | 73.6609 | 57.7816 | 70.2291 | 73.1066 | 59.1625 |
| 121 | 66.8498 | 54.9357 | 68.7433 | 77.6854 | 58.5739 |
| 122 | 71.5153 | 55.6149 | 68.108 | 75.1931 | 59.2869 |
| 123 | 68.5888 | 53.3372 | 77.7126 | 75.9841 | 61.4759 |
| 124 | 77.2293 | 58.0614 | 70.8315 | 63.7894 | 63.5967 |
| 125 | 72.1024 | 53.6384 | 70.1214 | 74.4028 | 60.1451 |
| 126 | 67.7836 | 56.9287 | 71.1988 | 65.3994 | 57.6164 |
| 127 | 67.0247 | 53.3778 | 65.057 | 68.4443 | 57.3721 |
| 128 | 66.5422 | 49.9702 | 67.5803 | 67.9781 | 61.1442 |
| 129 | 67.6581 | 49.8318 | 66.7322 | 72.7028 | 57.9337 |
| 130 | 64.9306 | 53.7279 | 71.1577 | 72.1226 | 63.8521 |
| 131 | 62.4304 | 48.9197 | 69.378 | 75.4197 | 62.0277 |
| 132 | 61.0749 | 51.7308 | 68.8987 | 72.5979 | 61.2938 |
| 133 | 55.4727 | 51.4158 | 63.8814 | 76.6277 | 60.1326 |
| 134 | 57.1584 | 54.8744 | 60.504 | 74.2585 | 60.8079 |
| 135 | 70.6894 | 58.7374 | 59.4985 | 67.2411 | 64.0533 |
| 136 | 69.2369 | 52.8334 | 63.7388 | 69.2673 | 59.8528 |
| 137 | 66.8006 | 53.0711 | 52.9863 | 72.3073 | 61.9764 |
| 138 | 78.0783 | 57.2575 | 56.7371 | 73.5482 | 57.1899 |
| 139 | 72.9839 | 54.9108 | 57.1248 | 63.0287 | 61.023 |
| 140 | 73.6203 | 54.7866 | 70.1889 | 72.1377 | 58.2369 |
| 141 | 69.2001 | 57.5131 | 56.6415 | 72.8291 | 58.4472 |
| 142 | 65.1881 | 52.2598 | 64.4204 | 71.3443 | 59.1935 |
| 143 | 69.0807 | 53.2654 | 67.9536 | 76.1034 | 58.4817 |
| 144 | 62.2642 | 58.9325 | 61.3669 | 70.3042 | 63.3291 |
| 145 | 71.0355 | 58.9007 | 61.4452 | 70.655 | 59.6734 |
| 146 | 71.3918 | 57.8819 | 64.5005 | 69.0594 | 59.3286 |
| 147 | 74.0825 | 56.0159 | 61.6225 | 70.2302 | 59.1854 |
| 148 | 63.739 | 58.9496 | 64.3819 | 70.6638 | 59.1497 |
| 149 | 56.899 | 58.5799 | 57.4309 | 77.8613 | 62.6468 |
| 150 | 56.2066 | 57.7484 | 59.6623 | 72.623 | 62.6974 |
| 151 | 55.8942 | 56.6717 | 54.5923 | 70.5544 | 60.1623 |
| 152 | 56.8873 | 57.845 | 63.7087 | 67.4426 | 56.5956 |
| 153 | 58.8247 | 55.8964 | 57.3073 | 65.2179 | 60.3848 |
| 154 | 63.9578 | 60.1536 | 62.9343 | 70.8699 | 60.7513 |
| 155 | 62.4284 | 59.9994 | 69.374 | 73.8206 | 59.366 |
| 156 | 69.1242 | 59.7521 | 62.5471 | 71.3858 | 58.6744 |
| 157 | 68.7389 | 57.5756 | 69.9549 | 77.5667 | 60.7968 |
| 158 | 66.5098 | 61.2783 | 59.8578 | 70.8706 | 59.1629 |
| 159 | 66.5961 | 55.9914 | 65.0334 | 69.7382 | 58.7675 |
| 160 | 63.0091 | 51.1891 | 66.6589 | 70.684 | 59.3231 |
| 161 | 71.463 | 56.7224 | 60.3027 | 73.6141 | 57.9395 |
| 162 | 63.0853 | 57.2074 | 65.7583 | 70.3099 | 59.6645 |
| 163 | 57.335 | 55.5167 | 62.4913 | 71.872 | 59.1588 |
| 164 | 72.413 | 59.3047 | 60.3073 | 69.6273 | 58.8479 |
| 165 | 67.9283 | 55.5847 | 63.5991 | 69.3429 | 58.8056 |
| 166 | 64.6748 | 56.8997 | 69.0945 | 70.6095 | 62.4414 |
| 167 | 65.399 | 56.9315 | 69.1897 | 60.5356 | 60.0668 |
| 168 | 66.6917 | 56.4823 | 72.1678 | 70.8779 | 58.7405 |
| 169 | 70.1662 | 57.6657 | 66.7351 | 68.1328 | 58.5057 |
| 170 | 66.2113 | 54.5445 | 55.0833 | 67.8705 | 60.7068 |
| 171 | 56.6835 | 57.9984 | 61.7691 | 79.151 | 58.1624 |
| 172 | 62.0541 | 58.5299 | 60.346 | 72.3579 | 60.2015 |
| 173 | 64.1476 | 54.5694 | 63.9387 | 70.3874 | 59.3534 |
| 174 | 68.1085 | 54.886 | 64.3247 | 70.7171 | 59.7293 |
| 175 | 68.0917 | 59.9936 | 68.6867 | 60.1246 | 60.1236 |
| 176 | 68.2988 | 55.6027 | 71.7126 | 66.5765 | 58.0855 |
| 177 | 72.98 | 56.2152 | 68.3346 | 67.644 | 61.468 |
| 178 | 57.0338 | 58.0423 | 73.4008 | 69.3906 | 58.9433 |
| 179 | 60.3704 | 56.5929 | 71.2418 | 72.8001 | 59.3954 |
| 180 | 71.9046 | 56.8413 | 67.3453 | 69.7962 | 59.0473 |
| 181 | 70.481 | 59.1275 | 71.5711 | 73.784 | 59.1052 |
| 182 | 61.5456 | 59.1224 | 69.1 | 69.2819 | 59.8866 |
| 183 | 68.4941 | 58.3095 | 67.3399 | 64.4032 | 59.0334 |
| 184 | 66.346 | 53.3307 | 73.932 | 64.2307 | 59.1261 |
| 185 | 66.3735 | 58.5396 | 68.1029 | 63.3756 | 62.17 |
| 186 | 62.9862 | 60.1315 | 68.2743 | 65.7245 | 60.8423 |
| 187 | 64.7095 | 54.18 | 69.0965 | 63.8802 | 58.2468 |
| 188 | 63.2747 | 56.335 | 67.0659 | 75.0695 | 59.698 |
| 189 | 65.5294 | 53.5317 | 68.4232 | 66.4825 | 60.1127 |
| 190 | 72.3391 | 56.3242 | 62.9523 | 64.4415 | 61.0819 |
| 191 | 65.5411 | 62.678 | 61.1982 | 63.4646 | 55.538 |
| 192 | 62.645 | 54.3939 | 60.3697 | 67.0134 | 59.0511 |
| 193 | 69.5918 | 58.4633 | 56.917 | 62.8004 | 65.6232 |
| 194 | 62.6767 | 50.7839 | 62.0133 | 64.5562 | 62.4996 |
| 195 | 54.0431 | 54.9439 | 58.1542 | 67.7146 | 57.7628 |
| 196 | 63.3766 | 58.1581 | 59.8649 | 71.4753 | 57.9986 |
| 197 | 66.9611 | 60.9182 | 56.6106 | 67.4289 | 57.3062 |
| 198 | 66.0123 | 58.1634 | 51.9319 | 59.026 | 59.6013 |
| 199 | 47.3415 | 55.3962 | 61.9764 | 65.7156 | 63.7495 |

ARG-10, ARG-11, ASP-21 (BMAL1S78)

Angle

| Frame | 1st | 2nd | 3rd | 4th | 5th |
| --- | --- | --- | --- | --- | --- |
| 0 | 6.20661 | 9.81246 | 107.406 | 2.18575 | 7.93413 |
| 1 | 6.20661 | 9.81246 | 107.406 | 72.0287 | 7.93412 |
| 2 | 79.2447 | 73.959 | 66.253 | 76.4188 | 71.3919 |
| 3 | 77.7689 | 74.0572 | 69.9228 | 76.722 | 58.5704 |
| 4 | 70.1027 | 81.2023 | 81.9228 | 88.5825 | 45.7402 |
| 5 | 65.4718 | 72.407 | 73.4815 | 76.6379 | 64.3434 |
| 6 | 67.1493 | 61.6278 | 70.3489 | 81.8271 | 58.7786 |
| 7 | 69.7205 | 66.806 | 61.2005 | 77.4815 | 59.3963 |
| 8 | 68.991 | 60.2263 | 65.5508 | 81.2737 | 61.2547 |
| 9 | 62.8389 | 59.8079 | 61.8313 | 85.8929 | 57.7262 |
| 10 | 54.4691 | 58.0344 | 65.7041 | 81.5005 | 57.3178 |
| 11 | 61.0439 | 53.8159 | 67.1482 | 78.2242 | 57.9666 |
| 12 | 56.9454 | 61.8727 | 57.2924 | 77.0827 | 57.5896 |
| 13 | 50.5685 | 72.3501 | 67.084 | 75.1781 | 60.657 |
| 14 | 55.588 | 80.9735 | 58.2952 | 75.5529 | 56.2328 |
| 15 | 49.9806 | 62.5398 | 58.151 | 73.5016 | 62.5594 |
| 16 | 54.0067 | 63.7613 | 48.6986 | 75.6471 | 70.1519 |
| 17 | 53.3691 | 60.9829 | 51.8428 | 74.4855 | 74.9799 |
| 18 | 57.3938 | 59.6542 | 55.0348 | 69.0756 | 73.3123 |
| 19 | 51.2939 | 54.9384 | 44.5353 | 70.6583 | 69.5318 |
| 20 | 51.3695 | 55.2652 | 51.4026 | 69.7858 | 81.0022 |
| 21 | 59.7679 | 55.2843 | 51.6615 | 75.6121 | 65.6726 |
| 22 | 60.7056 | 61.975 | 55.5713 | 72.5902 | 59.0639 |
| 23 | 59.1653 | 61.1853 | 49.3966 | 72.7234 | 68.4195 |
| 24 | 50.0745 | 60.7992 | 51.3077 | 69.0865 | 63.0397 |
| 25 | 60.8412 | 58.9289 | 48.5285 | 65.9199 | 66.4645 |
| 26 | 60.8021 | 62.5806 | 48.0531 | 66.9987 | 56.4025 |
| 27 | 62.5318 | 76.3682 | 55.1731 | 70.9658 | 62.1701 |
| 28 | 61.4784 | 52.3112 | 51.8254 | 69.7186 | 66.6167 |
| 29 | 58.3839 | 56.7159 | 55.4164 | 68.0891 | 66.1783 |
| 30 | 64.6314 | 58.0798 | 58.4768 | 60.6934 | 58.2904 |
| 31 | 61.0749 | 54.6071 | 57.6595 | 70.6218 | 60.0149 |
| 32 | 64.729 | 60.6541 | 52.6404 | 62.1611 | 58.71 |
| 33 | 56.2115 | 61.1249 | 58.8638 | 66.006 | 60.0039 |
| 34 | 56.7119 | 57.9081 | 45.5284 | 64.9281 | 64.4156 |
| 35 | 61.0084 | 58.4409 | 53.3264 | 62.7564 | 69.2938 |
| 36 | 58.0201 | 58.7569 | 46.3076 | 75.0969 | 69.3097 |
| 37 | 57.0188 | 63.5311 | 54.6594 | 69.4053 | 67.66 |
| 38 | 55.9188 | 57.0188 | 45.95 | 69.325 | 68.0575 |
| 39 | 50.0508 | 58.9584 | 48.5307 | 65.3823 | 60.6199 |
| 40 | 57.2052 | 51.4128 | 50.5674 | 67.578 | 60.6677 |
| 41 | 57.3067 | 58.7144 | 49.8118 | 64.1219 | 63.4873 |
| 42 | 54.1188 | 64.6014 | 50.8175 | 68.7298 | 65.3158 |
| 43 | 63.0837 | 56.2765 | 53.0174 | 66.9881 | 62.9013 |
| 44 | 56.9399 | 63.4142 | 49.7973 | 68.2221 | 61.1476 |
| 45 | 58.6469 | 75.1768 | 53.5264 | 73.5807 | 67.7324 |
| 46 | 61.8283 | 67.0506 | 52.3761 | 70.7369 | 56.0505 |
| 47 | 53.6842 | 67.0914 | 44.514 | 69.1173 | 64.2744 |
| 48 | 58.0646 | 66.4208 | 49.6972 | 65.637 | 60.5131 |
| 49 | 58.7579 | 64.5685 | 44.9568 | 67.6279 | 65.4739 |
| 50 | 58.9039 | 57.6365 | 48.6015 | 64.4745 | 66.3108 |
| 51 | 48.176 | 66.4535 | 42.2138 | 64.4187 | 63.6082 |
| 52 | 60.3502 | 63.0602 | 47.9626 | 53.6375 | 52.4469 |
| 53 | 60.9831 | 63.4442 | 47.0338 | 66.1172 | 49.6807 |
| 54 | 52.8422 | 66.6839 | 46.874 | 68.7882 | 50.4769 |
| 55 | 56.1057 | 60.7531 | 51.7597 | 70.1566 | 51.6254 |
| 56 | 57.8882 | 70.0455 | 54.725 | 63.4473 | 52.1024 |
| 57 | 64.1815 | 71.5069 | 52.5382 | 69.2585 | 48.0427 |
| 58 | 54.6157 | 62.9237 | 55.7505 | 73.0083 | 49.3247 |
| 59 | 65.4143 | 75.1123 | 57.8112 | 76.5501 | 56.7472 |
| 60 | 62.9824 | 71.1357 | 59.6865 | 72.1835 | 48.8494 |
| 61 | 65.8831 | 63.858 | 46.1974 | 68.6506 | 49.6464 |
| 62 | 59.0593 | 69.2672 | 46.5173 | 67.0035 | 49.0533 |
| 63 | 63.6417 | 62.2584 | 59.1986 | 72.6118 | 47.2798 |
| 64 | 62.7348 | 62.5722 | 59.0347 | 65.1173 | 38.1309 |
| 65 | 62.5403 | 67.9447 | 50.9407 | 72.9714 | 38.6335 |
| 66 | 58.4125 | 68.4366 | 52.9915 | 69.1817 | 43.9292 |
| 67 | 65.3952 | 65.111 | 51.7883 | 63.7808 | 47.4304 |
| 68 | 60.6871 | 68.7367 | 49.0957 | 59.7187 | 48.3408 |
| 69 | 61.1676 | 64.7591 | 50.4272 | 71.3203 | 53.1637 |
| 70 | 55.699 | 72.6994 | 55.4466 | 71.0593 | 48.0653 |
| 71 | 55.5938 | 78.3302 | 42.591 | 61.7552 | 45.4623 |
| 72 | 56.227 | 73.5104 | 48.6393 | 69.8517 | 50.1065 |
| 73 | 59.5372 | 64.1589 | 43.1986 | 62.8544 | 49.5 |
| 74 | 58.7791 | 69.8835 | 45.9636 | 67.5022 | 46.8198 |
| 75 | 59.3139 | 73.9653 | 42.4886 | 63.7818 | 47.1106 |
| 76 | 58.7045 | 64.7113 | 40.6127 | 64.8825 | 50.9557 |
| 77 | 59.7489 | 67.3652 | 45.9732 | 68.3125 | 48.7847 |
| 78 | 53.5197 | 73.7856 | 45.6696 | 68.226 | 47.5871 |
| 79 | 57.8338 | 68.3156 | 46.6319 | 63.7718 | 52.1372 |
| 80 | 57.8078 | 74.6688 | 45.4449 | 65.6582 | 48.3292 |
| 81 | 55.2965 | 69.7967 | 53.0502 | 68.5649 | 47.0839 |
| 82 | 61.4488 | 65.2105 | 52.6279 | 67.1547 | 43.715 |
| 83 | 57.5221 | 70.2154 | 43.0559 | 69.151 | 53.594 |
| 84 | 62.7524 | 65.3535 | 52.9153 | 62.4669 | 49.9281 |
| 85 | 58.4631 | 69.1956 | 53.9139 | 60.6912 | 46.8222 |
| 86 | 56.5109 | 69.3051 | 56.2136 | 57.5638 | 45.831 |
| 87 | 66.4109 | 74.5613 | 56.2907 | 59.5832 | 48.4232 |
| 88 | 59.2546 | 72.3014 | 51.9203 | 60.6376 | 41.5496 |
| 89 | 61.218 | 73.3573 | 50.046 | 63.7134 | 47.6179 |
| 90 | 60.4119 | 70.3144 | 51.1025 | 60.8138 | 45.8319 |
| 91 | 57.9611 | 63.0082 | 52.6384 | 60.6281 | 46.6589 |
| 92 | 54.0704 | 70.2403 | 47.2415 | 60.4631 | 46.1659 |
| 93 | 60.1676 | 74.402 | 53.0118 | 59.1527 | 45.7679 |
| 94 | 58.6732 | 67.8835 | 44.121 | 58.0728 | 48.525 |
| 95 | 60.1075 | 74.6885 | 52.712 | 59.4165 | 50.245 |
| 96 | 61.4919 | 68.2424 | 50.7418 | 59.9033 | 48.0988 |
| 97 | 57.385 | 69.3425 | 47.7209 | 57.4957 | 48.8388 |
| 98 | 54.5678 | 66.7521 | 43.7206 | 55.7532 | 50.6812 |
| 99 | 57.7119 | 76.1207 | 47.9042 | 75.0122 | 42.6773 |
| 100 | 59.0453 | 69.9674 | 43.7842 | 51.8473 | 41.4056 |
| 101 | 56.408 | 72.0025 | 46.0136 | 62.4692 | 35.812 |
| 102 | 58.4795 | 66.5258 | 45.0317 | 63.7623 | 50.4214 |
| 103 | 50.8792 | 63.3131 | 48.7594 | 60.7953 | 42.0993 |
| 104 | 58.433 | 70.5298 | 44.3559 | 55.7772 | 45.7051 |
| 105 | 52.0929 | 67.6931 | 45.1764 | 62.1438 | 44.85 |
| 106 | 60.3748 | 72.4778 | 49.9377 | 57.6345 | 54.0306 |
| 107 | 57.5558 | 70.5334 | 48.4929 | 61.4457 | 47.3407 |
| 108 | 58.1245 | 73.69 | 46.0413 | 59.7819 | 46.1965 |
| 109 | 60.2709 | 70.1689 | 40.4789 | 66.8234 | 41.3888 |
| 110 | 57.2549 | 67.8378 | 52.94 | 69.3709 | 43.2027 |
| 111 | 66.5554 | 66.7446 | 48.9555 | 65.624 | 46.3321 |
| 112 | 60.079 | 70.2505 | 47.2873 | 69.4011 | 46.6949 |
| 113 | 50.726 | 68.6531 | 52.2183 | 65.4602 | 47.2609 |
| 114 | 60.0397 | 76.5468 | 60.0958 | 66.032 | 45.8294 |
| 115 | 59.0834 | 70.7692 | 46.3312 | 66.6193 | 44.6793 |
| 116 | 60.1413 | 75.576 | 52.3133 | 66.498 | 39.7918 |
| 117 | 59.1007 | 64.2631 | 55.3223 | 61.8147 | 44.19 |
| 118 | 58.7195 | 70.3854 | 57.98 | 67.1716 | 44.78 |
| 119 | 61.8608 | 73.3079 | 58.4687 | 71.6169 | 41.4776 |
| 120 | 57.057 | 73.261 | 51.9574 | 75.7434 | 43.5027 |
| 121 | 57.9024 | 72.2317 | 52.2284 | 71.592 | 45.6464 |
| 122 | 66.6858 | 78.009 | 47.5664 | 72.1115 | 42.9831 |
| 123 | 61.2674 | 72.016 | 51.1967 | 71.7734 | 35.6007 |
| 124 | 62.7155 | 71.9935 | 45.3978 | 70.8089 | 38.2126 |
| 125 | 54.3444 | 74.3024 | 49.2381 | 70.1689 | 44.0183 |
| 126 | 57.7106 | 67.4546 | 47.8191 | 69.7793 | 43.5013 |
| 127 | 60.8733 | 72.9618 | 47.6897 | 70.6807 | 41.5478 |
| 128 | 58.6346 | 73.2155 | 49.0512 | 73.8829 | 44.6672 |
| 129 | 58.5999 | 75.1396 | 47.5026 | 70.7327 | 48.3326 |
| 130 | 60.9295 | 69.083 | 45.2529 | 64.1456 | 43.1924 |
| 131 | 63.728 | 76.5761 | 46.8655 | 68.1352 | 45.3602 |
| 132 | 69.7261 | 70.7935 | 47.9171 | 66.3488 | 37.7861 |
| 133 | 61.084 | 65.9467 | 51.2456 | 71.7688 | 40.1006 |
| 134 | 64.6332 | 70.5004 | 53.4019 | 62.8012 | 44.5053 |
| 135 | 67.082 | 78.2023 | 47.9353 | 68.0632 | 54.1527 |
| 136 | 65.0009 | 73.5331 | 53.5838 | 77.6316 | 55.0682 |
| 137 | 60.8892 | 71.0791 | 50.0758 | 80.2242 | 54.005 |
| 138 | 64.4182 | 72.1753 | 48.1876 | 63.7343 | 52.717 |
| 139 | 63.0922 | 76.6868 | 45.4008 | 61.6165 | 56.5423 |
| 140 | 61.8479 | 66.4696 | 50.0485 | 62.7776 | 52.0229 |
| 141 | 60.8426 | 70.5033 | 51.4567 | 68.7591 | 54.0963 |
| 142 | 64.6567 | 72.483 | 52.8725 | 62.6916 | 56.9379 |
| 143 | 66.6795 | 74.2432 | 49.0064 | 72.6249 | 53.855 |
| 144 | 66.9033 | 72.4522 | 52.6502 | 73.8049 | 55.5522 |
| 145 | 51.675 | 74.0999 | 52.5366 | 65.6544 | 51.7016 |
| 146 | 59.072 | 78.029 | 48.7954 | 61.584 | 46.8515 |
| 147 | 59.7826 | 75.1109 | 46.1873 | 62.9762 | 38.8969 |
| 148 | 61.2275 | 72.0281 | 51.6014 | 58.5093 | 53.6302 |
| 149 | 61.205 | 72.3201 | 54.0752 | 68.8919 | 46.965 |
| 150 | 63.7976 | 68.0422 | 49.8533 | 67.3755 | 41.1897 |
| 151 | 66.9662 | 73.8195 | 51.9282 | 63.5248 | 40.4931 |
| 152 | 59.3221 | 72.4643 | 54.7309 | 63.4494 | 41.8226 |
| 153 | 57.8502 | 67.6221 | 50.8248 | 67.5518 | 42.951 |
| 154 | 55.1951 | 63.4177 | 44.8134 | 58.7098 | 41.3716 |
| 155 | 59.1511 | 65.671 | 49.8542 | 58.2593 | 61.348 |
| 156 | 65.5055 | 72.2829 | 47.0984 | 57.2181 | 41.9216 |
| 157 | 62.4725 | 73.326 | 47.0347 | 61.8031 | 40.8129 |
| 158 | 67.063 | 70.6494 | 50.3547 | 59.6442 | 42.2429 |
| 159 | 57.1031 | 69.8329 | 47.9813 | 59.2714 | 43.4317 |
| 160 | 59.4419 | 72.7745 | 50.2757 | 58.4732 | 49.7061 |
| 161 | 56.6689 | 65.3546 | 49.0726 | 61.1651 | 46.7471 |
| 162 | 61.0166 | 65.7611 | 48.4689 | 60.1934 | 50.1233 |
| 163 | 58.0976 | 65.8974 | 52.4475 | 56.9825 | 47.6983 |
| 164 | 61.6099 | 64.9121 | 49.8618 | 62.3425 | 47.6734 |
| 165 | 62.9194 | 69.6546 | 48.6074 | 59.9781 | 46.7076 |
| 166 | 57.088 | 62.64 | 50.9674 | 59.9432 | 42.2996 |
| 167 | 53.4895 | 68.147 | 52.9571 | 64.0706 | 43.7262 |
| 168 | 58.3563 | 63.7812 | 48.2155 | 56.6298 | 42.7054 |
| 169 | 65.0209 | 61.0216 | 50.8447 | 53.5476 | 42.3227 |
| 170 | 58.2657 | 77.2499 | 46.2696 | 61.4058 | 38.7925 |
| 171 | 63.9293 | 63.1258 | 45.5634 | 57.3484 | 43.9471 |
| 172 | 59.2445 | 69.0889 | 43.4001 | 57.2568 | 54.4292 |
| 173 | 61.21 | 66.6289 | 46.2029 | 58.4846 | 48.0889 |
| 174 | 59.1076 | 67.6006 | 43.5191 | 59.8213 | 54.007 |
| 175 | 55.6669 | 60.5057 | 47.1389 | 62.7315 | 54.2612 |
| 176 | 54.3697 | 64.3054 | 50.9986 | 58.6401 | 44.9016 |
| 177 | 63.5288 | 65.285 | 47.8852 | 55.6173 | 44.7026 |
| 178 | 65.7156 | 65.6735 | 49.0271 | 63.692 | 46.4568 |
| 179 | 57.2823 | 61.0132 | 49.9075 | 60.6005 | 46.5529 |
| 180 | 70.5004 | 63.6052 | 49.2146 | 55.8531 | 46.7666 |
| 181 | 63.4699 | 66.5331 | 51.3323 | 57.125 | 53.718 |
| 182 | 56.4135 | 62.066 | 50.2745 | 63.4612 | 47.1843 |
| 183 | 51.6098 | 69.2883 | 46.7397 | 69.6855 | 48.8089 |
| 184 | 57.1345 | 71.4199 | 55.258 | 64.6531 | 57.3827 |
| 185 | 59.4109 | 71.5719 | 50.0133 | 62.3673 | 48.37 |
| 186 | 62.9138 | 71.9696 | 60.9243 | 59.1726 | 51.7089 |
| 187 | 57.7758 | 65.0051 | 57.1814 | 60.6009 | 38.6055 |
| 188 | 56.5806 | 68.8013 | 62.8038 | 63.723 | 40.388 |
| 189 | 56.6467 | 66.1468 | 55.1696 | 57.3908 | 40.2959 |
| 190 | 61.381 | 70.9982 | 53.3738 | 56.6919 | 43.9928 |
| 191 | 57.742 | 64.1491 | 47.7305 | 58.2069 | 48.5962 |
| 192 | 55.9535 | 59.9544 | 52.8668 | 61.2759 | 39.315 |
| 193 | 57.9048 | 71.1505 | 50.982 | 59.0258 | 44.6234 |
| 194 | 59.391 | 66.8032 | 55.3342 | 60.9361 | 41.5436 |
| 195 | 51.4382 | 61.459 | 47.7443 | 63.2578 | 45.6936 |
| 196 | 56.985 | 66.2023 | 52.3207 | 57.6319 | 43.035 |
| 197 | 56.8167 | 63.6983 | 38.6914 | 57.6718 | 43.7929 |
| 198 | 54.4879 | 65.2743 | 37.3421 | 58.932 | 42.2621 |
| 199 | 58.4783 | 56.5684 | 41.0174 | 61.1604 | 45.7122 |

ARG-10, ARG-11, ASP-21 (CLOCKS38/42)

Angle

| Frame | 1st | 2nd | 3rd | 4th | 5th |
| --- | --- | --- | --- | --- | --- |
| 0 | 19.3677 | 19.1635 | 111.254 | 3.79103 | 8.92769 |
| 1 | 19.3677 | 19.1635 | 111.254 | 3.79103 | 8.92769 |
| 2 | 65.2174 | 73.1323 | 64.3625 | 77.9131 | 78.6465 |
| 3 | 58.1688 | 76.5282 | 64.3053 | 71.7431 | 83.6776 |
| 4 | 59.992 | 69.7058 | 65.4775 | 75.209 | 75.0421 |
| 5 | 62.9497 | 77.2495 | 51.1842 | 74.1034 | 80.2996 |
| 6 | 55.0594 | 66.6191 | 64.3651 | 74.1368 | 67.5569 |
| 7 | 50.5823 | 65.6067 | 60.41 | 74.8209 | 63.2793 |
| 8 | 54.2973 | 70.0368 | 64.0118 | 76.1805 | 65.8758 |
| 9 | 55.9053 | 90.0652 | 62.3927 | 77.0094 | 63.5783 |
| 10 | 61.4607 | 71.9423 | 55.5536 | 78.636 | 67.749 |
| 11 | 55.2354 | 80.9513 | 53.0266 | 74.8153 | 64.9088 |
| 12 | 58.962 | 74.4953 | 56.1406 | 82.7101 | 66.5157 |
| 13 | 41.6247 | 74.0247 | 52.1451 | 83.2458 | 66.1305 |
| 14 | 42.9196 | 70.7173 | 45.5754 | 76.6976 | 66.1273 |
| 15 | 59.6379 | 73.0728 | 46.4353 | 76.769 | 67.3183 |
| 16 | 50.999 | 69.6157 | 62.9647 | 76.7406 | 66.8936 |
| 17 | 63.8802 | 65.1116 | 60.5805 | 77.7531 | 74.4745 |
| 18 | 51.1002 | 68.4461 | 60.0749 | 73.6035 | 74.0107 |
| 19 | 47.7446 | 61.5559 | 52.3736 | 77.1758 | 72.9922 |
| 20 | 47.6235 | 78.1094 | 58.3004 | 74.8597 | 72.4883 |
| 21 | 44.6253 | 76.9027 | 57.5026 | 75.0807 | 67.4972 |
| 22 | 47.9838 | 69.9747 | 58.886 | 76.0377 | 68.7986 |
| 23 | 48.8685 | 70.7606 | 55.4685 | 76.2288 | 69.0602 |
| 24 | 46.377 | 62.2154 | 59.1572 | 79.7653 | 67.867 |
| 25 | 42.8213 | 68.47 | 53.9533 | 81.1541 | 71.4092 |
| 26 | 45.7615 | 75.8817 | 47.5398 | 79.9472 | 71.701 |
| 27 | 44.3362 | 69.9236 | 51.9109 | 69.8648 | 72.3152 |
| 28 | 42.8013 | 65.2187 | 56.7822 | 71.9848 | 70.0339 |
| 29 | 42.2621 | 79.0604 | 60.9908 | 79.3143 | 72.2874 |
| 30 | 47.0987 | 78.714 | 53.6338 | 75.9905 | 74.5544 |
| 31 | 43.0494 | 89.3771 | 56.6623 | 73.0909 | 74.4103 |
| 32 | 46.7556 | 80.0192 | 60.805 | 70.4978 | 74.6049 |
| 33 | 46.535 | 73.413 | 55.2344 | 66.1044 | 81.0537 |
| 34 | 44.8976 | 73.3985 | 55.1693 | 67.5914 | 68.6077 |
| 35 | 48.4851 | 73.8333 | 55.2249 | 65.3595 | 83.978 |
| 36 | 50.9963 | 87.5819 | 60.0142 | 68.98 | 68.8136 |
| 37 | 46.5176 | 81.562 | 62.5068 | 63.127 | 69.6291 |
| 38 | 42.9547 | 87.9273 | 59.8889 | 66.31 | 67.4564 |
| 39 | 45.9871 | 88.0412 | 56.2837 | 71.8902 | 71.8085 |
| 40 | 42.3755 | 83.9072 | 60.6583 | 72.4528 | 77.2251 |
| 41 | 39.4443 | 83.5697 | 63.5574 | 77.1128 | 72.7219 |
| 42 | 51.4408 | 87.5104 | 61.525 | 73.6413 | 72.7046 |
| 43 | 48.2772 | 90.4934 | 60.9628 | 73.8204 | 63.6702 |
| 44 | 43.2492 | 84.6073 | 59.7581 | 74.8291 | 71.0411 |
| 45 | 47.7602 | 84.1455 | 66.7683 | 74.6981 | 67.3998 |
| 46 | 44.3726 | 86.9613 | 63.0645 | 72.4892 | 73.3259 |
| 47 | 43.7082 | 91.4257 | 71.8508 | 71.3698 | 69.7439 |
| 48 | 46.727 | 86.1025 | 60.1291 | 73.7235 | 77.2995 |
| 49 | 46.52 | 92.1932 | 61.0007 | 79.6892 | 74.9032 |
| 50 | 40.135 | 90.5429 | 60.6408 | 68.8463 | 71.1086 |
| 51 | 44.3936 | 78.4634 | 63.9022 | 71.2326 | 66.5961 |
| 52 | 41.4199 | 82.986 | 60.2038 | 74.4047 | 64.8646 |
| 53 | 47.3909 | 83.06 | 63.1583 | 66.8815 | 71.5317 |
| 54 | 50.8553 | 79.1982 | 58.5237 | 74.8152 | 69.8851 |
| 55 | 48.5188 | 72.7451 | 69.3946 | 75.8569 | 72.9136 |
| 56 | 50.757 | 89.4911 | 63.4215 | 67.06 | 74.862 |
| 57 | 47.9999 | 86.1876 | 59.5993 | 83.9726 | 73.5664 |
| 58 | 50.7233 | 83.4321 | 59.7958 | 70.5478 | 70.5399 |
| 59 | 47.9645 | 81.5516 | 58.6647 | 72.6588 | 65.1948 |
| 60 | 48.3261 | 78.9266 | 69.0318 | 76.0616 | 70.7343 |
| 61 | 47.7087 | 80.1105 | 58.8203 | 72.6678 | 65.4214 |
| 62 | 43.0288 | 84.3493 | 65.1916 | 77.1021 | 75.2402 |
| 63 | 49.5413 | 87.2485 | 59.1421 | 73.9028 | 75.6448 |
| 64 | 50.7717 | 84.5091 | 63.6154 | 70.1181 | 70.5408 |
| 65 | 49.4066 | 81.4409 | 63.9485 | 71.7631 | 70.8239 |
| 66 | 45.1204 | 79.3933 | 67.7131 | 67.973 | 71.5658 |
| 67 | 44.6405 | 82.6384 | 63.7564 | 72.8236 | 76.0225 |
| 68 | 54.1741 | 75.112 | 66.8029 | 69.8077 | 70.1154 |
| 69 | 49.3747 | 92.0214 | 66.697 | 74.3937 | 65.5793 |
| 70 | 43.7167 | 85.7807 | 61.0937 | 76.6387 | 62.3619 |
| 71 | 44.9381 | 79.179 | 60.889 | 71.3922 | 70.5594 |
| 72 | 54.3202 | 73.8558 | 63.6652 | 76.8532 | 73.2141 |
| 73 | 49.4593 | 73.3479 | 64.9454 | 76.785 | 70.8645 |
| 74 | 48.6005 | 81.8238 | 58.3047 | 71.6329 | 73.6142 |
| 75 | 48.0828 | 79.1316 | 55.1178 | 78.2327 | 74.3307 |
| 76 | 42.8942 | 72.2087 | 68.1069 | 69.0923 | 69.2916 |
| 77 | 46.6812 | 84.2821 | 60.3135 | 72.2441 | 66.9968 |
| 78 | 46.1411 | 90.782 | 52.0226 | 75.2996 | 72.657 |
| 79 | 52.1883 | 83.4573 | 58.4842 | 74.8721 | 69.4302 |
| 80 | 53.8227 | 88.7217 | 62.0363 | 74.8651 | 67.9805 |
| 81 | 40.9704 | 77.6136 | 62.1444 | 74.9898 | 70.3395 |
| 82 | 48.248 | 82.6242 | 65.1072 | 74.359 | 69.9577 |
| 83 | 49.4144 | 85.611 | 66.4083 | 76.042 | 71.8988 |
| 84 | 45.9699 | 94.6058 | 63.284 | 76.2735 | 68.4213 |
| 85 | 56.5649 | 90.0283 | 66.2025 | 76.4374 | 65.88 |
| 86 | 48.1808 | 91.0103 | 64.3666 | 77.9029 | 71.7846 |
| 87 | 52.3057 | 96.1643 | 62.6505 | 78.3027 | 80.2946 |
| 88 | 52.8332 | 89.7442 | 68.4971 | 75.9801 | 76.02 |
| 89 | 51.9017 | 87.9413 | 61.7471 | 76.5514 | 70.1049 |
| 90 | 49.6785 | 77.779 | 70.6075 | 73.9275 | 73.9613 |
| 91 | 49.5897 | 80.6346 | 59.4073 | 75.6685 | 68.2443 |
| 92 | 48.3749 | 86.3804 | 59.1571 | 75.5084 | 66.7139 |
| 93 | 51.1385 | 84.8887 | 51.0364 | 72.4646 | 76.0258 |
| 94 | 57.3077 | 90.1703 | 52.9465 | 73.5895 | 71.8164 |
| 95 | 58.2703 | 91.7366 | 49.5383 | 74.0129 | 73.4827 |
| 96 | 53.1696 | 80.2395 | 50.1808 | 72.6619 | 61.4846 |
| 97 | 56.2834 | 80.0558 | 55.3837 | 77.5363 | 80.8763 |
| 98 | 57.1878 | 83.7044 | 53.0298 | 83.1948 | 68.0185 |
| 99 | 48.247 | 82.422 | 50.9064 | 80.4971 | 69.6444 |
| 100 | 50.3536 | 79.9933 | 56.7995 | 78.0435 | 67.9939 |
| 101 | 53.3491 | 89.2578 | 52.2868 | 76.9155 | 73.4037 |
| 102 | 53.1028 | 76.6096 | 50.4497 | 75.9307 | 69.2403 |
| 103 | 48.8299 | 83.3381 | 59.1952 | 71.7002 | 68.6538 |
| 104 | 46.9837 | 77.0997 | 55.0519 | 75.1454 | 70.9131 |
| 105 | 48.7986 | 79.9355 | 58.4358 | 75.2099 | 69.2359 |
| 106 | 59.4945 | 88.3828 | 64.0555 | 71.5039 | 71.7449 |
| 107 | 72.9009 | 79.1782 | 58.1112 | 79.7063 | 81.0994 |
| 108 | 72.7194 | 85.5826 | 58.1562 | 72.3256 | 79.5104 |
| 109 | 69.139 | 87.0065 | 59.7891 | 73.5981 | 70.7714 |
| 110 | 69.6967 | 81.5475 | 61.6911 | 71.3417 | 72.3409 |
| 111 | 63.3548 | 88.959 | 62.8254 | 72.4638 | 84.4248 |
| 112 | 65.2748 | 94.6671 | 57.0848 | 74.1755 | 72.7533 |
| 113 | 63.6995 | 89.5636 | 56.5292 | 79.4112 | 73.918 |
| 114 | 70.4756 | 83.4613 | 61.4737 | 73.9231 | 75.3265 |
| 115 | 66.9439 | 87.0654 | 59.6765 | 67.1511 | 73.5618 |
| 116 | 63.8802 | 78.0132 | 59.6187 | 72.5587 | 72.2503 |
| 117 | 63.0676 | 81.8173 | 55.4942 | 70.0445 | 73.9083 |
| 118 | 64.8532 | 77.738 | 58.3138 | 70.5373 | 74.2268 |
| 119 | 64.2851 | 79.5049 | 60.0433 | 73.736 | 75.8623 |
| 120 | 62.9318 | 79.828 | 56.8097 | 71.547 | 72.3665 |
| 121 | 63.3111 | 79.3033 | 56.9317 | 67.2822 | 75.3999 |
| 122 | 67.3414 | 79.605 | 55.732 | 72.453 | 80.2837 |
| 123 | 66.064 | 82.8338 | 58.8056 | 75.4584 | 72.6823 |
| 124 | 65.9849 | 79.4829 | 60.6289 | 72.793 | 65.9151 |
| 125 | 66.7337 | 80.022 | 56.3879 | 77.3342 | 71.763 |
| 126 | 66.0106 | 81.8398 | 60.447 | 74.8406 | 78.9694 |
| 127 | 71.3882 | 77.3067 | 59.8031 | 75.2031 | 76.9036 |
| 128 | 64.353 | 80.6232 | 56.578 | 77.125 | 79.7485 |
| 129 | 69.5541 | 75.7944 | 57.8156 | 73.7419 | 74.2473 |
| 130 | 72.4856 | 78.9406 | 61.513 | 73.3705 | 72.3635 |
| 131 | 65.2571 | 76.2014 | 60.7789 | 74.3895 | 73.5326 |
| 132 | 77.4171 | 79.9187 | 60.1778 | 77.2327 | 77.3242 |
| 133 | 65.6446 | 78.4768 | 60.5641 | 70.6555 | 73.8906 |
| 134 | 68.8164 | 80.7223 | 59.862 | 73.5184 | 76.1103 |
| 135 | 59.9328 | 73.1314 | 57.2903 | 71.6602 | 72.5269 |
| 136 | 70.2756 | 77.2463 | 60.3481 | 77.0787 | 71.9642 |
| 137 | 66.0571 | 83.9338 | 58.2906 | 75.0889 | 70.9846 |
| 138 | 75.8092 | 79.5974 | 57.142 | 70.5026 | 74.3651 |
| 139 | 57.1728 | 81.7707 | 58.2219 | 70.8619 | 76.7647 |
| 140 | 67.27 | 82.8019 | 58.6342 | 71.3762 | 78.1445 |
| 141 | 72.1081 | 85.0305 | 56.2246 | 78.3524 | 71.2739 |
| 142 | 64.4986 | 81.315 | 54.4671 | 74.1791 | 77.0858 |
| 143 | 63.5315 | 78.2258 | 58.4283 | 69.7606 | 76.4212 |
| 144 | 70.8747 | 79.5776 | 50.6776 | 73.2002 | 73.076 |
| 145 | 54.851 | 78.6851 | 52.9861 | 73.8637 | 77.6981 |
| 146 | 56.5169 | 80.433 | 60.5623 | 74.2034 | 77.1359 |
| 147 | 67.0192 | 75.1814 | 56.9562 | 73.6012 | 74.5407 |
| 148 | 66.3052 | 78.0499 | 58.7681 | 69.6863 | 71.2721 |
| 149 | 63.0982 | 75.8666 | 52.96 | 69.6412 | 75.548 |
| 150 | 65.6394 | 79.5649 | 59.3091 | 72.7192 | 77.9227 |
| 151 | 65.5476 | 77.1719 | 60.7635 | 73.5497 | 74.4559 |
| 152 | 55.5491 | 76.5853 | 59.288 | 66.6682 | 69.6144 |
| 153 | 68.9566 | 77.4251 | 60.7894 | 76.0695 | 73.3343 |
| 154 | 68.3134 | 75.5693 | 53.5065 | 76.1958 | 71.4861 |
| 155 | 61.4593 | 80.8519 | 56.0817 | 73.4613 | 73.5599 |
| 156 | 64.0855 | 80.4405 | 58.2549 | 71.9616 | 76.6708 |
| 157 | 63.7525 | 76.4375 | 55.6078 | 75.4563 | 70.3432 |
| 158 | 68.8931 | 77.8064 | 61.3566 | 78.4047 | 68.7093 |
| 159 | 66.8824 | 75.7793 | 60.7098 | 76.8572 | 77.1969 |
| 160 | 64.9761 | 72.973 | 52.1846 | 68.7298 | 68.1591 |
| 161 | 66.5958 | 81.2261 | 52.5152 | 73.0193 | 69.9928 |
| 162 | 67.1414 | 77.8711 | 57.7214 | 74.0788 | 72.7325 |
| 163 | 62.8615 | 74.5196 | 51.9237 | 83.5119 | 72.4449 |
| 164 | 59.3038 | 89.3629 | 59.6443 | 73.9465 | 65.0976 |
| 165 | 66.0012 | 81.8712 | 61.8426 | 70.8793 | 77.4631 |
| 166 | 55.4923 | 77.4831 | 59.5101 | 75.7551 | 80.6338 |
| 167 | 62.1818 | 82.5478 | 55.1819 | 73.8092 | 77.7802 |
| 168 | 62.8667 | 79.3303 | 46.86 | 84.959 | 73.7091 |
| 169 | 56.5231 | 79.5735 | 57.0558 | 78.8727 | 66.364 |
| 170 | 61.5529 | 79.8843 | 57.2181 | 70.7701 | 62.8983 |
| 171 | 59.76 | 84.0767 | 54.2948 | 75.5462 | 66.5625 |
| 172 | 61.1798 | 83.7665 | 55.5125 | 78.3269 | 69.8934 |
| 173 | 62.0695 | 85.7622 | 49.7185 | 71.5802 | 71.3219 |
| 174 | 63.5045 | 86.9519 | 59.2523 | 80.9433 | 65.8506 |
| 175 | 70.04 | 88.586 | 59.4107 | 76.3335 | 74.6484 |
| 176 | 67.6113 | 78.9186 | 56.1969 | 73.1696 | 66.5103 |
| 177 | 60.8776 | 73.6132 | 51.128 | 71.8073 | 75.5587 |
| 178 | 63.0198 | 91.3842 | 53.0184 | 73.8752 | 74.4789 |
| 179 | 73.812 | 80.4765 | 54.3705 | 78.3872 | 79.5666 |
| 180 | 70.1321 | 76.6356 | 58.211 | 75.8081 | 72.3524 |
| 181 | 60.7646 | 87.2682 | 52.2165 | 71.6113 | 77.0743 |
| 182 | 59.5039 | 70.4871 | 50.1862 | 79.1797 | 69.1815 |
| 183 | 64.39 | 78.9679 | 52.2686 | 70.9979 | 73.0281 |
| 184 | 61.2648 | 87.2087 | 55.7202 | 76.288 | 78.219 |
| 185 | 66.8504 | 83.8785 | 53.6789 | 75.9035 | 81.7014 |
| 186 | 71.7691 | 80.2908 | 58.0989 | 76.4768 | 84.0274 |
| 187 | 68.9151 | 82.4695 | 56.6656 | 78.6151 | 73.489 |
| 188 | 70.1439 | 72.0856 | 58.9551 | 74.3913 | 66.3374 |
| 189 | 65.0149 | 77.6504 | 55.4807 | 70.9941 | 74.9624 |
| 190 | 66.3038 | 75.6104 | 57.2956 | 71.1477 | 70.7555 |
| 191 | 71.1978 | 80.4279 | 51.8971 | 70.8089 | 66.9817 |
| 192 | 66.2447 | 85.3294 | 55.6358 | 71.5496 | 72.8588 |
| 193 | 64.2776 | 87.0358 | 46.9854 | 74.5821 | 65.5714 |
| 194 | 67.4372 | 72.2587 | 58.5032 | 77.1684 | 68.0499 |
| 195 | 61.961 | 76.099 | 51.7656 | 73.9961 | 67.6612 |
| 196 | 63.1852 | 95.7995 | 57.6631 | 75.2012 | 65.3862 |
| 197 | 62.6272 | 86.6793 | 52.5976 | 74.7628 | 65.5081 |
| 198 | 64.2994 | 81.1413 | 50.9643 | 77.5782 | 67.1573 |
| 199 | 61.0296 | 79.12 | 49.6518 | 75.2246 | 66.0274 |

ARG-10, ARG-11, ASP-21 (CLOCKS38/42/BMAL1S78)

Angle

| Frame | 1st | 2nd | 3rd | 4th | 5th |
| --- | --- | --- | --- | --- | --- |
| 0 | 72.654 | 72.4159 | 69.1899 | 7.85121 | 10.7231 |
| 1 | 72.654 | 72.4159 | 69.19 | 7.85121 | 10.7231 |
| 2 | 72.6519 | 72.2722 | 69.0215 | 89.7717 | 70.7993 |
| 3 | 63.0201 | 53.9718 | 77.4531 | 97.2504 | 59.8807 |
| 4 | 78.4718 | 50.0195 | 74.7016 | 82.7333 | 71.8191 |
| 5 | 72.8765 | 47.2472 | 79.4086 | 81.5548 | 69.7547 |
| 6 | 72.2274 | 53.4598 | 75.7232 | 81.7126 | 71.4733 |
| 7 | 75.8209 | 46.1482 | 66.0821 | 81.024 | 74.533 |
| 8 | 76.3942 | 53.3302 | 61.5777 | 83.0304 | 76.4046 |
| 9 | 63.6049 | 57.5663 | 71.5914 | 79.4065 | 69.9757 |
| 10 | 60.7 | 60.0253 | 70.1419 | 83.7317 | 65.3114 |
| 11 | 60.7575 | 69.0387 | 77.0869 | 85.1535 | 61.8083 |
| 12 | 46.4881 | 59.3169 | 74.7571 | 83.4366 | 65.5831 |
| 13 | 56.8054 | 70.4407 | 69.6023 | 90.0138 | 50.7249 |
| 14 | 51.5156 | 74.1639 | 58.2415 | 92.006 | 59.2123 |
| 15 | 50.8703 | 72.255 | 63.6701 | 94.013 | 61.9484 |
| 16 | 55.0941 | 67.1222 | 66.1243 | 83.5472 | 57.1293 |
| 17 | 54.1265 | 59.7933 | 61.8057 | 86.1181 | 53.0025 |
| 18 | 53.3206 | 59.6989 | 57.519 | 87.971 | 58.1056 |
| 19 | 53.7568 | 61.1277 | 57.3002 | 82.6517 | 49.3532 |
| 20 | 60.8485 | 63.716 | 59.1564 | 85.875 | 46.0591 |
| 21 | 55.0465 | 61.4782 | 54.0163 | 89.5308 | 50.0024 |
| 22 | 47.6707 | 51.9674 | 56.8286 | 81.0858 | 60.6609 |
| 23 | 49.9715 | 61.632 | 55.9247 | 86.3953 | 66.3027 |
| 24 | 72.1891 | 59.7096 | 62.159 | 84.8889 | 51.3972 |
| 25 | 72.035 | 69.3326 | 59.6413 | 88.0679 | 60.2439 |
| 26 | 67.8414 | 63.3476 | 55.4666 | 91.7739 | 59.3352 |
| 27 | 80.1297 | 69.4493 | 58.436 | 80.8889 | 56.9326 |
| 28 | 69.7747 | 62.2441 | 62.2575 | 87.9289 | 56.2824 |
| 29 | 80.4334 | 68.8868 | 60.1445 | 85.7324 | 59.7386 |
| 30 | 74.6775 | 51.5092 | 53.1556 | 79.2064 | 52.7262 |
| 31 | 67.2196 | 48.7566 | 66.9405 | 88.5799 | 56.4169 |
| 32 | 67.6983 | 39.6904 | 61.7571 | 78.3851 | 58.2887 |
| 33 | 69.9112 | 33.6527 | 67.7318 | 84.9245 | 57.1823 |
| 34 | 69.5785 | 48.368 | 62.7958 | 77.8775 | 62.6118 |
| 35 | 71.6153 | 52.3276 | 63.6512 | 89.2391 | 56.6163 |
| 36 | 66.3611 | 57.5297 | 63.473 | 82.3541 | 60.5064 |
| 37 | 67.4991 | 52.6066 | 60.336 | 74.1355 | 55.0008 |
| 38 | 62.4059 | 50.1838 | 65.4585 | 90.7056 | 57.2731 |
| 39 | 64.4091 | 51.1053 | 63.7202 | 90.9375 | 59.9665 |
| 40 | 74.0991 | 56.9111 | 54.6829 | 85.7165 | 53.4291 |
| 41 | 74.8139 | 55.919 | 58.7761 | 84.0467 | 58.8763 |
| 42 | 75.5449 | 51.8193 | 61.5334 | 83.8802 | 62.645 |
| 43 | 72.5062 | 57.5624 | 61.01 | 82.9086 | 53.751 |
| 44 | 73.2732 | 53.4967 | 58.0327 | 82.2164 | 56.4444 |
| 45 | 75.0638 | 57.7544 | 59.599 | 87.9515 | 62.7964 |
| 46 | 71.43 | 49.2041 | 59.8886 | 87.1648 | 58.3494 |
| 47 | 78.6663 | 62.6213 | 63.3098 | 91.3045 | 48.0341 |
| 48 | 70.1913 | 52.4805 | 53.17 | 84.6083 | 53.8367 |
| 49 | 75.4921 | 65.1236 | 59.8381 | 90.3043 | 58.1949 |
| 50 | 68.0669 | 46.0305 | 56.7755 | 83.3589 | 52.3798 |
| 51 | 71.3284 | 61.2733 | 57.0923 | 87.6105 | 57.6348 |
| 52 | 76.0507 | 60.1075 | 56.4251 | 90.903 | 55.3261 |
| 53 | 74.6909 | 66.0293 | 54.2895 | 88.535 | 55.5708 |
| 54 | 73.6374 | 54.329 | 54.5503 | 101.883 | 55.2028 |
| 55 | 75.5279 | 50.7982 | 52.0396 | 82.0568 | 56.8602 |
| 56 | 71.9805 | 50.1302 | 56.2571 | 83.0273 | 48.2742 |
| 57 | 79.9879 | 54.0003 | 56.1664 | 87.2299 | 55.8298 |
| 58 | 74.3223 | 55.6716 | 58.7244 | 86.7188 | 62.1031 |
| 59 | 71.9012 | 58.3739 | 62.5716 | 90.0741 | 54.4888 |
| 60 | 77.773 | 48.3289 | 59.8421 | 86.7997 | 55.7042 |
| 61 | 72.2284 | 53.8151 | 62.0724 | 89.4726 | 56.3798 |
| 62 | 71.1329 | 49.8471 | 62.6292 | 86.4574 | 57.9902 |
| 63 | 68.9587 | 56.8593 | 59.5892 | 85.5319 | 58.1551 |
| 64 | 64.2394 | 59.5956 | 59.8859 | 84.8238 | 59.6009 |
| 65 | 76.4847 | 53.8336 | 63.6105 | 92.1115 | 55.8716 |
| 66 | 73.2727 | 60.1911 | 53.9952 | 83.083 | 55.2755 |
| 67 | 76.466 | 58.0091 | 64.4377 | 87.1565 | 55.7615 |
| 68 | 73.5218 | 55.7736 | 50.8287 | 91.7869 | 56.7416 |
| 69 | 79.6834 | 64.7641 | 59.6921 | 93.1167 | 56.3983 |
| 70 | 77.5679 | 53.7813 | 61.1719 | 83.6916 | 60.7203 |
| 71 | 71.7049 | 53.35 | 56.8007 | 85.5856 | 59.2548 |
| 72 | 69.6944 | 55.2892 | 64.6075 | 79.3215 | 57.7525 |
| 73 | 72.5539 | 58.8142 | 62.6856 | 87.4142 | 48.7817 |
| 74 | 71.3492 | 52.7514 | 59.9208 | 88.3805 | 54.8382 |
| 75 | 66.3879 | 65.5372 | 60.929 | 92.8264 | 62.2406 |
| 76 | 72.2598 | 60.5136 | 60.7403 | 86.1306 | 52.8176 |
| 77 | 71.2353 | 58.3843 | 59.2509 | 85.3116 | 59.1203 |
| 78 | 74.3113 | 59.6637 | 66.1021 | 85.3304 | 46.6864 |
| 79 | 72.7408 | 57.1356 | 59.8567 | 88.9485 | 55.8535 |
| 80 | 74.0188 | 59.7122 | 73.3825 | 88.0316 | 52.4721 |
| 81 | 72.0258 | 57.5711 | 59.7588 | 91.3401 | 55.5737 |
| 82 | 70.9966 | 60.462 | 57.5916 | 87.1066 | 59.7169 |
| 83 | 70.5204 | 63.4999 | 60.5038 | 82.6996 | 55.2814 |
| 84 | 70.0659 | 59.5786 | 64.2381 | 89.7545 | 49.7362 |
| 85 | 65.5278 | 50.1874 | 65.1965 | 77.9189 | 53.3105 |
| 86 | 69.679 | 56.0133 | 64.153 | 90.0869 | 49.3849 |
| 87 | 66.9096 | 57.1162 | 61.6073 | 86.2322 | 49.4162 |
| 88 | 70.2028 | 59.7515 | 60.0728 | 87.8869 | 52.1108 |
| 89 | 67.1028 | 56.422 | 61.6358 | 81.0169 | 54.9454 |
| 90 | 71.0405 | 60.3275 | 57.1276 | 88.757 | 56.1287 |
| 91 | 73.1219 | 61.0858 | 60.4667 | 85.3809 | 60.5438 |
| 92 | 78.4175 | 58.5805 | 60.4255 | 85.7169 | 56.9035 |
| 93 | 78.8932 | 56.6836 | 62.7735 | 86.5402 | 60.8229 |
| 94 | 83.1387 | 52.6949 | 69.872 | 89.6693 | 53.9911 |
| 95 | 78.5347 | 56.6059 | 60.3774 | 85.5235 | 50.8753 |
| 96 | 80.3149 | 56.6813 | 63.2854 | 82.4732 | 57.029 |
| 97 | 78.472 | 60.7274 | 69.8902 | 78.5902 | 59.118 |
| 98 | 80.8901 | 58.53 | 66.8692 | 80.5874 | 62.2195 |
| 99 | 74.1404 | 49.5873 | 63.5384 | 83.6209 | 60.2035 |
| 100 | 81.8235 | 65.5135 | 66.3498 | 88.4921 | 58.6011 |
| 101 | 78.2977 | 71.1571 | 61.7043 | 84.6065 | 55.5823 |
| 102 | 79.673 | 63.1571 | 62.3339 | 80.4296 | 57.4463 |
| 103 | 86.2397 | 57.9211 | 65.1297 | 81.9962 | 58.9191 |
| 104 | 80.5753 | 62.8638 | 68.5229 | 72.3268 | 58.0579 |
| 105 | 84.2779 | 58.6421 | 66.3867 | 86.3591 | 61.1593 |
| 106 | 80.5692 | 62.3002 | 68.9418 | 83.1632 | 56.1092 |
| 107 | 77.0898 | 52.2818 | 61.1623 | 83.2599 | 51.9424 |
| 108 | 73.1478 | 56.0048 | 70.2575 | 89.5022 | 58.5627 |
| 109 | 78.9309 | 53.6603 | 62.2234 | 82.705 | 65.0564 |
| 110 | 80.5359 | 53.5278 | 67.8372 | 84.4618 | 53.7159 |
| 111 | 85.532 | 56.7589 | 66.1988 | 84.2421 | 54.5953 |
| 112 | 83.7903 | 49.8351 | 60.766 | 86.8939 | 58.8065 |
| 113 | 74.1333 | 56.2059 | 56.9663 | 83.4984 | 57.7813 |
| 114 | 74.547 | 50.4975 | 61.6422 | 77.7673 | 55.1944 |
| 115 | 80.2399 | 46.4945 | 60.5907 | 78.7631 | 63.4454 |
| 116 | 81.624 | 61.7623 | 55.8429 | 81.6437 | 58.461 |
| 117 | 83.341 | 57.3688 | 56.6433 | 87.9644 | 58.5135 |
| 118 | 84.5921 | 54.8018 | 59.4527 | 78.33 | 54.4642 |
| 119 | 79.1432 | 61.9506 | 68.5279 | 81.6432 | 57.7524 |
| 120 | 79.36 | 62.273 | 61.0854 | 84.2949 | 65.5666 |
| 121 | 72.7192 | 59.8238 | 60.3087 | 82.82 | 62.7687 |
| 122 | 72.1642 | 64.7161 | 70.7006 | 75.9911 | 58.8678 |
| 123 | 69.6118 | 50.4937 | 64.2077 | 80.5029 | 60.8981 |
| 124 | 65.0821 | 47.0038 | 68.2007 | 81.1431 | 58.5451 |
| 125 | 67.0657 | 67.9546 | 52.4358 | 82.1277 | 55.2743 |
| 126 | 68.6757 | 53.057 | 56.0164 | 85.9168 | 58.0592 |
| 127 | 68.5672 | 61.6982 | 61.0169 | 81.1974 | 59.6241 |
| 128 | 71.3505 | 59.9087 | 55.3834 | 88.0192 | 58.1483 |
| 129 | 72.1806 | 49.7344 | 63.1674 | 83.5831 | 49.5869 |
| 130 | 73.4086 | 56.3238 | 56.1394 | 77.4535 | 54.9711 |
| 131 | 63.7165 | 56.861 | 59.1558 | 80.0481 | 54.2479 |
| 132 | 70.4979 | 54.6007 | 64.2703 | 80.5959 | 53.7908 |
| 133 | 65.7504 | 51.9085 | 52.4965 | 87.5927 | 55.6747 |
| 134 | 70.222 | 60.6489 | 56.8921 | 86.2844 | 54.4741 |
| 135 | 61.927 | 63.7097 | 54.6068 | 89.2302 | 54.2078 |
| 136 | 68.0503 | 57.0097 | 45.7957 | 93.0514 | 57.4427 |
| 137 | 69.4309 | 56.6927 | 58.5673 | 87.2902 | 62.1475 |
| 138 | 68.7409 | 59.3382 | 59.8724 | 90.7427 | 65.5877 |
| 139 | 62.2173 | 57.7407 | 59.0383 | 79.961 | 60.684 |
| 140 | 66.1091 | 53.1769 | 54.5464 | 89.9135 | 60.5475 |
| 141 | 70.1518 | 70.1175 | 64.8037 | 95.1658 | 60.9186 |
| 142 | 67.4762 | 63.6164 | 52.8209 | 83.7287 | 57.9321 |
| 143 | 71.1381 | 71.5967 | 56.9826 | 88.372 | 58.5887 |
| 144 | 65.3287 | 67.7576 | 52.3516 | 83.189 | 55.4369 |
| 145 | 70.2281 | 55.8274 | 50.8307 | 91.0769 | 55.7332 |
| 146 | 66.2385 | 59.9309 | 57.4069 | 89.3587 | 57.2624 |
| 147 | 69.2155 | 59.595 | 58.1892 | 86.3663 | 57.5602 |
| 148 | 70.2116 | 59.0493 | 62.6045 | 81.9792 | 55.7649 |
| 149 | 73.1888 | 55.8943 | 56.64 | 87.5579 | 52.8432 |
| 150 | 70.6403 | 60.0543 | 56.636 | 77.4812 | 58.9528 |
| 151 | 69.3678 | 65.6865 | 61.3005 | 76.7743 | 55.8726 |
| 152 | 75.7303 | 58.7004 | 60.2758 | 84.5987 | 66.2839 |
| 153 | 71.487 | 57.3831 | 60.3103 | 83.4538 | 67.9435 |
| 154 | 70.7019 | 62.1561 | 56.4775 | 84.553 | 63.1557 |
| 155 | 72.2964 | 57.4131 | 56.3892 | 89.163 | 63.5952 |
| 156 | 76.2312 | 62.4833 | 53.9247 | 82.2582 | 62.0157 |
| 157 | 67.8099 | 63.506 | 55.1287 | 92.0097 | 58.9172 |
| 158 | 69.0499 | 65.7002 | 56.9154 | 90.2099 | 65.2896 |
| 159 | 69.16 | 64.9322 | 46.4809 | 82.1559 | 69.1513 |
| 160 | 70.8113 | 58.4515 | 56.6757 | 88.6555 | 60.7499 |
| 161 | 68.6842 | 64.5475 | 61.4255 | 89.3288 | 65.6696 |
| 162 | 67.3167 | 58.3967 | 63.3022 | 89.9087 | 63.8264 |
| 163 | 71.7852 | 58.3861 | 69.0054 | 84.8429 | 65.0111 |
| 164 | 69.3869 | 58.1302 | 63.3072 | 81.6333 | 60.5369 |
| 165 | 70.6958 | 56.3527 | 66.4706 | 85.341 | 62.4916 |
| 166 | 71.6422 | 64.6104 | 51.2369 | 83.4457 | 67.4001 |
| 167 | 72.773 | 52.9286 | 59.5189 | 91.8793 | 65.5056 |
| 168 | 72.9757 | 52.5829 | 59.3977 | 78.1205 | 70.8409 |
| 169 | 71.752 | 58.1777 | 58.8842 | 80.1669 | 52.8676 |
| 170 | 68.6672 | 47.6433 | 56.4803 | 83.6316 | 61.0644 |
| 171 | 71.2341 | 52.9432 | 53.2788 | 84.6633 | 57.4278 |
| 172 | 70.6806 | 56.4865 | 55.7065 | 89.6046 | 66.4935 |
| 173 | 68.14 | 57.0477 | 52.7734 | 79.8201 | 65.1235 |
| 174 | 70.5905 | 55.5886 | 58.2504 | 81.5331 | 61.2607 |
| 175 | 66.3816 | 55.6611 | 58.687 | 85.8014 | 66.5307 |
| 176 | 68.1622 | 57.5255 | 62.1855 | 80.9254 | 59.9061 |
| 177 | 83.3885 | 51.8108 | 61.4667 | 81.6407 | 61.2998 |
| 178 | 79.1122 | 48.2732 | 62.8485 | 83.8887 | 65.6443 |
| 179 | 71.9176 | 49.2138 | 57.2571 | 85.8598 | 65.5242 |
| 180 | 76.5506 | 54.1984 | 63.7188 | 87.3735 | 62.8838 |
| 181 | 76.1114 | 57.548 | 59.2527 | 84.2278 | 61.7769 |
| 182 | 74.8988 | 59.1477 | 54.1765 | 75.466 | 59.8126 |
| 183 | 73.8266 | 54.3692 | 55.14 | 81.3614 | 68.4171 |
| 184 | 75.8472 | 50.2977 | 59.6192 | 90.1328 | 64.7957 |
| 185 | 73.3383 | 55.764 | 51.0129 | 85.2015 | 58.5206 |
| 186 | 72.7351 | 51.687 | 66.1618 | 89.8319 | 63.6326 |
| 187 | 72.3117 | 50.6703 | 65.5786 | 83.3713 | 67.0341 |
| 188 | 76.7653 | 58.9286 | 55.0063 | 86.4136 | 63.4163 |
| 189 | 68.7203 | 55.4667 | 59.8255 | 78.1102 | 60.1059 |
| 190 | 67.2873 | 51.3864 | 56.9984 | 93.6696 | 61.4538 |
| 191 | 71.3284 | 51.9409 | 60.9463 | 85.1889 | 61.9175 |
| 192 | 75.0802 | 55.8353 | 54.5879 | 85.2575 | 64.6577 |
| 193 | 69.714 | 49.6261 | 56.4069 | 86.2967 | 59.4276 |
| 194 | 79.5031 | 49.5791 | 56.562 | 91.4672 | 62.6366 |
| 195 | 76.8414 | 56.4545 | 56.5345 | 84.545 | 60.3799 |
| 196 | 76.6426 | 45.512 | 66.4385 | 86.59 | 60.808 |
| 197 | 74.1774 | 56.2718 | 55.3557 | 80.6704 | 67.0355 |
| 198 | 67.9219 | 54.8549 | 60.5896 | 80.5986 | 72.1574 |
| 199 | 73.1521 | 47.7025 | 64.2941 | 76.8814 | 77.3026 |

**Figure S4.** The angle described to quantify the relative orientation of the fork relative to the E-box.

Histones - CLOCK_bHLH (nonphosphorylated)

Number of contacts

| Frame | 1st | 2nd | 3rd | 4th | 5th |
| --- | --- | --- | --- | --- | --- |
| 0 | 8 | 1 | 13 | 18 | 5 |
| 0.1 | 8 | 1 | 13 | 18 | 0 |
| 0.2 | 8 | 1 | 11 | 27 | 0 |
| 0.3 | 8 | 1 | 19 | 17 | 0 |
| 0.4 | 33 | 0 | 21 | 25 | 0 |
| 0.5 | 39 | 0 | 25 | 28 | 0 |
| 0.6 | 37 | 14 | 36 | 28 | 0 |
| 0.7 | 48 | 16 | 35 | 19 | 1 |
| 0.8 | 37 | 15 | 33 | 21 | 1 |
| 0.9 | 38 | 13 | 35 | 27 | 3 |
| 1 | 41 | 10 | 33 | 26 | 14 |
| 1.1 | 43 | 16 | 32 | 20 | 3 |
| 1.2 | 39 | 8 | 31 | 20 | 4 |
| 1.3 | 41 | 8 | 35 | 14 | 3 |
| 1.4 | 37 | 10 | 37 | 18 | 4 |
| 1.5 | 27 | 18 | 32 | 21 | 17 |
| 1.6 | 28 | 12 | 40 | 14 | 12 |
| 1.7 | 19 | 6 | 32 | 14 | 15 |
| 1.8 | 24 | 11 | 53 | 15 | 5 |
| 1.9 | 25 | 17 | 44 | 10 | 10 |
| 2 | 24 | 7 | 39 | 10 | 6 |
| 2.1 | 26 | 8 | 31 | 13 | 8 |
| 2.2 | 25 | 17 | 30 | 9 | 22 |
| 2.3 | 38 | 14 | 31 | 20 | 7 |
| 2.4 | 24 | 12 | 32 | 16 | 8 |
| 2.5 | 28 | 43 | 40 | 14 | 8 |
| 2.6 | 25 | 34 | 32 | 23 | 5 |
| 2.7 | 25 | 32 | 33 | 15 | 10 |
| 2.8 | 39 | 27 | 34 | 16 | 10 |
| 2.9 | 21 | 32 | 31 | 19 | 14 |
| 3 | 18 | 33 | 43 | 10 | 12 |
| 3.1 | 19 | 29 | 17 | 15 | 5 |
| 3.2 | 27 | 30 | 29 | 13 | 10 |
| 3.3 | 29 | 25 | 29 | 12 | 18 |
| 3.4 | 31 | 33 | 31 | 13 | 20 |
| 3.5 | 22 | 29 | 32 | 19 | 14 |
| 3.6 | 43 | 25 | 36 | 16 | 15 |
| 3.7 | 32 | 24 | 30 | 16 | 14 |
| 3.8 | 37 | 31 | 26 | 16 | 14 |
| 3.9 | 33 | 35 | 37 | 19 | 14 |
| 4 | 37 | 27 | 28 | 12 | 9 |
| 4.1 | 33 | 36 | 25 | 17 | 13 |
| 4.2 | 31 | 41 | 27 | 15 | 16 |
| 4.3 | 38 | 36 | 34 | 17 | 7 |
| 4.4 | 30 | 36 | 36 | 21 | 2 |
| 4.5 | 35 | 36 | 42 | 20 | 0 |
| 4.6 | 40 | 32 | 34 | 13 | 1 |
| 4.7 | 35 | 35 | 38 | 14 | 0 |
| 4.8 | 31 | 34 | 39 | 17 | 1 |
| 4.9 | 34 | 33 | 39 | 13 | 0 |
| 5 | 38 | 27 | 39 | 13 | 0 |
| 5.1 | 30 | 38 | 32 | 17 | 0 |
| 5.2 | 36 | 23 | 41 | 13 | 0 |
| 5.3 | 39 | 31 | 39 | 16 | 0 |
| 5.4 | 35 | 37 | 32 | 12 | 2 |
| 5.5 | 37 | 30 | 45 | 31 | 3 |
| 5.6 | 36 | 23 | 46 | 31 | 0 |
| 5.7 | 41 | 28 | 41 | 25 | 0 |
| 5.8 | 39 | 20 | 39 | 17 | 0 |
| 5.9 | 38 | 23 | 48 | 14 | 1 |
| 6 | 31 | 26 | 41 | 26 | 1 |
| 6.1 | 28 | 21 | 36 | 9 | 0 |
| 6.2 | 40 | 13 | 26 | 8 | 0 |
| 6.3 | 37 | 12 | 32 | 11 | 0 |
| 6.4 | 36 | 17 | 34 | 16 | 1 |
| 6.5 | 40 | 9 | 26 | 17 | 1 |
| 6.6 | 46 | 8 | 32 | 15 | 0 |
| 6.7 | 50 | 13 | 29 | 14 | 0 |
| 6.8 | 36 | 13 | 32 | 17 | 1 |
| 6.9 | 45 | 16 | 31 | 25 | 5 |
| 7 | 43 | 3 | 27 | 15 | 1 |
| 7.1 | 37 | 4 | 37 | 17 | 1 |
| 7.2 | 42 | 7 | 38 | 5 | 9 |
| 7.3 | 27 | 5 | 36 | 9 | 3 |
| 7.4 | 35 | 5 | 37 | 11 | 8 |
| 7.5 | 39 | 6 | 36 | 11 | 1 |
| 7.6 | 24 | 5 | 41 | 13 | 0 |
| 7.7 | 35 | 10 | 33 | 7 | 2 |
| 7.8 | 28 | 14 | 37 | 8 | 1 |
| 7.9 | 32 | 22 | 39 | 1 | 2 |
| 8 | 31 | 21 | 31 | 13 | 0 |
| 8.1 | 28 | 38 | 44 | 15 | 0 |
| 8.2 | 41 | 32 | 42 | 4 | 0 |
| 8.3 | 27 | 27 | 32 | 7 | 1 |
| 8.4 | 34 | 32 | 37 | 6 | 0 |
| 8.5 | 36 | 36 | 30 | 7 | 0 |
| 8.6 | 33 | 32 | 42 | 6 | 0 |
| 8.7 | 28 | 18 | 39 | 8 | 1 |
| 8.8 | 34 | 28 | 31 | 3 | 3 |
| 8.9 | 30 | 31 | 37 | 2 | 5 |
| 9 | 44 | 24 | 34 | 3 | 0 |
| 9.1 | 45 | 28 | 37 | 3 | 0 |
| 9.2 | 42 | 23 | 35 | 4 | 1 |
| 9.3 | 39 | 22 | 24 | 6 | 1 |
| 9.4 | 25 | 27 | 42 | 2 | 4 |
| 9.5 | 52 | 30 | 31 | 2 | 1 |
| 9.6 | 39 | 19 | 29 | 6 | 2 |
| 9.7 | 39 | 21 | 34 | 2 | 0 |
| 9.8 | 39 | 24 | 26 | 2 | 1 |
| 9.9 | 27 | 23 | 31 | 2 | 0 |
| 10 | 25 | 22 | 36 | 2 | 0 |
| 10.1 | 23 | 25 | 35 | 1 | 0 |
| 10.2 | 28 | 29 | 30 | 2 | 0 |
| 10.3 | 32 | 27 | 39 | 2 | 0 |
| 10.4 | 29 | 1 | 35 | 2 | 0 |
| 10.5 | 9 | 19 | 38 | 1 | 1 |
| 10.6 | 16 | 28 | 47 | 1 | 2 |
| 10.7 | 18 | 23 | 38 | 1 | 0 |
| 10.8 | 17 | 20 | 48 | 1 | 0 |
| 10.9 | 14 | 20 | 50 | 1 | 0 |
| 11 | 30 | 25 | 43 | 6 | 0 |
| 11.1 | 13 | 26 | 53 | 1 | 0 |
| 11.2 | 17 | 25 | 41 | 2 | 0 |
| 11.3 | 16 | 24 | 43 | 5 | 0 |
| 11.4 | 17 | 26 | 49 | 4 | 0 |
| 11.5 | 14 | 20 | 49 | 5 | 0 |
| 11.6 | 17 | 26 | 60 | 4 | 0 |
| 11.7 | 19 | 33 | 59 | 6 | 0 |
| 11.8 | 28 | 28 | 66 | 3 | 0 |
| 11.9 | 26 | 23 | 39 | 6 | 5 |
| 12 | 29 | 26 | 54 | 7 | 2 |
| 12.1 | 29 | 22 | 41 | 3 | 2 |
| 12.2 | 26 | 32 | 41 | 6 | 1 |
| 12.3 | 31 | 32 | 47 | 3 | 0 |
| 12.4 | 17 | 29 | 50 | 4 | 0 |
| 12.5 | 25 | 29 | 54 | 2 | 0 |
| 12.6 | 21 | 34 | 56 | 2 | 0 |
| 12.7 | 19 | 37 | 54 | 3 | 0 |
| 12.8 | 20 | 31 | 49 | 2 | 1 |
| 12.9 | 19 | 35 | 40 | 6 | 1 |
| 13 | 20 | 30 | 63 | 3 | 0 |
| 13.1 | 22 | 30 | 57 | 2 | 1 |
| 13.2 | 21 | 26 | 64 | 1 | 1 |
| 13.3 | 18 | 24 | 39 | 3 | 1 |
| 13.4 | 20 | 30 | 23 | 1 | 0 |
| 13.5 | 17 | 23 | 44 | 6 | 0 |
| 13.6 | 23 | 23 | 48 | 4 | 1 |
| 13.7 | 17 | 23 | 37 | 2 | 4 |
| 13.8 | 20 | 25 | 40 | 1 | 0 |
| 13.9 | 12 | 30 | 42 | 4 | 0 |
| 14 | 13 | 27 | 37 | 3 | 1 |
| 14.1 | 11 | 23 | 44 | 2 | 1 |
| 14.2 | 9 | 28 | 47 | 3 | 2 |
| 14.3 | 16 | 19 | 43 | 4 | 3 |
| 14.4 | 15 | 33 | 37 | 5 | 2 |
| 14.5 | 17 | 23 | 45 | 2 | 3 |
| 14.6 | 17 | 24 | 40 | 3 | 4 |
| 14.7 | 9 | 28 | 41 | 3 | 4 |
| 14.8 | 6 | 32 | 36 | 4 | 0 |
| 14.9 | 6 | 21 | 40 | 2 | 1 |
| 15 | 10 | 29 | 45 | 3 | 1 |
| 15.1 | 7 | 19 | 45 | 5 | 0 |
| 15.2 | 15 | 25 | 50 | 0 | 0 |
| 15.3 | 16 | 27 | 37 | 4 | 1 |
| 15.4 | 9 | 18 | 31 | 2 | 0 |
| 15.5 | 17 | 26 | 37 | 1 | 0 |
| 15.6 | 25 | 28 | 39 | 0 | 0 |
| 15.7 | 25 | 24 | 46 | 2 | 1 |
| 15.8 | 31 | 24 | 42 | 2 | 1 |
| 15.9 | 20 | 23 | 39 | 1 | 2 |
| 16 | 19 | 22 | 29 | 4 | 3 |
| 16.1 | 11 | 26 | 40 | 3 | 3 |
| 16.2 | 11 | 33 | 48 | 5 | 1 |
| 16.3 | 7 | 29 | 46 | 2 | 1 |
| 16.4 | 18 | 24 | 39 | 0 | 3 |
| 16.5 | 23 | 18 | 37 | 1 | 1 |
| 16.6 | 19 | 24 | 47 | 1 | 0 |
| 16.7 | 20 | 24 | 47 | 3 | 0 |
| 16.8 | 20 | 25 | 54 | 0 | 1 |
| 16.9 | 17 | 30 | 43 | 1 | 0 |
| 17 | 21 | 30 | 53 | 2 | 0 |
| 17.1 | 23 | 28 | 42 | 0 | 0 |
| 17.2 | 24 | 30 | 36 | 3 | 1 |
| 17.3 | 23 | 29 | 41 | 2 | 2 |
| 17.4 | 20 | 21 | 41 | 2 | 1 |
| 17.5 | 22 | 26 | 46 | 1 | 1 |
| 17.6 | 27 | 26 | 30 | 4 | 0 |
| 17.7 | 28 | 23 | 37 | 3 | 3 |
| 17.8 | 31 | 32 | 42 | 2 | 2 |
| 17.9 | 22 | 19 | 44 | 3 | 0 |
| 18 | 18 | 25 | 42 | 6 | 0 |
| 18.1 | 20 | 28 | 34 | 1 | 0 |
| 18.2 | 19 | 27 | 40 | 3 | 1 |
| 18.3 | 9 | 25 | 38 | 0 | 1 |
| 18.4 | 11 | 26 | 47 | 2 | 0 |
| 18.5 | 14 | 28 | 43 | 1 | 0 |
| 18.6 | 24 | 27 | 45 | 0 | 0 |
| 18.7 | 18 | 25 | 40 | 0 | 1 |
| 18.8 | 15 | 30 | 31 | 0 | 0 |
| 18.9 | 11 | 24 | 33 | 2 | 1 |
| 19 | 11 | 33 | 28 | 3 | 0 |
| 19.1 | 7 | 33 | 26 | 0 | 2 |
| 19.2 | 17 | 30 | 29 | 2 | 3 |
| 19.3 | 20 | 27 | 27 | 2 | 0 |
| 19.4 | 10 | 32 | 22 | 1 | 2 |
| 19.5 | 11 | 22 | 28 | 0 | 1 |
| 19.6 | 11 | 26 | 27 | 2 | 3 |
| 19.7 | 22 | 26 | 25 | 1 | 1 |
| 19.8 | 12 | 29 | 31 | 10 | 1 |
| 19.9 | 21 | 20 | 39 | 1 | 0 |

Histones - CLOCK_bHLH (BMAL1S78)

Number of contacts

| Frame | 1st | 2nd | 3rd | 4th | 5th |
| --- | --- | --- | --- | --- | --- |
| 0 | 1 | 0 | 4 | 5 | 0 |
| 0.1 | 2 | 0 | 5 | 0 | 0 |
| 0.2 | 1 | 0 | 3 | 0 | 0 |
| 0.3 | 0 | 0 | 1 | 0 | 1 |
| 0.4 | 1 | 0 | 2 | 3 | 0 |
| 0.5 | 0 | 0 | 3 | 3 | 0 |
| 0.6 | 0 | 0 | 4 | 3 | 0 |
| 0.7 | 0 | 2 | 2 | 2 | 0 |
| 0.8 | 0 | 0 | 3 | 4 | 0 |
| 0.9 | 0 | 0 | 3 | 3 | 0 |
| 1 | 0 | 0 | 1 | 4 | 0 |
| 1.1 | 0 | 0 | 1 | 2 | 1 |
| 1.2 | 0 | 0 | 3 | 5 | 0 |
| 1.3 | 0 | 0 | 1 | 7 | 0 |
| 1.4 | 0 | 0 | 0 | 7 | 0 |
| 1.5 | 0 | 0 | 2 | 4 | 0 |
| 1.6 | 0 | 0 | 0 | 5 | 0 |
| 1.7 | 0 | 0 | 1 | 9 | 0 |
| 1.8 | 0 | 0 | 2 | 6 | 0 |
| 1.9 | 0 | 0 | 2 | 2 | 0 |
| 2 | 0 | 0 | 0 | 2 | 1 |
| 2.1 | 1 | 0 | 1 | 6 | 0 |
| 2.2 | 1 | 0 | 2 | 9 | 0 |
| 2.3 | 1 | 4 | 1 | 5 | 0 |
| 2.4 | 2 | 0 | 1 | 6 | 0 |
| 2.5 | 2 | 0 | 0 | 7 | 0 |
| 2.6 | 2 | 0 | 2 | 6 | 0 |
| 2.7 | 1 | 0 | 0 | 8 | 1 |
| 2.8 | 2 | 0 | 2 | 6 | 0 |
| 2.9 | 4 | 0 | 0 | 3 | 1 |
| 3 | 9 | 0 | 2 | 3 | 1 |
| 3.1 | 10 | 0 | 3 | 3 | 2 |
| 3.2 | 7 | 0 | 0 | 4 | 0 |
| 3.3 | 11 | 0 | 0 | 6 | 2 |
| 3.4 | 12 | 0 | 2 | 4 | 6 |
| 3.5 | 8 | 0 | 2 | 2 | 3 |
| 3.6 | 6 | 0 | 1 | 5 | 9 |
| 3.7 | 11 | 0 | 2 | 5 | 10 |
| 3.8 | 12 | 0 | 1 | 3 | 5 |
| 3.9 | 10 | 0 | 0 | 4 | 3 |
| 4 | 8 | 0 | 2 | 7 | 5 |
| 4.1 | 11 | 0 | 1 | 4 | 5 |
| 4.2 | 11 | 0 | 0 | 5 | 4 |
| 4.3 | 10 | 0 | 1 | 4 | 6 |
| 4.4 | 10 | 0 | 3 | 6 | 6 |
| 4.5 | 10 | 0 | 1 | 5 | 5 |
| 4.6 | 8 | 0 | 1 | 4 | 3 |
| 4.7 | 9 | 0 | 2 | 5 | 1 |
| 4.8 | 7 | 0 | 3 | 5 | 4 |
| 4.9 | 9 | 0 | 3 | 5 | 5 |
| 5 | 8 | 0 | 3 | 6 | 5 |
| 5.1 | 8 | 0 | 3 | 6 | 5 |
| 5.2 | 11 | 0 | 3 | 5 | 2 |
| 5.3 | 10 | 0 | 2 | 6 | 3 |
| 5.4 | 10 | 1 | 1 | 6 | 5 |
| 5.5 | 5 | 1 | 1 | 4 | 7 |
| 5.6 | 10 | 1 | 2 | 5 | 4 |
| 5.7 | 10 | 0 | 2 | 7 | 8 |
| 5.8 | 8 | 0 | 1 | 6 | 7 |
| 5.9 | 6 | 0 | 2 | 5 | 10 |
| 6 | 9 | 0 | 2 | 6 | 10 |
| 6.1 | 6 | 0 | 2 | 5 | 11 |
| 6.2 | 6 | 0 | 2 | 7 | 4 |
| 6.3 | 12 | 1 | 3 | 6 | 4 |
| 6.4 | 9 | 0 | 1 | 5 | 4 |
| 6.5 | 9 | 0 | 2 | 10 | 6 |
| 6.6 | 9 | 0 | 1 | 9 | 5 |
| 6.7 | 9 | 0 | 2 | 11 | 4 |
| 6.8 | 11 | 1 | 3 | 7 | 2 |
| 6.9 | 12 | 0 | 2 | 9 | 4 |
| 7 | 9 | 0 | 2 | 11 | 4 |
| 7.1 | 8 | 0 | 2 | 7 | 4 |
| 7.2 | 11 | 1 | 2 | 8 | 9 |
| 7.3 | 11 | 0 | 1 | 11 | 3 |
| 7.4 | 12 | 0 | 2 | 6 | 2 |
| 7.5 | 9 | 0 | 5 | 5 | 7 |
| 7.6 | 12 | 0 | 3 | 4 | 8 |
| 7.7 | 8 | 0 | 3 | 5 | 3 |
| 7.8 | 7 | 1 | 2 | 3 | 5 |
| 7.9 | 12 | 0 | 1 | 5 | 3 |
| 8 | 10 | 0 | 2 | 4 | 5 |
| 8.1 | 13 | 0 | 1 | 5 | 0 |
| 8.2 | 7 | 0 | 2 | 5 | 2 |
| 8.3 | 7 | 0 | 2 | 6 | 4 |
| 8.4 | 8 | 0 | 1 | 6 | 2 |
| 8.5 | 8 | 0 | 3 | 5 | 4 |
| 8.6 | 13 | 0 | 1 | 5 | 5 |
| 8.7 | 12 | 0 | 1 | 5 | 3 |
| 8.8 | 15 | 0 | 2 | 11 | 5 |
| 8.9 | 11 | 0 | 1 | 5 | 3 |
| 9 | 12 | 0 | 2 | 10 | 5 |
| 9.1 | 9 | 0 | 2 | 4 | 5 |
| 9.2 | 10 | 0 | 1 | 4 | 6 |
| 9.3 | 7 | 0 | 2 | 2 | 8 |
| 9.4 | 8 | 0 | 1 | 3 | 7 |
| 9.5 | 10 | 0 | 2 | 4 | 7 |
| 9.6 | 8 | 0 | 2 | 4 | 3 |
| 9.7 | 7 | 0 | 1 | 6 | 5 |
| 9.8 | 8 | 0 | 5 | 4 | 2 |
| 9.9 | 11 | 0 | 2 | 6 | 3 |
| 10 | 11 | 0 | 1 | 4 | 7 |
| 10.1 | 12 | 0 | 0 | 8 | 5 |
| 10.2 | 12 | 0 | 0 | 5 | 7 |
| 10.3 | 13 | 0 | 1 | 4 | 4 |
| 10.4 | 9 | 0 | 2 | 3 | 3 |
| 10.5 | 10 | 0 | 2 | 8 | 3 |
| 10.6 | 9 | 0 | 2 | 5 | 6 |
| 10.7 | 12 | 0 | 1 | 6 | 0 |
| 10.8 | 16 | 0 | 2 | 8 | 5 |
| 10.9 | 6 | 0 | 1 | 7 | 3 |
| 11 | 5 | 0 | 2 | 4 | 4 |
| 11.1 | 10 | 0 | 1 | 13 | 5 |
| 11.2 | 12 | 0 | 2 | 1 | 3 |
| 11.3 | 8 | 0 | 2 | 2 | 2 |
| 11.4 | 9 | 0 | 2 | 0 | 5 |
| 11.5 | 13 | 0 | 3 | 3 | 4 |
| 11.6 | 6 | 0 | 4 | 2 | 4 |
| 11.7 | 11 | 0 | 1 | 2 | 1 |
| 11.8 | 7 | 1 | 3 | 5 | 2 |
| 11.9 | 8 | 0 | 3 | 8 | 3 |
| 12 | 10 | 0 | 1 | 12 | 2 |
| 12.1 | 7 | 0 | 1 | 7 | 3 |
| 12.2 | 10 | 0 | 1 | 9 | 2 |
| 12.3 | 11 | 0 | 1 | 14 | 3 |
| 12.4 | 12 | 0 | 1 | 9 | 5 |
| 12.5 | 9 | 0 | 3 | 22 | 2 |
| 12.6 | 12 | 0 | 3 | 8 | 4 |
| 12.7 | 14 | 0 | 2 | 15 | 4 |
| 12.8 | 8 | 0 | 2 | 19 | 6 |
| 12.9 | 12 | 0 | 2 | 16 | 3 |
| 13 | 12 | 0 | 2 | 11 | 2 |
| 13.1 | 11 | 0 | 5 | 14 | 4 |
| 13.2 | 12 | 0 | 3 | 16 | 1 |
| 13.3 | 10 | 0 | 2 | 14 | 2 |
| 13.4 | 10 | 0 | 2 | 9 | 5 |
| 13.5 | 6 | 0 | 1 | 18 | 4 |
| 13.6 | 11 | 0 | 0 | 11 | 4 |
| 13.7 | 14 | 0 | 2 | 16 | 7 |
| 13.8 | 10 | 0 | 1 | 18 | 5 |
| 13.9 | 10 | 0 | 1 | 24 | 3 |
| 14 | 7 | 0 | 1 | 22 | 5 |
| 14.1 | 7 | 0 | 1 | 13 | 5 |
| 14.2 | 9 | 0 | 1 | 9 | 6 |
| 14.3 | 9 | 0 | 0 | 16 | 3 |
| 14.4 | 10 | 0 | 2 | 6 | 3 |
| 14.5 | 10 | 0 | 1 | 16 | 2 |
| 14.6 | 11 | 0 | 2 | 9 | 6 |
| 14.7 | 11 | 0 | 4 | 11 | 1 |
| 14.8 | 11 | 1 | 1 | 8 | 2 |
| 14.9 | 12 | 0 | 2 | 11 | 1 |
| 15 | 14 | 0 | 1 | 16 | 4 |
| 15.1 | 7 | 0 | 1 | 23 | 5 |
| 15.2 | 9 | 0 | 1 | 27 | 5 |
| 15.3 | 10 | 0 | 2 | 26 | 1 |
| 15.4 | 13 | 1 | 3 | 23 | 7 |
| 15.5 | 7 | 0 | 1 | 22 | 7 |
| 15.6 | 12 | 0 | 1 | 19 | 2 |
| 15.7 | 9 | 0 | 1 | 19 | 6 |
| 15.8 | 9 | 0 | 2 | 21 | 2 |
| 15.9 | 11 | 0 | 2 | 32 | 4 |
| 16 | 18 | 0 | 1 | 19 | 3 |
| 16.1 | 12 | 0 | 2 | 14 | 3 |
| 16.2 | 8 | 0 | 1 | 16 | 5 |
| 16.3 | 6 | 0 | 2 | 20 | 4 |
| 16.4 | 10 | 0 | 1 | 21 | 7 |
| 16.5 | 9 | 1 | 1 | 20 | 4 |
| 16.6 | 14 | 0 | 2 | 30 | 5 |
| 16.7 | 11 | 0 | 2 | 17 | 5 |
| 16.8 | 11 | 0 | 2 | 16 | 7 |
| 16.9 | 8 | 0 | 0 | 24 | 5 |
| 17 | 12 | 0 | 1 | 13 | 6 |
| 17.1 | 15 | 0 | 2 | 20 | 5 |
| 17.2 | 8 | 0 | 2 | 16 | 4 |
| 17.3 | 5 | 0 | 2 | 22 | 1 |
| 17.4 | 8 | 0 | 0 | 25 | 6 |
| 17.5 | 5 | 0 | 1 | 21 | 5 |
| 17.6 | 10 | 0 | 1 | 26 | 2 |
| 17.7 | 7 | 0 | 2 | 26 | 1 |
| 17.8 | 10 | 0 | 1 | 14 | 3 |
| 17.9 | 9 | 0 | 1 | 22 | 8 |
| 18 | 5 | 0 | 1 | 18 | 3 |
| 18.1 | 11 | 0 | 1 | 14 | 7 |
| 18.2 | 7 | 0 | 0 | 33 | 4 |
| 18.3 | 10 | 0 | 0 | 29 | 7 |
| 18.4 | 9 | 0 | 0 | 27 | 9 |
| 18.5 | 12 | 0 | 2 | 30 | 5 |
| 18.6 | 10 | 0 | 0 | 29 | 10 |
| 18.7 | 13 | 0 | 0 | 32 | 4 |
| 18.8 | 8 | 0 | 0 | 21 | 6 |
| 18.9 | 6 | 0 | 0 | 16 | 8 |
| 19 | 9 | 0 | 0 | 12 | 5 |
| 19.1 | 7 | 2 | 1 | 13 | 6 |
| 19.2 | 8 | 0 | 0 | 22 | 4 |
| 19.3 | 8 | 0 | 0 | 22 | 7 |
| 19.4 | 8 | 0 | 1 | 22 | 5 |
| 19.5 | 12 | 0 | 0 | 22 | 9 |
| 19.6 | 11 | 0 | 1 | 17 | 5 |
| 19.7 | 9 | 0 | 1 | 17 | 6 |
| 19.8 | 10 | 0 | 0 | 24 | 7 |
| 19.9 | 9 | 0 | 1 | 21 | 14 |

Histones - CLOCK_bHLH (CLOCKS38/42)

Number of contacts

| Frame | 1st | 2nd | 3rd | 4th | 5th |
| --- | --- | --- | --- | --- | --- |
| 0 | 11 | 12 | 0 | 4 | 2 |
| 0.1 | 11 | 13 | 0 | 4 | 2 |
| 0.2 | 6 | 5 | 0 | 10 | 3 |
| 0.3 | 8 | 3 | 1 | 6 | 2 |
| 0.4 | 0 | 2 | 1 | 8 | 4 |
| 0.5 | 0 | 0 | 0 | 10 | 4 |
| 0.6 | 0 | 0 | 0 | 6 | 5 |
| 0.7 | 2 | 0 | 2 | 5 | 4 |
| 0.8 | 6 | 0 | 0 | 5 | 4 |
| 0.9 | 2 | 0 | 0 | 6 | 3 |
| 1 | 2 | 0 | 1 | 5 | 21 |
| 1.1 | 3 | 0 | 2 | 1 | 19 |
| 1.2 | 5 | 0 | 2 | 1 | 23 |
| 1.3 | 5 | 0 | 4 | 1 | 23 |
| 1.4 | 5 | 0 | 0 | 1 | 12 |
| 1.5 | 22 | 0 | 1 | 1 | 8 |
| 1.6 | 36 | 0 | 2 | 0 | 22 |
| 1.7 | 46 | 0 | 1 | 0 | 18 |
| 1.8 | 26 | 0 | 1 | 4 | 4 |
| 1.9 | 17 | 0 | 0 | 6 | 28 |
| 2 | 14 | 0 | 0 | 7 | 9 |
| 2.1 | 2 | 0 | 1 | 6 | 10 |
| 2.2 | 16 | 0 | 0 | 4 | 6 |
| 2.3 | 25 | 0 | 0 | 4 | 16 |
| 2.4 | 26 | 0 | 1 | 6 | 8 |
| 2.5 | 28 | 0 | 0 | 6 | 12 |
| 2.6 | 26 | 0 | 0 | 3 | 8 |
| 2.7 | 26 | 0 | 0 | 3 | 8 |
| 2.8 | 23 | 0 | 3 | 8 | 8 |
| 2.9 | 33 | 0 | 1 | 9 | 7 |
| 3 | 16 | 0 | 1 | 6 | 23 |
| 3.1 | 20 | 0 | 1 | 7 | 35 |
| 3.2 | 22 | 0 | 0 | 2 | 29 |
| 3.3 | 17 | 0 | 1 | 2 | 32 |
| 3.4 | 1 | 0 | 1 | 4 | 27 |
| 3.5 | 4 | 0 | 1 | 3 | 29 |
| 3.6 | 4 | 0 | 0 | 5 | 29 |
| 3.7 | 2 | 0 | 1 | 8 | 37 |
| 3.8 | 3 | 0 | 1 | 6 | 43 |
| 3.9 | 2 | 0 | 1 | 4 | 33 |
| 4 | 9 | 0 | 1 | 9 | 29 |
| 4.1 | 0 | 0 | 1 | 7 | 33 |
| 4.2 | 1 | 0 | 1 | 7 | 24 |
| 4.3 | 26 | 0 | 0 | 7 | 39 |
| 4.4 | 13 | 0 | 1 | 7 | 42 |
| 4.5 | 18 | 0 | 0 | 5 | 34 |
| 4.6 | 18 | 0 | 0 | 8 | 37 |
| 4.7 | 15 | 0 | 0 | 7 | 40 |
| 4.8 | 17 | 0 | 1 | 7 | 28 |
| 4.9 | 6 | 0 | 1 | 7 | 37 |
| 5 | 2 | 0 | 0 | 2 | 36 |
| 5.1 | 13 | 0 | 0 | 1 | 40 |
| 5.2 | 1 | 0 | 0 | 2 | 46 |
| 5.3 | 1 | 0 | 0 | 1 | 42 |
| 5.4 | 1 | 0 | 1 | 7 | 40 |
| 5.5 | 0 | 0 | 1 | 7 | 37 |
| 5.6 | 0 | 0 | 2 | 8 | 39 |
| 5.7 | 5 | 0 | 0 | 7 | 42 |
| 5.8 | 8 | 0 | 0 | 12 | 37 |
| 5.9 | 4 | 0 | 0 | 4 | 42 |
| 6 | 8 | 0 | 0 | 2 | 40 |
| 6.1 | 5 | 0 | 0 | 7 | 39 |
| 6.2 | 7 | 0 | 0 | 6 | 37 |
| 6.3 | 4 | 0 | 1 | 6 | 39 |
| 6.4 | 5 | 0 | 1 | 9 | 42 |
| 6.5 | 6 | 0 | 0 | 6 | 44 |
| 6.6 | 9 | 0 | 2 | 6 | 43 |
| 6.7 | 6 | 0 | 0 | 6 | 43 |
| 6.8 | 4 | 0 | 1 | 3 | 44 |
| 6.9 | 2 | 0 | 1 | 6 | 37 |
| 7 | 7 | 0 | 3 | 4 | 41 |
| 7.1 | 2 | 0 | 0 | 6 | 29 |
| 7.2 | 6 | 0 | 0 | 6 | 37 |
| 7.3 | 4 | 0 | 2 | 9 | 33 |
| 7.4 | 3 | 1 | 4 | 5 | 32 |
| 7.5 | 7 | 0 | 1 | 10 | 37 |
| 7.6 | 10 | 0 | 0 | 5 | 38 |
| 7.7 | 4 | 0 | 4 | 6 | 41 |
| 7.8 | 4 | 0 | 1 | 10 | 38 |
| 7.9 | 9 | 0 | 1 | 7 | 41 |
| 8 | 1 | 0 | 3 | 5 | 39 |
| 8.1 | 2 | 0 | 3 | 7 | 39 |
| 8.2 | 11 | 1 | 5 | 11 | 38 |
| 8.3 | 11 | 0 | 9 | 9 | 50 |
| 8.4 | 7 | 0 | 5 | 7 | 41 |
| 8.5 | 0 | 1 | 6 | 7 | 45 |
| 8.6 | 3 | 2 | 6 | 8 | 41 |
| 8.7 | 2 | 1 | 5 | 10 | 46 |
| 8.8 | 8 | 0 | 10 | 9 | 39 |
| 8.9 | 3 | 0 | 6 | 3 | 44 |
| 9 | 2 | 0 | 7 | 8 | 46 |
| 9.1 | 6 | 1 | 3 | 7 | 44 |
| 9.2 | 16 | 0 | 6 | 7 | 42 |
| 9.3 | 9 | 0 | 6 | 16 | 40 |
| 9.4 | 8 | 0 | 8 | 10 | 41 |
| 9.5 | 19 | 0 | 5 | 13 | 45 |
| 9.6 | 24 | 0 | 5 | 18 | 37 |
| 9.7 | 21 | 0 | 4 | 19 | 37 |
| 9.8 | 13 | 0 | 5 | 16 | 32 |
| 9.9 | 25 | 0 | 5 | 13 | 37 |
| 10 | 25 | 2 | 5 | 19 | 33 |
| 10.1 | 22 | 0 | 7 | 8 | 40 |
| 10.2 | 22 | 0 | 5 | 9 | 35 |
| 10.3 | 17 | 0 | 6 | 16 | 40 |
| 10.4 | 8 | 0 | 4 | 21 | 45 |
| 10.5 | 20 | 0 | 4 | 8 | 38 |
| 10.6 | 16 | 0 | 6 | 7 | 37 |
| 10.7 | 35 | 1 | 6 | 6 | 36 |
| 10.8 | 29 | 0 | 3 | 8 | 37 |
| 10.9 | 23 | 0 | 4 | 9 | 33 |
| 11 | 20 | 0 | 3 | 5 | 50 |
| 11.1 | 19 | 0 | 5 | 8 | 41 |
| 11.2 | 19 | 1 | 3 | 7 | 48 |
| 11.3 | 21 | 0 | 5 | 6 | 41 |
| 11.4 | 20 | 0 | 6 | 6 | 39 |
| 11.5 | 22 | 0 | 4 | 6 | 50 |
| 11.6 | 25 | 0 | 4 | 6 | 42 |
| 11.7 | 26 | 0 | 5 | 5 | 43 |
| 11.8 | 19 | 0 | 1 | 4 | 50 |
| 11.9 | 24 | 0 | 3 | 3 | 51 |
| 12 | 28 | 0 | 2 | 5 | 43 |
| 12.1 | 17 | 2 | 1 | 5 | 35 |
| 12.2 | 18 | 0 | 1 | 8 | 36 |
| 12.3 | 19 | 0 | 1 | 5 | 39 |
| 12.4 | 22 | 0 | 1 | 5 | 41 |
| 12.5 | 22 | 0 | 1 | 4 | 36 |
| 12.6 | 21 | 0 | 1 | 6 | 51 |
| 12.7 | 20 | 0 | 1 | 6 | 39 |
| 12.8 | 28 | 0 | 2 | 6 | 39 |
| 12.9 | 20 | 0 | 1 | 4 | 19 |
| 13 | 25 | 0 | 1 | 4 | 28 |
| 13.1 | 27 | 0 | 3 | 5 | 32 |
| 13.2 | 22 | 0 | 1 | 8 | 23 |
| 13.3 | 18 | 0 | 1 | 5 | 34 |
| 13.4 | 20 | 0 | 1 | 5 | 22 |
| 13.5 | 19 | 0 | 2 | 5 | 24 |
| 13.6 | 20 | 1 | 1 | 7 | 33 |
| 13.7 | 18 | 0 | 1 | 6 | 33 |
| 13.8 | 15 | 0 | 1 | 7 | 34 |
| 13.9 | 20 | 0 | 1 | 8 | 34 |
| 14 | 21 | 0 | 1 | 6 | 32 |
| 14.1 | 25 | 0 | 1 | 2 | 28 |
| 14.2 | 18 | 0 | 2 | 7 | 30 |
| 14.3 | 19 | 0 | 1 | 7 | 17 |
| 14.4 | 19 | 5 | 1 | 7 | 24 |
| 14.5 | 17 | 11 | 1 | 7 | 16 |
| 14.6 | 22 | 16 | 3 | 5 | 42 |
| 14.7 | 12 | 1 | 1 | 6 | 29 |
| 14.8 | 26 | 0 | 1 | 5 | 35 |
| 14.9 | 22 | 3 | 1 | 3 | 32 |
| 15 | 25 | 2 | 1 | 3 | 37 |
| 15.1 | 23 | 1 | 1 | 3 | 40 |
| 15.2 | 28 | 1 | 1 | 2 | 32 |
| 15.3 | 27 | 2 | 1 | 3 | 34 |
| 15.4 | 31 | 1 | 0 | 7 | 26 |
| 15.5 | 23 | 1 | 2 | 6 | 32 |
| 15.6 | 14 | 1 | 1 | 8 | 43 |
| 15.7 | 22 | 2 | 1 | 5 | 32 |
| 15.8 | 13 | 11 | 1 | 9 | 32 |
| 15.9 | 16 | 1 | 1 | 6 | 34 |
| 16 | 22 | 11 | 0 | 5 | 34 |
| 16.1 | 36 | 7 | 3 | 5 | 25 |
| 16.2 | 13 | 1 | 5 | 6 | 32 |
| 16.3 | 26 | 2 | 3 | 7 | 35 |
| 16.4 | 14 | 2 | 2 | 5 | 35 |
| 16.5 | 20 | 10 | 1 | 6 | 39 |
| 16.6 | 19 | 4 | 0 | 4 | 39 |
| 16.7 | 19 | 16 | 1 | 7 | 40 |
| 16.8 | 18 | 1 | 1 | 4 | 33 |
| 16.9 | 20 | 2 | 1 | 7 | 37 |
| 17 | 17 | 1 | 1 | 4 | 35 |
| 17.1 | 23 | 0 | 1 | 5 | 38 |
| 17.2 | 25 | 2 | 1 | 3 | 48 |
| 17.3 | 27 | 3 | 1 | 3 | 43 |
| 17.4 | 17 | 4 | 3 | 4 | 43 |
| 17.5 | 23 | 0 | 3 | 5 | 40 |
| 17.6 | 24 | 7 | 3 | 6 | 37 |
| 17.7 | 21 | 7 | 1 | 5 | 40 |
| 17.8 | 17 | 7 | 1 | 3 | 36 |
| 17.9 | 22 | 7 | 1 | 3 | 44 |
| 18 | 16 | 18 | 0 | 6 | 39 |
| 18.1 | 25 | 7 | 1 | 5 | 31 |
| 18.2 | 23 | 20 | 1 | 4 | 48 |
| 18.3 | 25 | 18 | 2 | 5 | 37 |
| 18.4 | 28 | 8 | 0 | 4 | 36 |
| 18.5 | 22 | 18 | 1 | 3 | 45 |
| 18.6 | 23 | 19 | 0 | 4 | 31 |
| 18.7 | 36 | 9 | 1 | 3 | 33 |
| 18.8 | 30 | 0 | 3 | 4 | 34 |
| 18.9 | 36 | 3 | 4 | 4 | 42 |
| 19 | 29 | 14 | 3 | 5 | 41 |
| 19.1 | 14 | 21 | 1 | 4 | 48 |
| 19.2 | 23 | 5 | 3 | 8 | 38 |
| 19.3 | 30 | 11 | 1 | 3 | 44 |
| 19.4 | 25 | 15 | 3 | 8 | 37 |
| 19.5 | 27 | 37 | 0 | 3 | 36 |
| 19.6 | 31 | 29 | 1 | 5 | 32 |
| 19.7 | 24 | 32 | 1 | 5 | 38 |
| 19.8 | 33 | 22 | 1 | 3 | 35 |
| 19.9 | 23 | 15 | 1 | 4 | 32 |

Histones - CLOCK_bHLH (CLOCKS38/42/BMAL1S78)

Number of contacts

| Frame | 1st | 2nd | 3rd | 4th | 5th |
| --- | --- | --- | --- | --- | --- |
| 0 | 9 | 27 | 16 | 10 | 6 |
| 0.1 | 9 | 28 | 15 | 10 | 6 |
| 0.2 | 0 | 37 | 37 | 2 | 0 |
| 0.3 | 5 | 43 | 39 | 4 | 0 |
| 0.4 | 4 | 35 | 39 | 4 | 0 |
| 0.5 | 5 | 48 | 33 | 3 | 0 |
| 0.6 | 5 | 33 | 11 | 4 | 1 |
| 0.7 | 5 | 23 | 10 | 5 | 0 |
| 0.8 | 6 | 33 | 23 | 6 | 0 |
| 0.9 | 3 | 32 | 22 | 8 | 1 |
| 1 | 4 | 29 | 25 | 2 | 0 |
| 1.1 | 5 | 32 | 28 | 3 | 0 |
| 1.2 | 5 | 33 | 13 | 3 | 0 |
| 1.3 | 4 | 23 | 20 | 12 | 0 |
| 1.4 | 4 | 30 | 18 | 2 | 0 |
| 1.5 | 7 | 34 | 9 | 4 | 0 |
| 1.6 | 5 | 25 | 9 | 3 | 0 |
| 1.7 | 6 | 35 | 8 | 6 | 0 |
| 1.8 | 5 | 26 | 8 | 4 | 0 |
| 1.9 | 3 | 27 | 16 | 1 | 0 |
| 2 | 6 | 35 | 9 | 1 | 0 |
| 2.1 | 7 | 24 | 7 | 2 | 1 |
| 2.2 | 2 | 31 | 11 | 4 | 1 |
| 2.3 | 8 | 26 | 14 | 6 | 1 |
| 2.4 | 7 | 32 | 2 | 8 | 2 |
| 2.5 | 3 | 30 | 1 | 5 | 0 |
| 2.6 | 3 | 28 | 12 | 4 | 0 |
| 2.7 | 3 | 27 | 9 | 6 | 0 |
| 2.8 | 5 | 26 | 2 | 5 | 2 |
| 2.9 | 6 | 29 | 1 | 3 | 2 |
| 3 | 5 | 30 | 4 | 4 | 0 |
| 3.1 | 8 | 28 | 2 | 16 | 0 |
| 3.2 | 6 | 31 | 3 | 3 | 0 |
| 3.3 | 2 | 36 | 0 | 5 | 0 |
| 3.4 | 0 | 24 | 0 | 3 | 0 |
| 3.5 | 2 | 31 | 1 | 5 | 0 |
| 3.6 | 3 | 21 | 1 | 7 | 0 |
| 3.7 | 1 | 29 | 2 | 10 | 0 |
| 3.8 | 2 | 32 | 0 | 10 | 2 |
| 3.9 | 2 | 27 | 0 | 6 | 1 |
| 4 | 2 | 22 | 0 | 5 | 1 |
| 4.1 | 2 | 28 | 0 | 8 | 0 |
| 4.2 | 7 | 31 | 1 | 4 | 2 |
| 4.3 | 6 | 26 | 0 | 8 | 0 |
| 4.4 | 5 | 25 | 0 | 8 | 1 |
| 4.5 | 10 | 28 | 0 | 8 | 0 |
| 4.6 | 21 | 26 | 0 | 8 | 1 |
| 4.7 | 13 | 29 | 0 | 12 | 0 |
| 4.8 | 6 | 27 | 0 | 10 | 0 |
| 4.9 | 9 | 29 | 0 | 11 | 0 |
| 5 | 4 | 31 | 0 | 21 | 0 |
| 5.1 | 8 | 23 | 0 | 22 | 0 |
| 5.2 | 9 | 31 | 0 | 19 | 1 |
| 5.3 | 20 | 34 | 0 | 17 | 1 |
| 5.4 | 24 | 27 | 0 | 13 | 1 |
| 5.5 | 23 | 28 | 0 | 4 | 1 |
| 5.6 | 20 | 31 | 0 | 9 | 5 |
| 5.7 | 19 | 31 | 0 | 4 | 1 |
| 5.8 | 27 | 31 | 0 | 10 | 1 |
| 5.9 | 19 | 28 | 0 | 10 | 0 |
| 6 | 25 | 29 | 0 | 18 | 2 |
| 6.1 | 27 | 32 | 0 | 12 | 2 |
| 6.2 | 20 | 30 | 0 | 22 | 2 |
| 6.3 | 23 | 26 | 0 | 23 | 4 |
| 6.4 | 23 | 38 | 0 | 26 | 1 |
| 6.5 | 28 | 32 | 0 | 18 | 2 |
| 6.6 | 29 | 28 | 0 | 17 | 2 |
| 6.7 | 22 | 26 | 0 | 19 | 5 |
| 6.8 | 22 | 29 | 1 | 21 | 3 |
| 6.9 | 26 | 32 | 0 | 14 | 2 |
| 7 | 27 | 30 | 1 | 17 | 1 |
| 7.1 | 30 | 34 | 3 | 18 | 2 |
| 7.2 | 24 | 36 | 4 | 21 | 0 |
| 7.3 | 23 | 30 | 2 | 15 | 0 |
| 7.4 | 27 | 26 | 0 | 20 | 1 |
| 7.5 | 13 | 26 | 3 | 24 | 2 |
| 7.6 | 23 | 32 | 5 | 32 | 4 |
| 7.7 | 27 | 29 | 5 | 25 | 0 |
| 7.8 | 22 | 32 | 5 | 25 | 1 |
| 7.9 | 20 | 30 | 6 | 25 | 0 |
| 8 | 17 | 31 | 2 | 20 | 2 |
| 8.1 | 23 | 30 | 5 | 18 | 3 |
| 8.2 | 23 | 38 | 4 | 25 | 0 |
| 8.3 | 16 | 33 | 1 | 29 | 0 |
| 8.4 | 21 | 28 | 3 | 35 | 1 |
| 8.5 | 21 | 31 | 1 | 26 | 2 |
| 8.6 | 21 | 31 | 5 | 24 | 0 |
| 8.7 | 19 | 33 | 2 | 26 | 1 |
| 8.8 | 21 | 36 | 3 | 34 | 0 |
| 8.9 | 23 | 30 | 2 | 22 | 0 |
| 9 | 14 | 35 | 3 | 23 | 0 |
| 9.1 | 19 | 33 | 2 | 12 | 0 |
| 9.2 | 19 | 35 | 1 | 22 | 1 |
| 9.3 | 25 | 26 | 2 | 12 | 0 |
| 9.4 | 23 | 32 | 0 | 14 | 0 |
| 9.5 | 25 | 29 | 1 | 16 | 1 |
| 9.6 | 28 | 29 | 0 | 13 | 0 |
| 9.7 | 28 | 39 | 1 | 19 | 0 |
| 9.8 | 29 | 31 | 1 | 12 | 0 |
| 9.9 | 25 | 15 | 2 | 17 | 0 |
| 10 | 29 | 34 | 1 | 14 | 1 |
| 10.1 | 29 | 27 | 0 | 17 | 0 |
| 10.2 | 21 | 23 | 1 | 24 | 0 |
| 10.3 | 21 | 28 | 1 | 23 | 0 |
| 10.4 | 25 | 29 | 1 | 23 | 1 |
| 10.5 | 27 | 30 | 2 | 28 | 1 |
| 10.6 | 22 | 34 | 4 | 26 | 0 |
| 10.7 | 22 | 26 | 1 | 20 | 0 |
| 10.8 | 29 | 30 | 0 | 27 | 0 |
| 10.9 | 23 | 31 | 0 | 21 | 0 |
| 11 | 31 | 30 | 0 | 21 | 0 |
| 11.1 | 28 | 32 | 0 | 31 | 0 |
| 11.2 | 19 | 31 | 2 | 27 | 0 |
| 11.3 | 25 | 26 | 1 | 33 | 0 |
| 11.4 | 28 | 31 | 1 | 31 | 0 |
| 11.5 | 28 | 34 | 1 | 22 | 0 |
| 11.6 | 24 | 38 | 2 | 18 | 0 |
| 11.7 | 25 | 36 | 1 | 18 | 1 |
| 11.8 | 24 | 27 | 3 | 12 | 0 |
| 11.9 | 21 | 32 | 2 | 20 | 0 |
| 12 | 31 | 31 | 2 | 23 | 2 |
| 12.1 | 18 | 31 | 1 | 22 | 0 |
| 12.2 | 24 | 30 | 2 | 19 | 0 |
| 12.3 | 18 | 34 | 4 | 27 | 0 |
| 12.4 | 23 | 32 | 0 | 10 | 0 |
| 12.5 | 17 | 38 | 4 | 18 | 0 |
| 12.6 | 20 | 25 | 3 | 24 | 0 |
| 12.7 | 23 | 29 | 4 | 22 | 1 |
| 12.8 | 23 | 26 | 7 | 22 | 0 |
| 12.9 | 25 | 28 | 6 | 23 | 1 |
| 13 | 21 | 32 | 6 | 24 | 0 |
| 13.1 | 21 | 30 | 9 | 11 | 1 |
| 13.2 | 26 | 32 | 5 | 15 | 1 |
| 13.3 | 27 | 38 | 2 | 11 | 0 |
| 13.4 | 20 | 27 | 5 | 9 | 1 |
| 13.5 | 19 | 30 | 4 | 13 | 0 |
| 13.6 | 21 | 30 | 5 | 8 | 1 |
| 13.7 | 22 | 39 | 6 | 19 | 0 |
| 13.8 | 19 | 32 | 4 | 20 | 0 |
| 13.9 | 20 | 26 | 3 | 24 | 3 |
| 14 | 28 | 34 | 4 | 21 | 0 |
| 14.1 | 20 | 30 | 6 | 19 | 2 |
| 14.2 | 22 | 33 | 3 | 12 | 0 |
| 14.3 | 22 | 30 | 3 | 19 | 2 |
| 14.4 | 19 | 33 | 5 | 21 | 1 |
| 14.5 | 31 | 26 | 3 | 7 | 2 |
| 14.6 | 22 | 27 | 4 | 18 | 2 |
| 14.7 | 25 | 29 | 5 | 16 | 0 |
| 14.8 | 20 | 33 | 2 | 5 | 2 |
| 14.9 | 19 | 33 | 4 | 4 | 1 |
| 15 | 14 | 31 | 3 | 4 | 3 |
| 15.1 | 24 | 33 | 3 | 3 | 2 |
| 15.2 | 25 | 31 | 4 | 3 | 4 |
| 15.3 | 25 | 29 | 2 | 4 | 1 |
| 15.4 | 20 | 31 | 3 | 13 | 4 |
| 15.5 | 21 | 24 | 3 | 9 | 1 |
| 15.6 | 28 | 33 | 2 | 9 | 0 |
| 15.7 | 24 | 37 | 4 | 4 | 1 |
| 15.8 | 22 | 30 | 4 | 19 | 1 |
| 15.9 | 26 | 30 | 6 | 4 | 0 |
| 16 | 22 | 31 | 1 | 8 | 1 |
| 16.1 | 26 | 34 | 3 | 5 | 0 |
| 16.2 | 21 | 33 | 1 | 4 | 1 |
| 16.3 | 19 | 37 | 1 | 2 | 3 |
| 16.4 | 21 | 30 | 1 | 6 | 0 |
| 16.5 | 24 | 28 | 0 | 4 | 1 |
| 16.6 | 24 | 38 | 2 | 7 | 1 |
| 16.7 | 25 | 33 | 1 | 5 | 1 |
| 16.8 | 19 | 37 | 1 | 9 | 0 |
| 16.9 | 18 | 35 | 1 | 20 | 1 |
| 17 | 20 | 32 | 1 | 21 | 2 |
| 17.1 | 22 | 35 | 1 | 15 | 2 |
| 17.2 | 13 | 36 | 2 | 19 | 12 |
| 17.3 | 12 | 36 | 0 | 18 | 12 |
| 17.4 | 15 | 36 | 1 | 8 | 3 |
| 17.5 | 24 | 31 | 2 | 12 | 7 |
| 17.6 | 27 | 31 | 2 | 17 | 3 |
| 17.7 | 20 | 28 | 3 | 6 | 3 |
| 17.8 | 16 | 37 | 3 | 8 | 9 |
| 17.9 | 14 | 30 | 3 | 3 | 11 |
| 18 | 20 | 28 | 3 | 7 | 11 |
| 18.1 | 15 | 34 | 1 | 8 | 12 |
| 18.2 | 20 | 31 | 4 | 13 | 2 |
| 18.3 | 10 | 38 | 4 | 7 | 7 |
| 18.4 | 21 | 33 | 4 | 9 | 5 |
| 18.5 | 23 | 25 | 1 | 8 | 17 |
| 18.6 | 14 | 37 | 3 | 11 | 0 |
| 18.7 | 24 | 35 | 2 | 10 | 0 |
| 18.8 | 16 | 29 | 2 | 12 | 0 |
| 18.9 | 29 | 35 | 2 | 11 | 0 |
| 19 | 27 | 29 | 1 | 13 | 0 |
| 19.1 | 31 | 42 | 2 | 12 | 0 |
| 19.2 | 20 | 32 | 2 | 11 | 0 |
| 19.3 | 14 | 32 | 1 | 18 | 0 |
| 19.4 | 16 | 33 | 2 | 17 | 0 |
| 19.5 | 24 | 34 | 0 | 20 | 0 |
| 19.6 | 26 | 32 | 0 | 22 | 1 |
| 19.7 | 21 | 33 | 1 | 18 | 1 |
| 19.8 | 26 | 39 | 0 | 10 | 1 |
| 19.9 | 24 | 36 | 0 | 15 | 0 |

**Figure S5.** Timeline plots pertaining to number of contacts between histones and CLOCK bHLH for the systems studied.

Histones - CLOCK_PAS-B (nonphosphorylated)

Number of contacts

| Frame | 1st | 2nd | 3rd | 4th | 5th |
| --- | --- | --- | --- | --- | --- |
| 0 | 14 | 0 | 0 | 7 | 28 |
| 0.1 | 14 | 0 | 0 | 7 | 0 |
| 0.2 | 14 | 0 | 0 | 24 | 0 |
| 0.3 | 29 | 0 | 0 | 44 | 22 |
| 0.4 | 40 | 6 | 0 | 47 | 20 |
| 0.5 | 45 | 52 | 1 | 70 | 46 |
| 0.6 | 57 | 64 | 3 | 75 | 61 |
| 0.7 | 76 | 51 | 31 | 83 | 67 |
| 0.8 | 54 | 87 | 54 | 84 | 81 |
| 0.9 | 43 | 75 | 39 | 90 | 96 |
| 1 | 61 | 73 | 39 | 85 | 96 |
| 1.1 | 74 | 113 | 44 | 94 | 98 |
| 1.2 | 108 | 94 | 43 | 87 | 92 |
| 1.3 | 98 | 118 | 66 | 94 | 98 |
| 1.4 | 107 | 116 | 59 | 111 | 115 |
| 1.5 | 116 | 142 | 73 | 103 | 95 |
| 1.6 | 111 | 132 | 70 | 104 | 108 |
| 1.7 | 113 | 141 | 59 | 114 | 90 |
| 1.8 | 115 | 169 | 77 | 111 | 88 |
| 1.9 | 114 | 144 | 79 | 99 | 93 |
| 2 | 117 | 132 | 75 | 103 | 96 |
| 2.1 | 119 | 159 | 96 | 112 | 96 |
| 2.2 | 117 | 167 | 59 | 94 | 104 |
| 2.3 | 127 | 157 | 67 | 102 | 92 |
| 2.4 | 116 | 156 | 70 | 110 | 110 |
| 2.5 | 118 | 172 | 88 | 122 | 90 |
| 2.6 | 124 | 153 | 70 | 103 | 100 |
| 2.7 | 146 | 178 | 74 | 95 | 100 |
| 2.8 | 134 | 174 | 67 | 106 | 82 |
| 2.9 | 109 | 171 | 88 | 115 | 102 |
| 3 | 129 | 175 | 87 | 120 | 92 |
| 3.1 | 126 | 170 | 73 | 107 | 95 |
| 3.2 | 117 | 164 | 83 | 91 | 108 |
| 3.3 | 109 | 161 | 74 | 98 | 98 |
| 3.4 | 123 | 158 | 86 | 105 | 89 |
| 3.5 | 116 | 168 | 82 | 105 | 99 |
| 3.6 | 132 | 151 | 75 | 100 | 95 |
| 3.7 | 110 | 173 | 82 | 103 | 106 |
| 3.8 | 115 | 156 | 93 | 103 | 105 |
| 3.9 | 124 | 151 | 72 | 103 | 98 |
| 4 | 123 | 155 | 87 | 107 | 93 |
| 4.1 | 112 | 172 | 78 | 91 | 89 |
| 4.2 | 118 | 179 | 83 | 107 | 86 |
| 4.3 | 125 | 180 | 92 | 108 | 90 |
| 4.4 | 116 | 158 | 96 | 97 | 93 |
| 4.5 | 141 | 148 | 89 | 107 | 106 |
| 4.6 | 130 | 159 | 100 | 99 | 97 |
| 4.7 | 144 | 188 | 92 | 107 | 90 |
| 4.8 | 143 | 187 | 83 | 103 | 81 |
| 4.9 | 129 | 209 | 91 | 97 | 97 |
| 5 | 117 | 221 | 89 | 107 | 89 |
| 5.1 | 116 | 232 | 92 | 110 | 92 |
| 5.2 | 122 | 217 | 104 | 123 | 99 |
| 5.3 | 121 | 206 | 97 | 109 | 96 |
| 5.4 | 119 | 183 | 125 | 111 | 81 |
| 5.5 | 104 | 175 | 119 | 113 | 103 |
| 5.6 | 129 | 171 | 107 | 115 | 104 |
| 5.7 | 133 | 175 | 117 | 99 | 104 |
| 5.8 | 125 | 187 | 126 | 92 | 88 |
| 5.9 | 120 | 170 | 134 | 95 | 84 |
| 6 | 131 | 181 | 129 | 90 | 90 |
| 6.1 | 123 | 168 | 134 | 110 | 104 |
| 6.2 | 114 | 174 | 125 | 107 | 100 |
| 6.3 | 134 | 170 | 130 | 90 | 96 |
| 6.4 | 123 | 181 | 121 | 115 | 92 |
| 6.5 | 120 | 192 | 140 | 117 | 92 |
| 6.6 | 131 | 193 | 123 | 97 | 104 |
| 6.7 | 104 | 198 | 124 | 91 | 99 |
| 6.8 | 127 | 182 | 118 | 105 | 109 |
| 6.9 | 125 | 196 | 142 | 104 | 91 |
| 7 | 107 | 194 | 125 | 113 | 86 |
| 7.1 | 125 | 184 | 161 | 87 | 89 |
| 7.2 | 119 | 190 | 152 | 74 | 101 |
| 7.3 | 124 | 185 | 163 | 93 | 103 |
| 7.4 | 121 | 203 | 145 | 84 | 92 |
| 7.5 | 137 | 215 | 154 | 105 | 98 |
| 7.6 | 126 | 200 | 155 | 96 | 103 |
| 7.7 | 123 | 183 | 157 | 89 | 99 |
| 7.8 | 119 | 185 | 144 | 102 | 99 |
| 7.9 | 119 | 202 | 148 | 115 | 99 |
| 8 | 127 | 202 | 156 | 105 | 92 |
| 8.1 | 129 | 214 | 145 | 124 | 101 |
| 8.2 | 127 | 188 | 145 | 114 | 96 |
| 8.3 | 129 | 209 | 151 | 106 | 91 |
| 8.4 | 118 | 201 | 146 | 106 | 97 |
| 8.5 | 126 | 199 | 147 | 100 | 96 |
| 8.6 | 121 | 194 | 147 | 95 | 97 |
| 8.7 | 113 | 207 | 131 | 121 | 99 |
| 8.8 | 133 | 203 | 134 | 107 | 102 |
| 8.9 | 123 | 203 | 157 | 117 | 102 |
| 9 | 128 | 199 | 155 | 105 | 90 |
| 9.1 | 122 | 190 | 131 | 122 | 103 |
| 9.2 | 132 | 181 | 153 | 104 | 100 |
| 9.3 | 129 | 191 | 148 | 112 | 87 |
| 9.4 | 140 | 184 | 146 | 114 | 98 |
| 9.5 | 118 | 181 | 158 | 107 | 79 |
| 9.6 | 140 | 197 | 161 | 108 | 90 |
| 9.7 | 153 | 199 | 162 | 100 | 102 |
| 9.8 | 149 | 208 | 152 | 113 | 96 |
| 9.9 | 150 | 196 | 150 | 110 | 87 |
| 10 | 132 | 194 | 168 | 100 | 89 |
| 10.1 | 144 | 209 | 149 | 122 | 86 |
| 10.2 | 151 | 217 | 153 | 107 | 85 |
| 10.3 | 150 | 220 | 148 | 109 | 98 |
| 10.4 | 148 | 220 | 138 | 106 | 92 |
| 10.5 | 141 | 231 | 151 | 111 | 105 |
| 10.6 | 136 | 229 | 144 | 108 | 88 |
| 10.7 | 134 | 233 | 143 | 104 | 99 |
| 10.8 | 182 | 240 | 159 | 92 | 92 |
| 10.9 | 177 | 219 | 150 | 115 | 73 |
| 11 | 169 | 210 | 153 | 106 | 96 |
| 11.1 | 170 | 237 | 171 | 120 | 79 |
| 11.2 | 159 | 207 | 150 | 114 | 90 |
| 11.3 | 162 | 235 | 142 | 104 | 86 |
| 11.4 | 172 | 242 | 148 | 109 | 101 |
| 11.5 | 160 | 235 | 143 | 111 | 95 |
| 11.6 | 181 | 209 | 144 | 104 | 87 |
| 11.7 | 182 | 235 | 141 | 110 | 102 |
| 11.8 | 185 | 212 | 145 | 100 | 83 |
| 11.9 | 185 | 236 | 162 | 113 | 95 |
| 12 | 192 | 217 | 135 | 111 | 93 |
| 12.1 | 183 | 215 | 127 | 105 | 93 |
| 12.2 | 180 | 212 | 139 | 103 | 92 |
| 12.3 | 191 | 219 | 140 | 103 | 80 |
| 12.4 | 181 | 220 | 150 | 133 | 96 |
| 12.5 | 192 | 229 | 162 | 109 | 92 |
| 12.6 | 191 | 213 | 161 | 109 | 86 |
| 12.7 | 186 | 223 | 163 | 100 | 86 |
| 12.8 | 199 | 204 | 157 | 105 | 82 |
| 12.9 | 182 | 223 | 163 | 112 | 94 |
| 13 | 175 | 235 | 167 | 95 | 95 |
| 13.1 | 175 | 239 | 152 | 105 | 100 |
| 13.2 | 167 | 217 | 181 | 120 | 102 |
| 13.3 | 162 | 240 | 172 | 117 | 95 |
| 13.4 | 186 | 216 | 189 | 122 | 89 |
| 13.5 | 175 | 217 | 181 | 109 | 99 |
| 13.6 | 159 | 222 | 177 | 100 | 88 |
| 13.7 | 207 | 198 | 178 | 116 | 84 |
| 13.8 | 179 | 212 | 164 | 114 | 88 |
| 13.9 | 160 | 219 | 177 | 107 | 97 |
| 14 | 170 | 218 | 164 | 114 | 87 |
| 14.1 | 160 | 213 | 174 | 93 | 100 |
| 14.2 | 179 | 222 | 177 | 117 | 94 |
| 14.3 | 200 | 203 | 174 | 116 | 101 |
| 14.4 | 192 | 211 | 177 | 100 | 90 |
| 14.5 | 189 | 208 | 174 | 98 | 96 |
| 14.6 | 187 | 235 | 192 | 95 | 96 |
| 14.7 | 172 | 228 | 180 | 105 | 93 |
| 14.8 | 171 | 228 | 192 | 106 | 94 |
| 14.9 | 179 | 222 | 183 | 106 | 83 |
| 15 | 178 | 229 | 181 | 112 | 89 |
| 15.1 | 175 | 225 | 166 | 107 | 95 |
| 15.2 | 178 | 231 | 165 | 107 | 99 |
| 15.3 | 184 | 224 | 176 | 117 | 96 |
| 15.4 | 169 | 226 | 160 | 114 | 98 |
| 15.5 | 178 | 232 | 171 | 97 | 95 |
| 15.6 | 159 | 270 | 175 | 110 | 97 |
| 15.7 | 180 | 238 | 167 | 105 | 89 |
| 15.8 | 169 | 222 | 178 | 105 | 104 |
| 15.9 | 175 | 234 | 178 | 97 | 97 |
| 16 | 172 | 235 | 164 | 118 | 94 |
| 16.1 | 191 | 234 | 170 | 98 | 98 |
| 16.2 | 188 | 226 | 168 | 119 | 88 |
| 16.3 | 189 | 224 | 196 | 119 | 93 |
| 16.4 | 177 | 225 | 169 | 109 | 98 |
| 16.5 | 170 | 229 | 180 | 104 | 82 |
| 16.6 | 187 | 227 | 177 | 118 | 105 |
| 16.7 | 165 | 220 | 173 | 90 | 99 |
| 16.8 | 195 | 220 | 153 | 108 | 90 |
| 16.9 | 169 | 212 | 191 | 107 | 81 |
| 17 | 170 | 226 | 166 | 105 | 89 |
| 17.1 | 180 | 220 | 161 | 117 | 108 |
| 17.2 | 194 | 234 | 186 | 89 | 99 |
| 17.3 | 180 | 225 | 180 | 119 | 97 |
| 17.4 | 184 | 230 | 183 | 124 | 100 |
| 17.5 | 174 | 220 | 171 | 103 | 89 |
| 17.6 | 178 | 211 | 184 | 109 | 90 |
| 17.7 | 179 | 221 | 180 | 113 | 88 |
| 17.8 | 179 | 210 | 170 | 111 | 109 |
| 17.9 | 186 | 205 | 162 | 107 | 103 |
| 18 | 175 | 210 | 200 | 104 | 106 |
| 18.1 | 180 | 248 | 185 | 101 | 91 |
| 18.2 | 168 | 201 | 193 | 114 | 97 |
| 18.3 | 171 | 216 | 185 | 110 | 88 |
| 18.4 | 185 | 224 | 183 | 115 | 103 |
| 18.5 | 164 | 236 | 193 | 109 | 107 |
| 18.6 | 183 | 242 | 182 | 119 | 87 |
| 18.7 | 164 | 228 | 173 | 121 | 97 |
| 18.8 | 182 | 229 | 185 | 95 | 82 |
| 18.9 | 165 | 225 | 193 | 100 | 90 |
| 19 | 164 | 250 | 197 | 117 | 95 |
| 19.1 | 163 | 217 | 167 | 102 | 108 |
| 19.2 | 165 | 222 | 195 | 113 | 107 |
| 19.3 | 177 | 211 | 177 | 114 | 102 |
| 19.4 | 172 | 207 | 191 | 113 | 99 |
| 19.5 | 175 | 226 | 177 | 110 | 97 |
| 19.6 | 166 | 228 | 184 | 104 | 99 |
| 19.7 | 169 | 208 | 188 | 110 | 110 |
| 19.8 | 167 | 214 | 188 | 109 | 109 |
| 19.9 | 168 | 220 | 187 | 108 | 100 |

Histones - CLOCK_PAS-B (BMAL1S78)

Number of contacts

| Frame | 1st | 2nd | 3rd | 4th | 5th |
| --- | --- | --- | --- | --- | --- |
| 0 | 2 | 0 | 0 | 27 | 0 |
| 0.1 | 2 | 0 | 0 | 0 | 0 |
| 0.2 | 85 | 0 | 23 | 4 | 0 |
| 0.3 | 111 | 1 | 98 | 13 | 15 |
| 0.4 | 104 | 28 | 84 | 25 | 0 |
| 0.5 | 93 | 23 | 86 | 102 | 1 |
| 0.6 | 117 | 27 | 76 | 107 | 21 |
| 0.7 | 102 | 29 | 80 | 95 | 35 |
| 0.8 | 109 | 26 | 82 | 82 | 40 |
| 0.9 | 120 | 37 | 77 | 84 | 51 |
| 1 | 114 | 39 | 78 | 91 | 36 |
| 1.1 | 115 | 46 | 86 | 89 | 39 |
| 1.2 | 110 | 52 | 85 | 94 | 42 |
| 1.3 | 107 | 56 | 82 | 93 | 38 |
| 1.4 | 108 | 53 | 87 | 97 | 31 |
| 1.5 | 169 | 66 | 94 | 82 | 36 |
| 1.6 | 156 | 80 | 86 | 110 | 49 |
| 1.7 | 157 | 69 | 87 | 97 | 33 |
| 1.8 | 143 | 61 | 89 | 114 | 41 |
| 1.9 | 151 | 67 | 79 | 107 | 27 |
| 2 | 160 | 66 | 84 | 102 | 31 |
| 2.1 | 152 | 56 | 96 | 114 | 33 |
| 2.2 | 161 | 61 | 85 | 105 | 31 |
| 2.3 | 162 | 83 | 86 | 110 | 36 |
| 2.4 | 152 | 102 | 82 | 102 | 34 |
| 2.5 | 165 | 95 | 97 | 109 | 42 |
| 2.6 | 156 | 94 | 80 | 88 | 37 |
| 2.7 | 158 | 92 | 82 | 92 | 32 |
| 2.8 | 158 | 103 | 90 | 113 | 40 |
| 2.9 | 146 | 98 | 88 | 100 | 27 |
| 3 | 157 | 84 | 74 | 121 | 31 |
| 3.1 | 151 | 78 | 91 | 121 | 45 |
| 3.2 | 155 | 96 | 87 | 111 | 31 |
| 3.3 | 149 | 90 | 98 | 136 | 31 |
| 3.4 | 167 | 75 | 89 | 102 | 35 |
| 3.5 | 153 | 71 | 75 | 115 | 39 |
| 3.6 | 147 | 82 | 79 | 102 | 36 |
| 3.7 | 155 | 77 | 82 | 103 | 41 |
| 3.8 | 162 | 63 | 88 | 121 | 41 |
| 3.9 | 163 | 77 | 71 | 114 | 30 |
| 4 | 136 | 67 | 87 | 90 | 34 |
| 4.1 | 147 | 80 | 97 | 100 | 47 |
| 4.2 | 159 | 72 | 83 | 105 | 41 |
| 4.3 | 163 | 83 | 88 | 116 | 39 |
| 4.4 | 153 | 85 | 84 | 109 | 48 |
| 4.5 | 163 | 82 | 83 | 107 | 42 |
| 4.6 | 162 | 74 | 84 | 112 | 45 |
| 4.7 | 159 | 66 | 89 | 128 | 50 |
| 4.8 | 156 | 96 | 115 | 134 | 56 |
| 4.9 | 144 | 67 | 118 | 123 | 61 |
| 5 | 149 | 80 | 112 | 107 | 53 |
| 5.1 | 164 | 120 | 87 | 96 | 51 |
| 5.2 | 138 | 99 | 92 | 97 | 56 |
| 5.3 | 147 | 115 | 77 | 111 | 50 |
| 5.4 | 137 | 99 | 81 | 109 | 39 |
| 5.5 | 142 | 94 | 89 | 114 | 39 |
| 5.6 | 144 | 112 | 90 | 116 | 39 |
| 5.7 | 143 | 118 | 74 | 112 | 35 |
| 5.8 | 160 | 114 | 82 | 100 | 42 |
| 5.9 | 152 | 115 | 95 | 111 | 35 |
| 6 | 137 | 110 | 91 | 115 | 51 |
| 6.1 | 134 | 106 | 91 | 126 | 52 |
| 6.2 | 148 | 112 | 112 | 119 | 47 |
| 6.3 | 145 | 118 | 117 | 125 | 45 |
| 6.4 | 150 | 105 | 123 | 140 | 57 |
| 6.5 | 145 | 95 | 111 | 124 | 52 |
| 6.6 | 137 | 99 | 127 | 112 | 47 |
| 6.7 | 147 | 111 | 109 | 109 | 44 |
| 6.8 | 151 | 111 | 105 | 123 | 57 |
| 6.9 | 156 | 100 | 101 | 101 | 45 |
| 7 | 157 | 106 | 126 | 112 | 45 |
| 7.1 | 149 | 103 | 114 | 120 | 50 |
| 7.2 | 152 | 102 | 109 | 87 | 49 |
| 7.3 | 155 | 125 | 96 | 111 | 40 |
| 7.4 | 149 | 105 | 98 | 114 | 39 |
| 7.5 | 143 | 110 | 107 | 114 | 37 |
| 7.6 | 146 | 115 | 105 | 118 | 53 |
| 7.7 | 141 | 101 | 103 | 100 | 53 |
| 7.8 | 137 | 103 | 110 | 109 | 44 |
| 7.9 | 143 | 104 | 105 | 101 | 46 |
| 8 | 148 | 107 | 109 | 114 | 49 |
| 8.1 | 143 | 109 | 107 | 107 | 49 |
| 8.2 | 155 | 105 | 86 | 125 | 53 |
| 8.3 | 132 | 95 | 98 | 124 | 48 |
| 8.4 | 141 | 102 | 97 | 124 | 43 |
| 8.5 | 159 | 108 | 107 | 135 | 54 |
| 8.6 | 148 | 97 | 116 | 127 | 47 |
| 8.7 | 150 | 90 | 113 | 133 | 56 |
| 8.8 | 153 | 114 | 108 | 139 | 34 |
| 8.9 | 164 | 107 | 114 | 98 | 57 |
| 9 | 161 | 99 | 103 | 125 | 54 |
| 9.1 | 149 | 99 | 111 | 122 | 49 |
| 9.2 | 151 | 102 | 123 | 123 | 50 |
| 9.3 | 147 | 105 | 104 | 124 | 44 |
| 9.4 | 149 | 96 | 106 | 107 | 50 |
| 9.5 | 139 | 101 | 95 | 118 | 51 |
| 9.6 | 144 | 108 | 101 | 108 | 40 |
| 9.7 | 151 | 94 | 109 | 120 | 49 |
| 9.8 | 152 | 105 | 97 | 108 | 50 |
| 9.9 | 129 | 108 | 106 | 111 | 33 |
| 10 | 153 | 100 | 123 | 112 | 44 |
| 10.1 | 147 | 108 | 109 | 115 | 41 |
| 10.2 | 152 | 102 | 111 | 111 | 37 |
| 10.3 | 138 | 102 | 119 | 110 | 36 |
| 10.4 | 158 | 98 | 111 | 139 | 52 |
| 10.5 | 153 | 102 | 129 | 116 | 40 |
| 10.6 | 143 | 99 | 115 | 108 | 45 |
| 10.7 | 145 | 106 | 108 | 128 | 48 |
| 10.8 | 160 | 108 | 110 | 125 | 52 |
| 10.9 | 147 | 98 | 114 | 118 | 57 |
| 11 | 160 | 116 | 112 | 121 | 59 |
| 11.1 | 158 | 97 | 122 | 118 | 68 |
| 11.2 | 147 | 100 | 112 | 115 | 78 |
| 11.3 | 150 | 100 | 118 | 119 | 85 |
| 11.4 | 146 | 97 | 111 | 106 | 74 |
| 11.5 | 167 | 91 | 103 | 109 | 78 |
| 11.6 | 165 | 93 | 95 | 117 | 74 |
| 11.7 | 156 | 100 | 109 | 121 | 82 |
| 11.8 | 146 | 107 | 101 | 126 | 73 |
| 11.9 | 144 | 109 | 113 | 119 | 85 |
| 12 | 141 | 112 | 116 | 109 | 70 |
| 12.1 | 151 | 99 | 122 | 114 | 69 |
| 12.2 | 151 | 106 | 107 | 109 | 76 |
| 12.3 | 145 | 111 | 103 | 107 | 78 |
| 12.4 | 143 | 112 | 112 | 119 | 71 |
| 12.5 | 152 | 114 | 121 | 114 | 89 |
| 12.6 | 140 | 98 | 104 | 116 | 81 |
| 12.7 | 138 | 95 | 106 | 119 | 84 |
| 12.8 | 146 | 114 | 106 | 115 | 82 |
| 12.9 | 144 | 107 | 108 | 116 | 81 |
| 13 | 150 | 105 | 115 | 113 | 75 |
| 13.1 | 134 | 100 | 109 | 129 | 68 |
| 13.2 | 148 | 110 | 110 | 122 | 66 |
| 13.3 | 150 | 118 | 124 | 120 | 71 |
| 13.4 | 136 | 106 | 106 | 126 | 76 |
| 13.5 | 149 | 107 | 105 | 123 | 66 |
| 13.6 | 146 | 105 | 113 | 123 | 60 |
| 13.7 | 162 | 116 | 112 | 125 | 65 |
| 13.8 | 142 | 114 | 110 | 119 | 70 |
| 13.9 | 153 | 105 | 89 | 117 | 79 |
| 14 | 159 | 112 | 117 | 105 | 78 |
| 14.1 | 144 | 133 | 97 | 115 | 71 |
| 14.2 | 142 | 110 | 115 | 133 | 69 |
| 14.3 | 148 | 101 | 122 | 127 | 74 |
| 14.4 | 152 | 107 | 106 | 123 | 58 |
| 14.5 | 151 | 109 | 109 | 136 | 54 |
| 14.6 | 141 | 110 | 110 | 134 | 68 |
| 14.7 | 140 | 113 | 104 | 139 | 67 |
| 14.8 | 142 | 111 | 110 | 129 | 69 |
| 14.9 | 148 | 113 | 111 | 132 | 64 |
| 15 | 144 | 107 | 96 | 131 | 67 |
| 15.1 | 154 | 106 | 107 | 146 | 77 |
| 15.2 | 150 | 121 | 104 | 137 | 78 |
| 15.3 | 153 | 124 | 104 | 130 | 74 |
| 15.4 | 137 | 104 | 109 | 119 | 91 |
| 15.5 | 131 | 109 | 92 | 128 | 76 |
| 15.6 | 147 | 112 | 90 | 135 | 82 |
| 15.7 | 153 | 120 | 131 | 143 | 81 |
| 15.8 | 154 | 101 | 118 | 137 | 70 |
| 15.9 | 159 | 125 | 114 | 150 | 92 |
| 16 | 151 | 122 | 117 | 153 | 111 |
| 16.1 | 145 | 118 | 118 | 141 | 112 |
| 16.2 | 141 | 117 | 117 | 159 | 95 |
| 16.3 | 160 | 107 | 115 | 170 | 86 |
| 16.4 | 136 | 105 | 101 | 143 | 86 |
| 16.5 | 142 | 105 | 119 | 156 | 84 |
| 16.6 | 141 | 97 | 113 | 164 | 107 |
| 16.7 | 140 | 118 | 105 | 142 | 87 |
| 16.8 | 146 | 93 | 105 | 117 | 94 |
| 16.9 | 133 | 104 | 124 | 129 | 100 |
| 17 | 148 | 96 | 113 | 171 | 102 |
| 17.1 | 145 | 108 | 84 | 162 | 94 |
| 17.2 | 159 | 95 | 101 | 171 | 114 |
| 17.3 | 160 | 104 | 100 | 144 | 105 |
| 17.4 | 141 | 100 | 97 | 151 | 108 |
| 17.5 | 137 | 95 | 104 | 164 | 92 |
| 17.6 | 145 | 110 | 106 | 158 | 71 |
| 17.7 | 155 | 104 | 110 | 157 | 102 |
| 17.8 | 140 | 92 | 106 | 160 | 108 |
| 17.9 | 160 | 105 | 107 | 171 | 102 |
| 18 | 127 | 97 | 125 | 178 | 102 |
| 18.1 | 161 | 101 | 108 | 172 | 119 |
| 18.2 | 155 | 97 | 106 | 141 | 109 |
| 18.3 | 143 | 113 | 107 | 133 | 93 |
| 18.4 | 158 | 108 | 103 | 154 | 101 |
| 18.5 | 123 | 97 | 121 | 141 | 91 |
| 18.6 | 152 | 119 | 121 | 152 | 87 |
| 18.7 | 149 | 111 | 99 | 152 | 82 |
| 18.8 | 135 | 108 | 108 | 141 | 89 |
| 18.9 | 149 | 113 | 115 | 137 | 95 |
| 19 | 134 | 100 | 108 | 147 | 88 |
| 19.1 | 160 | 114 | 112 | 154 | 99 |
| 19.2 | 135 | 100 | 113 | 144 | 69 |
| 19.3 | 142 | 93 | 102 | 132 | 91 |
| 19.4 | 159 | 104 | 104 | 145 | 71 |
| 19.5 | 148 | 118 | 112 | 128 | 78 |
| 19.6 | 161 | 112 | 106 | 126 | 65 |
| 19.7 | 138 | 97 | 129 | 144 | 85 |
| 19.8 | 155 | 95 | 120 | 141 | 71 |
| 19.9 | 147 | 100 | 138 | 148 | 81 |

Histones - CLOCK_PAS-B (CLOCKS38/42)

Number of contacts

| Frame | 1st | 2nd | 3rd | 4th | 5th |
| --- | --- | --- | --- | --- | --- |
| 0 | 0 | 0 | 4 | 0 | 49 |
| 0.1 | 0 | 0 | 4 | 0 | 50 |
| 0.2 | 20 | 31 | 23 | 0 | 80 |
| 0.3 | 66 | 82 | 55 | 1 | 89 |
| 0.4 | 75 | 87 | 47 | 3 | 104 |
| 0.5 | 66 | 78 | 48 | 95 | 106 |
| 0.6 | 71 | 76 | 62 | 71 | 86 |
| 0.7 | 62 | 79 | 55 | 59 | 85 |
| 0.8 | 69 | 77 | 55 | 81 | 91 |
| 0.9 | 79 | 68 | 57 | 58 | 75 |
| 1 | 66 | 81 | 68 | 67 | 89 |
| 1.1 | 71 | 84 | 39 | 69 | 75 |
| 1.2 | 78 | 75 | 55 | 58 | 91 |
| 1.3 | 91 | 81 | 58 | 85 | 93 |
| 1.4 | 69 | 90 | 72 | 64 | 90 |
| 1.5 | 108 | 88 | 70 | 65 | 123 |
| 1.6 | 111 | 86 | 60 | 68 | 112 |
| 1.7 | 138 | 92 | 68 | 83 | 116 |
| 1.8 | 112 | 91 | 66 | 74 | 107 |
| 1.9 | 106 | 90 | 66 | 78 | 111 |
| 2 | 98 | 85 | 66 | 86 | 113 |
| 2.1 | 85 | 79 | 59 | 87 | 111 |
| 2.2 | 81 | 83 | 75 | 86 | 114 |
| 2.3 | 97 | 93 | 56 | 80 | 118 |
| 2.4 | 79 | 86 | 60 | 75 | 108 |
| 2.5 | 72 | 80 | 61 | 70 | 119 |
| 2.6 | 87 | 72 | 66 | 68 | 107 |
| 2.7 | 81 | 85 | 66 | 100 | 118 |
| 2.8 | 91 | 81 | 65 | 94 | 115 |
| 2.9 | 92 | 78 | 64 | 101 | 108 |
| 3 | 85 | 77 | 66 | 114 | 106 |
| 3.1 | 80 | 79 | 67 | 97 | 121 |
| 3.2 | 89 | 80 | 65 | 99 | 109 |
| 3.3 | 91 | 95 | 64 | 103 | 100 |
| 3.4 | 82 | 82 | 67 | 110 | 133 |
| 3.5 | 82 | 94 | 61 | 118 | 122 |
| 3.6 | 84 | 89 | 64 | 116 | 119 |
| 3.7 | 80 | 81 | 62 | 116 | 105 |
| 3.8 | 92 | 89 | 52 | 111 | 127 |
| 3.9 | 67 | 99 | 67 | 118 | 121 |
| 4 | 56 | 85 | 73 | 112 | 138 |
| 4.1 | 63 | 81 | 67 | 109 | 140 |
| 4.2 | 70 | 82 | 64 | 117 | 120 |
| 4.3 | 72 | 81 | 73 | 115 | 128 |
| 4.4 | 88 | 85 | 72 | 98 | 126 |
| 4.5 | 87 | 87 | 59 | 96 | 116 |
| 4.6 | 94 | 86 | 72 | 83 | 124 |
| 4.7 | 64 | 80 | 78 | 86 | 113 |
| 4.8 | 63 | 76 | 76 | 105 | 111 |
| 4.9 | 73 | 75 | 83 | 101 | 125 |
| 5 | 73 | 80 | 81 | 124 | 102 |
| 5.1 | 89 | 91 | 74 | 110 | 113 |
| 5.2 | 79 | 81 | 76 | 107 | 112 |
| 5.3 | 88 | 81 | 86 | 108 | 123 |
| 5.4 | 98 | 80 | 68 | 103 | 105 |
| 5.5 | 64 | 66 | 71 | 114 | 110 |
| 5.6 | 67 | 86 | 77 | 113 | 120 |
| 5.7 | 62 | 78 | 77 | 120 | 102 |
| 5.8 | 103 | 81 | 69 | 108 | 111 |
| 5.9 | 84 | 82 | 71 | 105 | 107 |
| 6 | 84 | 100 | 71 | 110 | 122 |
| 6.1 | 81 | 86 | 74 | 108 | 121 |
| 6.2 | 90 | 88 | 66 | 118 | 143 |
| 6.3 | 94 | 81 | 71 | 130 | 123 |
| 6.4 | 74 | 75 | 70 | 110 | 134 |
| 6.5 | 79 | 92 | 67 | 100 | 132 |
| 6.6 | 86 | 77 | 76 | 101 | 107 |
| 6.7 | 94 | 82 | 77 | 109 | 107 |
| 6.8 | 71 | 85 | 70 | 114 | 116 |
| 6.9 | 86 | 79 | 68 | 104 | 105 |
| 7 | 79 | 86 | 71 | 100 | 105 |
| 7.1 | 105 | 90 | 75 | 113 | 105 |
| 7.2 | 94 | 80 | 79 | 107 | 108 |
| 7.3 | 97 | 92 | 65 | 110 | 107 |
| 7.4 | 94 | 79 | 63 | 125 | 114 |
| 7.5 | 86 | 94 | 69 | 104 | 109 |
| 7.6 | 104 | 83 | 63 | 106 | 104 |
| 7.7 | 94 | 89 | 61 | 104 | 116 |
| 7.8 | 80 | 91 | 64 | 99 | 114 |
| 7.9 | 96 | 84 | 59 | 92 | 107 |
| 8 | 78 | 93 | 68 | 104 | 107 |
| 8.1 | 87 | 81 | 59 | 97 | 106 |
| 8.2 | 83 | 75 | 59 | 114 | 123 |
| 8.3 | 77 | 101 | 63 | 117 | 110 |
| 8.4 | 72 | 80 | 58 | 113 | 109 |
| 8.5 | 74 | 74 | 72 | 108 | 111 |
| 8.6 | 82 | 84 | 64 | 112 | 120 |
| 8.7 | 79 | 93 | 67 | 103 | 104 |
| 8.8 | 73 | 79 | 95 | 97 | 105 |
| 8.9 | 84 | 96 | 69 | 112 | 122 |
| 9 | 91 | 85 | 82 | 98 | 110 |
| 9.1 | 83 | 95 | 81 | 122 | 110 |
| 9.2 | 83 | 103 | 66 | 109 | 131 |
| 9.3 | 81 | 86 | 64 | 118 | 108 |
| 9.4 | 93 | 99 | 71 | 111 | 110 |
| 9.5 | 85 | 83 | 69 | 109 | 125 |
| 9.6 | 79 | 89 | 75 | 119 | 106 |
| 9.7 | 93 | 95 | 83 | 115 | 104 |
| 9.8 | 96 | 102 | 63 | 117 | 127 |
| 9.9 | 93 | 83 | 68 | 122 | 143 |
| 10 | 89 | 80 | 63 | 121 | 140 |
| 10.1 | 90 | 89 | 66 | 128 | 124 |
| 10.2 | 82 | 93 | 77 | 133 | 111 |
| 10.3 | 97 | 93 | 63 | 118 | 126 |
| 10.4 | 102 | 78 | 62 | 126 | 148 |
| 10.5 | 78 | 77 | 60 | 130 | 139 |
| 10.6 | 87 | 92 | 69 | 147 | 127 |
| 10.7 | 89 | 79 | 76 | 158 | 144 |
| 10.8 | 104 | 99 | 69 | 136 | 152 |
| 10.9 | 86 | 85 | 77 | 138 | 132 |
| 11 | 107 | 78 | 64 | 131 | 147 |
| 11.1 | 113 | 85 | 67 | 125 | 140 |
| 11.2 | 111 | 93 | 73 | 126 | 135 |
| 11.3 | 97 | 87 | 68 | 113 | 143 |
| 11.4 | 99 | 83 | 71 | 122 | 141 |
| 11.5 | 87 | 95 | 65 | 115 | 144 |
| 11.6 | 104 | 88 | 56 | 126 | 146 |
| 11.7 | 101 | 74 | 64 | 129 | 147 |
| 11.8 | 88 | 75 | 66 | 123 | 156 |
| 11.9 | 98 | 80 | 65 | 120 | 134 |
| 12 | 98 | 59 | 75 | 130 | 162 |
| 12.1 | 108 | 85 | 60 | 105 | 140 |
| 12.2 | 68 | 73 | 78 | 125 | 141 |
| 12.3 | 63 | 88 | 66 | 130 | 145 |
| 12.4 | 88 | 84 | 61 | 127 | 134 |
| 12.5 | 79 | 58 | 71 | 143 | 154 |
| 12.6 | 85 | 78 | 82 | 133 | 135 |
| 12.7 | 74 | 74 | 71 | 131 | 116 |
| 12.8 | 81 | 86 | 77 | 123 | 117 |
| 12.9 | 75 | 71 | 81 | 131 | 106 |
| 13 | 75 | 89 | 75 | 133 | 111 |
| 13.1 | 84 | 87 | 68 | 127 | 120 |
| 13.2 | 85 | 85 | 65 | 125 | 109 |
| 13.3 | 90 | 80 | 61 | 130 | 115 |
| 13.4 | 89 | 87 | 68 | 126 | 106 |
| 13.5 | 91 | 70 | 70 | 136 | 113 |
| 13.6 | 81 | 82 | 75 | 140 | 111 |
| 13.7 | 88 | 85 | 67 | 125 | 123 |
| 13.8 | 79 | 85 | 50 | 117 | 131 |
| 13.9 | 83 | 81 | 60 | 146 | 129 |
| 14 | 101 | 65 | 84 | 142 | 135 |
| 14.1 | 77 | 81 | 66 | 137 | 121 |
| 14.2 | 83 | 80 | 58 | 128 | 132 |
| 14.3 | 89 | 88 | 73 | 142 | 118 |
| 14.4 | 98 | 83 | 62 | 139 | 133 |
| 14.5 | 86 | 73 | 68 | 143 | 123 |
| 14.6 | 96 | 74 | 80 | 128 | 142 |
| 14.7 | 85 | 82 | 77 | 148 | 145 |
| 14.8 | 76 | 72 | 68 | 127 | 137 |
| 14.9 | 95 | 85 | 72 | 148 | 130 |
| 15 | 83 | 79 | 65 | 135 | 120 |
| 15.1 | 90 | 84 | 72 | 138 | 125 |
| 15.2 | 85 | 79 | 72 | 141 | 111 |
| 15.3 | 87 | 77 | 75 | 137 | 107 |
| 15.4 | 81 | 76 | 75 | 141 | 121 |
| 15.5 | 79 | 78 | 71 | 134 | 117 |
| 15.6 | 78 | 85 | 73 | 130 | 129 |
| 15.7 | 106 | 87 | 66 | 135 | 131 |
| 15.8 | 83 | 72 | 65 | 116 | 131 |
| 15.9 | 88 | 83 | 72 | 134 | 139 |
| 16 | 96 | 79 | 63 | 135 | 146 |
| 16.1 | 90 | 61 | 64 | 135 | 133 |
| 16.2 | 88 | 73 | 56 | 118 | 107 |
| 16.3 | 86 | 82 | 70 | 130 | 114 |
| 16.4 | 96 | 84 | 72 | 130 | 135 |
| 16.5 | 80 | 91 | 59 | 132 | 127 |
| 16.6 | 85 | 83 | 80 | 144 | 145 |
| 16.7 | 85 | 78 | 65 | 143 | 130 |
| 16.8 | 63 | 85 | 56 | 124 | 118 |
| 16.9 | 59 | 80 | 69 | 138 | 134 |
| 17 | 67 | 79 | 63 | 130 | 146 |
| 17.1 | 76 | 69 | 69 | 133 | 136 |
| 17.2 | 69 | 67 | 69 | 134 | 147 |
| 17.3 | 75 | 75 | 72 | 144 | 146 |
| 17.4 | 70 | 75 | 71 | 147 | 136 |
| 17.5 | 85 | 82 | 68 | 135 | 136 |
| 17.6 | 85 | 77 | 69 | 140 | 127 |
| 17.7 | 64 | 85 | 74 | 163 | 113 |
| 17.8 | 79 | 71 | 62 | 134 | 121 |
| 17.9 | 101 | 92 | 61 | 146 | 151 |
| 18 | 90 | 75 | 73 | 143 | 112 |
| 18.1 | 82 | 80 | 72 | 125 | 131 |
| 18.2 | 78 | 88 | 60 | 129 | 143 |
| 18.3 | 80 | 86 | 67 | 136 | 134 |
| 18.4 | 82 | 83 | 67 | 127 | 141 |
| 18.5 | 83 | 81 | 78 | 141 | 154 |
| 18.6 | 74 | 76 | 74 | 125 | 131 |
| 18.7 | 55 | 90 | 61 | 146 | 163 |
| 18.8 | 44 | 93 | 72 | 138 | 174 |
| 18.9 | 65 | 89 | 79 | 152 | 139 |
| 19 | 55 | 82 | 75 | 150 | 118 |
| 19.1 | 63 | 79 | 60 | 141 | 131 |
| 19.2 | 60 | 94 | 66 | 155 | 137 |
| 19.3 | 51 | 86 | 70 | 140 | 121 |
| 19.4 | 58 | 78 | 75 | 157 | 154 |
| 19.5 | 66 | 73 | 62 | 141 | 160 |
| 19.6 | 67 | 81 | 64 | 141 | 133 |
| 19.7 | 73 | 94 | 73 | 140 | 125 |
| 19.8 | 65 | 82 | 85 | 158 | 141 |
| 19.9 | 67 | 77 | 61 | 146 | 141 |

Histones - CLOCK_PAS-B (CLOCKS38/42/BMAL1S78)

Number of contacts

| Frame | 1st | 2nd | 3rd | 4th | 5th |
| --- | --- | --- | --- | --- | --- |
| 0 | 3 | 0 | 0 | 15 | 11 |
| 0.1 | 3 | 0 | 0 | 16 | 11 |
| 0.2 | 22 | 0 | 0 | 81 | 72 |
| 0.3 | 64 | 0 | 10 | 100 | 88 |
| 0.4 | 66 | 0 | 21 | 84 | 92 |
| 0.5 | 96 | 0 | 55 | 99 | 96 |
| 0.6 | 105 | 0 | 99 | 102 | 66 |
| 0.7 | 113 | 2 | 91 | 118 | 58 |
| 0.8 | 111 | 0 | 96 | 106 | 67 |
| 0.9 | 99 | 0 | 98 | 111 | 65 |
| 1 | 113 | 0 | 127 | 114 | 91 |
| 1.1 | 95 | 0 | 124 | 97 | 77 |
| 1.2 | 109 | 0 | 117 | 115 | 77 |
| 1.3 | 107 | 0 | 134 | 106 | 87 |
| 1.4 | 95 | 0 | 120 | 90 | 89 |
| 1.5 | 93 | 0 | 126 | 96 | 86 |
| 1.6 | 80 | 0 | 122 | 113 | 90 |
| 1.7 | 86 | 0 | 120 | 123 | 69 |
| 1.8 | 89 | 0 | 128 | 132 | 79 |
| 1.9 | 96 | 0 | 122 | 114 | 80 |
| 2 | 98 | 0 | 109 | 122 | 79 |
| 2.1 | 105 | 0 | 125 | 115 | 76 |
| 2.2 | 94 | 0 | 123 | 105 | 86 |
| 2.3 | 103 | 0 | 126 | 113 | 82 |
| 2.4 | 91 | 0 | 124 | 92 | 71 |
| 2.5 | 107 | 0 | 128 | 110 | 75 |
| 2.6 | 90 | 0 | 103 | 110 | 79 |
| 2.7 | 93 | 22 | 132 | 117 | 66 |
| 2.8 | 107 | 32 | 139 | 101 | 76 |
| 2.9 | 96 | 62 | 114 | 102 | 73 |
| 3 | 106 | 53 | 126 | 118 | 79 |
| 3.1 | 107 | 59 | 108 | 134 | 60 |
| 3.2 | 100 | 62 | 124 | 128 | 63 |
| 3.3 | 102 | 59 | 124 | 136 | 80 |
| 3.4 | 100 | 63 | 111 | 125 | 71 |
| 3.5 | 112 | 61 | 108 | 126 | 71 |
| 3.6 | 121 | 59 | 119 | 111 | 53 |
| 3.7 | 109 | 65 | 115 | 132 | 62 |
| 3.8 | 122 | 58 | 110 | 124 | 65 |
| 3.9 | 114 | 61 | 118 | 109 | 67 |
| 4 | 102 | 66 | 116 | 114 | 73 |
| 4.1 | 95 | 57 | 126 | 117 | 81 |
| 4.2 | 98 | 77 | 113 | 110 | 70 |
| 4.3 | 97 | 59 | 120 | 112 | 68 |
| 4.4 | 107 | 59 | 105 | 105 | 96 |
| 4.5 | 99 | 62 | 109 | 105 | 95 |
| 4.6 | 104 | 67 | 96 | 123 | 101 |
| 4.7 | 94 | 64 | 99 | 105 | 106 |
| 4.8 | 107 | 67 | 93 | 105 | 93 |
| 4.9 | 109 | 49 | 111 | 127 | 76 |
| 5 | 110 | 77 | 120 | 127 | 66 |
| 5.1 | 113 | 69 | 122 | 106 | 87 |
| 5.2 | 125 | 57 | 130 | 105 | 85 |
| 5.3 | 110 | 70 | 118 | 112 | 97 |
| 5.4 | 112 | 75 | 126 | 118 | 98 |
| 5.5 | 95 | 74 | 120 | 127 | 88 |
| 5.6 | 98 | 79 | 126 | 107 | 87 |
| 5.7 | 87 | 79 | 124 | 113 | 85 |
| 5.8 | 94 | 67 | 128 | 117 | 83 |
| 5.9 | 114 | 76 | 126 | 145 | 81 |
| 6 | 97 | 89 | 124 | 109 | 84 |
| 6.1 | 96 | 91 | 119 | 119 | 95 |
| 6.2 | 110 | 67 | 132 | 120 | 98 |
| 6.3 | 103 | 73 | 129 | 110 | 110 |
| 6.4 | 107 | 87 | 142 | 116 | 88 |
| 6.5 | 94 | 78 | 128 | 111 | 90 |
| 6.6 | 102 | 69 | 126 | 122 | 105 |
| 6.7 | 89 | 83 | 147 | 133 | 87 |
| 6.8 | 90 | 81 | 144 | 137 | 96 |
| 6.9 | 81 | 82 | 127 | 136 | 97 |
| 7 | 94 | 83 | 125 | 147 | 85 |
| 7.1 | 98 | 88 | 132 | 123 | 91 |
| 7.2 | 106 | 71 | 127 | 141 | 87 |
| 7.3 | 116 | 82 | 110 | 142 | 83 |
| 7.4 | 101 | 80 | 125 | 144 | 95 |
| 7.5 | 87 | 69 | 135 | 145 | 92 |
| 7.6 | 109 | 82 | 132 | 118 | 100 |
| 7.7 | 101 | 85 | 131 | 126 | 116 |
| 7.8 | 119 | 84 | 124 | 135 | 92 |
| 7.9 | 104 | 82 | 149 | 125 | 94 |
| 8 | 92 | 77 | 145 | 139 | 88 |
| 8.1 | 119 | 79 | 148 | 123 | 86 |
| 8.2 | 112 | 67 | 149 | 133 | 90 |
| 8.3 | 117 | 90 | 141 | 123 | 79 |
| 8.4 | 120 | 97 | 160 | 127 | 79 |
| 8.5 | 119 | 78 | 142 | 134 | 74 |
| 8.6 | 96 | 79 | 140 | 124 | 73 |
| 8.7 | 98 | 75 | 150 | 115 | 85 |
| 8.8 | 111 | 87 | 125 | 140 | 70 |
| 8.9 | 96 | 94 | 134 | 137 | 81 |
| 9 | 122 | 101 | 126 | 130 | 86 |
| 9.1 | 114 | 79 | 151 | 137 | 85 |
| 9.2 | 107 | 93 | 148 | 136 | 94 |
| 9.3 | 122 | 90 | 137 | 128 | 86 |
| 9.4 | 119 | 85 | 135 | 140 | 78 |
| 9.5 | 109 | 90 | 163 | 138 | 96 |
| 9.6 | 118 | 90 | 130 | 135 | 107 |
| 9.7 | 106 | 92 | 136 | 136 | 95 |
| 9.8 | 120 | 97 | 132 | 127 | 80 |
| 9.9 | 117 | 104 | 158 | 136 | 94 |
| 10 | 119 | 127 | 125 | 145 | 80 |
| 10.1 | 106 | 119 | 137 | 146 | 91 |
| 10.2 | 111 | 117 | 140 | 119 | 98 |
| 10.3 | 113 | 119 | 150 | 131 | 97 |
| 10.4 | 122 | 115 | 133 | 128 | 84 |
| 10.5 | 115 | 112 | 117 | 132 | 65 |
| 10.6 | 115 | 116 | 143 | 138 | 89 |
| 10.7 | 127 | 122 | 136 | 132 | 92 |
| 10.8 | 101 | 100 | 133 | 144 | 97 |
| 10.9 | 121 | 110 | 135 | 141 | 91 |
| 11 | 108 | 106 | 135 | 132 | 98 |
| 11.1 | 115 | 131 | 122 | 140 | 85 |
| 11.2 | 119 | 119 | 119 | 154 | 86 |
| 11.3 | 125 | 115 | 146 | 145 | 95 |
| 11.4 | 104 | 113 | 164 | 136 | 80 |
| 11.5 | 110 | 108 | 158 | 137 | 96 |
| 11.6 | 114 | 106 | 150 | 139 | 94 |
| 11.7 | 105 | 109 | 164 | 138 | 93 |
| 11.8 | 115 | 111 | 145 | 145 | 102 |
| 11.9 | 117 | 114 | 152 | 146 | 89 |
| 12 | 112 | 117 | 141 | 142 | 97 |
| 12.1 | 122 | 121 | 143 | 134 | 79 |
| 12.2 | 106 | 108 | 156 | 151 | 101 |
| 12.3 | 118 | 105 | 152 | 145 | 77 |
| 12.4 | 131 | 127 | 149 | 138 | 81 |
| 12.5 | 105 | 134 | 162 | 132 | 97 |
| 12.6 | 101 | 124 | 152 | 147 | 81 |
| 12.7 | 100 | 117 | 136 | 157 | 90 |
| 12.8 | 103 | 112 | 144 | 138 | 82 |
| 12.9 | 103 | 125 | 139 | 136 | 89 |
| 13 | 103 | 121 | 137 | 144 | 91 |
| 13.1 | 115 | 133 | 136 | 139 | 98 |
| 13.2 | 116 | 132 | 133 | 154 | 100 |
| 13.3 | 106 | 127 | 133 | 142 | 78 |
| 13.4 | 112 | 128 | 147 | 138 | 87 |
| 13.5 | 125 | 120 | 115 | 126 | 91 |
| 13.6 | 112 | 120 | 129 | 132 | 95 |
| 13.7 | 115 | 133 | 135 | 140 | 83 |
| 13.8 | 115 | 119 | 151 | 150 | 88 |
| 13.9 | 109 | 127 | 131 | 128 | 99 |
| 14 | 111 | 116 | 133 | 133 | 103 |
| 14.1 | 128 | 120 | 136 | 143 | 102 |
| 14.2 | 109 | 123 | 126 | 152 | 99 |
| 14.3 | 112 | 137 | 144 | 141 | 94 |
| 14.4 | 112 | 126 | 125 | 147 | 104 |
| 14.5 | 132 | 134 | 146 | 150 | 89 |
| 14.6 | 112 | 118 | 127 | 137 | 77 |
| 14.7 | 109 | 130 | 114 | 135 | 87 |
| 14.8 | 117 | 122 | 138 | 154 | 68 |
| 14.9 | 120 | 141 | 135 | 129 | 97 |
| 15 | 103 | 119 | 131 | 143 | 71 |
| 15.1 | 92 | 91 | 145 | 143 | 93 |
| 15.2 | 120 | 121 | 130 | 143 | 71 |
| 15.3 | 121 | 112 | 128 | 145 | 83 |
| 15.4 | 91 | 109 | 136 | 135 | 94 |
| 15.5 | 113 | 108 | 134 | 150 | 86 |
| 15.6 | 104 | 134 | 131 | 148 | 89 |
| 15.7 | 117 | 109 | 125 | 132 | 79 |
| 15.8 | 103 | 121 | 129 | 139 | 84 |
| 15.9 | 109 | 120 | 127 | 151 | 104 |
| 16 | 113 | 104 | 134 | 146 | 71 |
| 16.1 | 107 | 119 | 117 | 135 | 73 |
| 16.2 | 111 | 132 | 109 | 147 | 99 |
| 16.3 | 105 | 126 | 147 | 144 | 84 |
| 16.4 | 105 | 125 | 132 | 136 | 77 |
| 16.5 | 108 | 117 | 122 | 149 | 83 |
| 16.6 | 98 | 113 | 120 | 141 | 79 |
| 16.7 | 100 | 119 | 134 | 146 | 76 |
| 16.8 | 94 | 125 | 118 | 146 | 89 |
| 16.9 | 98 | 108 | 118 | 143 | 87 |
| 17 | 112 | 112 | 132 | 139 | 89 |
| 17.1 | 102 | 117 | 144 | 137 | 89 |
| 17.2 | 112 | 137 | 145 | 155 | 71 |
| 17.3 | 112 | 128 | 146 | 144 | 88 |
| 17.4 | 111 | 121 | 147 | 136 | 81 |
| 17.5 | 111 | 121 | 135 | 143 | 102 |
| 17.6 | 89 | 109 | 128 | 141 | 77 |
| 17.7 | 115 | 118 | 138 | 134 | 80 |
| 17.8 | 124 | 125 | 122 | 137 | 100 |
| 17.9 | 116 | 120 | 136 | 136 | 77 |
| 18 | 130 | 119 | 138 | 154 | 90 |
| 18.1 | 125 | 134 | 141 | 161 | 81 |
| 18.2 | 118 | 108 | 146 | 145 | 84 |
| 18.3 | 124 | 116 | 133 | 135 | 90 |
| 18.4 | 115 | 131 | 136 | 135 | 89 |
| 18.5 | 123 | 130 | 128 | 150 | 86 |
| 18.6 | 130 | 122 | 138 | 154 | 97 |
| 18.7 | 105 | 122 | 135 | 155 | 95 |
| 18.8 | 131 | 121 | 139 | 146 | 89 |
| 18.9 | 120 | 134 | 143 | 145 | 96 |
| 19 | 115 | 124 | 138 | 155 | 80 |
| 19.1 | 106 | 108 | 136 | 141 | 84 |
| 19.2 | 120 | 126 | 144 | 133 | 82 |
| 19.3 | 104 | 117 | 125 | 130 | 98 |
| 19.4 | 111 | 116 | 134 | 137 | 88 |
| 19.5 | 122 | 134 | 141 | 127 | 79 |
| 19.6 | 119 | 117 | 138 | 159 | 99 |
| 19.7 | 136 | 113 | 135 | 144 | 84 |
| 19.8 | 117 | 132 | 145 | 156 | 85 |
| 19.9 | 109 | 125 | 138 | 139 | 77 |

**Figure S6.** Timeline plots pertaining to number of contacts between histones and CLOCK PASB domain for the systems studied.

Histones - BMAL1_PAS-A (nonphosphorylated)

Number of contacts

| Frame | 1st | 2nd | 3rd | 4th | 5th |
| --- | --- | --- | --- | --- | --- |
| 0 | 8 | 0 | 4 | 7 | 30 |
| 0.1 | 8 | 0 | 4 | 7 | 2 |
| 0.2 | 8 | 0 | 0 | 4 | 0 |
| 0.3 | 11 | 0 | 0 | 0 | 15 |
| 0.4 | 26 | 11 | 0 | 10 | 1 |
| 0.5 | 28 | 17 | 1 | 0 | 14 |
| 0.6 | 22 | 20 | 0 | 0 | 15 |
| 0.7 | 25 | 20 | 0 | 0 | 27 |
| 0.8 | 22 | 14 | 14 | 4 | 27 |
| 0.9 | 33 | 12 | 9 | 4 | 9 |
| 1 | 34 | 24 | 8 | 17 | 31 |
| 1.1 | 28 | 18 | 24 | 16 | 20 |
| 1.2 | 30 | 6 | 24 | 4 | 25 |
| 1.3 | 46 | 18 | 21 | 4 | 13 |
| 1.4 | 36 | 16 | 25 | 1 | 15 |
| 1.5 | 42 | 13 | 19 | 1 | 28 |
| 1.6 | 40 | 12 | 16 | 11 | 16 |
| 1.7 | 38 | 1 | 3 | 8 | 30 |
| 1.8 | 36 | 16 | 20 | 4 | 29 |
| 1.9 | 35 | 4 | 2 | 6 | 15 |
| 2 | 45 | 4 | 14 | 2 | 0 |
| 2.1 | 36 | 2 | 16 | 1 | 0 |
| 2.2 | 40 | 2 | 15 | 8 | 0 |
| 2.3 | 35 | 4 | 23 | 8 | 26 |
| 2.4 | 41 | 2 | 19 | 21 | 18 |
| 2.5 | 40 | 7 | 18 | 10 | 23 |
| 2.6 | 36 | 13 | 21 | 11 | 19 |
| 2.7 | 36 | 8 | 27 | 8 | 4 |
| 2.8 | 47 | 11 | 19 | 19 | 7 |
| 2.9 | 33 | 19 | 19 | 18 | 23 |
| 3 | 32 | 23 | 18 | 23 | 21 |
| 3.1 | 28 | 16 | 17 | 25 | 23 |
| 3.2 | 33 | 21 | 20 | 17 | 24 |
| 3.3 | 48 | 17 | 15 | 10 | 16 |
| 3.4 | 50 | 15 | 13 | 12 | 23 |
| 3.5 | 33 | 18 | 9 | 15 | 17 |
| 3.6 | 35 | 20 | 2 | 16 | 16 |
| 3.7 | 28 | 27 | 0 | 13 | 23 |
| 3.8 | 31 | 14 | 4 | 19 | 23 |
| 3.9 | 32 | 2 | 15 | 17 | 22 |
| 4 | 31 | 2 | 13 | 22 | 22 |
| 4.1 | 33 | 0 | 8 | 24 | 18 |
| 4.2 | 28 | 24 | 9 | 18 | 5 |
| 4.3 | 28 | 14 | 14 | 19 | 13 |
| 4.4 | 33 | 13 | 10 | 21 | 23 |
| 4.5 | 25 | 12 | 7 | 20 | 26 |
| 4.6 | 28 | 11 | 14 | 29 | 16 |
| 4.7 | 36 | 13 | 12 | 8 | 28 |
| 4.8 | 50 | 12 | 13 | 12 | 26 |
| 4.9 | 39 | 14 | 10 | 23 | 26 |
| 5 | 36 | 15 | 9 | 27 | 27 |
| 5.1 | 34 | 18 | 11 | 24 | 25 |
| 5.2 | 47 | 23 | 9 | 17 | 21 |
| 5.3 | 58 | 9 | 10 | 18 | 28 |
| 5.4 | 52 | 19 | 7 | 10 | 25 |
| 5.5 | 52 | 17 | 13 | 11 | 29 |
| 5.6 | 64 | 20 | 10 | 19 | 21 |
| 5.7 | 51 | 22 | 13 | 18 | 16 |
| 5.8 | 48 | 18 | 11 | 6 | 24 |
| 5.9 | 51 | 19 | 8 | 7 | 26 |
| 6 | 46 | 17 | 11 | 9 | 26 |
| 6.1 | 51 | 19 | 13 | 24 | 19 |
| 6.2 | 59 | 25 | 16 | 16 | 16 |
| 6.3 | 31 | 23 | 14 | 20 | 15 |
| 6.4 | 49 | 23 | 19 | 19 | 10 |
| 6.5 | 39 | 23 | 11 | 24 | 15 |
| 6.6 | 24 | 25 | 12 | 12 | 21 |
| 6.7 | 16 | 25 | 9 | 14 | 13 |
| 6.8 | 32 | 27 | 22 | 11 | 26 |
| 6.9 | 25 | 24 | 14 | 14 | 14 |
| 7 | 42 | 18 | 18 | 28 | 19 |
| 7.1 | 41 | 22 | 12 | 17 | 21 |
| 7.2 | 49 | 18 | 14 | 13 | 20 |
| 7.3 | 44 | 26 | 24 | 25 | 17 |
| 7.4 | 45 | 20 | 19 | 18 | 22 |
| 7.5 | 50 | 26 | 13 | 23 | 15 |
| 7.6 | 60 | 26 | 19 | 23 | 20 |
| 7.7 | 60 | 22 | 17 | 21 | 17 |
| 7.8 | 55 | 28 | 16 | 24 | 24 |
| 7.9 | 54 | 23 | 14 | 34 | 21 |
| 8 | 56 | 27 | 14 | 29 | 27 |
| 8.1 | 23 | 19 | 12 | 26 | 25 |
| 8.2 | 39 | 18 | 13 | 24 | 29 |
| 8.3 | 34 | 28 | 22 | 18 | 27 |
| 8.4 | 23 | 27 | 17 | 27 | 25 |
| 8.5 | 27 | 31 | 18 | 17 | 28 |
| 8.6 | 24 | 26 | 19 | 1 | 14 |
| 8.7 | 20 | 31 | 19 | 0 | 23 |
| 8.8 | 49 | 22 | 16 | 20 | 31 |
| 8.9 | 22 | 25 | 15 | 19 | 18 |
| 9 | 29 | 24 | 14 | 15 | 17 |
| 9.1 | 50 | 18 | 20 | 21 | 21 |
| 9.2 | 41 | 19 | 20 | 27 | 13 |
| 9.3 | 36 | 15 | 22 | 19 | 12 |
| 9.4 | 20 | 21 | 25 | 22 | 20 |
| 9.5 | 40 | 24 | 18 | 21 | 29 |
| 9.6 | 32 | 22 | 18 | 15 | 22 |
| 9.7 | 37 | 24 | 22 | 19 | 18 |
| 9.8 | 40 | 18 | 22 | 21 | 27 |
| 9.9 | 52 | 17 | 19 | 22 | 27 |
| 10 | 37 | 16 | 20 | 23 | 33 |
| 10.1 | 34 | 17 | 25 | 24 | 23 |
| 10.2 | 49 | 21 | 20 | 23 | 27 |
| 10.3 | 57 | 20 | 21 | 27 | 40 |
| 10.4 | 43 | 0 | 22 | 22 | 50 |
| 10.5 | 28 | 19 | 20 | 23 | 33 |
| 10.6 | 36 | 24 | 19 | 18 | 35 |
| 10.7 | 42 | 23 | 16 | 23 | 41 |
| 10.8 | 37 | 22 | 21 | 24 | 42 |
| 10.9 | 33 | 18 | 21 | 23 | 32 |
| 11 | 14 | 19 | 21 | 12 | 38 |
| 11.1 | 16 | 18 | 20 | 20 | 36 |
| 11.2 | 21 | 23 | 17 | 24 | 32 |
| 11.3 | 14 | 20 | 20 | 22 | 33 |
| 11.4 | 19 | 21 | 22 | 27 | 36 |
| 11.5 | 20 | 25 | 19 | 21 | 32 |
| 11.6 | 15 | 18 | 21 | 21 | 36 |
| 11.7 | 26 | 19 | 24 | 18 | 40 |
| 11.8 | 25 | 17 | 17 | 31 | 42 |
| 11.9 | 20 | 21 | 15 | 17 | 14 |
| 12 | 25 | 22 | 19 | 21 | 20 |
| 12.1 | 21 | 22 | 18 | 31 | 13 |
| 12.2 | 25 | 20 | 20 | 22 | 26 |
| 12.3 | 23 | 21 | 26 | 21 | 33 |
| 12.4 | 24 | 21 | 22 | 20 | 25 |
| 12.5 | 27 | 18 | 20 | 32 | 24 |
| 12.6 | 26 | 21 | 21 | 29 | 29 |
| 12.7 | 26 | 15 | 26 | 26 | 31 |
| 12.8 | 25 | 16 | 24 | 24 | 37 |
| 12.9 | 24 | 18 | 20 | 25 | 29 |
| 13 | 32 | 22 | 21 | 32 | 31 |
| 13.1 | 27 | 21 | 21 | 31 | 35 |
| 13.2 | 21 | 23 | 18 | 33 | 32 |
| 13.3 | 16 | 33 | 21 | 30 | 35 |
| 13.4 | 20 | 32 | 21 | 25 | 35 |
| 13.5 | 18 | 34 | 17 | 24 | 32 |
| 13.6 | 19 | 37 | 23 | 27 | 24 |
| 13.7 | 17 | 37 | 20 | 24 | 29 |
| 13.8 | 21 | 40 | 19 | 15 | 27 |
| 13.9 | 20 | 35 | 21 | 17 | 25 |
| 14 | 19 | 19 | 16 | 20 | 25 |
| 14.1 | 16 | 28 | 18 | 12 | 28 |
| 14.2 | 22 | 29 | 24 | 20 | 28 |
| 14.3 | 18 | 34 | 21 | 15 | 33 |
| 14.4 | 19 | 29 | 22 | 25 | 28 |
| 14.5 | 11 | 36 | 19 | 19 | 29 |
| 14.6 | 15 | 35 | 24 | 22 | 23 |
| 14.7 | 22 | 31 | 14 | 22 | 28 |
| 14.8 | 19 | 29 | 19 | 23 | 33 |
| 14.9 | 21 | 39 | 17 | 21 | 28 |
| 15 | 18 | 31 | 15 | 16 | 25 |
| 15.1 | 20 | 38 | 18 | 8 | 37 |
| 15.2 | 22 | 41 | 24 | 9 | 22 |
| 15.3 | 24 | 34 | 19 | 13 | 26 |
| 15.4 | 17 | 33 | 22 | 11 | 31 |
| 15.5 | 19 | 32 | 21 | 18 | 22 |
| 15.6 | 21 | 36 | 20 | 20 | 25 |
| 15.7 | 21 | 36 | 20 | 19 | 23 |
| 15.8 | 24 | 37 | 21 | 23 | 21 |
| 15.9 | 17 | 31 | 18 | 4 | 25 |
| 16 | 21 | 38 | 18 | 32 | 16 |
| 16.1 | 19 | 37 | 17 | 31 | 27 |
| 16.2 | 19 | 35 | 19 | 23 | 24 |
| 16.3 | 19 | 36 | 20 | 32 | 26 |
| 16.4 | 19 | 40 | 16 | 25 | 37 |
| 16.5 | 22 | 33 | 19 | 22 | 28 |
| 16.6 | 21 | 26 | 20 | 16 | 25 |
| 16.7 | 21 | 30 | 17 | 18 | 23 |
| 16.8 | 16 | 29 | 17 | 22 | 29 |
| 16.9 | 17 | 31 | 17 | 10 | 22 |
| 17 | 23 | 36 | 19 | 21 | 22 |
| 17.1 | 20 | 35 | 19 | 23 | 16 |
| 17.2 | 21 | 32 | 19 | 20 | 6 |
| 17.3 | 20 | 34 | 23 | 25 | 5 |
| 17.4 | 19 | 36 | 19 | 24 | 17 |
| 17.5 | 17 | 32 | 23 | 23 | 16 |
| 17.6 | 17 | 37 | 20 | 9 | 16 |
| 17.7 | 16 | 34 | 21 | 18 | 18 |
| 17.8 | 17 | 30 | 21 | 18 | 14 |
| 17.9 | 21 | 28 | 19 | 12 | 19 |
| 18 | 19 | 30 | 19 | 14 | 15 |
| 18.1 | 19 | 31 | 19 | 16 | 13 |
| 18.2 | 21 | 32 | 21 | 24 | 15 |
| 18.3 | 17 | 33 | 14 | 5 | 15 |
| 18.4 | 18 | 35 | 20 | 23 | 14 |
| 18.5 | 19 | 29 | 18 | 20 | 15 |
| 18.6 | 21 | 39 | 21 | 24 | 23 |
| 18.7 | 19 | 32 | 14 | 13 | 20 |
| 18.8 | 18 | 35 | 17 | 14 | 13 |
| 18.9 | 19 | 36 | 16 | 13 | 16 |
| 19 | 18 | 33 | 16 | 5 | 18 |
| 19.1 | 20 | 33 | 19 | 11 | 12 |
| 19.2 | 18 | 35 | 19 | 3 | 18 |
| 19.3 | 15 | 37 | 17 | 11 | 14 |
| 19.4 | 21 | 31 | 18 | 10 | 13 |
| 19.5 | 19 | 31 | 20 | 15 | 35 |
| 19.6 | 18 | 30 | 18 | 9 | 29 |
| 19.7 | 17 | 31 | 13 | 12 | 33 |
| 19.8 | 17 | 32 | 20 | 5 | 16 |
| 19.9 | 20 | 27 | 15 | 4 | 17 |

Histones - BMAL1_PAS-A (BMAL1S78)

Number of contacts

| Frame | 1st | 2nd | 3rd | 4th | 5th |
| --- | --- | --- | --- | --- | --- |
| 0 | 0 | 5 | 9 | 32 | 22 |
| 0.1 | 0 | 5 | 9 | 0 | 21 |
| 0.2 | 0 | 22 | 7 | 0 | 35 |
| 0.3 | 0 | 11 | 17 | 0 | 42 |
| 0.4 | 9 | 12 | 14 | 5 | 37 |
| 0.5 | 13 | 0 | 18 | 17 | 39 |
| 0.6 | 17 | 19 | 16 | 16 | 26 |
| 0.7 | 26 | 21 | 6 | 17 | 22 |
| 0.8 | 33 | 9 | 14 | 17 | 19 |
| 0.9 | 15 | 27 | 14 | 24 | 17 |
| 1 | 29 | 27 | 17 | 34 | 28 |
| 1.1 | 25 | 22 | 1 | 16 | 20 |
| 1.2 | 26 | 23 | 15 | 32 | 25 |
| 1.3 | 25 | 20 | 24 | 32 | 21 |
| 1.4 | 27 | 29 | 6 | 26 | 25 |
| 1.5 | 20 | 29 | 7 | 29 | 25 |
| 1.6 | 22 | 24 | 4 | 18 | 28 |
| 1.7 | 25 | 26 | 6 | 27 | 13 |
| 1.8 | 35 | 24 | 4 | 36 | 2 |
| 1.9 | 27 | 29 | 1 | 33 | 15 |
| 2 | 18 | 27 | 7 | 31 | 20 |
| 2.1 | 21 | 26 | 12 | 30 | 13 |
| 2.2 | 44 | 25 | 15 | 30 | 23 |
| 2.3 | 28 | 26 | 10 | 30 | 1 |
| 2.4 | 25 | 22 | 10 | 32 | 15 |
| 2.5 | 17 | 20 | 4 | 27 | 17 |
| 2.6 | 18 | 21 | 14 | 32 | 4 |
| 2.7 | 13 | 21 | 17 | 32 | 16 |
| 2.8 | 12 | 17 | 23 | 31 | 18 |
| 2.9 | 14 | 25 | 20 | 25 | 5 |
| 3 | 16 | 21 | 12 | 44 | 13 |
| 3.1 | 17 | 29 | 13 | 48 | 17 |
| 3.2 | 13 | 22 | 14 | 36 | 16 |
| 3.3 | 19 | 24 | 19 | 34 | 18 |
| 3.4 | 23 | 26 | 15 | 42 | 12 |
| 3.5 | 26 | 27 | 17 | 33 | 16 |
| 3.6 | 58 | 28 | 21 | 30 | 16 |
| 3.7 | 38 | 19 | 16 | 26 | 13 |
| 3.8 | 48 | 18 | 11 | 25 | 22 |
| 3.9 | 43 | 27 | 13 | 30 | 16 |
| 4 | 36 | 24 | 20 | 20 | 22 |
| 4.1 | 44 | 20 | 17 | 23 | 6 |
| 4.2 | 58 | 17 | 16 | 29 | 17 |
| 4.3 | 62 | 26 | 13 | 23 | 18 |
| 4.4 | 58 | 24 | 14 | 19 | 25 |
| 4.5 | 59 | 18 | 1 | 26 | 16 |
| 4.6 | 51 | 18 | 6 | 17 | 17 |
| 4.7 | 47 | 25 | 5 | 18 | 17 |
| 4.8 | 62 | 20 | 7 | 12 | 15 |
| 4.9 | 56 | 27 | 2 | 17 | 17 |
| 5 | 57 | 28 | 1 | 16 | 6 |
| 5.1 | 64 | 27 | 0 | 22 | 12 |
| 5.2 | 54 | 17 | 2 | 17 | 8 |
| 5.3 | 59 | 11 | 0 | 17 | 12 |
| 5.4 | 56 | 16 | 2 | 17 | 14 |
| 5.5 | 66 | 18 | 5 | 23 | 20 |
| 5.6 | 57 | 26 | 1 | 32 | 24 |
| 5.7 | 68 | 20 | 1 | 34 | 21 |
| 5.8 | 64 | 14 | 2 | 26 | 27 |
| 5.9 | 56 | 31 | 10 | 35 | 27 |
| 6 | 65 | 35 | 3 | 32 | 23 |
| 6.1 | 68 | 21 | 8 | 28 | 16 |
| 6.2 | 55 | 22 | 0 | 23 | 13 |
| 6.3 | 64 | 26 | 0 | 27 | 0 |
| 6.4 | 66 | 19 | 0 | 21 | 8 |
| 6.5 | 61 | 19 | 2 | 23 | 22 |
| 6.6 | 55 | 28 | 1 | 32 | 22 |
| 6.7 | 69 | 15 | 0 | 33 | 19 |
| 6.8 | 62 | 31 | 0 | 21 | 18 |
| 6.9 | 64 | 26 | 0 | 33 | 23 |
| 7 | 70 | 23 | 0 | 31 | 30 |
| 7.1 | 63 | 44 | 2 | 29 | 24 |
| 7.2 | 55 | 24 | 0 | 19 | 19 |
| 7.3 | 46 | 25 | 0 | 21 | 19 |
| 7.4 | 61 | 15 | 0 | 25 | 32 |
| 7.5 | 63 | 25 | 0 | 21 | 25 |
| 7.6 | 69 | 32 | 0 | 22 | 9 |
| 7.7 | 68 | 14 | 0 | 28 | 11 |
| 7.8 | 54 | 32 | 0 | 27 | 10 |
| 7.9 | 61 | 32 | 0 | 34 | 25 |
| 8 | 70 | 33 | 0 | 23 | 24 |
| 8.1 | 58 | 28 | 0 | 26 | 21 |
| 8.2 | 68 | 34 | 0 | 21 | 21 |
| 8.3 | 70 | 23 | 2 | 36 | 31 |
| 8.4 | 54 | 22 | 2 | 18 | 24 |
| 8.5 | 61 | 25 | 0 | 28 | 20 |
| 8.6 | 59 | 27 | 2 | 28 | 27 |
| 8.7 | 64 | 21 | 0 | 29 | 19 |
| 8.8 | 66 | 24 | 0 | 31 | 14 |
| 8.9 | 57 | 33 | 0 | 24 | 21 |
| 9 | 58 | 25 | 0 | 27 | 22 |
| 9.1 | 57 | 26 | 11 | 41 | 18 |
| 9.2 | 66 | 27 | 0 | 39 | 22 |
| 9.3 | 72 | 26 | 2 | 40 | 27 |
| 9.4 | 63 | 30 | 0 | 33 | 24 |
| 9.5 | 58 | 31 | 3 | 25 | 23 |
| 9.6 | 64 | 25 | 2 | 36 | 30 |
| 9.7 | 60 | 23 | 0 | 35 | 20 |
| 9.8 | 62 | 27 | 0 | 37 | 10 |
| 9.9 | 65 | 18 | 0 | 26 | 22 |
| 10 | 69 | 19 | 0 | 22 | 27 |
| 10.1 | 66 | 25 | 1 | 27 | 15 |
| 10.2 | 68 | 18 | 0 | 32 | 22 |
| 10.3 | 61 | 26 | 0 | 36 | 11 |
| 10.4 | 67 | 26 | 0 | 35 | 20 |
| 10.5 | 57 | 27 | 0 | 34 | 20 |
| 10.6 | 65 | 7 | 0 | 33 | 25 |
| 10.7 | 72 | 8 | 0 | 37 | 12 |
| 10.8 | 60 | 14 | 0 | 30 | 28 |
| 10.9 | 55 | 13 | 0 | 34 | 23 |
| 11 | 60 | 11 | 2 | 33 | 23 |
| 11.1 | 62 | 16 | 2 | 38 | 18 |
| 11.2 | 65 | 13 | 14 | 41 | 21 |
| 11.3 | 65 | 13 | 0 | 26 | 26 |
| 11.4 | 61 | 12 | 5 | 44 | 25 |
| 11.5 | 64 | 25 | 3 | 39 | 29 |
| 11.6 | 73 | 28 | 6 | 42 | 24 |
| 11.7 | 66 | 28 | 6 | 26 | 28 |
| 11.8 | 68 | 36 | 0 | 34 | 22 |
| 11.9 | 57 | 31 | 0 | 32 | 22 |
| 12 | 67 | 24 | 0 | 29 | 23 |
| 12.1 | 57 | 27 | 0 | 28 | 28 |
| 12.2 | 67 | 34 | 0 | 33 | 16 |
| 12.3 | 57 | 28 | 0 | 28 | 25 |
| 12.4 | 60 | 28 | 0 | 27 | 22 |
| 12.5 | 61 | 19 | 0 | 33 | 18 |
| 12.6 | 67 | 29 | 0 | 37 | 23 |
| 12.7 | 65 | 29 | 0 | 34 | 24 |
| 12.8 | 64 | 31 | 0 | 40 | 20 |
| 12.9 | 65 | 18 | 1 | 31 | 8 |
| 13 | 78 | 22 | 0 | 42 | 12 |
| 13.1 | 71 | 25 | 0 | 35 | 17 |
| 13.2 | 60 | 27 | 2 | 35 | 22 |
| 13.3 | 65 | 23 | 0 | 44 | 18 |
| 13.4 | 58 | 27 | 0 | 41 | 21 |
| 13.5 | 60 | 30 | 0 | 32 | 18 |
| 13.6 | 62 | 21 | 0 | 25 | 14 |
| 13.7 | 75 | 22 | 0 | 32 | 8 |
| 13.8 | 70 | 5 | 0 | 41 | 16 |
| 13.9 | 57 | 24 | 0 | 19 | 21 |
| 14 | 66 | 21 | 0 | 37 | 18 |
| 14.1 | 56 | 22 | 0 | 26 | 3 |
| 14.2 | 54 | 22 | 0 | 25 | 24 |
| 14.3 | 70 | 33 | 0 | 32 | 16 |
| 14.4 | 68 | 24 | 0 | 20 | 17 |
| 14.5 | 64 | 26 | 0 | 33 | 18 |
| 14.6 | 66 | 25 | 0 | 59 | 18 |
| 14.7 | 64 | 25 | 0 | 59 | 15 |
| 14.8 | 72 | 29 | 0 | 49 | 26 |
| 14.9 | 64 | 29 | 0 | 52 | 23 |
| 15 | 67 | 35 | 0 | 71 | 22 |
| 15.1 | 67 | 21 | 0 | 58 | 17 |
| 15.2 | 52 | 32 | 0 | 73 | 21 |
| 15.3 | 62 | 20 | 0 | 65 | 23 |
| 15.4 | 71 | 26 | 0 | 63 | 24 |
| 15.5 | 69 | 27 | 0 | 63 | 20 |
| 15.6 | 61 | 8 | 0 | 73 | 21 |
| 15.7 | 57 | 24 | 0 | 66 | 17 |
| 15.8 | 77 | 21 | 0 | 53 | 17 |
| 15.9 | 54 | 24 | 0 | 57 | 23 |
| 16 | 63 | 22 | 0 | 60 | 23 |
| 16.1 | 59 | 26 | 1 | 66 | 20 |
| 16.2 | 62 | 22 | 0 | 54 | 23 |
| 16.3 | 62 | 17 | 1 | 62 | 22 |
| 16.4 | 67 | 15 | 0 | 51 | 30 |
| 16.5 | 64 | 24 | 2 | 51 | 26 |
| 16.6 | 47 | 32 | 0 | 53 | 25 |
| 16.7 | 60 | 27 | 0 | 54 | 15 |
| 16.8 | 78 | 21 | 9 | 66 | 23 |
| 16.9 | 66 | 21 | 16 | 55 | 9 |
| 17 | 62 | 29 | 23 | 59 | 23 |
| 17.1 | 60 | 17 | 13 | 48 | 24 |
| 17.2 | 68 | 16 | 21 | 54 | 22 |
| 17.3 | 65 | 23 | 20 | 46 | 27 |
| 17.4 | 72 | 15 | 13 | 38 | 22 |
| 17.5 | 72 | 27 | 23 | 39 | 24 |
| 17.6 | 70 | 30 | 14 | 46 | 18 |
| 17.7 | 66 | 19 | 20 | 47 | 13 |
| 17.8 | 80 | 15 | 21 | 53 | 3 |
| 17.9 | 66 | 13 | 16 | 52 | 2 |
| 18 | 61 | 25 | 22 | 52 | 3 |
| 18.1 | 68 | 19 | 20 | 59 | 21 |
| 18.2 | 55 | 29 | 4 | 72 | 20 |
| 18.3 | 60 | 19 | 10 | 68 | 30 |
| 18.4 | 65 | 18 | 7 | 77 | 21 |
| 18.5 | 55 | 19 | 13 | 71 | 19 |
| 18.6 | 71 | 7 | 21 | 61 | 4 |
| 18.7 | 54 | 15 | 10 | 69 | 14 |
| 18.8 | 68 | 30 | 10 | 53 | 17 |
| 18.9 | 62 | 24 | 20 | 62 | 18 |
| 19 | 59 | 33 | 24 | 62 | 5 |
| 19.1 | 60 | 18 | 22 | 59 | 3 |
| 19.2 | 61 | 17 | 17 | 54 | 3 |
| 19.3 | 53 | 18 | 22 | 60 | 2 |
| 19.4 | 64 | 16 | 13 | 53 | 5 |
| 19.5 | 67 | 23 | 14 | 55 | 18 |
| 19.6 | 62 | 21 | 2 | 59 | 9 |
| 19.7 | 68 | 17 | 2 | 54 | 16 |
| 19.8 | 61 | 25 | 0 | 50 | 22 |
| 19.9 | 66 | 24 | 0 | 57 | 23 |

Histones - BMAL1_PAS-A (CLOCKS38/42)

Number of contacts

| Frame | 1st | 2nd | 3rd | 4th | 5th |
| --- | --- | --- | --- | --- | --- |
| 0 | 6 | 6 | 17 | 12 | 0 |
| 0.1 | 6 | 7 | 17 | 12 | 0 |
| 0.2 | 3 | 18 | 2 | 22 | 31 |
| 0.3 | 0 | 23 | 12 | 19 | 37 |
| 0.4 | 6 | 14 | 19 | 23 | 46 |
| 0.5 | 8 | 13 | 12 | 34 | 47 |
| 0.6 | 13 | 20 | 27 | 28 | 44 |
| 0.7 | 2 | 22 | 22 | 29 | 29 |
| 0.8 | 0 | 25 | 21 | 7 | 56 |
| 0.9 | 1 | 23 | 21 | 15 | 37 |
| 1 | 0 | 19 | 18 | 13 | 34 |
| 1.1 | 1 | 15 | 29 | 6 | 39 |
| 1.2 | 0 | 23 | 31 | 7 | 39 |
| 1.3 | 0 | 12 | 25 | 12 | 47 |
| 1.4 | 2 | 21 | 30 | 11 | 46 |
| 1.5 | 1 | 12 | 27 | 5 | 39 |
| 1.6 | 1 | 24 | 31 | 16 | 33 |
| 1.7 | 1 | 17 | 24 | 13 | 53 |
| 1.8 | 0 | 23 | 31 | 10 | 35 |
| 1.9 | 0 | 26 | 32 | 9 | 31 |
| 2 | 0 | 23 | 29 | 15 | 27 |
| 2.1 | 0 | 18 | 30 | 16 | 27 |
| 2.2 | 0 | 19 | 26 | 17 | 31 |
| 2.3 | 0 | 22 | 29 | 20 | 33 |
| 2.4 | 0 | 22 | 26 | 17 | 18 |
| 2.5 | 0 | 25 | 30 | 13 | 14 |
| 2.6 | 0 | 14 | 27 | 14 | 20 |
| 2.7 | 0 | 21 | 26 | 12 | 10 |
| 2.8 | 0 | 18 | 34 | 12 | 20 |
| 2.9 | 0 | 23 | 28 | 15 | 22 |
| 3 | 0 | 20 | 28 | 14 | 27 |
| 3.1 | 0 | 12 | 28 | 12 | 28 |
| 3.2 | 0 | 11 | 29 | 14 | 35 |
| 3.3 | 0 | 23 | 38 | 18 | 18 |
| 3.4 | 0 | 22 | 35 | 14 | 10 |
| 3.5 | 0 | 21 | 31 | 14 | 14 |
| 3.6 | 0 | 25 | 23 | 16 | 11 |
| 3.7 | 0 | 28 | 23 | 14 | 24 |
| 3.8 | 0 | 18 | 32 | 12 | 22 |
| 3.9 | 0 | 23 | 38 | 16 | 21 |
| 4 | 0 | 17 | 27 | 17 | 13 |
| 4.1 | 0 | 19 | 33 | 16 | 16 |
| 4.2 | 0 | 22 | 29 | 17 | 14 |
| 4.3 | 0 | 28 | 22 | 17 | 23 |
| 4.4 | 0 | 17 | 21 | 17 | 15 |
| 4.5 | 0 | 7 | 29 | 13 | 5 |
| 4.6 | 0 | 30 | 27 | 21 | 15 |
| 4.7 | 0 | 24 | 19 | 31 | 31 |
| 4.8 | 0 | 23 | 27 | 14 | 39 |
| 4.9 | 0 | 25 | 28 | 10 | 24 |
| 5 | 0 | 22 | 29 | 17 | 38 |
| 5.1 | 0 | 28 | 29 | 19 | 40 |
| 5.2 | 0 | 20 | 30 | 20 | 34 |
| 5.3 | 0 | 20 | 23 | 26 | 41 |
| 5.4 | 0 | 18 | 14 | 17 | 41 |
| 5.5 | 0 | 27 | 20 | 16 | 40 |
| 5.6 | 0 | 14 | 39 | 17 | 28 |
| 5.7 | 0 | 30 | 39 | 20 | 43 |
| 5.8 | 0 | 50 | 35 | 21 | 49 |
| 5.9 | 0 | 42 | 17 | 18 | 56 |
| 6 | 0 | 48 | 30 | 11 | 28 |
| 6.1 | 0 | 45 | 26 | 22 | 35 |
| 6.2 | 0 | 53 | 25 | 22 | 51 |
| 6.3 | 0 | 43 | 27 | 24 | 22 |
| 6.4 | 0 | 43 | 38 | 24 | 29 |
| 6.5 | 0 | 51 | 16 | 13 | 30 |
| 6.6 | 0 | 30 | 28 | 13 | 34 |
| 6.7 | 0 | 42 | 26 | 24 | 52 |
| 6.8 | 0 | 43 | 38 | 17 | 27 |
| 6.9 | 0 | 41 | 27 | 26 | 21 |
| 7 | 0 | 47 | 29 | 11 | 10 |
| 7.1 | 0 | 48 | 27 | 16 | 27 |
| 7.2 | 0 | 46 | 24 | 21 | 19 |
| 7.3 | 0 | 44 | 29 | 15 | 22 |
| 7.4 | 0 | 45 | 15 | 12 | 10 |
| 7.5 | 0 | 48 | 26 | 22 | 37 |
| 7.6 | 0 | 45 | 30 | 17 | 27 |
| 7.7 | 0 | 50 | 25 | 24 | 26 |
| 7.8 | 0 | 44 | 31 | 20 | 31 |
| 7.9 | 0 | 45 | 28 | 19 | 13 |
| 8 | 0 | 40 | 27 | 19 | 17 |
| 8.1 | 0 | 42 | 34 | 21 | 38 |
| 8.2 | 0 | 31 | 31 | 15 | 15 |
| 8.3 | 0 | 37 | 25 | 21 | 21 |
| 8.4 | 0 | 41 | 27 | 21 | 17 |
| 8.5 | 0 | 41 | 32 | 20 | 33 |
| 8.6 | 0 | 53 | 27 | 17 | 20 |
| 8.7 | 0 | 50 | 28 | 25 | 26 |
| 8.8 | 0 | 71 | 25 | 21 | 44 |
| 8.9 | 0 | 64 | 24 | 20 | 32 |
| 9 | 0 | 65 | 27 | 23 | 20 |
| 9.1 | 0 | 57 | 31 | 22 | 23 |
| 9.2 | 0 | 64 | 29 | 21 | 18 |
| 9.3 | 0 | 61 | 27 | 16 | 35 |
| 9.4 | 0 | 57 | 29 | 23 | 31 |
| 9.5 | 0 | 71 | 28 | 17 | 41 |
| 9.6 | 0 | 58 | 24 | 20 | 28 |
| 9.7 | 0 | 55 | 29 | 22 | 37 |
| 9.8 | 0 | 69 | 31 | 22 | 28 |
| 9.9 | 0 | 73 | 28 | 23 | 26 |
| 10 | 0 | 60 | 30 | 25 | 26 |
| 10.1 | 0 | 60 | 25 | 20 | 19 |
| 10.2 | 0 | 64 | 23 | 15 | 14 |
| 10.3 | 0 | 70 | 32 | 20 | 18 |
| 10.4 | 0 | 65 | 28 | 24 | 28 |
| 10.5 | 0 | 48 | 25 | 26 | 20 |
| 10.6 | 0 | 52 | 18 | 26 | 21 |
| 10.7 | 0 | 48 | 20 | 26 | 9 |
| 10.8 | 0 | 55 | 21 | 22 | 12 |
| 10.9 | 0 | 61 | 26 | 33 | 23 |
| 11 | 0 | 48 | 27 | 37 | 19 |
| 11.1 | 0 | 47 | 34 | 24 | 11 |
| 11.2 | 0 | 57 | 23 | 25 | 5 |
| 11.3 | 0 | 61 | 21 | 28 | 15 |
| 11.4 | 0 | 47 | 26 | 30 | 14 |
| 11.5 | 0 | 59 | 20 | 29 | 14 |
| 11.6 | 0 | 65 | 17 | 30 | 13 |
| 11.7 | 0 | 54 | 28 | 27 | 10 |
| 11.8 | 0 | 44 | 20 | 28 | 10 |
| 11.9 | 0 | 63 | 23 | 22 | 11 |
| 12 | 0 | 63 | 24 | 18 | 20 |
| 12.1 | 0 | 61 | 21 | 28 | 26 |
| 12.2 | 0 | 70 | 29 | 22 | 23 |
| 12.3 | 0 | 62 | 20 | 37 | 12 |
| 12.4 | 0 | 57 | 24 | 26 | 28 |
| 12.5 | 0 | 63 | 29 | 30 | 26 |
| 12.6 | 0 | 50 | 26 | 28 | 14 |
| 12.7 | 0 | 62 | 21 | 18 | 12 |
| 12.8 | 0 | 39 | 25 | 32 | 21 |
| 12.9 | 0 | 50 | 24 | 26 | 27 |
| 13 | 0 | 47 | 29 | 29 | 26 |
| 13.1 | 0 | 39 | 21 | 28 | 17 |
| 13.2 | 0 | 61 | 29 | 29 | 28 |
| 13.3 | 0 | 60 | 29 | 24 | 32 |
| 13.4 | 0 | 59 | 29 | 41 | 34 |
| 13.5 | 0 | 68 | 25 | 23 | 53 |
| 13.6 | 0 | 64 | 28 | 22 | 44 |
| 13.7 | 0 | 50 | 25 | 25 | 23 |
| 13.8 | 0 | 57 | 25 | 31 | 22 |
| 13.9 | 0 | 65 | 26 | 25 | 30 |
| 14 | 0 | 49 | 27 | 33 | 41 |
| 14.1 | 0 | 62 | 26 | 37 | 64 |
| 14.2 | 0 | 65 | 23 | 34 | 54 |
| 14.3 | 0 | 75 | 21 | 37 | 87 |
| 14.4 | 0 | 68 | 25 | 36 | 37 |
| 14.5 | 0 | 70 | 17 | 34 | 36 |
| 14.6 | 0 | 57 | 28 | 34 | 21 |
| 14.7 | 0 | 52 | 22 | 35 | 22 |
| 14.8 | 0 | 54 | 27 | 28 | 21 |
| 14.9 | 0 | 67 | 25 | 27 | 24 |
| 15 | 0 | 58 | 18 | 33 | 17 |
| 15.1 | 0 | 61 | 25 | 28 | 26 |
| 15.2 | 0 | 52 | 21 | 24 | 41 |
| 15.3 | 0 | 65 | 23 | 33 | 42 |
| 15.4 | 0 | 65 | 27 | 26 | 32 |
| 15.5 | 0 | 69 | 27 | 35 | 28 |
| 15.6 | 0 | 64 | 24 | 28 | 30 |
| 15.7 | 0 | 68 | 22 | 24 | 29 |
| 15.8 | 0 | 66 | 22 | 25 | 13 |
| 15.9 | 0 | 73 | 25 | 32 | 19 |
| 16 | 0 | 65 | 21 | 29 | 29 |
| 16.1 | 0 | 60 | 26 | 26 | 15 |
| 16.2 | 0 | 84 | 21 | 23 | 25 |
| 16.3 | 0 | 72 | 24 | 31 | 22 |
| 16.4 | 0 | 58 | 24 | 24 | 12 |
| 16.5 | 0 | 54 | 20 | 27 | 12 |
| 16.6 | 0 | 54 | 17 | 35 | 16 |
| 16.7 | 0 | 62 | 17 | 26 | 8 |
| 16.8 | 0 | 63 | 19 | 18 | 20 |
| 16.9 | 0 | 56 | 18 | 30 | 11 |
| 17 | 0 | 71 | 26 | 22 | 10 |
| 17.1 | 0 | 69 | 18 | 25 | 7 |
| 17.2 | 0 | 62 | 17 | 21 | 11 |
| 17.3 | 0 | 59 | 28 | 27 | 18 |
| 17.4 | 0 | 60 | 30 | 32 | 10 |
| 17.5 | 0 | 54 | 26 | 25 | 13 |
| 17.6 | 0 | 53 | 15 | 40 | 11 |
| 17.7 | 0 | 70 | 20 | 31 | 12 |
| 17.8 | 0 | 62 | 20 | 36 | 7 |
| 17.9 | 0 | 61 | 23 | 35 | 18 |
| 18 | 0 | 73 | 19 | 29 | 10 |
| 18.1 | 0 | 66 | 25 | 29 | 9 |
| 18.2 | 0 | 71 | 27 | 34 | 21 |
| 18.3 | 0 | 62 | 25 | 39 | 23 |
| 18.4 | 0 | 55 | 29 | 21 | 46 |
| 18.5 | 0 | 76 | 30 | 31 | 41 |
| 18.6 | 0 | 59 | 30 | 27 | 33 |
| 18.7 | 0 | 68 | 31 | 30 | 27 |
| 18.8 | 0 | 67 | 26 | 30 | 25 |
| 18.9 | 0 | 72 | 26 | 27 | 10 |
| 19 | 0 | 70 | 29 | 31 | 11 |
| 19.1 | 0 | 50 | 23 | 31 | 13 |
| 19.2 | 0 | 57 | 28 | 27 | 22 |
| 19.3 | 0 | 55 | 28 | 46 | 13 |
| 19.4 | 0 | 55 | 29 | 25 | 10 |
| 19.5 | 0 | 61 | 28 | 37 | 22 |
| 19.6 | 0 | 56 | 25 | 33 | 11 |
| 19.7 | 0 | 54 | 31 | 27 | 12 |
| 19.8 | 0 | 44 | 26 | 34 | 16 |
| 19.9 | 0 | 46 | 28 | 38 | 13 |

Histones - BMAL1_PAS-A (CLOCKS38/42/BMAL1S78)

Number of contacts

| Frame | 1st | 2nd | 3rd | 4th | 5th |
| --- | --- | --- | --- | --- | --- |
| 0 | 19 | 2 | 0 | 17 | 22 |
| 0.1 | 18 | 2 | 0 | 17 | 22 |
| 0.2 | 16 | 16 | 0 | 26 | 23 |
| 0.3 | 5 | 17 | 0 | 33 | 21 |
| 0.4 | 9 | 20 | 0 | 13 | 24 |
| 0.5 | 15 | 12 | 0 | 26 | 21 |
| 0.6 | 13 | 27 | 0 | 27 | 24 |
| 0.7 | 17 | 30 | 0 | 30 | 26 |
| 0.8 | 22 | 24 | 0 | 25 | 30 |
| 0.9 | 21 | 8 | 0 | 34 | 21 |
| 1 | 11 | 12 | 0 | 31 | 14 |
| 1.1 | 20 | 13 | 0 | 28 | 27 |
| 1.2 | 9 | 25 | 0 | 36 | 4 |
| 1.3 | 23 | 18 | 0 | 29 | 17 |
| 1.4 | 12 | 13 | 0 | 27 | 19 |
| 1.5 | 14 | 12 | 0 | 34 | 13 |
| 1.6 | 20 | 12 | 0 | 27 | 17 |
| 1.7 | 15 | 21 | 0 | 31 | 27 |
| 1.8 | 17 | 19 | 0 | 44 | 21 |
| 1.9 | 15 | 13 | 0 | 40 | 26 |
| 2 | 26 | 17 | 0 | 38 | 17 |
| 2.1 | 4 | 21 | 0 | 45 | 12 |
| 2.2 | 8 | 9 | 0 | 46 | 25 |
| 2.3 | 7 | 21 | 0 | 44 | 24 |
| 2.4 | 0 | 40 | 0 | 51 | 17 |
| 2.5 | 19 | 53 | 0 | 49 | 16 |
| 2.6 | 13 | 62 | 0 | 57 | 21 |
| 2.7 | 14 | 59 | 0 | 49 | 15 |
| 2.8 | 14 | 65 | 0 | 53 | 24 |
| 2.9 | 17 | 58 | 0 | 47 | 29 |
| 3 | 11 | 66 | 0 | 53 | 22 |
| 3.1 | 20 | 63 | 0 | 46 | 21 |
| 3.2 | 23 | 57 | 0 | 45 | 24 |
| 3.3 | 21 | 61 | 0 | 45 | 27 |
| 3.4 | 24 | 63 | 0 | 46 | 18 |
| 3.5 | 21 | 67 | 0 | 43 | 26 |
| 3.6 | 6 | 78 | 0 | 40 | 22 |
| 3.7 | 19 | 72 | 0 | 48 | 22 |
| 3.8 | 17 | 72 | 0 | 48 | 21 |
| 3.9 | 23 | 68 | 0 | 40 | 30 |
| 4 | 17 | 72 | 0 | 50 | 25 |
| 4.1 | 16 | 70 | 0 | 44 | 24 |
| 4.2 | 20 | 82 | 0 | 47 | 15 |
| 4.3 | 16 | 69 | 0 | 40 | 29 |
| 4.4 | 19 | 55 | 0 | 47 | 25 |
| 4.5 | 22 | 68 | 0 | 34 | 17 |
| 4.6 | 10 | 69 | 0 | 36 | 31 |
| 4.7 | 25 | 59 | 0 | 34 | 32 |
| 4.8 | 20 | 72 | 0 | 45 | 27 |
| 4.9 | 20 | 64 | 0 | 40 | 22 |
| 5 | 12 | 57 | 0 | 34 | 28 |
| 5.1 | 23 | 76 | 0 | 30 | 23 |
| 5.2 | 10 | 73 | 0 | 33 | 20 |
| 5.3 | 6 | 60 | 0 | 34 | 20 |
| 5.4 | 10 | 75 | 0 | 37 | 22 |
| 5.5 | 10 | 68 | 0 | 40 | 19 |
| 5.6 | 6 | 60 | 0 | 35 | 21 |
| 5.7 | 13 | 70 | 0 | 40 | 23 |
| 5.8 | 12 | 76 | 0 | 38 | 22 |
| 5.9 | 14 | 66 | 0 | 40 | 22 |
| 6 | 9 | 69 | 0 | 31 | 27 |
| 6.1 | 26 | 75 | 0 | 42 | 22 |
| 6.2 | 18 | 82 | 0 | 37 | 29 |
| 6.3 | 15 | 73 | 0 | 32 | 26 |
| 6.4 | 24 | 81 | 0 | 34 | 26 |
| 6.5 | 16 | 59 | 0 | 33 | 20 |
| 6.6 | 28 | 74 | 0 | 39 | 23 |
| 6.7 | 10 | 79 | 0 | 38 | 29 |
| 6.8 | 22 | 74 | 0 | 37 | 24 |
| 6.9 | 12 | 70 | 0 | 32 | 20 |
| 7 | 23 | 81 | 0 | 33 | 17 |
| 7.1 | 21 | 57 | 0 | 34 | 17 |
| 7.2 | 19 | 80 | 0 | 31 | 22 |
| 7.3 | 11 | 76 | 0 | 40 | 11 |
| 7.4 | 14 | 65 | 0 | 38 | 17 |
| 7.5 | 12 | 75 | 0 | 32 | 17 |
| 7.6 | 8 | 57 | 0 | 44 | 10 |
| 7.7 | 6 | 65 | 0 | 43 | 27 |
| 7.8 | 14 | 83 | 0 | 38 | 24 |
| 7.9 | 17 | 70 | 0 | 41 | 12 |
| 8 | 16 | 69 | 0 | 40 | 32 |
| 8.1 | 15 | 68 | 0 | 37 | 35 |
| 8.2 | 9 | 74 | 0 | 41 | 33 |
| 8.3 | 3 | 81 | 0 | 33 | 31 |
| 8.4 | 12 | 77 | 0 | 41 | 24 |
| 8.5 | 15 | 75 | 0 | 32 | 15 |
| 8.6 | 14 | 88 | 0 | 25 | 19 |
| 8.7 | 18 | 80 | 0 | 20 | 17 |
| 8.8 | 7 | 95 | 0 | 18 | 25 |
| 8.9 | 7 | 84 | 0 | 23 | 24 |
| 9 | 8 | 75 | 0 | 21 | 26 |
| 9.1 | 6 | 103 | 0 | 23 | 23 |
| 9.2 | 13 | 80 | 0 | 23 | 26 |
| 9.3 | 9 | 83 | 0 | 25 | 26 |
| 9.4 | 2 | 77 | 0 | 18 | 25 |
| 9.5 | 18 | 91 | 0 | 22 | 20 |
| 9.6 | 19 | 102 | 0 | 29 | 26 |
| 9.7 | 16 | 83 | 0 | 26 | 31 |
| 9.8 | 11 | 75 | 0 | 24 | 21 |
| 9.9 | 24 | 88 | 0 | 23 | 25 |
| 10 | 18 | 73 | 0 | 23 | 28 |
| 10.1 | 18 | 70 | 0 | 17 | 20 |
| 10.2 | 9 | 81 | 0 | 16 | 24 |
| 10.3 | 7 | 84 | 0 | 18 | 19 |
| 10.4 | 11 | 83 | 0 | 20 | 14 |
| 10.5 | 9 | 81 | 0 | 14 | 17 |
| 10.6 | 10 | 78 | 0 | 19 | 21 |
| 10.7 | 14 | 73 | 0 | 19 | 19 |
| 10.8 | 10 | 81 | 0 | 18 | 27 |
| 10.9 | 17 | 76 | 0 | 17 | 32 |
| 11 | 21 | 68 | 0 | 11 | 30 |
| 11.1 | 21 | 78 | 0 | 13 | 34 |
| 11.2 | 20 | 76 | 0 | 14 | 34 |
| 11.3 | 17 | 95 | 0 | 11 | 25 |
| 11.4 | 19 | 93 | 0 | 12 | 29 |
| 11.5 | 34 | 84 | 0 | 10 | 26 |
| 11.6 | 37 | 81 | 0 | 9 | 34 |
| 11.7 | 24 | 82 | 0 | 8 | 35 |
| 11.8 | 17 | 91 | 0 | 10 | 26 |
| 11.9 | 12 | 82 | 0 | 10 | 30 |
| 12 | 6 | 94 | 0 | 13 | 34 |
| 12.1 | 28 | 87 | 0 | 11 | 26 |
| 12.2 | 26 | 83 | 2 | 13 | 33 |
| 12.3 | 27 | 75 | 2 | 11 | 28 |
| 12.4 | 28 | 81 | 0 | 11 | 30 |
| 12.5 | 31 | 65 | 0 | 9 | 31 |
| 12.6 | 21 | 80 | 0 | 12 | 30 |
| 12.7 | 18 | 85 | 0 | 13 | 38 |
| 12.8 | 25 | 87 | 0 | 10 | 32 |
| 12.9 | 27 | 82 | 0 | 13 | 29 |
| 13 | 29 | 76 | 0 | 9 | 28 |
| 13.1 | 28 | 76 | 0 | 9 | 33 |
| 13.2 | 16 | 71 | 0 | 11 | 32 |
| 13.3 | 19 | 60 | 0 | 8 | 26 |
| 13.4 | 21 | 76 | 0 | 8 | 36 |
| 13.5 | 22 | 68 | 0 | 13 | 36 |
| 13.6 | 17 | 57 | 0 | 8 | 26 |
| 13.7 | 23 | 58 | 0 | 4 | 29 |
| 13.8 | 25 | 50 | 0 | 5 | 29 |
| 13.9 | 25 | 64 | 0 | 4 | 33 |
| 14 | 15 | 56 | 0 | 5 | 34 |
| 14.1 | 27 | 61 | 0 | 4 | 25 |
| 14.2 | 20 | 67 | 0 | 4 | 28 |
| 14.3 | 30 | 56 | 0 | 11 | 34 |
| 14.4 | 22 | 50 | 0 | 8 | 27 |
| 14.5 | 23 | 51 | 0 | 6 | 29 |
| 14.6 | 28 | 54 | 0 | 6 | 20 |
| 14.7 | 21 | 53 | 0 | 6 | 26 |
| 14.8 | 29 | 58 | 0 | 3 | 26 |
| 14.9 | 30 | 45 | 0 | 0 | 29 |
| 15 | 25 | 43 | 0 | 0 | 33 |
| 15.1 | 21 | 56 | 0 | 0 | 32 |
| 15.2 | 27 | 51 | 0 | 0 | 32 |
| 15.3 | 23 | 63 | 0 | 6 | 33 |
| 15.4 | 23 | 68 | 0 | 7 | 33 |
| 15.5 | 29 | 68 | 0 | 7 | 30 |
| 15.6 | 21 | 71 | 0 | 6 | 28 |
| 15.7 | 28 | 45 | 0 | 3 | 29 |
| 15.8 | 26 | 53 | 0 | 6 | 29 |
| 15.9 | 20 | 50 | 0 | 1 | 34 |
| 16 | 30 | 74 | 0 | 10 | 26 |
| 16.1 | 23 | 67 | 0 | 6 | 31 |
| 16.2 | 27 | 69 | 0 | 7 | 35 |
| 16.3 | 27 | 70 | 0 | 3 | 30 |
| 16.4 | 30 | 68 | 0 | 5 | 29 |
| 16.5 | 17 | 62 | 0 | 8 | 30 |
| 16.6 | 28 | 59 | 0 | 12 | 29 |
| 16.7 | 24 | 65 | 0 | 2 | 27 |
| 16.8 | 25 | 74 | 0 | 9 | 35 |
| 16.9 | 24 | 63 | 0 | 8 | 27 |
| 17 | 20 | 67 | 0 | 7 | 28 |
| 17.1 | 10 | 60 | 0 | 6 | 32 |
| 17.2 | 10 | 65 | 0 | 8 | 32 |
| 17.3 | 11 | 69 | 0 | 6 | 33 |
| 17.4 | 4 | 65 | 1 | 12 | 31 |
| 17.5 | 10 | 70 | 1 | 8 | 31 |
| 17.6 | 15 | 68 | 0 | 13 | 28 |
| 17.7 | 4 | 64 | 0 | 0 | 28 |
| 17.8 | 7 | 63 | 1 | 3 | 22 |
| 17.9 | 6 | 69 | 1 | 2 | 28 |
| 18 | 15 | 56 | 2 | 6 | 31 |
| 18.1 | 18 | 70 | 2 | 2 | 29 |
| 18.2 | 22 | 69 | 0 | 2 | 33 |
| 18.3 | 23 | 57 | 0 | 3 | 27 |
| 18.4 | 10 | 59 | 0 | 3 | 35 |
| 18.5 | 7 | 69 | 0 | 4 | 27 |
| 18.6 | 18 | 57 | 0 | 2 | 34 |
| 18.7 | 15 | 63 | 0 | 2 | 32 |
| 18.8 | 14 | 64 | 0 | 3 | 38 |
| 18.9 | 12 | 67 | 0 | 2 | 36 |
| 19 | 21 | 63 | 0 | 6 | 32 |
| 19.1 | 21 | 65 | 0 | 9 | 31 |
| 19.2 | 11 | 62 | 0 | 10 | 29 |
| 19.3 | 10 | 61 | 3 | 4 | 28 |
| 19.4 | 2 | 54 | 0 | 3 | 31 |
| 19.5 | 8 | 59 | 0 | 5 | 29 |
| 19.6 | 5 | 63 | 1 | 2 | 25 |
| 19.7 | 3 | 58 | 0 | 4 | 34 |
| 19.8 | 0 | 66 | 0 | 3 | 34 |
| 19.9 | 5 | 58 | 0 | 4 | 30 |

**Figure S7.** Timeline plots pertaining to number of contacts between histones and BMAL1 PASA domain for the systems studied.

DNA-Histones (nonphosphorylated)

Number of contacts

| Frame | 1st | 2nd | 3rd | 4th | 5th |
| --- | --- | --- | --- | --- | --- |
| 0 | 267 | 284 | 295 | 264 | 308 |
| 0.1 | 267 | 284 | 295 | 263 | 298 |
| 0.2 | 267 | 284 | 282 | 277 | 305 |
| 0.3 | 248 | 284 | 316 | 282 | 295 |
| 0.4 | 270 | 285 | 288 | 264 | 294 |
| 0.5 | 280 | 262 | 305 | 267 | 294 |
| 0.6 | 253 | 241 | 282 | 263 | 281 |
| 0.7 | 255 | 259 | 297 | 246 | 286 |
| 0.8 | 265 | 245 | 288 | 252 | 298 |
| 0.9 | 251 | 248 | 315 | 242 | 288 |
| 1 | 256 | 240 | 287 | 234 | 283 |
| 1.1 | 253 | 254 | 298 | 235 | 282 |
| 1.2 | 271 | 250 | 284 | 241 | 305 |
| 1.3 | 259 | 253 | 287 | 247 | 311 |
| 1.4 | 247 | 231 | 269 | 230 | 315 |
| 1.5 | 244 | 250 | 279 | 261 | 328 |
| 1.6 | 231 | 245 | 287 | 270 | 341 |
| 1.7 | 273 | 232 | 283 | 240 | 345 |
| 1.8 | 255 | 235 | 266 | 259 | 356 |
| 1.9 | 263 | 242 | 295 | 272 | 303 |
| 2 | 260 | 256 | 284 | 284 | 336 |
| 2.1 | 274 | 250 | 293 | 271 | 330 |
| 2.2 | 262 | 223 | 298 | 285 | 354 |
| 2.3 | 255 | 243 | 278 | 308 | 321 |
| 2.4 | 282 | 264 | 306 | 272 | 340 |
| 2.5 | 274 | 254 | 315 | 279 | 314 |
| 2.6 | 272 | 270 | 302 | 264 | 347 |
| 2.7 | 293 | 253 | 318 | 236 | 348 |
| 2.8 | 293 | 272 | 320 | 275 | 344 |
| 2.9 | 288 | 249 | 323 | 270 | 328 |
| 3 | 282 | 272 | 316 | 275 | 338 |
| 3.1 | 293 | 267 | 307 | 276 | 336 |
| 3.2 | 293 | 264 | 305 | 267 | 337 |
| 3.3 | 331 | 283 | 297 | 278 | 349 |
| 3.4 | 319 | 271 | 319 | 266 | 334 |
| 3.5 | 281 | 270 | 332 | 265 | 339 |
| 3.6 | 313 | 272 | 313 | 276 | 345 |
| 3.7 | 296 | 269 | 342 | 275 | 341 |
| 3.8 | 293 | 283 | 337 | 274 | 365 |
| 3.9 | 302 | 267 | 338 | 260 | 356 |
| 4 | 299 | 251 | 308 | 267 | 350 |
| 4.1 | 324 | 257 | 333 | 266 | 341 |
| 4.2 | 299 | 244 | 320 | 269 | 360 |
| 4.3 | 304 | 237 | 317 | 262 | 359 |
| 4.4 | 308 | 243 | 315 | 284 | 357 |
| 4.5 | 305 | 241 | 321 | 280 | 373 |
| 4.6 | 278 | 222 | 357 | 301 | 360 |
| 4.7 | 294 | 251 | 313 | 281 | 365 |
| 4.8 | 315 | 245 | 325 | 261 | 372 |
| 4.9 | 321 | 265 | 332 | 265 | 361 |
| 5 | 302 | 262 | 315 | 270 | 361 |
| 5.1 | 302 | 273 | 308 | 285 | 376 |
| 5.2 | 321 | 237 | 290 | 294 | 362 |
| 5.3 | 312 | 245 | 301 | 274 | 377 |
| 5.4 | 286 | 246 | 319 | 286 | 376 |
| 5.5 | 298 | 242 | 306 | 316 | 354 |
| 5.6 | 328 | 239 | 331 | 322 | 362 |
| 5.7 | 326 | 260 | 307 | 299 | 357 |
| 5.8 | 307 | 243 | 316 | 301 | 351 |
| 5.9 | 309 | 240 | 313 | 330 | 373 |
| 6 | 314 | 260 | 342 | 302 | 350 |
| 6.1 | 295 | 233 | 313 | 296 | 340 |
| 6.2 | 302 | 235 | 320 | 316 | 366 |
| 6.3 | 300 | 243 | 324 | 302 | 355 |
| 6.4 | 308 | 236 | 344 | 299 | 364 |
| 6.5 | 307 | 252 | 348 | 299 | 375 |
| 6.6 | 316 | 255 | 344 | 274 | 370 |
| 6.7 | 307 | 278 | 321 | 298 | 362 |
| 6.8 | 304 | 239 | 333 | 300 | 353 |
| 6.9 | 309 | 232 | 376 | 288 | 382 |
| 7 | 299 | 256 | 356 | 290 | 342 |
| 7.1 | 283 | 252 | 347 | 308 | 344 |
| 7.2 | 332 | 242 | 332 | 286 | 350 |
| 7.3 | 301 | 269 | 350 | 282 | 350 |
| 7.4 | 311 | 259 | 338 | 302 | 345 |
| 7.5 | 304 | 276 | 330 | 318 | 351 |
| 7.6 | 328 | 261 | 316 | 289 | 345 |
| 7.7 | 319 | 266 | 348 | 299 | 360 |
| 7.8 | 326 | 260 | 353 | 328 | 364 |
| 7.9 | 294 | 267 | 338 | 329 | 376 |
| 8 | 318 | 274 | 325 | 315 | 346 |
| 8.1 | 326 | 253 | 325 | 334 | 349 |
| 8.2 | 305 | 243 | 318 | 329 | 355 |
| 8.3 | 293 | 254 | 318 | 315 | 344 |
| 8.4 | 295 | 272 | 336 | 301 | 333 |
| 8.5 | 313 | 254 | 319 | 295 | 360 |
| 8.6 | 304 | 255 | 319 | 316 | 350 |
| 8.7 | 302 | 281 | 348 | 323 | 348 |
| 8.8 | 309 | 291 | 341 | 305 | 350 |
| 8.9 | 299 | 259 | 350 | 314 | 346 |
| 9 | 305 | 249 | 344 | 308 | 343 |
| 9.1 | 311 | 278 | 339 | 306 | 343 |
| 9.2 | 295 | 277 | 332 | 316 | 361 |
| 9.3 | 297 | 265 | 329 | 319 | 354 |
| 9.4 | 330 | 278 | 344 | 304 | 344 |
| 9.5 | 285 | 254 | 332 | 306 | 358 |
| 9.6 | 316 | 266 | 342 | 307 | 371 |
| 9.7 | 313 | 262 | 324 | 304 | 346 |
| 9.8 | 314 | 289 | 338 | 311 | 362 |
| 9.9 | 320 | 273 | 339 | 319 | 361 |
| 10 | 317 | 294 | 353 | 290 | 330 |
| 10.1 | 292 | 297 | 328 | 314 | 348 |
| 10.2 | 306 | 290 | 353 | 312 | 368 |
| 10.3 | 262 | 299 | 348 | 288 | 329 |
| 10.4 | 312 | 284 | 348 | 310 | 321 |
| 10.5 | 316 | 299 | 322 | 310 | 336 |
| 10.6 | 298 | 275 | 325 | 307 | 328 |
| 10.7 | 323 | 295 | 332 | 298 | 315 |
| 10.8 | 305 | 271 | 320 | 319 | 317 |
| 10.9 | 290 | 282 | 318 | 303 | 329 |
| 11 | 319 | 262 | 338 | 315 | 321 |
| 11.1 | 309 | 260 | 332 | 313 | 344 |
| 11.2 | 283 | 280 | 340 | 300 | 337 |
| 11.3 | 285 | 290 | 320 | 302 | 336 |
| 11.4 | 282 | 252 | 317 | 303 | 320 |
| 11.5 | 269 | 258 | 315 | 314 | 344 |
| 11.6 | 278 | 276 | 318 | 313 | 348 |
| 11.7 | 292 | 274 | 309 | 310 | 342 |
| 11.8 | 282 | 267 | 323 | 305 | 336 |
| 11.9 | 275 | 280 | 319 | 287 | 351 |
| 12 | 288 | 292 | 305 | 310 | 323 |
| 12.1 | 281 | 280 | 316 | 311 | 340 |
| 12.2 | 313 | 280 | 312 | 296 | 340 |
| 12.3 | 300 | 269 | 318 | 318 | 327 |
| 12.4 | 297 | 265 | 325 | 302 | 335 |
| 12.5 | 293 | 271 | 310 | 306 | 334 |
| 12.6 | 303 | 267 | 307 | 312 | 327 |
| 12.7 | 317 | 302 | 302 | 311 | 348 |
| 12.8 | 299 | 283 | 326 | 320 | 349 |
| 12.9 | 301 | 280 | 327 | 326 | 340 |
| 13 | 299 | 293 | 323 | 332 | 359 |
| 13.1 | 296 | 290 | 331 | 321 | 349 |
| 13.2 | 288 | 270 | 326 | 316 | 336 |
| 13.3 | 317 | 289 | 353 | 308 | 329 |
| 13.4 | 302 | 270 | 322 | 314 | 341 |
| 13.5 | 307 | 280 | 337 | 300 | 349 |
| 13.6 | 315 | 281 | 340 | 293 | 326 |
| 13.7 | 307 | 287 | 337 | 307 | 356 |
| 13.8 | 296 | 287 | 351 | 311 | 343 |
| 13.9 | 309 | 306 | 331 | 318 | 359 |
| 14 | 301 | 290 | 341 | 325 | 356 |
| 14.1 | 298 | 283 | 339 | 323 | 342 |
| 14.2 | 301 | 286 | 319 | 334 | 338 |
| 14.3 | 284 | 254 | 323 | 329 | 333 |
| 14.4 | 282 | 285 | 324 | 332 | 347 |
| 14.5 | 271 | 290 | 325 | 330 | 340 |
| 14.6 | 287 | 281 | 320 | 324 | 373 |
| 14.7 | 260 | 259 | 335 | 344 | 359 |
| 14.8 | 281 | 282 | 339 | 332 | 371 |
| 14.9 | 277 | 300 | 359 | 349 | 353 |
| 15 | 285 | 302 | 340 | 340 | 359 |
| 15.1 | 309 | 272 | 344 | 345 | 343 |
| 15.2 | 308 | 285 | 340 | 328 | 366 |
| 15.3 | 289 | 292 | 335 | 337 | 375 |
| 15.4 | 293 | 295 | 328 | 328 | 348 |
| 15.5 | 278 | 263 | 341 | 340 | 366 |
| 15.6 | 304 | 290 | 344 | 332 | 366 |
| 15.7 | 272 | 273 | 341 | 333 | 334 |
| 15.8 | 284 | 293 | 327 | 316 | 348 |
| 15.9 | 279 | 294 | 354 | 340 | 369 |
| 16 | 267 | 265 | 327 | 314 | 350 |
| 16.1 | 270 | 295 | 340 | 335 | 366 |
| 16.2 | 285 | 255 | 335 | 329 | 366 |
| 16.3 | 285 | 265 | 332 | 322 | 347 |
| 16.4 | 292 | 262 | 354 | 319 | 356 |
| 16.5 | 302 | 273 | 361 | 320 | 361 |
| 16.6 | 275 | 273 | 345 | 341 | 358 |
| 16.7 | 284 | 260 | 348 | 333 | 354 |
| 16.8 | 274 | 266 | 350 | 354 | 333 |
| 16.9 | 297 | 263 | 364 | 350 | 337 |
| 17 | 283 | 281 | 354 | 338 | 312 |
| 17.1 | 293 | 276 | 350 | 321 | 346 |
| 17.2 | 280 | 246 | 334 | 325 | 323 |
| 17.3 | 298 | 289 | 312 | 323 | 328 |
| 17.4 | 292 | 292 | 335 | 335 | 323 |
| 17.5 | 283 | 274 | 357 | 335 | 339 |
| 17.6 | 277 | 276 | 340 | 337 | 333 |
| 17.7 | 284 | 262 | 347 | 335 | 345 |
| 17.8 | 305 | 282 | 354 | 322 | 330 |
| 17.9 | 298 | 277 | 325 | 341 | 340 |
| 18 | 288 | 262 | 342 | 345 | 324 |
| 18.1 | 285 | 253 | 338 | 335 | 343 |
| 18.2 | 301 | 258 | 333 | 328 | 353 |
| 18.3 | 298 | 256 | 331 | 321 | 321 |
| 18.4 | 310 | 263 | 361 | 316 | 348 |
| 18.5 | 286 | 257 | 371 | 328 | 337 |
| 18.6 | 292 | 258 | 348 | 352 | 313 |
| 18.7 | 296 | 272 | 348 | 342 | 337 |
| 18.8 | 323 | 273 | 334 | 336 | 327 |
| 18.9 | 294 | 265 | 331 | 324 | 336 |
| 19 | 277 | 243 | 341 | 338 | 353 |
| 19.1 | 291 | 248 | 329 | 338 | 329 |
| 19.2 | 299 | 263 | 325 | 324 | 338 |
| 19.3 | 309 | 244 | 339 | 325 | 335 |
| 19.4 | 280 | 241 | 339 | 351 | 327 |
| 19.5 | 280 | 241 | 348 | 341 | 342 |
| 19.6 | 304 | 260 | 340 | 346 | 346 |
| 19.7 | 303 | 251 | 350 | 317 | 316 |
| 19.8 | 300 | 278 | 326 | 332 | 332 |
| 19.9 | 303 | 247 | 353 | 300 | 336 |

DNA-Histones (BMAL1S78)

Number of contacts

| Frame | 1st | 2nd | 3rd | 4th | 5th |
| --- | --- | --- | --- | --- | --- |
| 0 | 284 | 291 | 278 | 297 | 263 |
| 0.1 | 304 | 301 | 307 | 297 | 276 |
| 0.2 | 338 | 289 | 291 | 318 | 313 |
| 0.3 | 348 | 262 | 255 | 318 | 305 |
| 0.4 | 338 | 266 | 268 | 318 | 318 |
| 0.5 | 337 | 281 | 283 | 302 | 324 |
| 0.6 | 335 | 281 | 284 | 308 | 309 |
| 0.7 | 324 | 275 | 290 | 294 | 296 |
| 0.8 | 309 | 287 | 304 | 310 | 311 |
| 0.9 | 337 | 265 | 307 | 303 | 323 |
| 1 | 305 | 275 | 304 | 290 | 326 |
| 1.1 | 310 | 267 | 309 | 307 | 292 |
| 1.2 | 329 | 258 | 325 | 319 | 296 |
| 1.3 | 300 | 272 | 335 | 293 | 310 |
| 1.4 | 336 | 264 | 318 | 293 | 298 |
| 1.5 | 320 | 263 | 312 | 296 | 303 |
| 1.6 | 320 | 276 | 338 | 317 | 286 |
| 1.7 | 309 | 252 | 317 | 280 | 295 |
| 1.8 | 344 | 250 | 333 | 299 | 286 |
| 1.9 | 320 | 241 | 311 | 316 | 278 |
| 2 | 308 | 278 | 316 | 316 | 305 |
| 2.1 | 300 | 267 | 317 | 297 | 300 |
| 2.2 | 279 | 250 | 304 | 295 | 286 |
| 2.3 | 304 | 267 | 326 | 312 | 301 |
| 2.4 | 307 | 259 | 304 | 300 | 310 |
| 2.5 | 279 | 266 | 297 | 301 | 313 |
| 2.6 | 289 | 271 | 281 | 292 | 299 |
| 2.7 | 308 | 305 | 295 | 306 | 282 |
| 2.8 | 300 | 293 | 326 | 301 | 306 |
| 2.9 | 302 | 291 | 303 | 314 | 311 |
| 3 | 300 | 287 | 302 | 314 | 304 |
| 3.1 | 288 | 271 | 298 | 301 | 311 |
| 3.2 | 304 | 277 | 298 | 322 | 304 |
| 3.3 | 295 | 270 | 313 | 300 | 312 |
| 3.4 | 309 | 281 | 331 | 322 | 273 |
| 3.5 | 300 | 269 | 304 | 308 | 304 |
| 3.6 | 301 | 301 | 290 | 310 | 338 |
| 3.7 | 304 | 316 | 299 | 310 | 312 |
| 3.8 | 286 | 318 | 308 | 322 | 327 |
| 3.9 | 294 | 283 | 318 | 318 | 321 |
| 4 | 325 | 325 | 315 | 314 | 318 |
| 4.1 | 310 | 305 | 330 | 302 | 295 |
| 4.2 | 301 | 309 | 303 | 324 | 291 |
| 4.3 | 296 | 320 | 287 | 310 | 325 |
| 4.4 | 302 | 307 | 313 | 325 | 300 |
| 4.5 | 301 | 314 | 300 | 332 | 315 |
| 4.6 | 266 | 315 | 290 | 308 | 299 |
| 4.7 | 306 | 306 | 308 | 313 | 324 |
| 4.8 | 306 | 334 | 282 | 302 | 321 |
| 4.9 | 299 | 314 | 305 | 298 | 324 |
| 5 | 287 | 331 | 290 | 296 | 337 |
| 5.1 | 286 | 308 | 283 | 309 | 301 |
| 5.2 | 294 | 309 | 287 | 308 | 317 |
| 5.3 | 296 | 310 | 284 | 292 | 298 |
| 5.4 | 300 | 309 | 296 | 294 | 314 |
| 5.5 | 310 | 308 | 298 | 312 | 328 |
| 5.6 | 287 | 331 | 320 | 318 | 318 |
| 5.7 | 308 | 336 | 307 | 299 | 298 |
| 5.8 | 286 | 315 | 287 | 303 | 305 |
| 5.9 | 310 | 312 | 296 | 309 | 284 |
| 6 | 287 | 309 | 301 | 312 | 306 |
| 6.1 | 307 | 310 | 288 | 323 | 300 |
| 6.2 | 286 | 302 | 298 | 314 | 305 |
| 6.3 | 314 | 302 | 303 | 295 | 303 |
| 6.4 | 310 | 320 | 263 | 316 | 281 |
| 6.5 | 311 | 300 | 283 | 294 | 320 |
| 6.6 | 289 | 298 | 316 | 282 | 327 |
| 6.7 | 305 | 306 | 308 | 289 | 316 |
| 6.8 | 304 | 328 | 303 | 290 | 298 |
| 6.9 | 298 | 311 | 290 | 298 | 300 |
| 7 | 299 | 326 | 292 | 307 | 313 |
| 7.1 | 303 | 314 | 290 | 306 | 312 |
| 7.2 | 274 | 301 | 301 | 304 | 324 |
| 7.3 | 281 | 308 | 306 | 276 | 306 |
| 7.4 | 267 | 291 | 295 | 300 | 306 |
| 7.5 | 291 | 307 | 335 | 288 | 295 |
| 7.6 | 288 | 291 | 314 | 298 | 298 |
| 7.7 | 293 | 286 | 299 | 276 | 309 |
| 7.8 | 296 | 315 | 281 | 290 | 331 |
| 7.9 | 279 | 278 | 292 | 292 | 334 |
| 8 | 300 | 267 | 301 | 300 | 307 |
| 8.1 | 284 | 302 | 295 | 273 | 299 |
| 8.2 | 295 | 310 | 308 | 297 | 324 |
| 8.3 | 291 | 279 | 293 | 310 | 302 |
| 8.4 | 287 | 312 | 302 | 289 | 293 |
| 8.5 | 280 | 289 | 309 | 279 | 299 |
| 8.6 | 278 | 290 | 288 | 316 | 306 |
| 8.7 | 284 | 294 | 317 | 302 | 306 |
| 8.8 | 283 | 270 | 302 | 284 | 305 |
| 8.9 | 314 | 314 | 320 | 325 | 301 |
| 9 | 283 | 305 | 307 | 329 | 308 |
| 9.1 | 302 | 298 | 314 | 330 | 317 |
| 9.2 | 305 | 292 | 298 | 314 | 319 |
| 9.3 | 286 | 292 | 307 | 310 | 325 |
| 9.4 | 308 | 300 | 276 | 309 | 322 |
| 9.5 | 340 | 299 | 303 | 323 | 304 |
| 9.6 | 311 | 266 | 318 | 297 | 307 |
| 9.7 | 307 | 284 | 289 | 321 | 321 |
| 9.8 | 326 | 291 | 301 | 315 | 319 |
| 9.9 | 311 | 288 | 287 | 313 | 323 |
| 10 | 303 | 289 | 302 | 332 | 324 |
| 10.1 | 310 | 289 | 277 | 314 | 303 |
| 10.2 | 296 | 272 | 295 | 322 | 330 |
| 10.3 | 315 | 268 | 313 | 323 | 356 |
| 10.4 | 291 | 266 | 265 | 322 | 341 |
| 10.5 | 320 | 273 | 290 | 317 | 334 |
| 10.6 | 310 | 270 | 285 | 317 | 353 |
| 10.7 | 312 | 258 | 280 | 321 | 331 |
| 10.8 | 298 | 284 | 293 | 331 | 356 |
| 10.9 | 287 | 277 | 301 | 313 | 331 |
| 11 | 308 | 258 | 298 | 321 | 341 |
| 11.1 | 310 | 274 | 302 | 315 | 319 |
| 11.2 | 293 | 266 | 311 | 312 | 342 |
| 11.3 | 292 | 284 | 302 | 335 | 324 |
| 11.4 | 316 | 278 | 307 | 315 | 328 |
| 11.5 | 304 | 290 | 289 | 296 | 311 |
| 11.6 | 286 | 294 | 270 | 320 | 314 |
| 11.7 | 307 | 284 | 277 | 333 | 315 |
| 11.8 | 302 | 293 | 286 | 310 | 332 |
| 11.9 | 306 | 282 | 288 | 301 | 308 |
| 12 | 297 | 279 | 285 | 324 | 301 |
| 12.1 | 291 | 276 | 275 | 324 | 281 |
| 12.2 | 305 | 281 | 310 | 317 | 312 |
| 12.3 | 293 | 271 | 291 | 304 | 326 |
| 12.4 | 320 | 296 | 281 | 317 | 317 |
| 12.5 | 308 | 288 | 319 | 299 | 304 |
| 12.6 | 317 | 271 | 313 | 307 | 305 |
| 12.7 | 310 | 293 | 321 | 306 | 279 |
| 12.8 | 332 | 290 | 312 | 314 | 299 |
| 12.9 | 321 | 283 | 327 | 316 | 315 |
| 13 | 296 | 282 | 300 | 309 | 313 |
| 13.1 | 306 | 271 | 303 | 334 | 307 |
| 13.2 | 318 | 316 | 310 | 307 | 303 |
| 13.3 | 316 | 286 | 304 | 325 | 316 |
| 13.4 | 322 | 293 | 299 | 326 | 293 |
| 13.5 | 302 | 284 | 301 | 326 | 320 |
| 13.6 | 310 | 282 | 303 | 346 | 303 |
| 13.7 | 313 | 291 | 302 | 329 | 328 |
| 13.8 | 319 | 279 | 303 | 329 | 298 |
| 13.9 | 320 | 293 | 271 | 326 | 311 |
| 14 | 303 | 284 | 269 | 332 | 318 |
| 14.1 | 303 | 272 | 298 | 339 | 308 |
| 14.2 | 286 | 301 | 286 | 326 | 326 |
| 14.3 | 291 | 291 | 306 | 316 | 324 |
| 14.4 | 299 | 290 | 314 | 324 | 311 |
| 14.5 | 290 | 296 | 299 | 336 | 315 |
| 14.6 | 303 | 277 | 312 | 326 | 316 |
| 14.7 | 296 | 302 | 294 | 351 | 310 |
| 14.8 | 284 | 280 | 307 | 325 | 329 |
| 14.9 | 282 | 286 | 304 | 324 | 335 |
| 15 | 299 | 283 | 283 | 319 | 335 |
| 15.1 | 294 | 290 | 267 | 331 | 326 |
| 15.2 | 309 | 292 | 314 | 347 | 318 |
| 15.3 | 310 | 280 | 280 | 331 | 310 |
| 15.4 | 279 | 278 | 289 | 332 | 324 |
| 15.5 | 283 | 275 | 297 | 324 | 332 |
| 15.6 | 296 | 287 | 299 | 345 | 317 |
| 15.7 | 273 | 285 | 305 | 315 | 319 |
| 15.8 | 292 | 276 | 305 | 315 | 324 |
| 15.9 | 287 | 278 | 297 | 313 | 327 |
| 16 | 283 | 281 | 300 | 356 | 303 |
| 16.1 | 299 | 277 | 313 | 322 | 291 |
| 16.2 | 300 | 278 | 327 | 329 | 290 |
| 16.3 | 298 | 256 | 304 | 334 | 315 |
| 16.4 | 282 | 273 | 282 | 328 | 330 |
| 16.5 | 292 | 261 | 283 | 333 | 297 |
| 16.6 | 298 | 285 | 313 | 332 | 299 |
| 16.7 | 311 | 284 | 300 | 321 | 301 |
| 16.8 | 311 | 272 | 307 | 325 | 300 |
| 16.9 | 291 | 285 | 322 | 324 | 313 |
| 17 | 319 | 282 | 312 | 308 | 323 |
| 17.1 | 301 | 276 | 292 | 337 | 309 |
| 17.2 | 300 | 276 | 299 | 337 | 330 |
| 17.3 | 295 | 281 | 310 | 353 | 322 |
| 17.4 | 297 | 286 | 320 | 326 | 310 |
| 17.5 | 301 | 272 | 321 | 308 | 295 |
| 17.6 | 313 | 278 | 304 | 336 | 305 |
| 17.7 | 277 | 277 | 312 | 329 | 302 |
| 17.8 | 297 | 290 | 319 | 331 | 284 |
| 17.9 | 309 | 281 | 314 | 349 | 301 |
| 18 | 288 | 294 | 331 | 347 | 311 |
| 18.1 | 293 | 248 | 332 | 343 | 297 |
| 18.2 | 311 | 258 | 329 | 368 | 296 |
| 18.3 | 303 | 304 | 335 | 319 | 308 |
| 18.4 | 301 | 288 | 317 | 355 | 302 |
| 18.5 | 282 | 287 | 336 | 334 | 298 |
| 18.6 | 299 | 298 | 312 | 335 | 304 |
| 18.7 | 308 | 297 | 316 | 324 | 303 |
| 18.8 | 297 | 274 | 294 | 333 | 303 |
| 18.9 | 305 | 291 | 340 | 329 | 312 |
| 19 | 313 | 287 | 316 | 338 | 300 |
| 19.1 | 279 | 279 | 340 | 351 | 298 |
| 19.2 | 304 | 281 | 345 | 352 | 294 |
| 19.3 | 318 | 295 | 320 | 346 | 296 |
| 19.4 | 336 | 282 | 331 | 356 | 326 |
| 19.5 | 293 | 273 | 358 | 360 | 325 |
| 19.6 | 312 | 288 | 344 | 364 | 328 |
| 19.7 | 310 | 276 | 338 | 345 | 328 |
| 19.8 | 294 | 298 | 333 | 353 | 291 |
| 19.9 | 310 | 278 | 352 | 343 | 329 |

DNA-Histones (CLOCKS38/42)

Number of contacts

| Frame | 1st | 2nd | 3rd | 4th | 5th |
| --- | --- | --- | --- | --- | --- |
| 0 | 288 | 311 | 259 | 318 | 299 |
| 0.1 | 296 | 332 | 267 | 332 | 305 |
| 0.2 | 288 | 298 | 263 | 300 | 327 |
| 0.3 | 294 | 269 | 293 | 294 | 322 |
| 0.4 | 290 | 255 | 306 | 268 | 322 |
| 0.5 | 284 | 265 | 319 | 262 | 336 |
| 0.6 | 235 | 297 | 300 | 252 | 320 |
| 0.7 | 269 | 277 | 282 | 265 | 342 |
| 0.8 | 248 | 265 | 261 | 287 | 366 |
| 0.9 | 240 | 284 | 272 | 284 | 346 |
| 1 | 256 | 305 | 262 | 290 | 340 |
| 1.1 | 262 | 316 | 282 | 281 | 314 |
| 1.2 | 260 | 331 | 276 | 290 | 330 |
| 1.3 | 222 | 322 | 295 | 291 | 302 |
| 1.4 | 227 | 348 | 281 | 296 | 330 |
| 1.5 | 238 | 331 | 269 | 274 | 332 |
| 1.6 | 229 | 322 | 296 | 317 | 316 |
| 1.7 | 233 | 318 | 277 | 325 | 325 |
| 1.8 | 243 | 326 | 270 | 293 | 305 |
| 1.9 | 243 | 325 | 269 | 283 | 304 |
| 2 | 239 | 341 | 259 | 296 | 318 |
| 2.1 | 236 | 344 | 257 | 300 | 327 |
| 2.2 | 244 | 351 | 275 | 299 | 348 |
| 2.3 | 251 | 333 | 245 | 297 | 322 |
| 2.4 | 275 | 338 | 285 | 271 | 319 |
| 2.5 | 306 | 344 | 265 | 308 | 355 |
| 2.6 | 300 | 339 | 279 | 285 | 354 |
| 2.7 | 275 | 339 | 277 | 285 | 338 |
| 2.8 | 271 | 331 | 297 | 284 | 330 |
| 2.9 | 264 | 334 | 265 | 287 | 328 |
| 3 | 311 | 321 | 275 | 296 | 347 |
| 3.1 | 299 | 360 | 265 | 315 | 340 |
| 3.2 | 288 | 327 | 277 | 302 | 339 |
| 3.3 | 300 | 324 | 297 | 307 | 342 |
| 3.4 | 278 | 321 | 279 | 305 | 323 |
| 3.5 | 261 | 322 | 261 | 331 | 337 |
| 3.6 | 299 | 342 | 259 | 311 | 337 |
| 3.7 | 297 | 310 | 245 | 306 | 328 |
| 3.8 | 307 | 311 | 270 | 309 | 329 |
| 3.9 | 305 | 339 | 274 | 338 | 340 |
| 4 | 330 | 336 | 299 | 320 | 299 |
| 4.1 | 315 | 318 | 293 | 308 | 298 |
| 4.2 | 321 | 330 | 274 | 324 | 301 |
| 4.3 | 314 | 325 | 270 | 328 | 310 |
| 4.4 | 315 | 350 | 297 | 323 | 297 |
| 4.5 | 325 | 358 | 283 | 333 | 328 |
| 4.6 | 332 | 356 | 282 | 326 | 299 |
| 4.7 | 301 | 338 | 286 | 333 | 306 |
| 4.8 | 345 | 364 | 318 | 327 | 309 |
| 4.9 | 314 | 368 | 317 | 328 | 309 |
| 5 | 334 | 353 | 291 | 326 | 311 |
| 5.1 | 338 | 327 | 308 | 317 | 294 |
| 5.2 | 326 | 327 | 312 | 322 | 293 |
| 5.3 | 296 | 332 | 322 | 318 | 289 |
| 5.4 | 305 | 363 | 342 | 329 | 306 |
| 5.5 | 302 | 335 | 304 | 332 | 317 |
| 5.6 | 306 | 319 | 329 | 312 | 330 |
| 5.7 | 312 | 354 | 338 | 324 | 319 |
| 5.8 | 301 | 355 | 328 | 338 | 307 |
| 5.9 | 336 | 347 | 305 | 318 | 298 |
| 6 | 321 | 331 | 302 | 351 | 296 |
| 6.1 | 307 | 344 | 299 | 336 | 301 |
| 6.2 | 331 | 341 | 313 | 335 | 318 |
| 6.3 | 310 | 346 | 315 | 333 | 301 |
| 6.4 | 316 | 352 | 330 | 327 | 298 |
| 6.5 | 312 | 330 | 307 | 351 | 335 |
| 6.6 | 298 | 337 | 318 | 366 | 333 |
| 6.7 | 281 | 334 | 350 | 355 | 317 |
| 6.8 | 307 | 359 | 331 | 330 | 320 |
| 6.9 | 309 | 338 | 345 | 333 | 322 |
| 7 | 300 | 349 | 342 | 355 | 313 |
| 7.1 | 312 | 354 | 321 | 340 | 344 |
| 7.2 | 302 | 358 | 324 | 354 | 363 |
| 7.3 | 312 | 322 | 326 | 337 | 347 |
| 7.4 | 313 | 351 | 344 | 329 | 354 |
| 7.5 | 315 | 350 | 351 | 337 | 353 |
| 7.6 | 322 | 374 | 344 | 326 | 346 |
| 7.7 | 322 | 359 | 328 | 326 | 349 |
| 7.8 | 318 | 350 | 348 | 332 | 335 |
| 7.9 | 338 | 328 | 331 | 328 | 338 |
| 8 | 321 | 332 | 331 | 334 | 349 |
| 8.1 | 323 | 330 | 315 | 341 | 325 |
| 8.2 | 332 | 329 | 337 | 327 | 355 |
| 8.3 | 324 | 327 | 339 | 333 | 382 |
| 8.4 | 305 | 350 | 321 | 323 | 347 |
| 8.5 | 312 | 339 | 341 | 351 | 366 |
| 8.6 | 289 | 339 | 357 | 341 | 370 |
| 8.7 | 307 | 346 | 337 | 350 | 360 |
| 8.8 | 320 | 330 | 369 | 320 | 361 |
| 8.9 | 293 | 327 | 342 | 340 | 375 |
| 9 | 309 | 334 | 333 | 335 | 377 |
| 9.1 | 300 | 321 | 340 | 335 | 392 |
| 9.2 | 304 | 328 | 359 | 333 | 383 |
| 9.3 | 322 | 362 | 342 | 343 | 375 |
| 9.4 | 301 | 376 | 352 | 313 | 389 |
| 9.5 | 289 | 333 | 340 | 327 | 397 |
| 9.6 | 311 | 339 | 324 | 337 | 376 |
| 9.7 | 301 | 336 | 347 | 342 | 377 |
| 9.8 | 300 | 336 | 357 | 336 | 391 |
| 9.9 | 294 | 329 | 348 | 349 | 404 |
| 10 | 328 | 338 | 339 | 339 | 355 |
| 10.1 | 290 | 362 | 338 | 316 | 374 |
| 10.2 | 307 | 335 | 332 | 348 | 357 |
| 10.3 | 285 | 317 | 340 | 365 | 364 |
| 10.4 | 292 | 340 | 339 | 370 | 366 |
| 10.5 | 263 | 327 | 323 | 385 | 398 |
| 10.6 | 278 | 346 | 357 | 333 | 357 |
| 10.7 | 269 | 319 | 324 | 363 | 356 |
| 10.8 | 275 | 330 | 349 | 370 | 358 |
| 10.9 | 304 | 307 | 334 | 347 | 372 |
| 11 | 276 | 329 | 342 | 324 | 356 |
| 11.1 | 279 | 339 | 341 | 353 | 355 |
| 11.2 | 291 | 318 | 329 | 347 | 327 |
| 11.3 | 264 | 337 | 346 | 340 | 352 |
| 11.4 | 277 | 329 | 340 | 358 | 339 |
| 11.5 | 275 | 354 | 341 | 361 | 347 |
| 11.6 | 292 | 370 | 316 | 350 | 343 |
| 11.7 | 264 | 332 | 336 | 355 | 367 |
| 11.8 | 281 | 339 | 327 | 369 | 348 |
| 11.9 | 278 | 329 | 324 | 340 | 341 |
| 12 | 277 | 324 | 347 | 347 | 380 |
| 12.1 | 284 | 332 | 345 | 357 | 373 |
| 12.2 | 283 | 340 | 339 | 343 | 328 |
| 12.3 | 301 | 352 | 329 | 372 | 358 |
| 12.4 | 283 | 321 | 341 | 345 | 358 |
| 12.5 | 280 | 365 | 353 | 371 | 325 |
| 12.6 | 282 | 325 | 335 | 350 | 336 |
| 12.7 | 296 | 333 | 352 | 362 | 350 |
| 12.8 | 262 | 321 | 363 | 343 | 333 |
| 12.9 | 294 | 346 | 357 | 377 | 336 |
| 13 | 287 | 309 | 348 | 356 | 361 |
| 13.1 | 308 | 313 | 342 | 359 | 367 |
| 13.2 | 304 | 337 | 350 | 369 | 366 |
| 13.3 | 285 | 336 | 336 | 346 | 369 |
| 13.4 | 283 | 358 | 348 | 371 | 379 |
| 13.5 | 304 | 329 | 319 | 347 | 338 |
| 13.6 | 292 | 359 | 336 | 357 | 357 |
| 13.7 | 287 | 365 | 339 | 376 | 346 |
| 13.8 | 285 | 353 | 364 | 352 | 355 |
| 13.9 | 289 | 353 | 342 | 339 | 369 |
| 14 | 301 | 369 | 342 | 374 | 363 |
| 14.1 | 298 | 359 | 357 | 392 | 334 |
| 14.2 | 319 | 333 | 343 | 383 | 349 |
| 14.3 | 270 | 345 | 342 | 376 | 354 |
| 14.4 | 293 | 354 | 354 | 395 | 312 |
| 14.5 | 281 | 355 | 331 | 371 | 341 |
| 14.6 | 258 | 347 | 342 | 355 | 336 |
| 14.7 | 273 | 369 | 351 | 378 | 322 |
| 14.8 | 266 | 356 | 337 | 383 | 364 |
| 14.9 | 276 | 352 | 344 | 380 | 317 |
| 15 | 270 | 325 | 350 | 341 | 344 |
| 15.1 | 270 | 356 | 354 | 375 | 345 |
| 15.2 | 280 | 343 | 339 | 345 | 327 |
| 15.3 | 294 | 352 | 357 | 354 | 342 |
| 15.4 | 290 | 364 | 355 | 357 | 337 |
| 15.5 | 265 | 375 | 334 | 355 | 369 |
| 15.6 | 264 | 363 | 357 | 356 | 355 |
| 15.7 | 267 | 363 | 357 | 344 | 326 |
| 15.8 | 291 | 350 | 346 | 351 | 350 |
| 15.9 | 267 | 358 | 353 | 355 | 323 |
| 16 | 297 | 322 | 350 | 358 | 349 |
| 16.1 | 297 | 345 | 355 | 353 | 325 |
| 16.2 | 286 | 340 | 350 | 371 | 332 |
| 16.3 | 282 | 341 | 363 | 346 | 335 |
| 16.4 | 295 | 328 | 344 | 372 | 344 |
| 16.5 | 269 | 346 | 339 | 358 | 346 |
| 16.6 | 290 | 351 | 346 | 353 | 323 |
| 16.7 | 295 | 321 | 344 | 367 | 347 |
| 16.8 | 304 | 376 | 324 | 345 | 375 |
| 16.9 | 306 | 352 | 354 | 342 | 387 |
| 17 | 272 | 348 | 328 | 359 | 380 |
| 17.1 | 294 | 323 | 348 | 347 | 385 |
| 17.2 | 280 | 353 | 351 | 357 | 347 |
| 17.3 | 283 | 332 | 347 | 349 | 317 |
| 17.4 | 284 | 371 | 355 | 358 | 331 |
| 17.5 | 282 | 338 | 351 | 348 | 335 |
| 17.6 | 284 | 353 | 361 | 356 | 340 |
| 17.7 | 264 | 326 | 335 | 347 | 359 |
| 17.8 | 269 | 349 | 348 | 358 | 348 |
| 17.9 | 282 | 366 | 345 | 355 | 360 |
| 18 | 267 | 366 | 348 | 344 | 339 |
| 18.1 | 282 | 341 | 362 | 331 | 359 |
| 18.2 | 277 | 349 | 357 | 341 | 332 |
| 18.3 | 269 | 351 | 334 | 327 | 358 |
| 18.4 | 289 | 345 | 335 | 342 | 335 |
| 18.5 | 284 | 359 | 353 | 298 | 348 |
| 18.6 | 267 | 357 | 353 | 342 | 371 |
| 18.7 | 282 | 315 | 363 | 340 | 356 |
| 18.8 | 262 | 340 | 354 | 351 | 340 |
| 18.9 | 251 | 345 | 365 | 330 | 354 |
| 19 | 256 | 348 | 364 | 320 | 351 |
| 19.1 | 274 | 309 | 371 | 319 | 350 |
| 19.2 | 282 | 333 | 344 | 315 | 331 |
| 19.3 | 250 | 315 | 348 | 336 | 346 |
| 19.4 | 260 | 336 | 360 | 333 | 341 |
| 19.5 | 279 | 351 | 346 | 320 | 336 |
| 19.6 | 253 | 333 | 348 | 326 | 346 |
| 19.7 | 250 | 363 | 358 | 312 | 337 |
| 19.8 | 255 | 337 | 335 | 316 | 341 |
| 19.9 | 257 | 361 | 341 | 322 | 346 |

DNA-Histones (CLOCKS38/42/BMAL1S78)

Number of contacts

| Frame | 1st | 2nd | 3rd | 4th | 5th |
| --- | --- | --- | --- | --- | --- |
| 0 | 304 | 272 | 296 | 310 | 311 |
| 0.1 | 305 | 273 | 302 | 327 | 329 |
| 0.2 | 264 | 305 | 310 | 337 | 341 |
| 0.3 | 259 | 291 | 297 | 303 | 342 |
| 0.4 | 244 | 312 | 277 | 326 | 352 |
| 0.5 | 251 | 300 | 291 | 307 | 338 |
| 0.6 | 263 | 297 | 292 | 309 | 314 |
| 0.7 | 252 | 306 | 265 | 317 | 320 |
| 0.8 | 259 | 306 | 279 | 312 | 297 |
| 0.9 | 270 | 293 | 306 | 329 | 319 |
| 1 | 265 | 259 | 276 | 321 | 293 |
| 1.1 | 281 | 260 | 301 | 322 | 330 |
| 1.2 | 278 | 258 | 300 | 341 | 313 |
| 1.3 | 254 | 252 | 295 | 341 | 333 |
| 1.4 | 256 | 246 | 297 | 335 | 334 |
| 1.5 | 262 | 252 | 317 | 336 | 350 |
| 1.6 | 283 | 243 | 296 | 320 | 319 |
| 1.7 | 297 | 239 | 316 | 306 | 313 |
| 1.8 | 305 | 239 | 306 | 316 | 320 |
| 1.9 | 284 | 229 | 297 | 336 | 331 |
| 2 | 290 | 250 | 306 | 331 | 333 |
| 2.1 | 279 | 246 | 287 | 304 | 319 |
| 2.2 | 294 | 238 | 289 | 330 | 349 |
| 2.3 | 302 | 237 | 285 | 316 | 330 |
| 2.4 | 288 | 244 | 265 | 315 | 359 |
| 2.5 | 287 | 223 | 284 | 310 | 332 |
| 2.6 | 296 | 244 | 281 | 327 | 346 |
| 2.7 | 279 | 228 | 283 | 302 | 322 |
| 2.8 | 283 | 236 | 304 | 319 | 326 |
| 2.9 | 295 | 229 | 275 | 332 | 337 |
| 3 | 284 | 224 | 287 | 307 | 326 |
| 3.1 | 278 | 235 | 278 | 332 | 349 |
| 3.2 | 305 | 221 | 274 | 309 | 323 |
| 3.3 | 290 | 213 | 259 | 320 | 319 |
| 3.4 | 270 | 215 | 278 | 335 | 330 |
| 3.5 | 266 | 235 | 287 | 331 | 342 |
| 3.6 | 272 | 218 | 287 | 296 | 311 |
| 3.7 | 284 | 216 | 298 | 308 | 324 |
| 3.8 | 252 | 220 | 271 | 322 | 326 |
| 3.9 | 247 | 218 | 263 | 308 | 337 |
| 4 | 259 | 238 | 285 | 309 | 320 |
| 4.1 | 274 | 194 | 275 | 306 | 324 |
| 4.2 | 264 | 216 | 281 | 336 | 341 |
| 4.3 | 243 | 203 | 280 | 310 | 328 |
| 4.4 | 253 | 196 | 294 | 336 | 323 |
| 4.5 | 261 | 207 | 282 | 314 | 321 |
| 4.6 | 258 | 205 | 285 | 311 | 320 |
| 4.7 | 249 | 205 | 272 | 318 | 334 |
| 4.8 | 259 | 193 | 280 | 328 | 333 |
| 4.9 | 261 | 185 | 270 | 341 | 320 |
| 5 | 266 | 187 | 286 | 311 | 329 |
| 5.1 | 269 | 181 | 299 | 325 | 341 |
| 5.2 | 283 | 201 | 293 | 332 | 315 |
| 5.3 | 290 | 189 | 326 | 335 | 341 |
| 5.4 | 305 | 229 | 309 | 324 | 375 |
| 5.5 | 298 | 227 | 313 | 338 | 340 |
| 5.6 | 284 | 215 | 315 | 321 | 330 |
| 5.7 | 300 | 219 | 318 | 326 | 343 |
| 5.8 | 284 | 215 | 321 | 332 | 318 |
| 5.9 | 292 | 220 | 308 | 332 | 352 |
| 6 | 289 | 225 | 322 | 340 | 323 |
| 6.1 | 291 | 235 | 325 | 332 | 331 |
| 6.2 | 262 | 223 | 331 | 365 | 349 |
| 6.3 | 285 | 221 | 346 | 347 | 339 |
| 6.4 | 285 | 242 | 340 | 351 | 330 |
| 6.5 | 278 | 219 | 338 | 349 | 324 |
| 6.6 | 281 | 236 | 332 | 349 | 312 |
| 6.7 | 315 | 225 | 336 | 328 | 323 |
| 6.8 | 286 | 225 | 308 | 317 | 318 |
| 6.9 | 314 | 218 | 310 | 342 | 340 |
| 7 | 291 | 217 | 325 | 322 | 351 |
| 7.1 | 291 | 223 | 332 | 300 | 357 |
| 7.2 | 299 | 229 | 329 | 296 | 325 |
| 7.3 | 303 | 213 | 330 | 327 | 316 |
| 7.4 | 310 | 220 | 340 | 311 | 318 |
| 7.5 | 301 | 226 | 346 | 336 | 330 |
| 7.6 | 312 | 214 | 327 | 358 | 316 |
| 7.7 | 303 | 237 | 339 | 329 | 323 |
| 7.8 | 295 | 219 | 342 | 352 | 330 |
| 7.9 | 314 | 221 | 317 | 344 | 292 |
| 8 | 300 | 225 | 312 | 340 | 319 |
| 8.1 | 280 | 213 | 318 | 350 | 339 |
| 8.2 | 280 | 216 | 291 | 315 | 362 |
| 8.3 | 290 | 247 | 316 | 337 | 363 |
| 8.4 | 274 | 236 | 310 | 340 | 359 |
| 8.5 | 297 | 221 | 320 | 347 | 353 |
| 8.6 | 284 | 236 | 311 | 321 | 337 |
| 8.7 | 307 | 222 | 329 | 352 | 354 |
| 8.8 | 283 | 236 | 321 | 330 | 368 |
| 8.9 | 271 | 217 | 310 | 328 | 366 |
| 9 | 270 | 225 | 330 | 353 | 368 |
| 9.1 | 294 | 234 | 312 | 382 | 408 |
| 9.2 | 283 | 235 | 313 | 360 | 374 |
| 9.3 | 278 | 227 | 316 | 361 | 366 |
| 9.4 | 264 | 231 | 301 | 350 | 344 |
| 9.5 | 306 | 263 | 334 | 339 | 385 |
| 9.6 | 273 | 239 | 315 | 342 | 360 |
| 9.7 | 288 | 234 | 318 | 362 | 363 |
| 9.8 | 266 | 239 | 295 | 344 | 388 |
| 9.9 | 290 | 228 | 312 | 350 | 417 |
| 10 | 284 | 247 | 319 | 341 | 388 |
| 10.1 | 304 | 243 | 330 | 360 | 379 |
| 10.2 | 275 | 213 | 320 | 352 | 379 |
| 10.3 | 315 | 240 | 319 | 341 | 382 |
| 10.4 | 295 | 226 | 322 | 363 | 370 |
| 10.5 | 278 | 244 | 337 | 370 | 398 |
| 10.6 | 280 | 238 | 325 | 366 | 382 |
| 10.7 | 306 | 221 | 326 | 368 | 384 |
| 10.8 | 310 | 237 | 295 | 346 | 397 |
| 10.9 | 299 | 228 | 305 | 380 | 384 |
| 11 | 315 | 244 | 308 | 392 | 420 |
| 11.1 | 306 | 244 | 296 | 416 | 407 |
| 11.2 | 298 | 231 | 335 | 381 | 409 |
| 11.3 | 297 | 249 | 316 | 394 | 405 |
| 11.4 | 277 | 256 | 321 | 398 | 397 |
| 11.5 | 293 | 260 | 310 | 383 | 423 |
| 11.6 | 275 | 258 | 298 | 378 | 410 |
| 11.7 | 278 | 248 | 315 | 400 | 424 |
| 11.8 | 285 | 242 | 306 | 397 | 401 |
| 11.9 | 289 | 234 | 324 | 386 | 428 |
| 12 | 279 | 236 | 310 | 393 | 419 |
| 12.1 | 293 | 254 | 287 | 391 | 408 |
| 12.2 | 280 | 252 | 298 | 378 | 407 |
| 12.3 | 280 | 238 | 335 | 381 | 428 |
| 12.4 | 257 | 255 | 318 | 398 | 404 |
| 12.5 | 278 | 247 | 317 | 377 | 403 |
| 12.6 | 265 | 252 | 321 | 385 | 397 |
| 12.7 | 283 | 271 | 308 | 382 | 400 |
| 12.8 | 266 | 266 | 322 | 365 | 415 |
| 12.9 | 282 | 272 | 316 | 386 | 410 |
| 13 | 286 | 273 | 331 | 390 | 427 |
| 13.1 | 279 | 262 | 301 | 354 | 405 |
| 13.2 | 287 | 284 | 314 | 367 | 414 |
| 13.3 | 314 | 301 | 301 | 363 | 388 |
| 13.4 | 291 | 284 | 327 | 360 | 406 |
| 13.5 | 288 | 267 | 299 | 378 | 408 |
| 13.6 | 281 | 278 | 307 | 390 | 440 |
| 13.7 | 283 | 266 | 317 | 369 | 412 |
| 13.8 | 287 | 279 | 309 | 395 | 406 |
| 13.9 | 275 | 289 | 317 | 409 | 411 |
| 14 | 289 | 269 | 332 | 410 | 397 |
| 14.1 | 275 | 282 | 318 | 405 | 421 |
| 14.2 | 284 | 275 | 322 | 400 | 432 |
| 14.3 | 286 | 277 | 313 | 394 | 420 |
| 14.4 | 280 | 281 | 306 | 423 | 409 |
| 14.5 | 280 | 301 | 316 | 407 | 446 |
| 14.6 | 294 | 274 | 313 | 404 | 438 |
| 14.7 | 285 | 281 | 319 | 399 | 425 |
| 14.8 | 288 | 280 | 344 | 413 | 419 |
| 14.9 | 287 | 282 | 330 | 406 | 423 |
| 15 | 318 | 303 | 312 | 396 | 415 |
| 15.1 | 270 | 271 | 326 | 403 | 398 |
| 15.2 | 297 | 289 | 322 | 423 | 413 |
| 15.3 | 282 | 238 | 323 | 434 | 422 |
| 15.4 | 286 | 261 | 361 | 405 | 414 |
| 15.5 | 292 | 269 | 337 | 385 | 398 |
| 15.6 | 279 | 243 | 327 | 377 | 420 |
| 15.7 | 284 | 247 | 314 | 402 | 415 |
| 15.8 | 299 | 240 | 326 | 387 | 439 |
| 15.9 | 290 | 270 | 310 | 392 | 422 |
| 16 | 272 | 259 | 319 | 404 | 412 |
| 16.1 | 283 | 252 | 328 | 399 | 398 |
| 16.2 | 289 | 240 | 288 | 403 | 462 |
| 16.3 | 292 | 265 | 273 | 421 | 430 |
| 16.4 | 266 | 258 | 279 | 410 | 420 |
| 16.5 | 275 | 255 | 258 | 409 | 424 |
| 16.6 | 295 | 253 | 272 | 390 | 412 |
| 16.7 | 312 | 239 | 272 | 395 | 414 |
| 16.8 | 275 | 275 | 279 | 404 | 408 |
| 16.9 | 290 | 285 | 277 | 401 | 447 |
| 17 | 291 | 291 | 263 | 379 | 432 |
| 17.1 | 305 | 264 | 281 | 417 | 408 |
| 17.2 | 271 | 281 | 250 | 412 | 446 |
| 17.3 | 295 | 271 | 285 | 433 | 437 |
| 17.4 | 283 | 279 | 270 | 410 | 429 |
| 17.5 | 286 | 279 | 273 | 410 | 427 |
| 17.6 | 279 | 284 | 274 | 408 | 453 |
| 17.7 | 299 | 280 | 264 | 407 | 431 |
| 17.8 | 286 | 257 | 262 | 409 | 446 |
| 17.9 | 290 | 278 | 248 | 409 | 436 |
| 18 | 282 | 273 | 271 | 401 | 437 |
| 18.1 | 290 | 307 | 263 | 395 | 410 |
| 18.2 | 287 | 298 | 263 | 417 | 426 |
| 18.3 | 276 | 291 | 243 | 417 | 410 |
| 18.4 | 293 | 256 | 252 | 430 | 380 |
| 18.5 | 285 | 276 | 272 | 430 | 427 |
| 18.6 | 286 | 281 | 269 | 415 | 433 |
| 18.7 | 286 | 283 | 262 | 436 | 428 |
| 18.8 | 282 | 274 | 246 | 434 | 441 |
| 18.9 | 288 | 251 | 265 | 387 | 441 |
| 19 | 292 | 272 | 251 | 393 | 421 |
| 19.1 | 279 | 295 | 255 | 406 | 411 |
| 19.2 | 298 | 278 | 253 | 382 | 410 |
| 19.3 | 298 | 277 | 249 | 406 | 387 |
| 19.4 | 296 | 278 | 256 | 413 | 389 |
| 19.5 | 277 | 244 | 281 | 418 | 376 |
| 19.6 | 275 | 271 | 258 | 398 | 392 |
| 19.7 | 278 | 278 | 254 | 404 | 389 |
| 19.8 | 303 | 258 | 244 | 413 | 367 |
| 19.9 | 286 | 267 | 260 | 385 | 393 |

**Figure S8.** Timeline plots pertaining to number of contacts between histones and DNA for the systems studied.

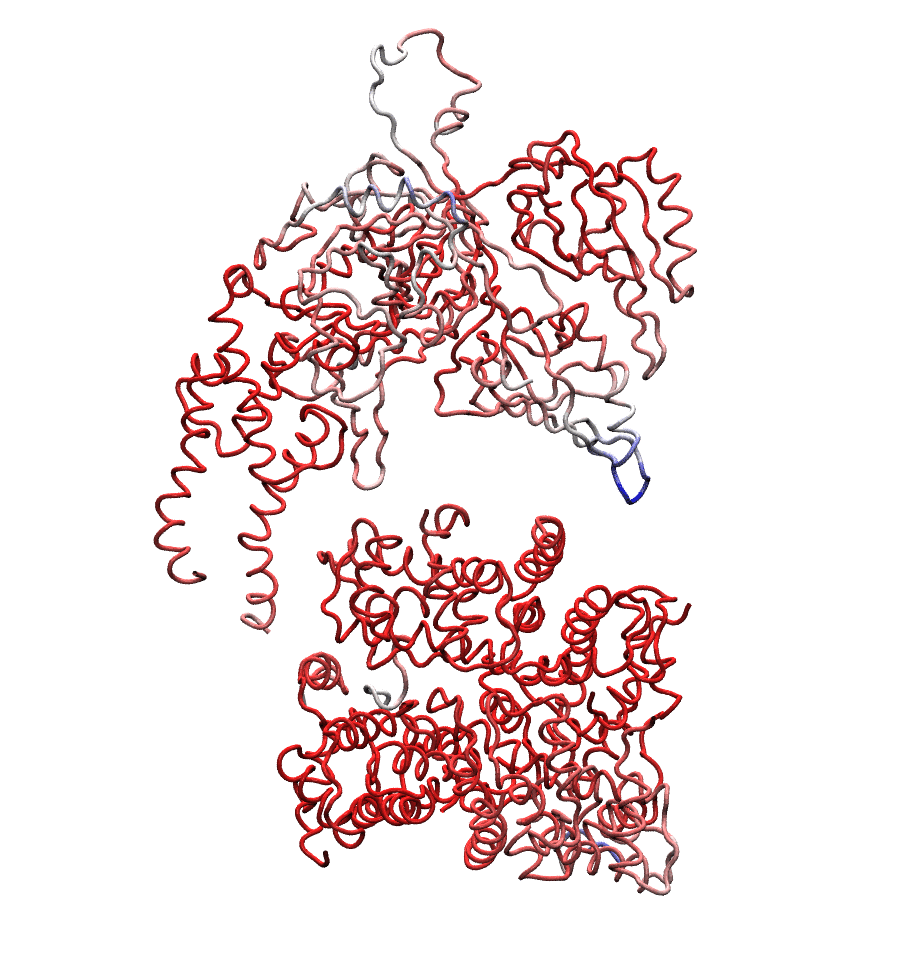

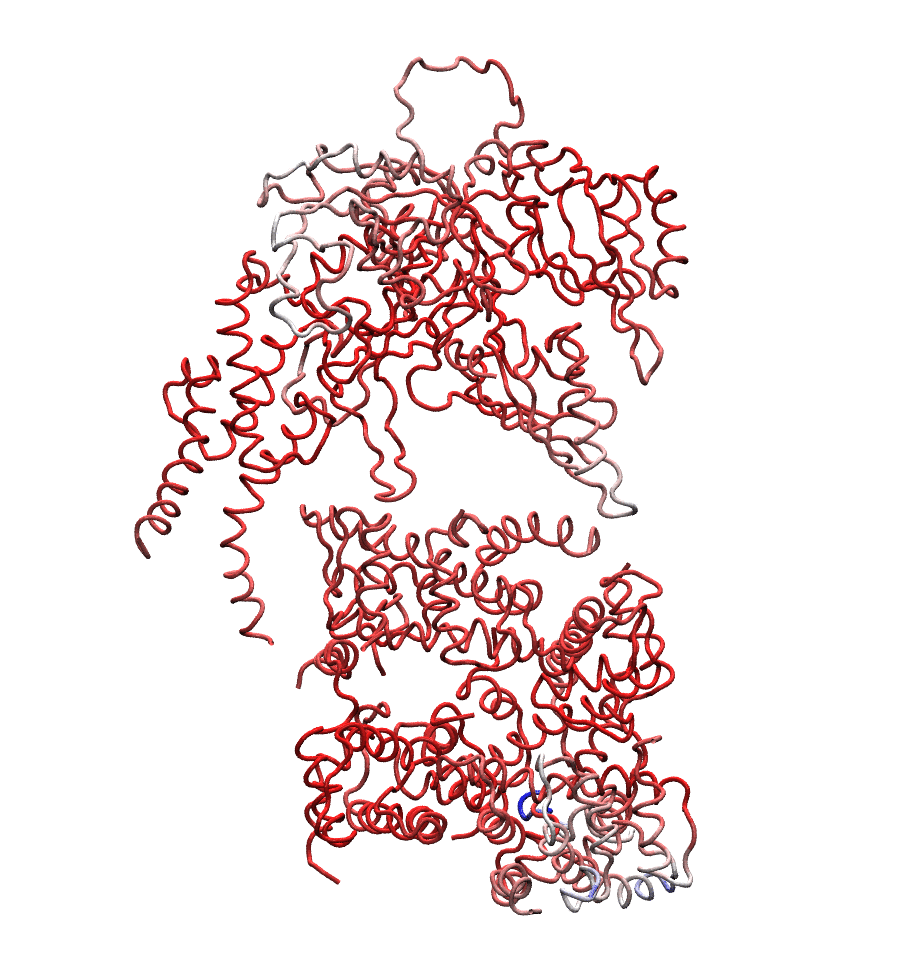

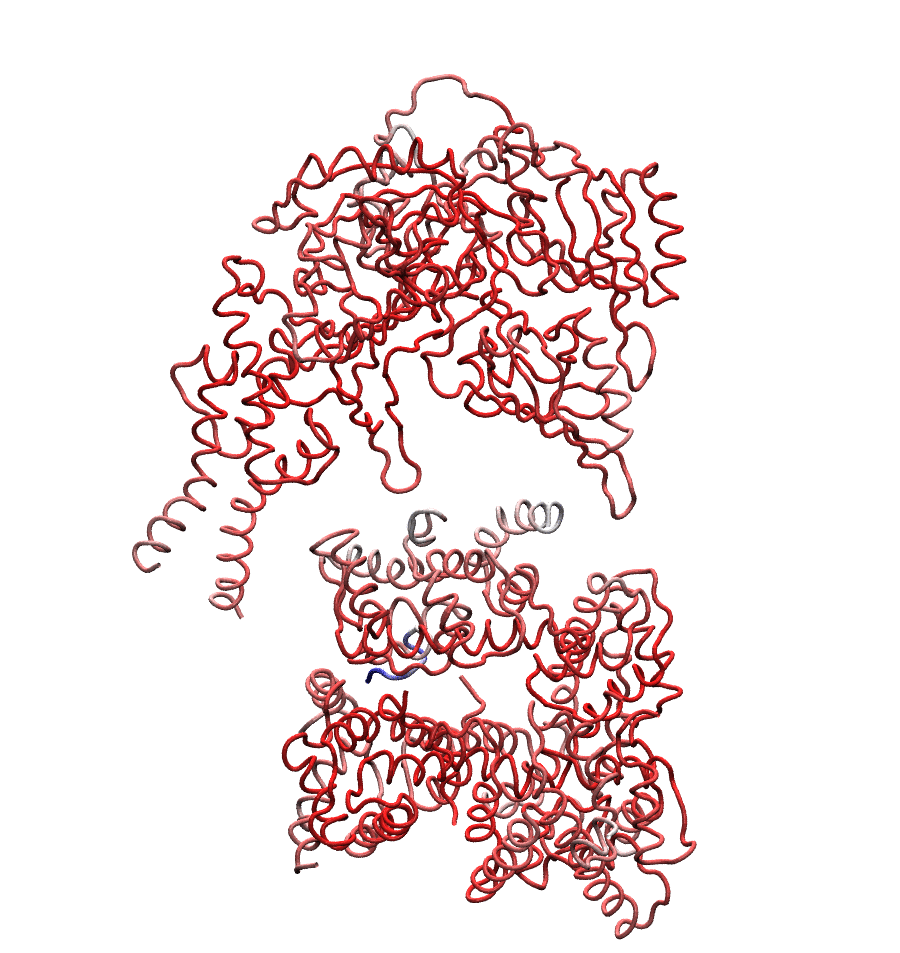

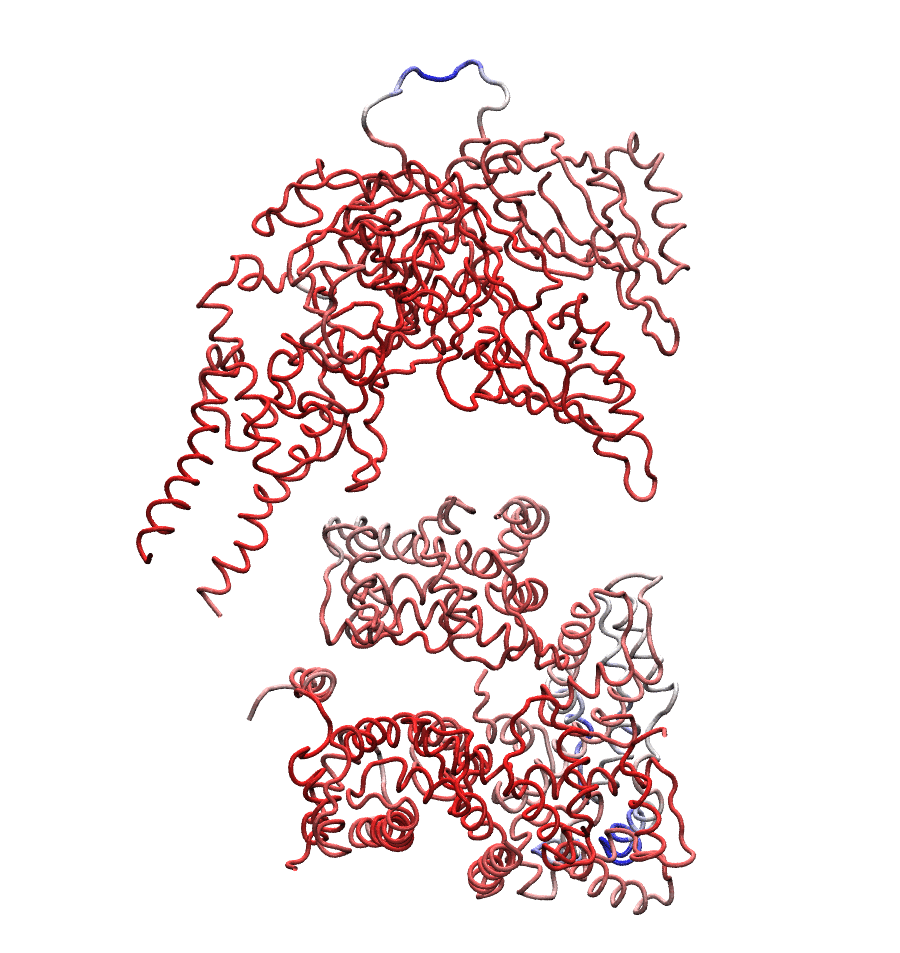

**Figure S9.** Movies illustrating the global motions along the first eigenvector for the systems: nonphosphorylated (upper left), CLOCK^S38/42^ (upper right), BMAL1^S78^ (lower left), and CLOCK^S38/42^/BMAL1^S78^ (lower right).

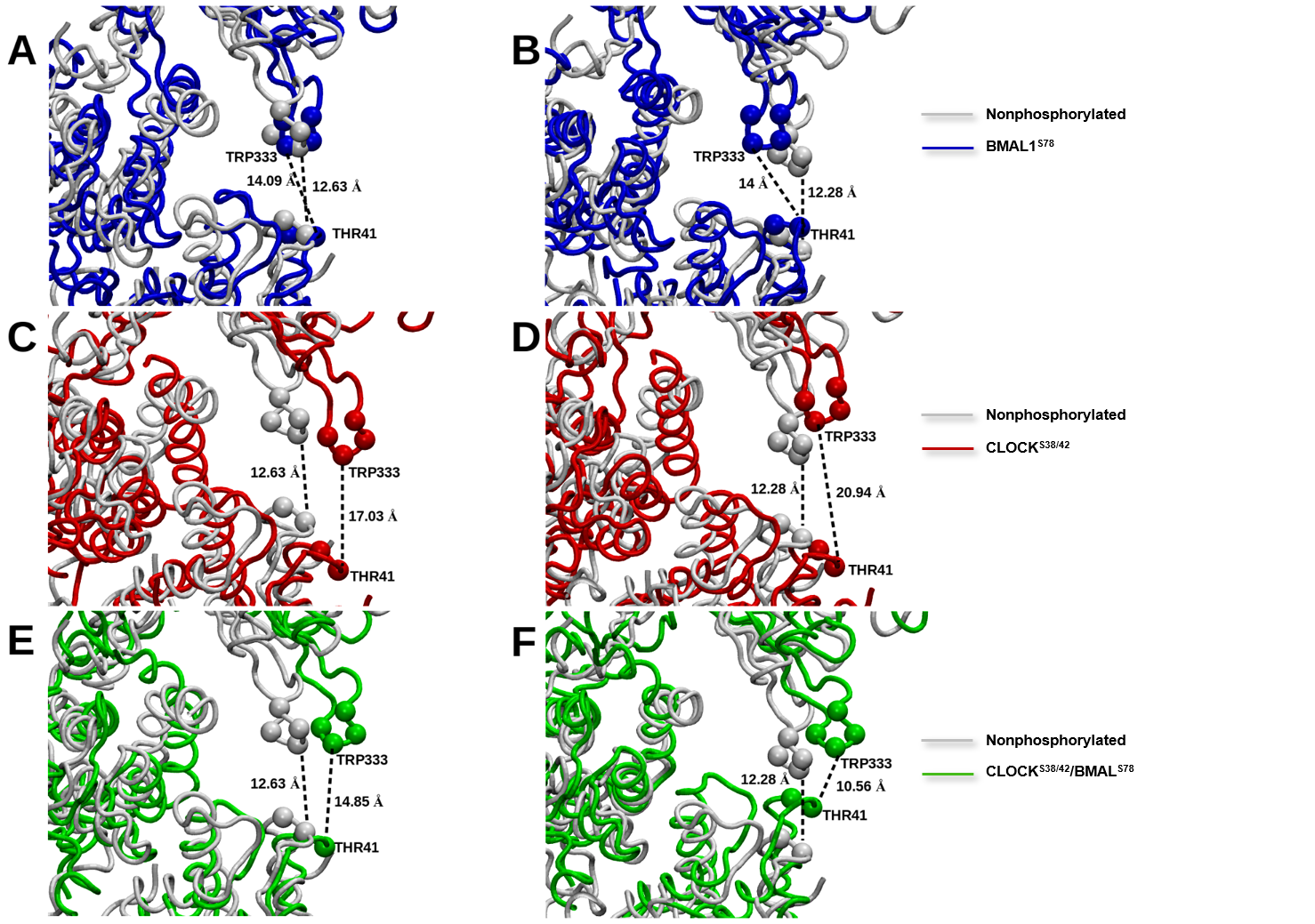

**Figure S10.** Average distance between residues chosen from CLOCK HI loop (TRP333) and H3α1–L1 elbow (THR41) measured from PCA trajectories. The initial and final PCA states of the phosphorylated systems were compared with the nonphosphorylated system: **A.** BMAL1^S78^ initial PCA state **B.** BMAL1^S78^ final PCA state **C.** CLOCK^S38/42^ initial PCA state **D.** CLOCK^S38/42^ final PCA state **E.** CLOCK^S38/42^/BMAL1^S78^ initial PCA state **F.** CLOCK^S38/42^/BMAL1^S78^ final PCA state.

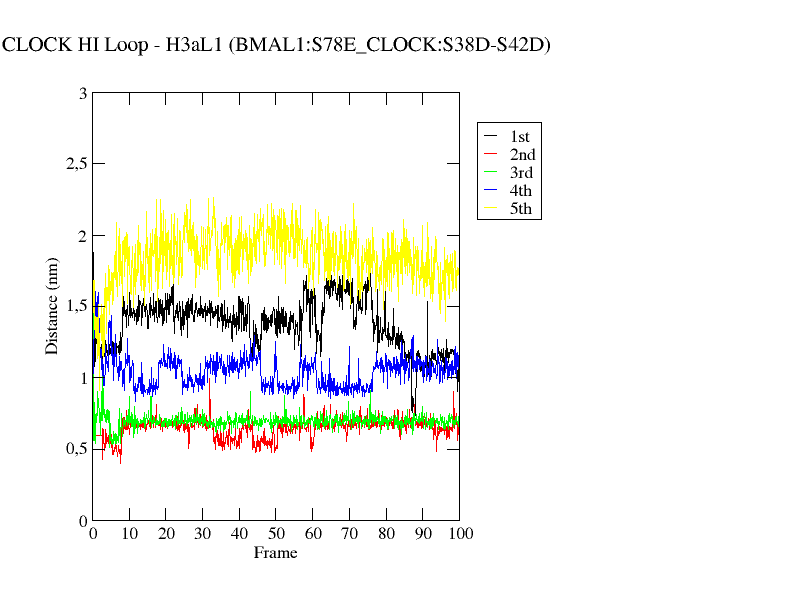

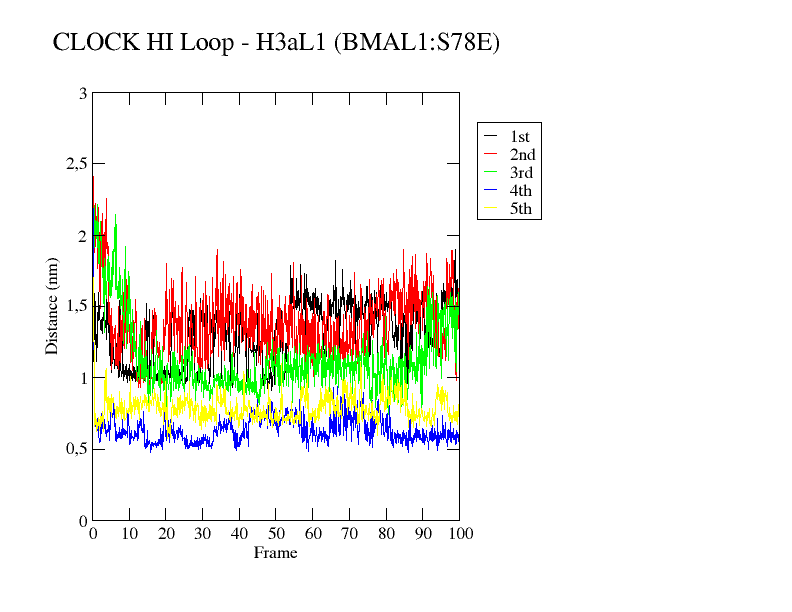

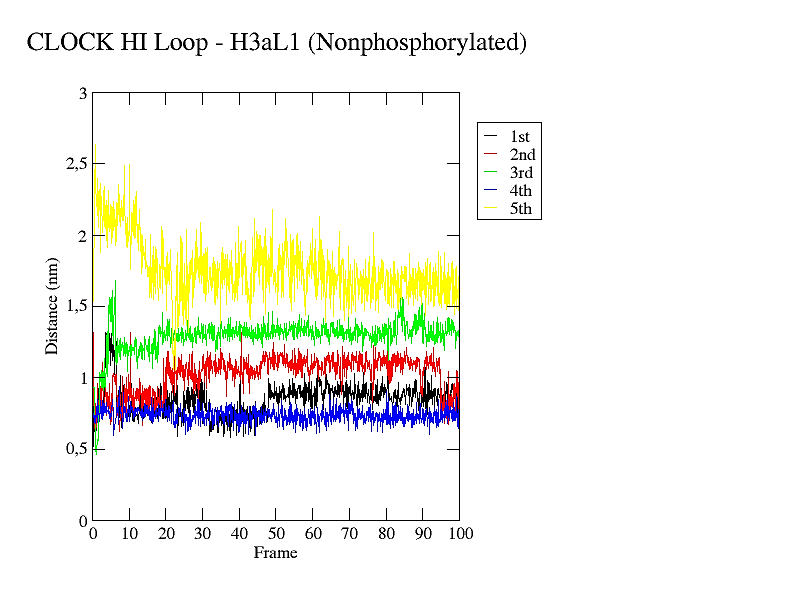

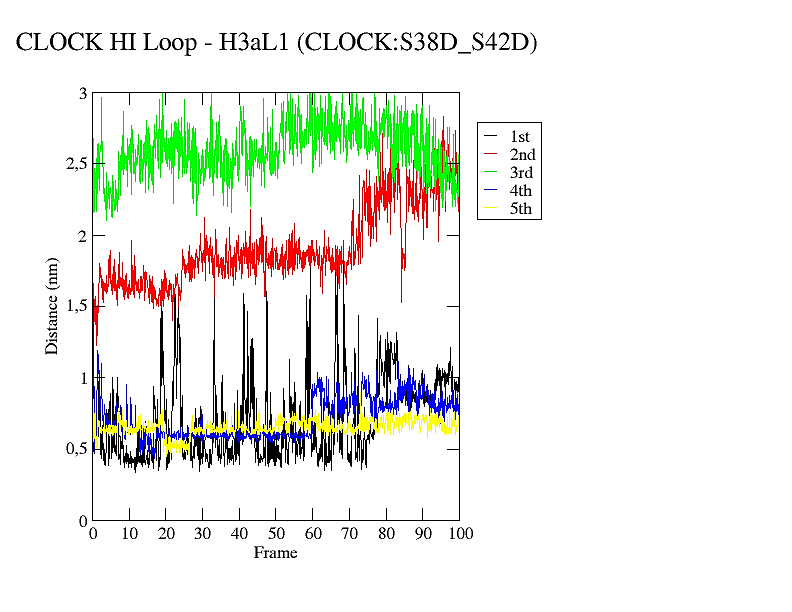

**Figure S11.** Timeline plots pertaining to distances between the center of mass of CLOCK HI loop and the H3α1–L1 elbow for the systems studied.

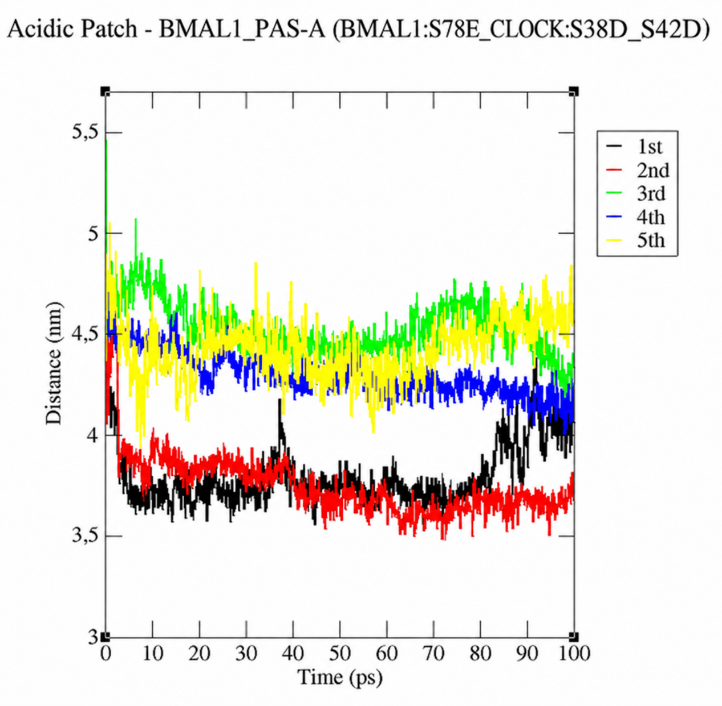

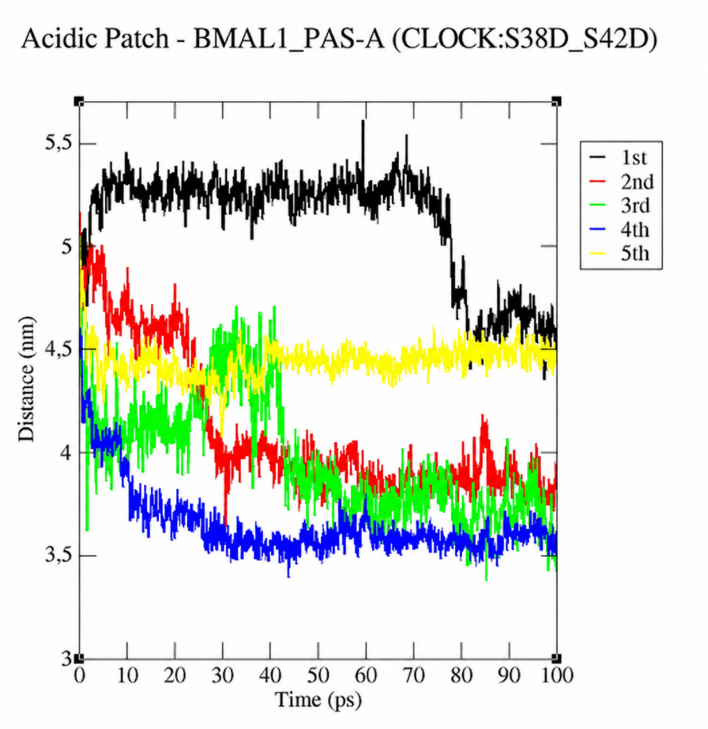

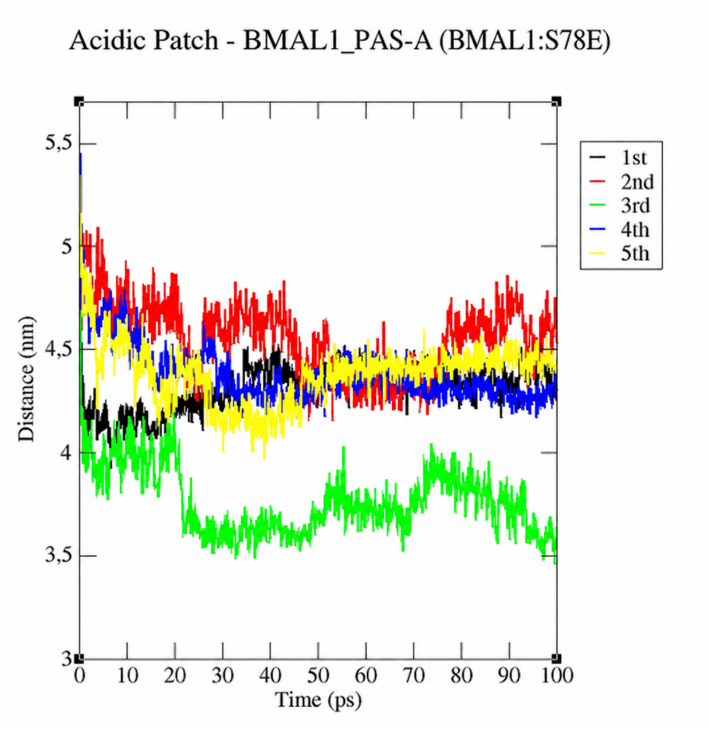

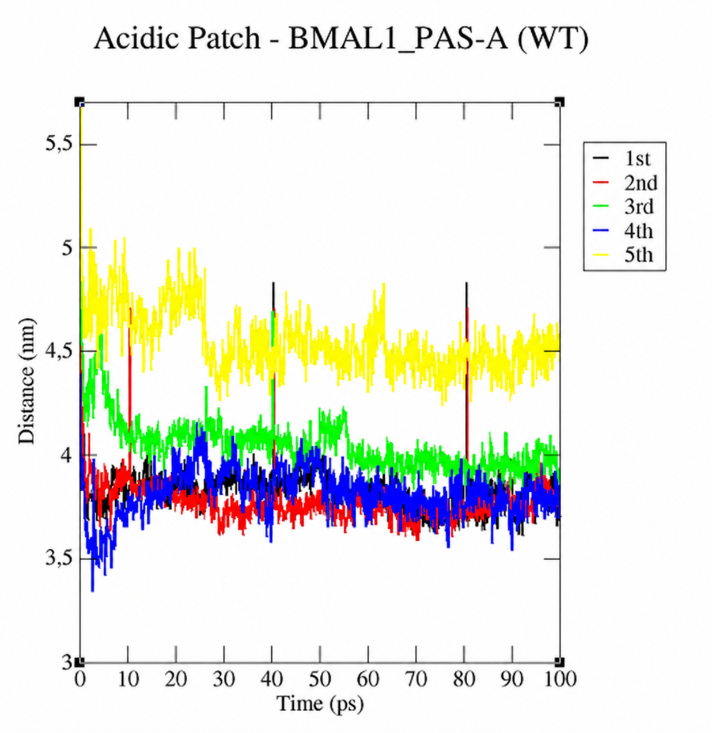

**Figure S12.** Timeline plots pertaining to distances between the center of mass of BMAL1 PAS-A and the acidic patch for the systems studied.

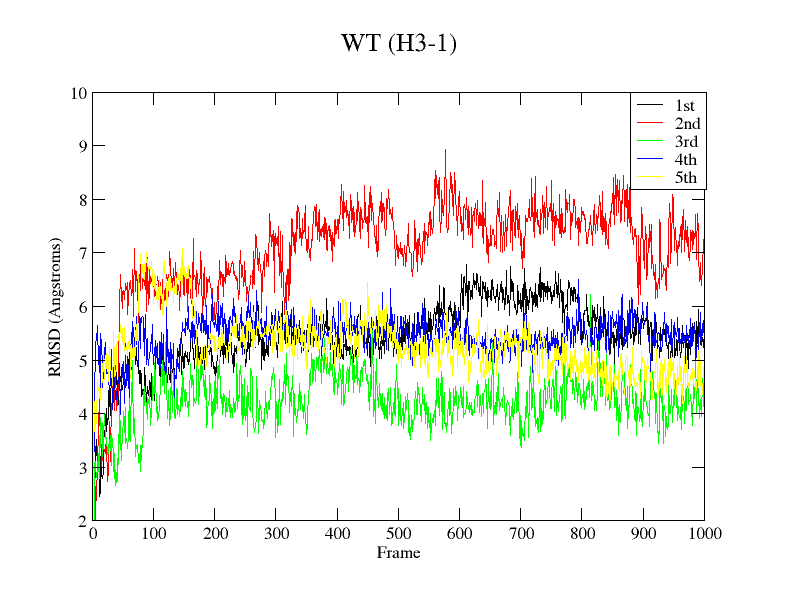

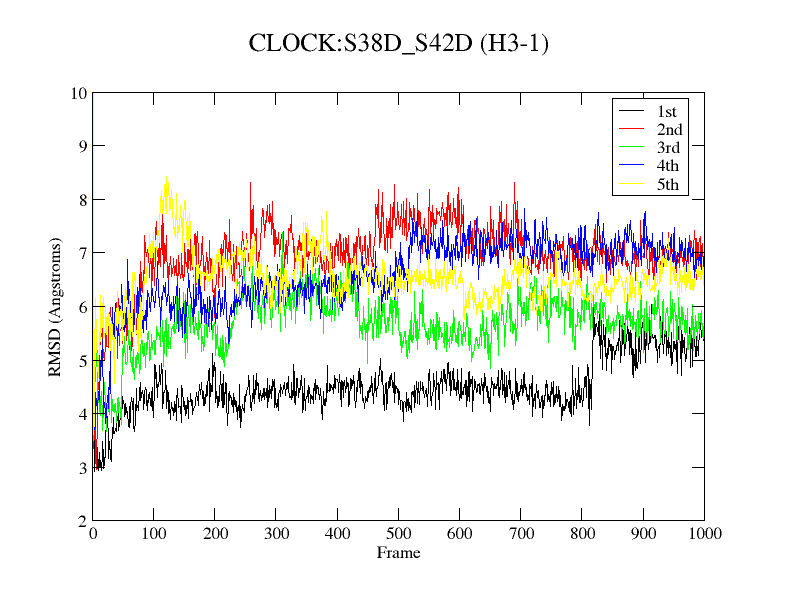

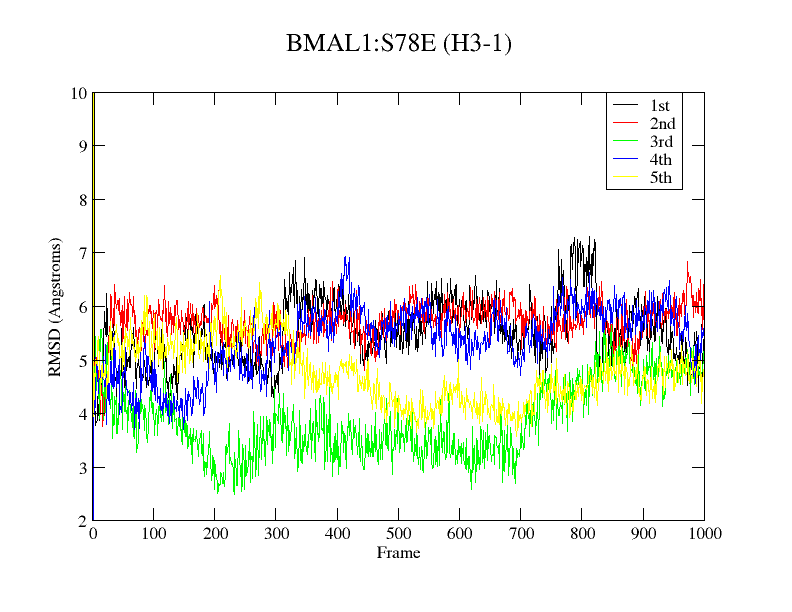

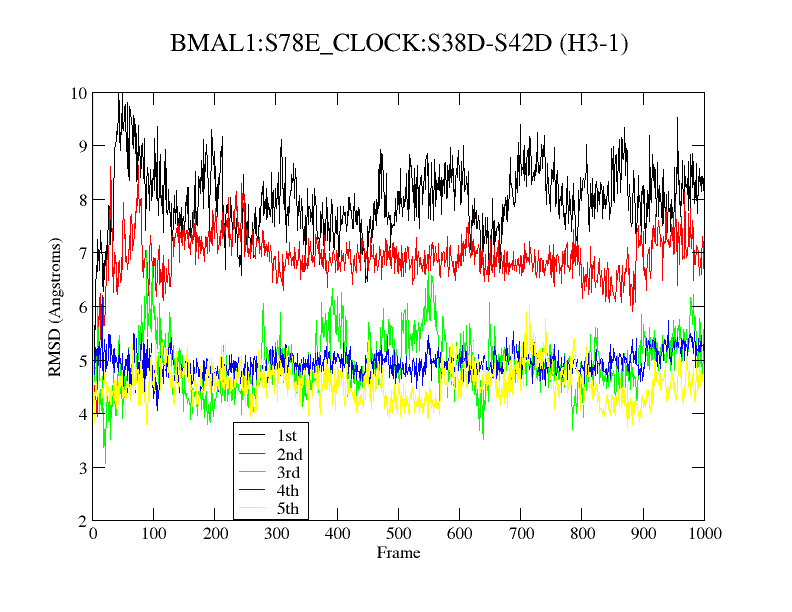

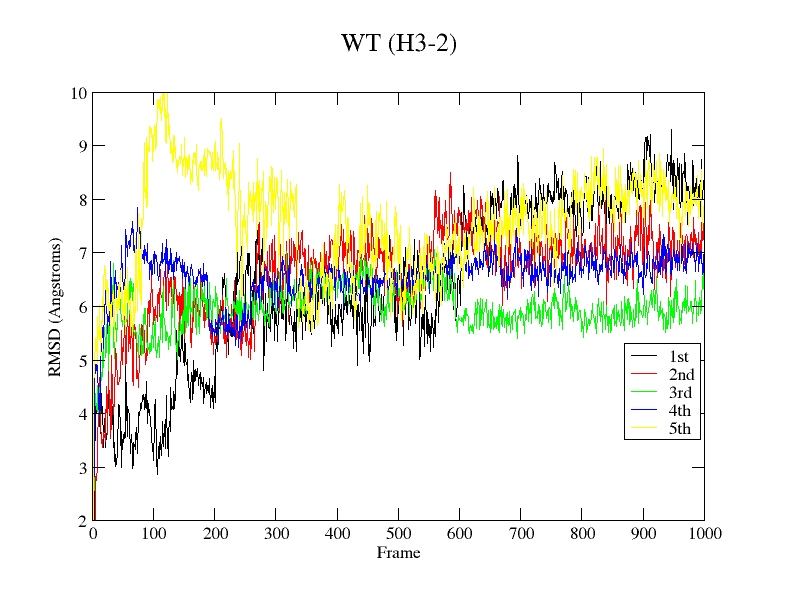

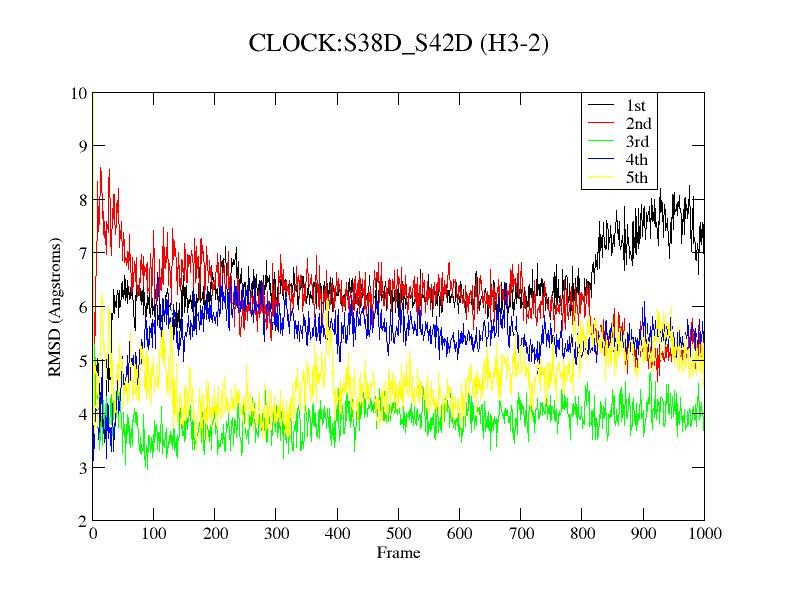

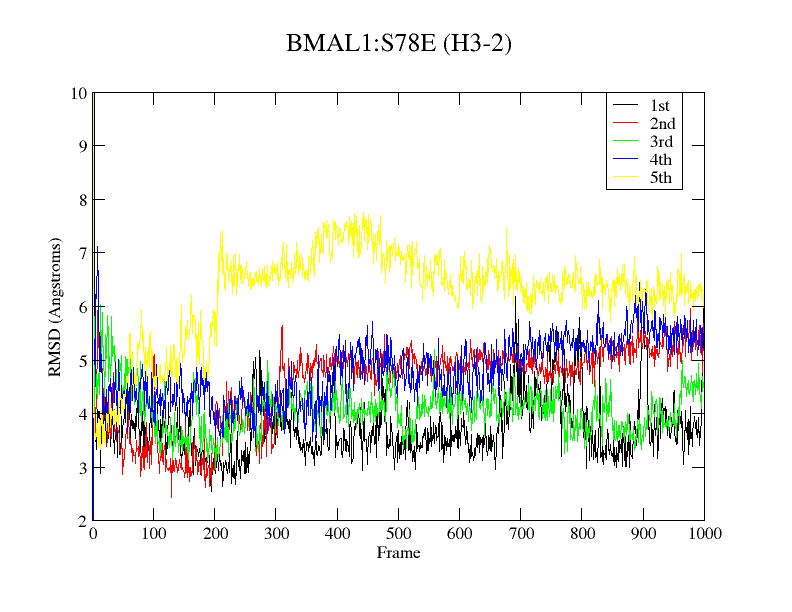

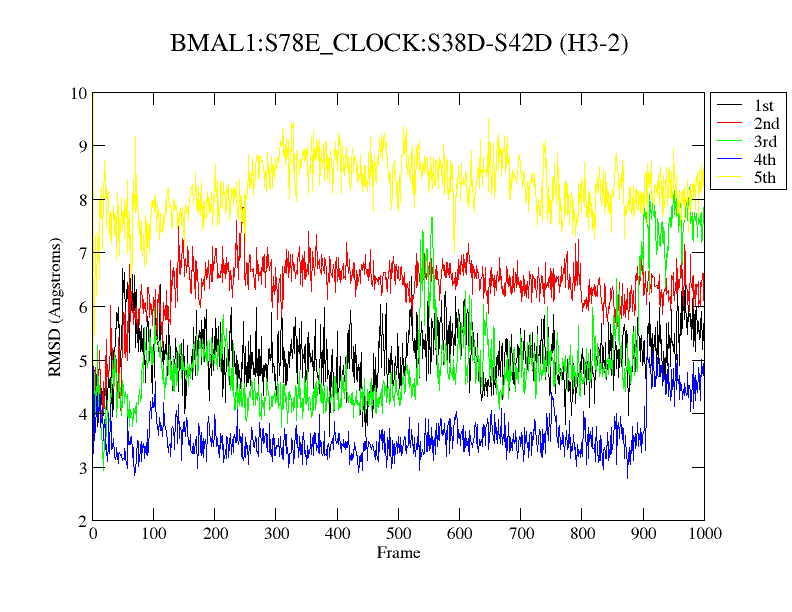

**Figure S13.** RMSD of histones H3-1 and H3-2 for the systems studied.

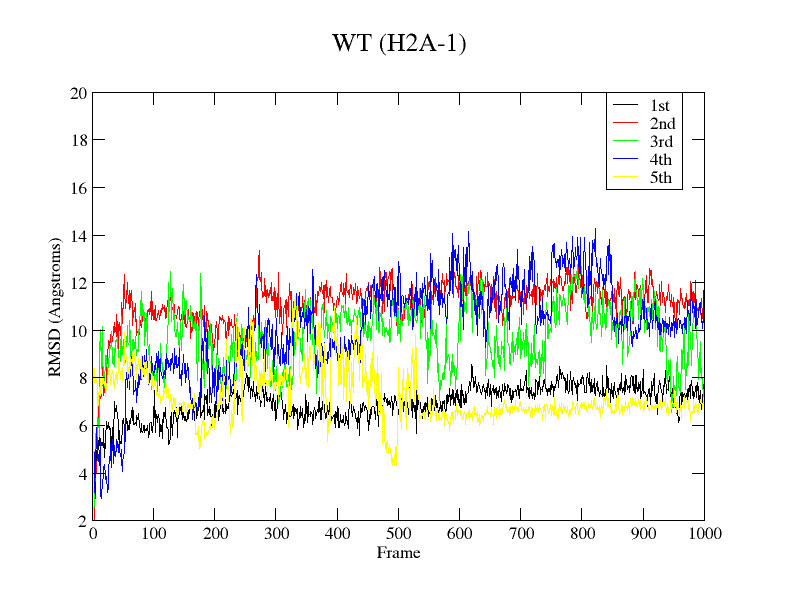

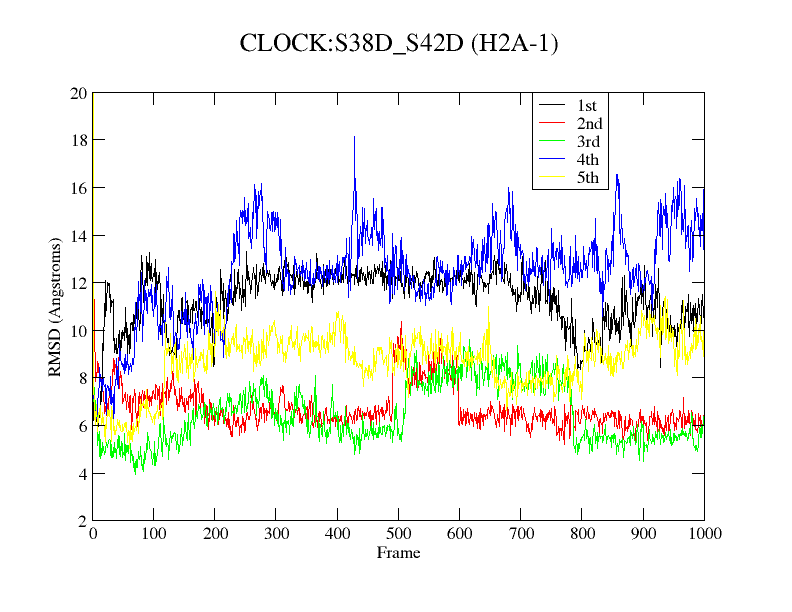

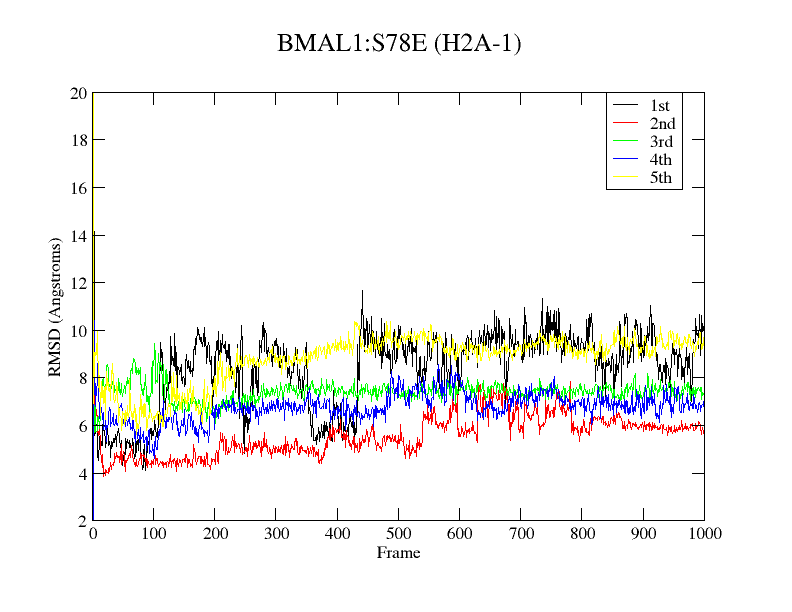

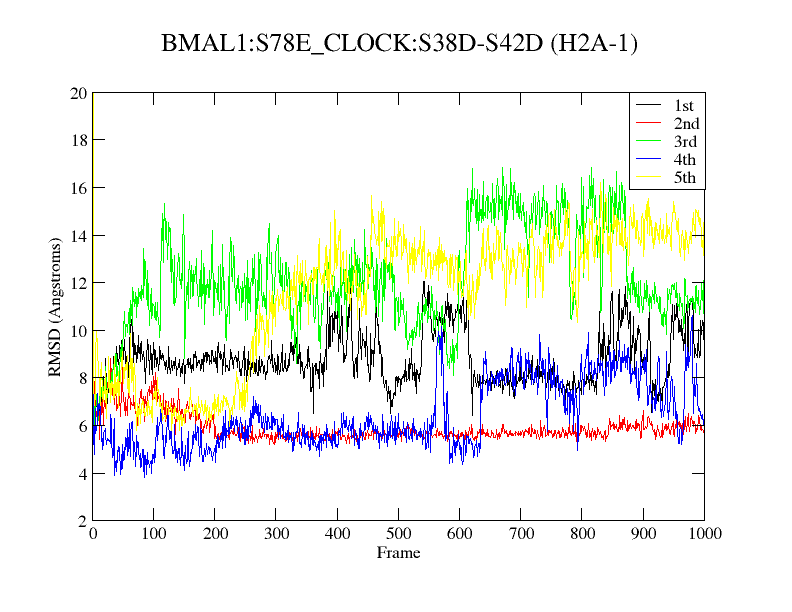

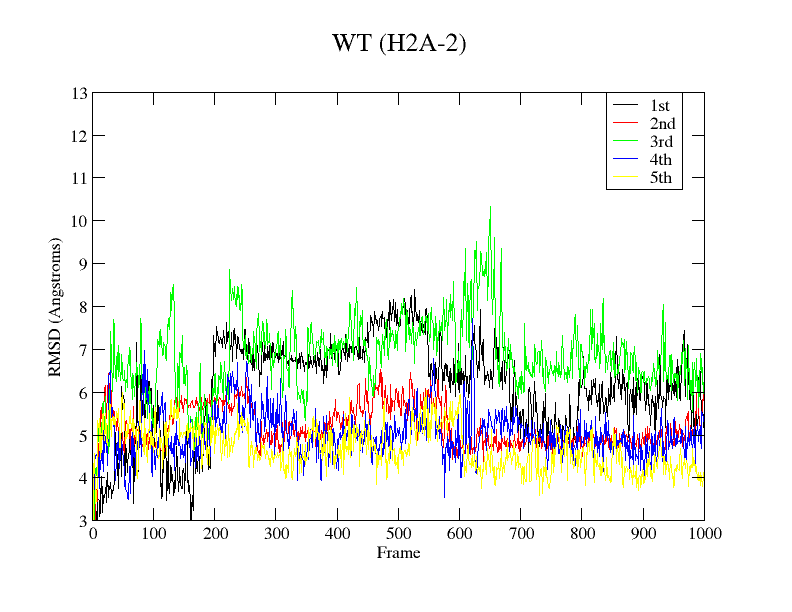

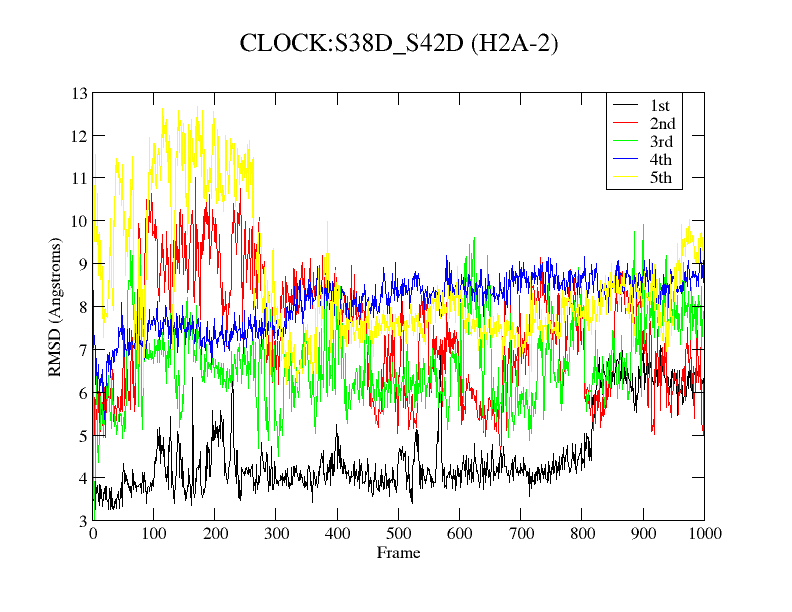

**Figure S14.** RMSD of histones H2A-1 and H2A-2 for the systems studie
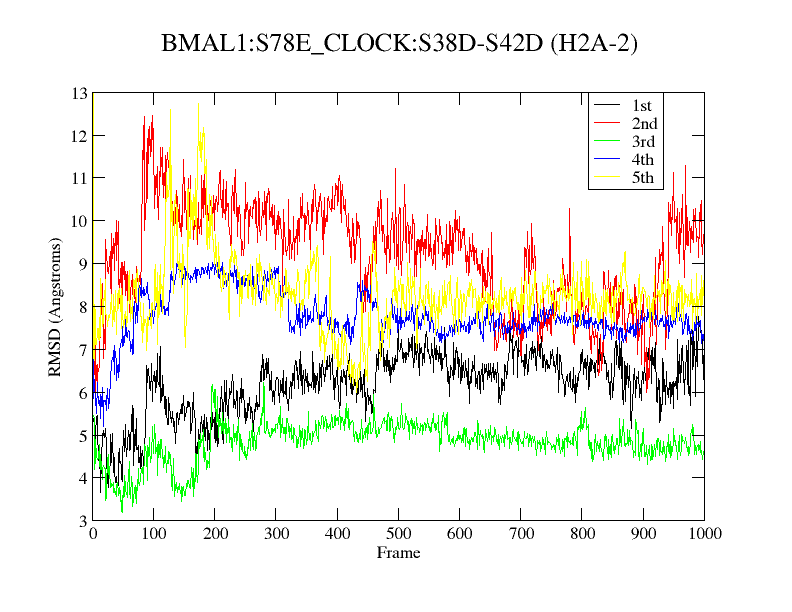

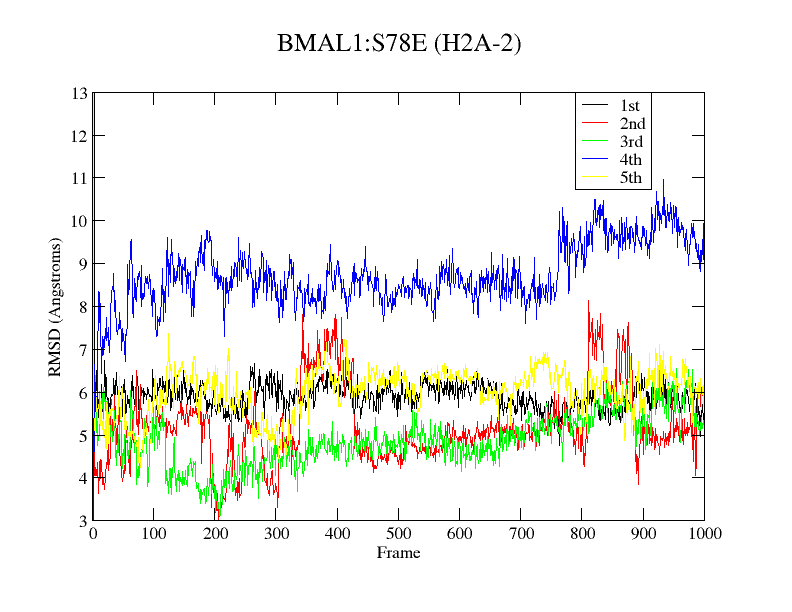
d.

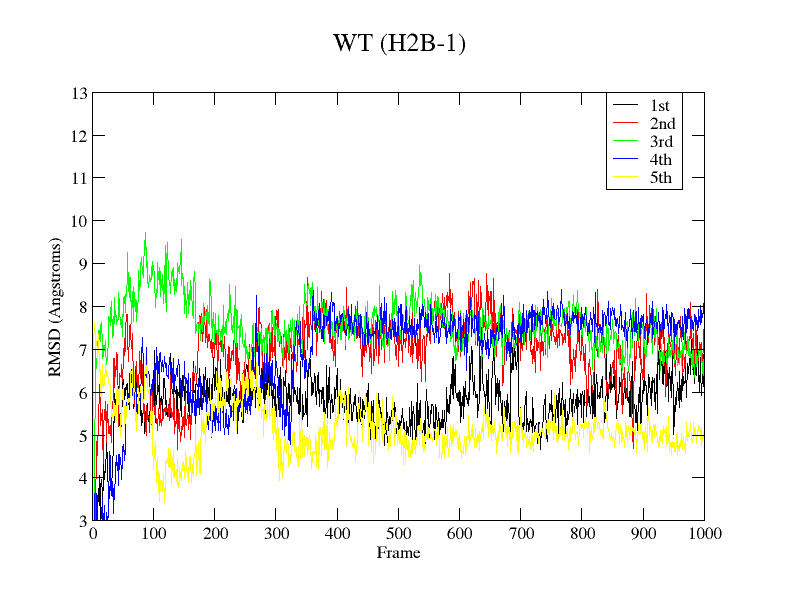

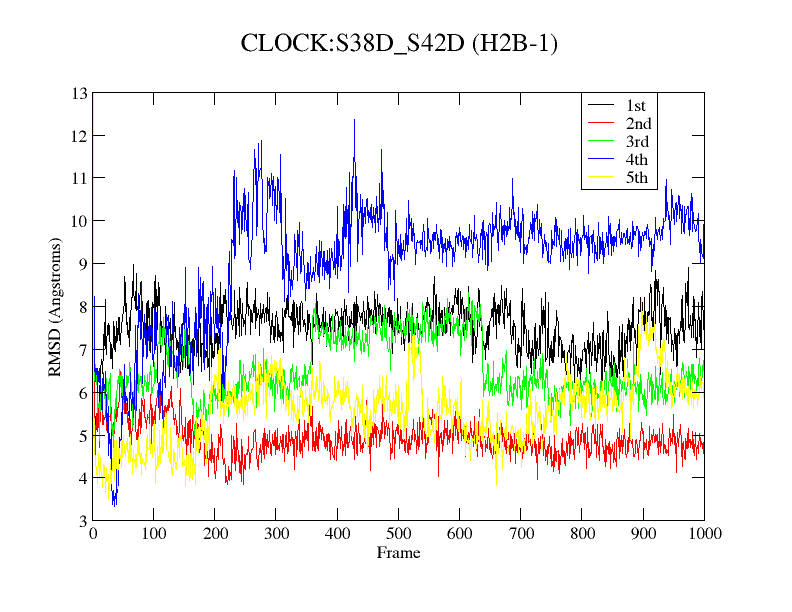

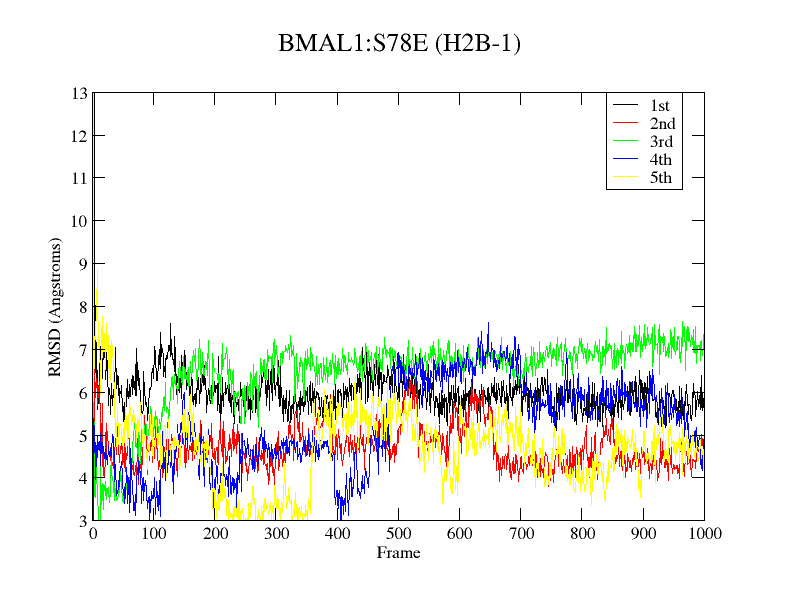

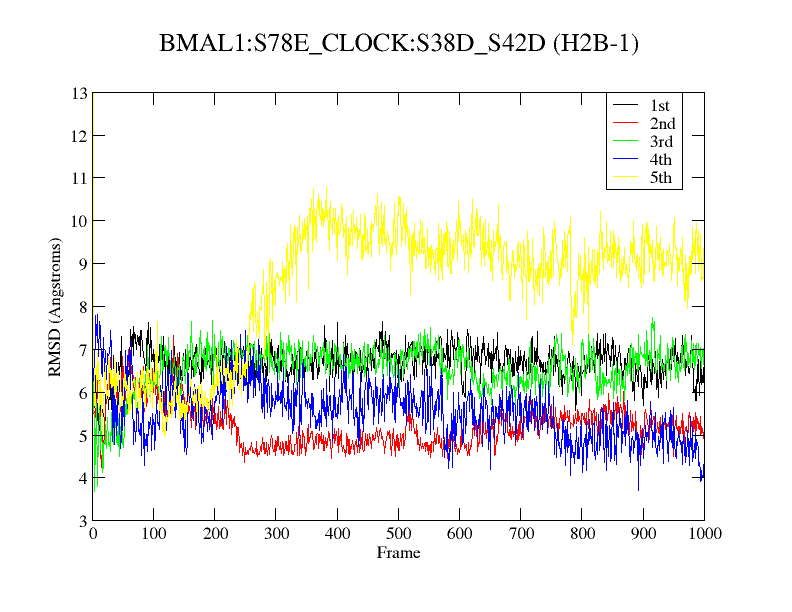

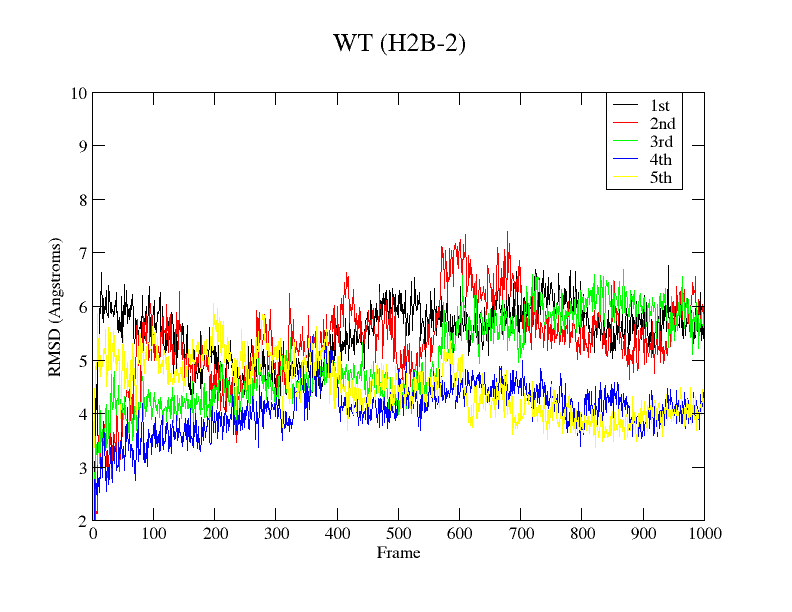

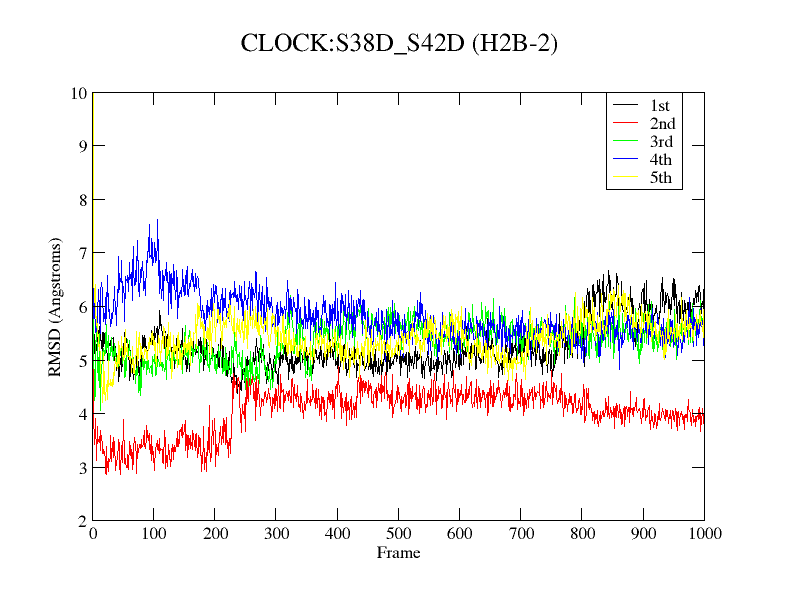

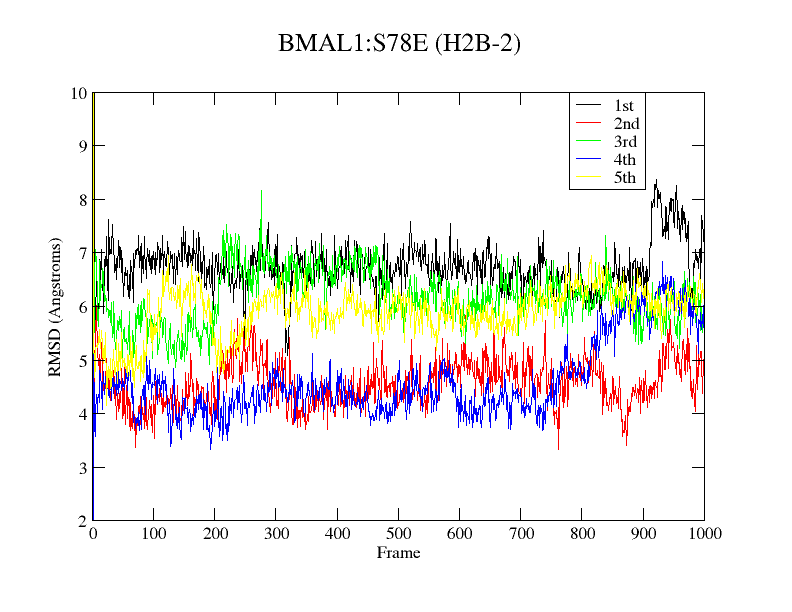

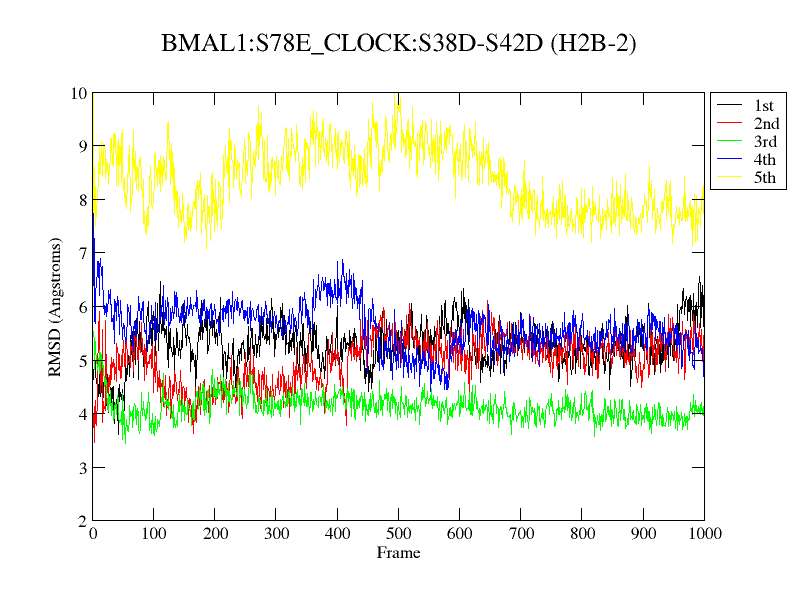

**Figure S15.** RMSD of histones H2B-1 and H2B-2 for the systems studied.
